# Rapid cyclic stretching induces a synthetic, proinflammatory phenotype in cultured human intestinal smooth muscle, with the potential to alter signaling to adjacent bowel cells

**DOI:** 10.1101/2024.10.12.617767

**Authors:** Sharon M. Wolfson, Katherine Beigel, Sierra E. Anderson, Brooke Deal, Molly Weiner, Se-Hwan Lee, Deanne Taylor, Su Chin Heo, Robert O. Heuckeroth, Sohaib K. Hashmi

## Abstract

**Background and Aims:** Bowel smooth muscle experiences mechanical stress constantly during normal function, and pathologic mechanical stressors in disease states. We tested the hypothesis that pathologic mechanical stress could alter transcription to induce smooth muscle phenotypic class switching.

**Methods:** Primary human intestinal smooth muscle cells (HISMCs), seeded on electrospun aligned poly-ε-caprolactone nano-fibrous scaffolds, were subjected to pathologic, high frequency (1 Hz) uniaxial 3% cyclic stretch (loaded) or kept unloaded in culture for 6 hours. Total RNA sequencing, qRT-PCR, and quantitative immunohistochemistry defined loading-induced changes in gene expression. NicheNet predicted how differentially expressed genes might impact HISMCs and other bowel cells.

**Results:** Loading induced differential expression of 4537 genes in HISMCs. Loaded HISMCs had a less contractile phenotype, with increased expression of synthetic SMC genes, proinflammatory cytokines, and altered expression of axon guidance molecules, growth factors and morphogens. Many differentially expressed genes encode secreted ligands that could act cell-autonomously on smooth muscle and on other cells in the bowel wall.

**Discussion:** HISMCs demonstrate remarkably rapid phenotypic plasticity in response to mechanical stress that may convert contractile HISMCs into proliferative, fibroblast-like cells or proinflammatory cells. These mechanical stress-induced changes in HISMC gene expression may be relevant for human bowel disease.

## Introduction

The gastrointestinal (GI) tract is constantly moving to digest and absorb nutrients, and to eliminate waste. This movement creates mechanical stress that can be sensed by cells and may alter gene expression through mechanotransduction pathways (1–5). In the bowel, pathologic radial or longitudinal force occurs from mechanical obstruction (stricture, web, volvulus, adhesions), motility disorders (achalasia, gastroparesis, Hirschsprung disease, chronic intestinal pseudo-obstruction (CIPO)), and surgical manipulation. Similar to physiologic mechanical forces, pathological mechanical forces may also induce transcriptional changes that alter cell phenotypes (2, 5, 6). While many bowel cell types respond to mechanical cues (5), we hypothesized that unusual mechanical stressors might particularly impact gene expression in bowel (visceral) smooth muscle cells (SMCs), altering cell fate. These changes in visceral SMC fate were predicted based on decreased contractile smooth muscle marker expression in pediatric CIPO bowel (7), and by extrapolating from vascular smooth muscle cells, which undergo “phenotypic class switching” to a synthetic, proliferative phenotype in response to injury (7–9). This phenotypic class switching for vascular SMCs is thought to be protective, but is also an important pathophysiologic mechanism in hypertension, ischemic vascular disease, and atherosclerosis.

Phenotypic class switching is not as well studied in visceral SMCs, and many differences between visceral and vascular SMCs may make extrapolation inappropriate. However, prior studies suggest visceral SMCs also change fate in response to specific physiologic mechanical stressors. For example, partial intestinal obstruction increases SMC expression of COX-2 (PTGS2), mPGES-1, and PGE2 (10) *in vivo*, while stretch *in vitro* of primary colon SMCs increases levels of IL-8, IL-6, MCP1, iNOS, COX2, BDNF, and NGF (11). Mechanical stress also induces human fetal visceral SMC expression of pro-fibrotic mediators, including TGFβ1 and α1 collagen (12), markers of synthetic SMCs.

To test the hypothesis that mechanical stress could rapidly alter gene expression in visceral SMCs, and to gain insight into early changes in SMC phenotype in response to mechanical stress, we evaluated gene expression in cultured human intestinal smooth muscle cells (HISMCs) after only 6 hours in culture with or without cyclic stretching. We used a low amplitude, high frequency mechanical stress (3%, 1 Hz), a frequency up to an order of magnitude greater than physiologic bowel contraction. This pathologic stress rapidly altered expression of 4537 genes (adjusted p-value < 0.05 for loaded versus unloaded cells). Of these genes, 2500 had log2 fold change > 0.48 or < -0.48. Compared to unloaded HISMCs, loaded cells had increased expression of genes typically produced in synthetic phenotype SMCs, increased production of many cytokines, chemokines, cytokine receptors, axon guidance molecules, junctional proteins, and altered levels of many signaling molecules predicted to act on nearby cells in the bowel. Collectively, these data suggest that bowel SMC phenotype, in part, depends on the unique physical forces these cells experience as nutrients move through the bowel, waste is eliminated, and in response to bowel injury or disease. This suggests that even a brief pathologic mechanical insult may profoundly affect visceral smooth muscle phenotype. Furthermore, our NicheNet analyses suggest mechanotransduction-induced phenotypic changes in smooth muscle gene expression may lead to secretion of ligands that subsequently alter function of most bowel cells on a timescale much longer than the duration of the original insult.

## Methods

### Sex as a biological variable

Our study used human HISMCs that express *XIST* (gene count 1536 to 5403 based on our RNAseq data), indicating they were derived from a human female.

### Preparation of Nanofibrous Scaffolds

Aligned and non-aligned poly(ε-caprolactone) (PCL) nanofibrous scaffolds (Mol. Wt. 80kDa, Shenzhen Bright China Industrial Co., Ltd., China) were fabricated via electrospinning, as described (13). Scaffolds were hydrated and sterilized in ethanol diluted in distilled water (100%, 70%, 50%, 30%; 30 min/step), and then incubated in a laminin (20 μg/mL) (14, 15) solution in 1x Phosphate Buffered Saline (PBS) (Invitrogen, Catalog# 14190136) overnight at 37°C to enhance cell attachment.

### HISMC Preparation and expansion

5x10^5^ smooth muscle cells from human small intestine (cryopreserved at passage 1, ScienCell Research Laboratories, Catalog# 2910) were plated on 10 cm tissue culture dishes coated with 0.1% gelatin (MilliporeSigma, Catalog# G1890) and cultured (37°C, humidified incubator, 5% CO_2_) in HISMC culture media (SMC medium (Sciencell, Catalog# 1101), 2% FBS (fetal bovine serum, Sciencell, #0010), 1% Penicillin/Streptomycin (Sciencell, Catalog# 0503), 1% smooth muscle cell growth supplement (Sciencell, Catalog# 1152)). HISMC culture media was changed every other day. Cells were passaged at 90% confluence. After 1-2 passages, confluent HISMCs were cryopreserved in 90% FBS/10% dimethyl sulfoxide (DMSO) at 5x10^5^ cells/mL. All experiments then used HISMCs at passage 3-5.

### Dynamic mechanical loading of HISMC-seeded scaffolds

Frozen HISMCs were thawed in a 37°C water bath for 2-3 minutes and then added to 10 mL of Iscove’s Modification of DMEM (Corning, Catalog# 10-016-CM), and pelleted (270 x g, 3 minutes). Aligned laminin-coated PCL scaffolds (30 mm x 5 mm) were seeded with 350,000 HISMCs resuspended in 80 μL HISMC culture media. Cell-seeded scaffolds were maintained free-floating in HISMC culture media for 72 hours. “Loaded” scaffolds then experienced cyclic stretch (3% uniaxial stretch, 1 Hz, parallel to the long axis of HISMCs) for 6 hours in fresh HISMC culture media using a custom bioreactor (16). In parallel, “unloaded” HISMCs were maintained free-floating on scaffolds in fresh HISMC culture media for 6 hours. All cells were maintained at 37°C, 5% CO_2_ in a humidified incubator. Scaffolds were then cut in half. One half was dissolved in Trizol (Ambion, Catalog# 15596018) for RNA extraction and the other half fixed for immunohistochemistry (see below).

### RNA extraction and purification

To isolate RNA from HISMCs on PCL scaffolds, each scaffold was minced in 500 μL TRizol (Ambion, Catalog# 15596018) using sharp scissors and then vortexed for up to 15 minutes until the scaffold dissolved. RNA was purified from cells lysed in TRIzol using the RNeasy Plus Mini kit (QIAGEN, Catalog# 74134), with RNase Free DNase Set (QIAGEN, Catalog# 79254) to remove residual DNA. RNA concentrations were measured by NanoDrop (ND-2000, Thermo Fisher Scientific).

### Quantitative real-time PCR

Quantitative real-time PCR (qRT-PCR) was performed using SsoFast Evagreen Supermix with Low ROX (Bio-Rad, Catalog# 172-684 5211) and primers in Supplemental Table 1. Cycle threshold (C_t_) values were normalized to *YWHAZ* mRNA.

### Immunofluorescent staining

Scaffolds were washed once with 1x PBS, fixed (4% paraformaldehyde, 30 minutes, room temperature), washed twice with 1x PBS (5 minutes each, room temperature), blocked (5% normal donkey serum [NDS], 0.5% Triton X-100 in PBS (0.5% PBST), 1 hour, room temperature), incubated in primary antibodies (5% NDS, 0.5% PBST 1 hour, room temperature) (Supplemental Table 2), washed 3 times for 5 minutes (0.5% PBST), and then incubated in secondary antibodies (Supplemental Table 2) (0.5% PBST, 30 minutes, dark, room temperature). Phalloidin staining was performed after secondary antibody staining by washing 3 times for 5 minutes (PBS) and incubating (1 hour, dark, room temperature) in Alexa Fluor–conjugated phalloidin (488 nm, 555 nm, or 647 nm; Invitrogen Catalog# A12379, A34055, and A2287) diluted 1:1000 in PBS. Cells were washed 2 times in 1x PBS, incubated in 1:30,000 SYTOX^TM^ green (Thermo Fisher Scientific, Catalog# S7020) diluted in Hanks Balanced Salt Solution (30 minutes, dark, room temperature), washed twice in 1x PBS, mounted in Prolong^TM^ Diamond AntiFade Mountant (Thermo Fisher Scientific, Catalog#P36961), and allowed to set (overnight, dark, room temperature) before long-term storage in PBS at 4°C.

### Immunofluorescence microscopy

Scaffolds were imaged using a Zeiss LSM 710 (Zeiss ZEN 2.3 SP1 FP3 (Black; (version 14.0.18.201; data in Figure 1) or LSM 980 (Zeiss Zen Blue 3.5 software; data in all other figures) laser scanning confocal microscopes. Images were acquired with a x20/0.8 air or x63/1.4 oil DIC M27 Plan-Apochromat objective. Confocal images were processed in ImageJ (NIH) to crop, scale, and uniformly color adjust. Confocal images are represented as “sum of slices” or “maximum intensity” projections after ImageJ processing.

**Figure 1:**
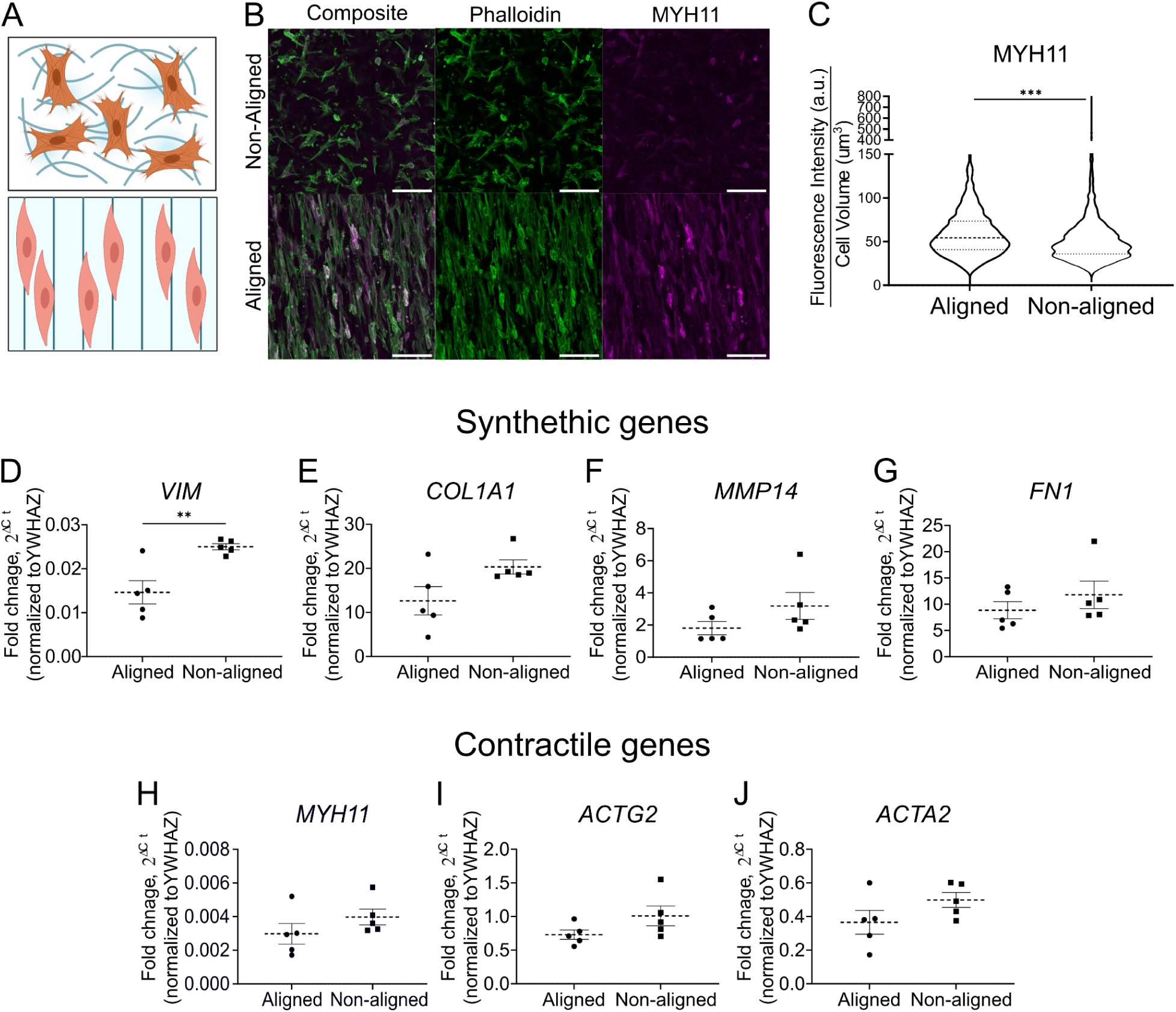
Aligned nanofibrous spun scaffolds enhanced abundance of contractile phenotype mRNA and proteins in HISMC. (A) Schematic of non-aligned (top) and aligned (bottom) PCL scaffolds (Created with Biorender).(B) Confocal Z-stack maximum intensity projections of HISMCs stained with antibodies to smooth muscle myosin (MYH11, magenta) and for F-actin (Phalloidin-Alexa Fluor 488, green) after culture on non-aligned (top) or aligned (bottom) scaffolds immersed in culture media for 72 hours. (scale bar: 100 μm). (C) MYH11 antibody staining was brighter in HISMCs cultured 72 hours on aligned scaffolds compared to HISMC cultured on non-aligned scaffolds (median [interquartile range] Aligned: 54.33 AU [32.6 AU], non-aligned: 47.6 AU [28.28 AU], Mann-Whitney test, P<0.0001, n=3). (D-J) qRT-PCR analyses for mRNA levels of smooth muscle synthetic genes (*VIM, COL1A1, MMP14, FN1*) showed increased VIM expression in HISMCs grown on non-aligned scaffolds (D), with similar expression of the other synthetic genes. qRT-PCR analyses demonstrated that mRNA levels of smooth muscle contractile genes (*MYH11, ACTG2, ACTA2*) were similar for HISMCs grown on non-aligned and aligned scaffolds. *VIM* (mean+/- SEM aligned: 0.01462 +/- 0.002633, mean non-aligned: .025 +/-0.0002020, P=0.0052, n=5). *ACTA2* (mean +/- SEM aligned: 0.3656 +/- 0.07053, mean non-aligned: .4986 +/- 0.04450, P=0.1494, n=5). *MYH11* (mean+/- SEM aligned: 0.002982 +/- 0.0006099, mean non-aligned: 0.003978 +/- 0.0004698, P=0.2322, n=5). *COL1A1* (median [interquartile range], aligned: 10.38 [12.679], non-aligned: 18.95 [4.61], P=0.0952, n=5). *FN1* (median [interquartile range], aligned: 6.985 [6.93], non-aligned: 10.21 [8.483], P=0.4206, n=5). *ACTG2* (mean +/- SEM aligned: 0.7499 +/- 0.06910, mean non-aligned: 1.009 +/ 0.1477, P=0.1249, n=5). *MMP14* (median [interquartile range], aligned: 1.16 [1.624], non-aligned: 2.319 [2.898], P=0.2222, n=5).

### Quantitative image analysis

MYH11 fluorescence intensity quantitative analysis employed Imaris software (version 9.02, Bitplane Inc.). Phalloidin staining was used to generate isosurfaces corresponding to individual cells. Cell volume and total MYH11 fluorescence intensity were obtained from the isosurfaces and fluorescence intensity was normalized to cell volume, as previously reported (17). For nuclear to cytoplasmic intensity ratio calculations, images were processed using ImageJ (NIH). Z stacks containing identified cells were condensed to “sum of slices” projections. Cells and their nuclei were outlined using the “freehand selection tool” and intensity was measured for each antibody stain of interest. Parameters calculated included raw intensity and volume of the cell and the nuclei.

### Statistical Analysis

GraphPad Prism (version 9.5.1) was used for statistical analysis of qRT-PCR and quantitative image analysis data. Two-tailed Student’s t-test (parametric data) or Mann-Whitney rank sum tests (nonparametric data) were used for comparisons between 2 groups. P<0.05 was considered significant. Data are represented as mean +/- SEM for parametric data and median [interquartile range] for nonparametric data. Statistical analysis of RNA-seq data was performed as discussed below.

### Bulk RNA Sequencing Analyses

Libraries were prepped using TruSeq total RNA sequencing kit (Illumina, 20020596; TruSeq® Stranded Total RNA Library Prep Human/Mouse/Rat; 48 Samples) and samples were sequenced (paired-end) on Illumina NovaSeq 6000. The bioinformatics pipeline nf-core/rnaseq (18) (reference genome: GRCh38, aligner: STAR (19), quantifier: RSEM (20)) provided counts for 29,972 genes. Additional analyses of bulk RNA-sequencing gene count data were performed in R (v4.4) (21) using RStudio Server (2023.06.1 Build 524) (22). Gene count data was filtered using the *WGCNA* (23) function goodSamplesGenes() with default parameters to remove genes with too many missing entries across samples, resulting in 20,410 remaining genes. Additional filtering removed genes with fewer than 50 counts across all samples. The remaining 14,892 genes were used for downstream analysis. *DESeq2* (v1.44, Wald test method, contrasting loaded versus unloaded samples, and using the log2 fold change (log_2_FC) shrinkage method; Benjamini–Hochberg procedure for FDR) was used for differential gene expression analysis to compare loaded versus unloaded conditions (24).

### Gene Set Enrichment Analysis (GSEA)

Gene Set Enrichment Analysis employed *fgsea* (v3.17) (25) in R. Genes were ranked based on log_2_FC from the differential expression analysis of loaded versus unloaded HISMCs. Normalized enrichment scores (NES) reflect the degree to which genes are overrepresented at the top or bottom of the entire ranked gene list, normalized to the mean enrichment of random samples of the same size. The Human MSigDB Collections Hallmark (MSigDB v7.5.1) (26, 27)) gene sets were used for this analysis (26–28). MSigDB human gene sets were downloaded via the R package *msigdbr* (v7.5.1) (27).

### STRING Analyses

STRING is a database of known and predicted protein-protein interactions. This includes direct/physical and indirect/functional associations (29). STRING (v11.5) was used to examine possible interactions between differentially expressed genes based on DESeq2 analyses. STRING input included 500 genes with the lowest adjusted p-values filtered for log_2_FC < -0.48 or > 0.48). Pathway enrichment analysis in the STRING software used the “whole genome” background option in STRING for statistical comparison employing KEGG and GO Pathways.

### Ligand-receptor association analyses using NicheNet

NicheNet (*nichenetr*, v2.0.0) was employed to characterize potential interactions between HISMC-expressed ligands and receptors present in various bowel cell types (30). NicheNet prioritizes ligands in the “sender/niche” population that are most likely to affect (according to the NicheNet model) the transcriptional state of a gene set of interest in the “target/receiver” population. For our analyses, “sender/niche” genes were genes differentially expressed (identified in DESeq2 analysis) in HISMCs cultured for 6 hours on loaded versus unloaded scaffolds, filtered for adjusted p-value less than 0.05 and log_2_FC < -0.48 (for “up in unloaded”; 14 ligands) or > 0.48 (for “up in loaded”; 57 ligands). Gene symbols were converted to official HUGO Gene Nomenclature Committee at the University of Cambridge (HGNC) symbols using the GeneSymbolThesarus() function in *Seurat* (v4.4.0) (31) prior to NicheNet analysis. The NicheNet analysis was run separately for ligands “up in loaded” and ligands “up in unloaded”. Selected cell types from the human single nucleus RNA-seq (droplet-based MIRACL-seq) dataset published by Drokhlyansky *et al*. were used as the “receiver/target” populations (32). Patient-specific clusters ("H3", "MHC.I_H1", "MHC.I_H9", "OXPHOS_H3") described by Drokhlyansky *et al.* were removed from the dataset prior to analysis. The following cell types from the Drokhlyansky *et al*. paper were used for this analysis with their original cell type annotations: Epithelial, Fibroblast_1, Fibroblast_2, ICCs, Macrophage, and Neuron. Myocyte clusters from Drokhlyansky *et al*. were reannotated for our analysis based on differential gene expression reported by Drokhlyanksy et al. 2020. According to the average log_2_FC in Drokhlyansky et al. in their Supplemental Table 4, Myocyte_3, Myocyte_4, and Myocyte_5 all had MYH11, suggesting that these myocyte clusters are smooth muscle clusters. Myocyte_3 and Myocyte_5 had more ACTG2 than ACTA2, suggesting that these are a visceral SMC clusters (reannotated as VisceralSMC_1 and VisceralSMC_2, respectively), and Myocyte_4 had more ACTA2 than ACTG2, suggesting that this is a vascular SMC cluster (reannotated as VascularSMC). The following cell type groups were generated and used for this analysis by grouping and renaming the original cell type annotations: Glia (includes Glia_1, Glia_2, and Glia_3) and Vascular (includes Vascular_1 and Vascular_2, probably endothelial cells based on marker gene expression). The gene set of interest for each “receiver/target” cell type was defined as differentially expressed genes identified using the FindMarkers function (Wilcoxon Rank Sum test) in Seurat (31). Genes were only considered for differential expression testing if expressed in at least 10% of cells in that population, and differential expression for each cell type compared each cell type against all other cell types in the Drokhlyansky *et al*. dataset. The resulting differentially expressed genes were filtered to keep only genes with adjusted p-value < 0.05 and average log_2_FC > 0.25 or < -0.25. These filtered results were used as the receiver gene sets of interest in the NicheNet analysis. The NicheNet prior model (v2) was modified according to the developer “model construction” instructions to keep only data sources classified as “literature” and “comprehensive_db”. NicheNet ranks potential ligands based on the presence of receptors and target genes in the gene set of interest that are associated with each ligand in the NicheNet model (compared to the background of genes for that cell type; background genes were identified for each cell type individually via the get_expressed_genes() function in NicheNet with default parameters). For each cell type, the results of the ligand activity analysis were filtered based on their log_2_FC (from the DESeq2 differential expression analysis), keeping only the “top 20” ligands (or fewer if not more than 20) by absolute value of log_2_FC for each receiver cell type. To infer potential ligand-to-target signaling paths, the get_ligand_signaling_path() function in NicheNet was run for each set of ligands and receiver cell type. The inferred signaling network for each receiver cell type was filtered to remove target genes if they were also identified as receptors within a particular cell type and to keep only ligand-receptor, ligand-target, and receptor-target links that contributed to a complete ligand-receptor-target signaling path. The Sankey plots of inferred ligand-receptor-target paths were generated using subsets of the “top 20” ligands results for each receiver cell type, separated into “top 10” (ligands 1-10, ranked by greatest fold change) and “next 10” (ligands 11-20, ranked by greatest fold change). As these subsets of ligands were generated individually for each receiver cell type, the “top 10” and “next 10” refers to the top ligands prioritized for each receiver cell type, and the prioritized ligands are not identical for all receiver cell types. A schematic for how NicheNet was used for this analysis is shown in Supplemental Figure 1. Sankey plots were generated using R package *sankeyD3* (v0.3.2) (33) and manually edited in Adobe Illustrator (2023) for legibility.

## Data availability

A Supporting Data Values file is available in supplemental materials. Full data sets are deposited in Gene Expression Omnibus (GEO) accession: GSE264225. Differentially expressed gene lists can be accessed as Supplemental Data. All code and package version information used for DESeq2, GSEA, and ligand-receptor-target NicheNet analyses is available on GitHub at github.com/HeuckerothLab/Mechanobiology_Wolfson2024/.

## Results

### HISMCs grown on aligned scaffolds have more smooth muscle myosin heavy chain 11 (MYH11) protein and less vimentin (*VIM*) mRNA than HISMCs grown on non-aligned scaffolds

Our initial goal was to test the hypothesis that pathological mechanical stress might acutely alter gene expression in contractile bowel smooth muscle cells. One challenge is that SMCs cultured on hard plastic rapidly undergo phenotypic class switching from a “contractile” (MYH11-expressing) phenotype to a “synthetic” phenotype that produces extracellular matrix (ECM), migrates, and proliferates (8, 34). To study the effects of mechanical stress in a more contractile phenotype cell, we seeded HISMCs onto electrospun poly-caprolactone (PCL) scaffolds (35) coated with laminin, an extracellular matrix protein that promotes the contractile SMC phenotype (14, 36) (Figure 1A, B). One set of PCL scaffolds was spun to have aligned fibers that promote growth of elongated spindle-shaped SMCs reported to be more contractile (37, 38). In parallel, HISMCs were cultured on laminin-coated PCL scaffolds with non-aligned fibers (Figure 1A, B). After 72 hours with scaffolds floating freely in HISMC culture media, cells were fixed and stained with antibodies to MYH11, a contractile apparatus protein prominently produced in mature contractile phenotype SMCs. Pixel intensity measurements showed HISMCs grown on aligned scaffolds averaged (mean) 14% more MYH11 protein than HISMC grown on non-aligned scaffolds (Aligned: 54.33 arbitrary units (AU) [32.6 AU]; Non-aligned: 47.6 AU [28.28 AU], median [interquartile range]) (Mann-Whitney test, P<0.0001, n=3) (Figure 1B, C). Vimentin (*VIM*) mRNA, a synthetic SMC marker (39), was also less abundant in HISMCs cultured on aligned compared to non-aligned scaffolds (P=0.0052, n=5) (Figure 1D). In contrast, mRNA for extracellular matrix related genes (*COL1A1*, *MMP14*, *FN1*) and contractile apparatus genes (*MYH11, ACTG2, ACTA2*) were equivalent in HISMCs cultured on aligned versus non-aligned scaffolds (Figure 1 E-J). Based on these findings, further experiments used HISMCs cultured on aligned scaffolds.

### Dynamic loading (cyclic stretching) of HISMCs leads to marked changes in gene expression

To identify early gene expression changes in response to pathologic stretch, HISMCs cultured for 72 hours on aligned PCL scaffolds were subjected to cyclic uniaxial stretch (loaded) along the long axis of the cell (3% stretch, 1 Hz, 6 hours). Unloaded control scaffolds were maintained free-floating in fresh culture media for 6 hours (Figure 2A). Scaffolds were then stained (Figure 2B) or dissolved in Trizol for RNA sequencing. Bulk RNA sequencing demonstrated clear separation of loaded (stretched) versus unloaded (free-floating) HISMCs using Principal Component Analysis (PCA) (Figure 2C). Differential expression analysis using DESeq2 identified 1239 mRNA that were more abundant (log_2_FC > 0.48) and 1261 mRNA that were less abundant (log_2_FC < -0.48) in loaded compared to unloaded HISMCs (adjusted p-value<0.05, DESeq2). This gene set includes markers of the contractile or synthetic SMC phenotypes, inflammatory mediators, TGFβ superfamily genes, axon guidance molecules, cytoskeletal proteins, cell-cell junctional proteins, and cell-ECM interacting proteins (Figure 2D). In addition, Gene Set Enrichment Analysis (GSEA) using the Hallmark gene sets from the Human Molecular Signatures Database (MSigDB) highlighted several cytokine and inflammation pathways with a normalized enrichment score (NES) greater than 2 (Supplemental Figure 2), suggesting that cyclic mechanical stress induces gene expression changes associated with pro-inflammatory states. These pathways included “TNF alpha signaling via NFκB”, “Inflammatory Response”, “Allograft Rejection”, “IL6-JAK-STAT3 Signaling”, “IL2-STAT5 Signaling”, and “Interferon Gamma Response”.

**Figure 2:**
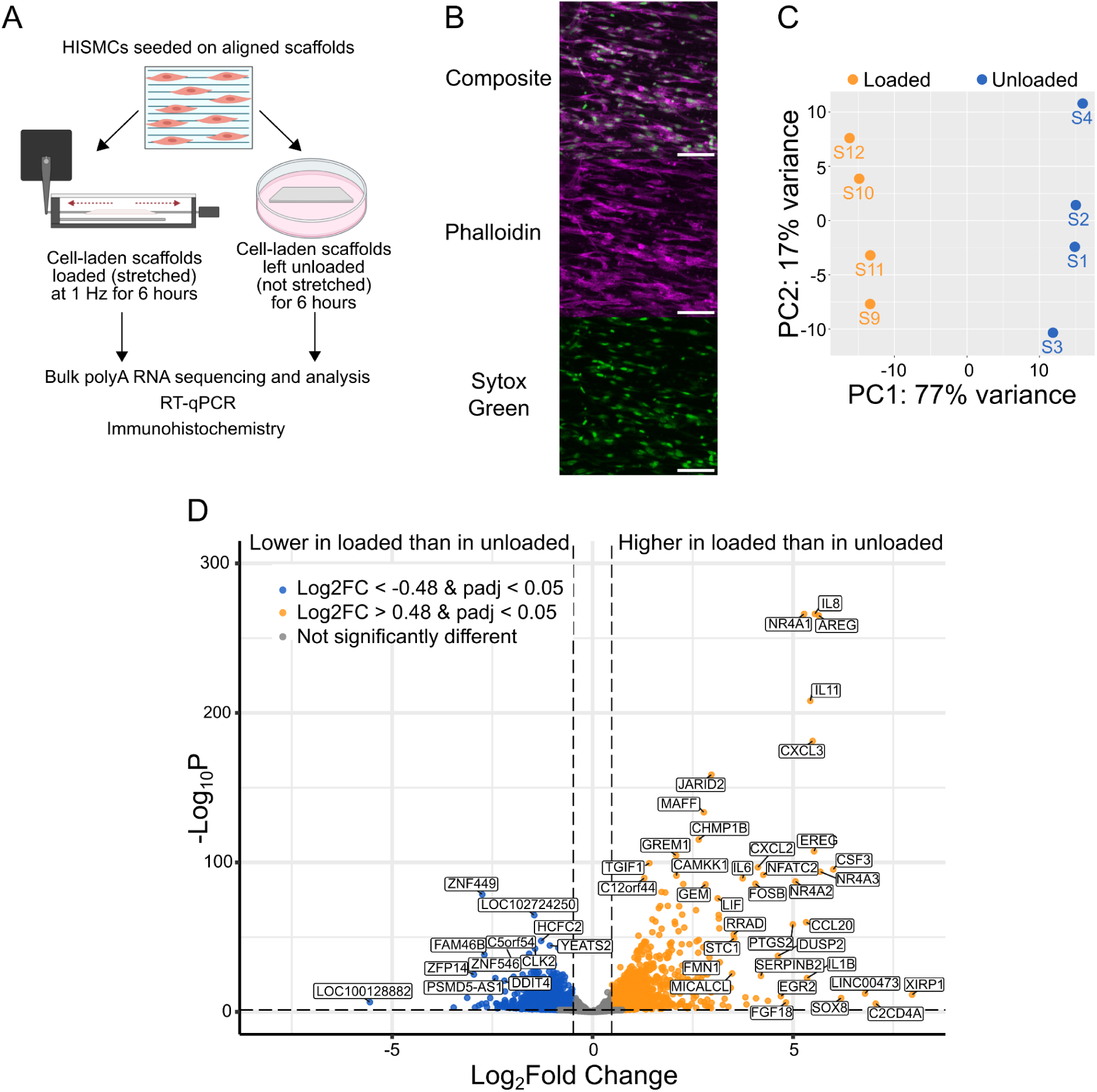
Dynamic loading for 6 hours led to many changes in gene expression. (A) Schematic of experimental design with primary outputs. (B) Sum-of-slices Z-projection of 20x confocal image of HISMCs on aligned scaffolds stained for F-actin (Phalloidin, magenta) with nuclei labeled with Sytox Green. Scale bar = 100 μm. (C) PCA plot (based on the top 500 most variable genes) showing that loaded versus unloaded HISMCs differ significantly in gene expression. Orange dots = loaded samples, blue dots = unloaded samples. (D) Volcano plot of differentially expressed genes between loaded vs unloaded HISMCs, from DESeq2 analysis. Log_2_fold change (log_2_FC) cutoff > 0.48 or < -0.48 (vertical dotted lines). False Discovery Rate (FDR) cutoff = 0.05 (horizontal dotted line). Orange dots indicate differentially expressed genes (FDR<0.05) with log_2_FC>0.48 (1239 genes up in loaded), and blue dots indicate differential expressed genes (FDR<0.05) with log_2_FC<-0.48 (1261 genes up in unloaded).

Consistent with the hypothesis that loading induced a pro-inflammatory state, STRING classification using KEGG pathways to characterize the 500 most differentially regulated genes (based on adjusted p-values) identified 13 NFκB pathway genes more abundant in loaded than in unloaded HISMCs (Figure 3A). To determine if this reflected increased NFκB signaling, we used immunohistochemistry and discovered more nuclear NFκB in loaded compared to unloaded HISMCs (Figure 3B,C). Since NFκB (NFKB1, NFKB2) also promotes a synthetic phenotype in SMCs by repressing myocardin, the master regulator for SMC contractile phenotype (40), we hypothesized loaded HISMCs might have a more synthetic phenotype than unloaded HISMCs. Consistent with this hypothesis, loaded HISMCs had higher levels of mRNA for many synthetic SMC phenotype genes compared to unloaded HISMCs, including *EREG* (41)*, AREG* (42)*, KLF4* (43), *PDGFA* (44), *EPHA2* (*45*), *ETS1*, *ETS2*, *ELF1* (*46*), *POU2F2* (47), and *THBS1* (48) (Figure 3D). Loaded HISMCs also had less MKL2 protein in the nucleus compared to unloaded HISMCs (Figures 3E, F) and less mRNA for MKL2 (log_2_FC=-0.5, p-val adj=0.0029). MKL2 is a myocardin transcription factor family gene that promotes the SMC contractile phenotype (49). Furthermore, loaded HISMCs had less mRNA for *CARMN* (reported as *MIR143HG* in Supplemental Table 3) (log_2_FC=-2.06, p-val adj=9.68E-05), a long noncoding RNA critical for maintaining visceral SMC contractile function (50) (Figure 3G). Collectively, these data show our cyclic stretching paradigm promotes a synthetic, pro-inflammatory state in HISMCs, instead of a contractile phenotype.

**Figure 3:**
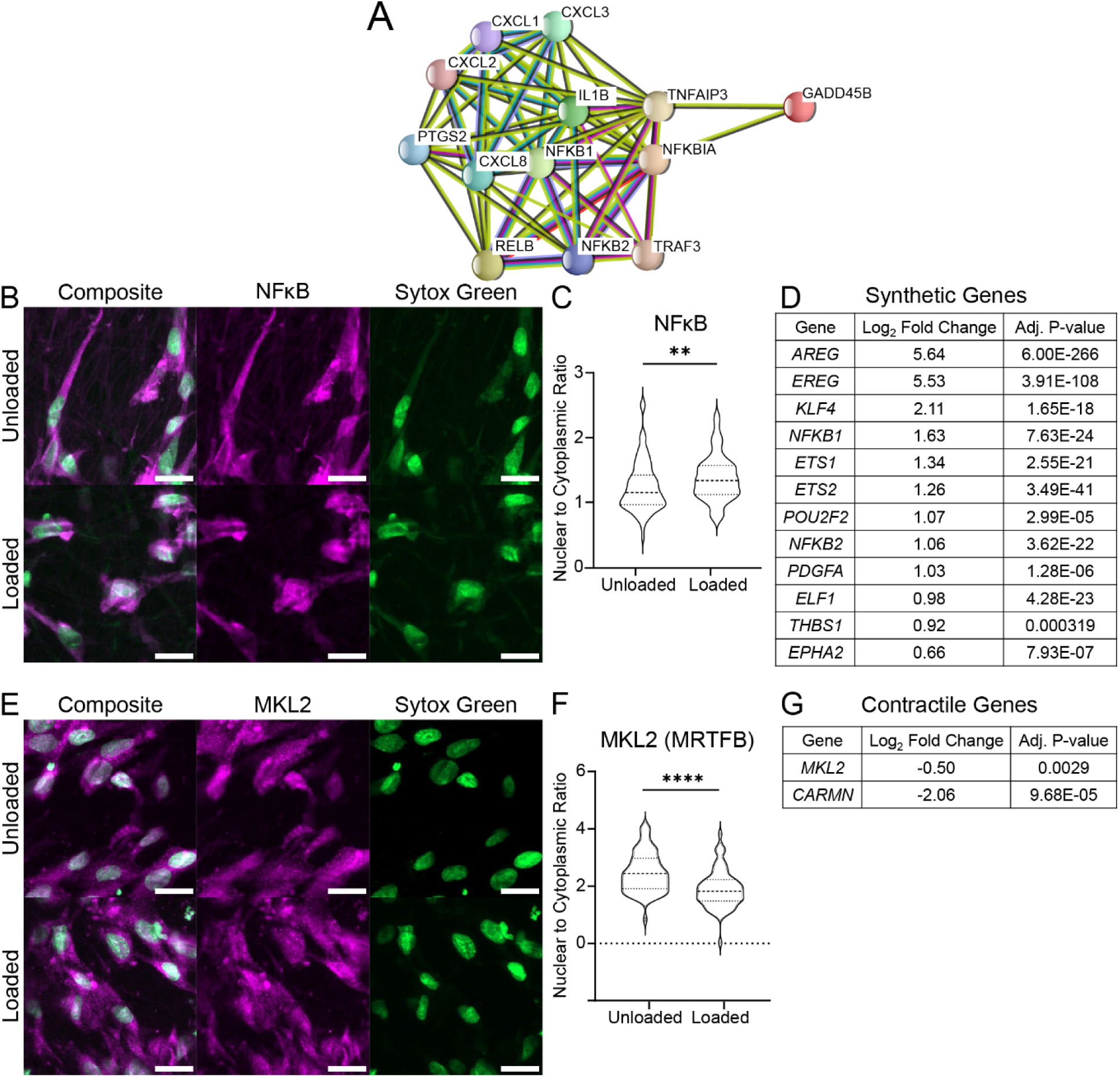
Loaded HISMCs have increased synthetic gene expression and activation of NFκB signaling. **(A)** STRING diagram of KEGG NFκB pathway analysis. Input included mRNA more abundant in loaded HISMCs with Log2 fold change (log_2_FC) >0.48 based on bulk RNA sequencing. (**B**) Representative sum-of-slices Z-projections of confocal images (63X oil objective) showing NFκB antibody staining (red) and Sytox green nuclear staining. Top: Unloaded HISMCs grown on an aligned scaffold. Bottom: Loaded HISMCs grown on an aligned scaffold. Scale bar = 20 μm. **(C)** Quantitative analysis of antibody staining demonstrated increased NFκB nuclear to cytoplasmic ratio in loaded compared to unloaded HISMCs (median [Interquartile range] unloaded 1.152 [0.4651], loaded 1.336 [0.8271] (P=.0081, Mann Whitney), n=79 cells for both groups. (**D**) Smooth muscle synthetic genes differentially expressed in loaded versus unloaded HISMC based on bulk RNA sequencing. (**E**) Representative sum-of-slices Z-projections of confocal images (63X oil objective) of MKL2 stained HISMCs (red) and Sytox green nuclear staining. Top: Unloaded HISMCs grown on an aligned scaffold. Bottom: Loaded HISMCs grown on an aligned scaffold. (**F**) Quantitative analysis of antibody staining demonstrated an increased MKL2 nuclear to cytoplasmic ratio in unloaded compared to loaded HISMCs. (median [Interquartile range] unloaded 2.435 [1.066], loaded 1.820 [0.731] (P<.0001, Mann Whitney), n=65 cells for unloaded and n=70 cells for loaded HISMCs. **(G)** Smooth muscle contractile genes differentially expressed in loaded versus unloaded HISMCs based on bulk RNA sequencing.

These findings are reinforced by STRING classification of the top 500 genes (by adjusted p-value) with absolute value log_2_FC > 0.48 using the Gene Ontology (GO) Biological Process pathways (51, 52). In the STRING analysis, 30 of the top 500 genes were involved in cytokine signaling (cytokine-mediated signaling pathway, GO:0019221) including *IL8, CXCL3, IL11, IL1B, CCL20, PTSG2, CXCL2, IL6, LIF, IL24, CXCL1, CXCL5, and CLCF1* (Figure 4A-B).

**Figure 4:**
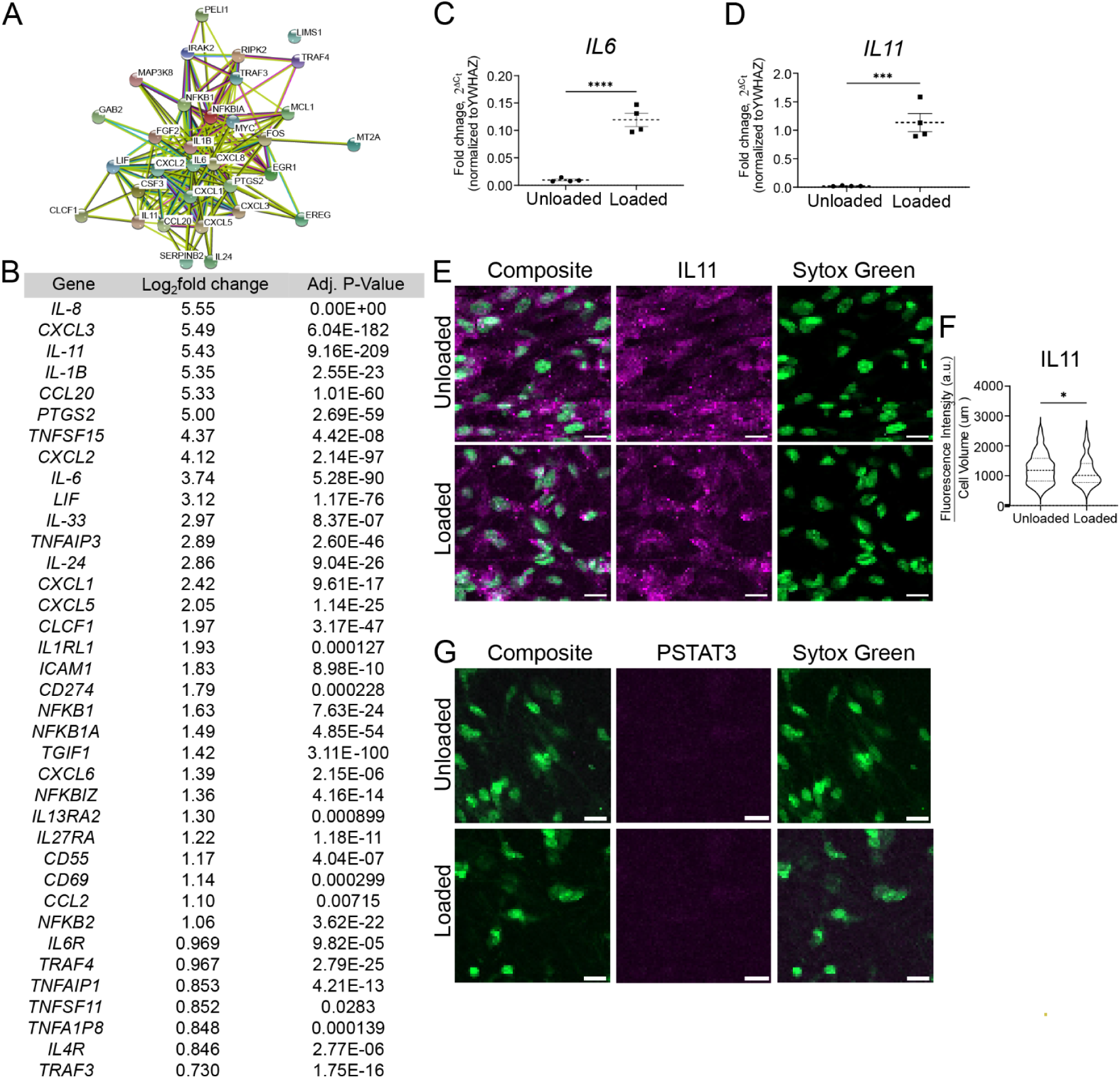
Supraphysiologic cyclical stretching stimulates production of cytokines, cytokine receptors, and chemokines. **(A)** STRING diagram of 32 genes (from the top 500 differentially expressed genes with padj <10^-6 genes and absolute value log_2_FC > 0.48, out of 1113 genes meeting this criteria) from bulk RNA sequencing that were annotated as genes in the GO Biological Process cytokine-mediated signaling pathway (GO:0019221) in the STRING analysis. **(B)** Genes for cytokines, cytokine receptors, and chemokines significantly differentially expressed from bulk RNA sequencing (P<0.05). All were significantly upregulated in loaded HISMCs. **(C)** qRT-PCR shows 12-fold more *IL-6* mRNA in loaded compared to unloaded HISMCs. (mean +/- SEM unloaded: 0.009601 +/- 0,001695, loaded: 0.1195 +/- 0.01191, [0.01203], P<.0001, n=4). (**D**) qRT-PCR shows 55-fold more *IL11* mRNA in loaded compared to unloaded HISMCs. (mean +/- SEM unloaded: 0.02060 +/- 0.002797, loaded: 1.136 +/- 0.1585, [0.1585], P=0.0004, n=4). (**E**) Representative sum-of-slices Z-projections of confocal images of loaded or unloaded HISMCs stained with antibodies to IL11 (63X oil objective, confocal Z-stack). (**F**) Quantification of IL11 immunohistochemistry showed a small, but statistically significant increase in pixel intensity in unloaded compared to loaded HISMCs (median [interquartile range], unloaded: 1184 [762.6], n=122; loaded: 1012 [631.2], n=99; P=0.0469, Mann-Whitney). (**G**) Representative sum-of-slices Z-projections of confocal images of loaded or unloaded HISMCs stained with antibodies to phosphorylated STAT3 (PSTAT3) (63X oil objective, confocal Z-stack). PSTAT3 was not detected in HISMCs under either condition, but PSTAT3 was readily detectable in the human monocyte cell line THP-1 (Supplemental Figure 3).

Many of these cytokines may impair intestinal motility (53–55). To validate RNA sequencing, we used qRT-PCR to analyze mRNA abundance for *IL6* (Figure 4C), a major proinflammatory cytokine (56), and *IL11* (Figure 4D), which promotes a synthetic phenotype in vascular smooth muscle (55). The qRT-PCR showed *IL6* mRNA was 12-fold more abundant (P<0.001, n=4) in loaded versus unloaded HISMCs and *IL11* mRNA was 55-fold (P=0.004, n=4) more abundant in loaded HISMCs. This is similar to the 13.4-fold (log2FC=3.74) elevation in *IL-6* and 43.1-fold (log2FC=5.43) elevation in *IL11* based on RNA sequencing (Figure 4B). In contrast to mRNA data, IL11 immunohistochemistry revealed lower protein levels in loaded than in unloaded HISMCs (Figures 4E, F). In addition, phospho-STAT3, a key IL6 signaling protein, was not detected in either loaded or unloaded HISMCs by antibody staining (Figure 4G) although our antibody readily detected phospho-STAT3 in human THP-1 macrophages (Supplemental Figure 3). Collectively, these data show dramatic increases in many proinflammatory signaling molecules at the mRNA level after only 6 hours of pathologic stretching.

### TGFβ superfamily genes are differentially expressed in loaded versus unloaded HISMC

TGFβ signaling has been shown to have roles in smooth muscle embryogenesis and phenotypic class switching (57). Many TGFβ superfamily genes were differentially regulated by 6 hours of cyclic HISMC loading based on RNA sequencing. Loaded HISMCs had higher levels of *INHBB, TGFBR1, TGFBR3, SMAD7, TGFB1, BMP2, GREM1,* and *SMAD1* mRNA, and lower levels of *TMEM100, SMAD6, BAMBI*, *BMP4, SMAD6,* and *BAMBI* mRNA, compared to unloaded HISMCs (Figure 5A). qRT-PCR confirmed higher levels of *BMP2* (Figure 5B) and *GREM1* (Figure 5C) in loaded HISMCs and reduced *BMP4* mRNA (Figure 5D) compared to unloaded cells. Since TGFβ and BMP can alter SMC phenotype, we evaluated nuclear to cytoplasmic ratios of signaling proteins that localize to the nucleus after BMP (phospho-SMAD1/5/8) or TGBβ (phospho-SMAD2/3) receptor activation (Figure 5E). Quantitative analysis of immunohistochemistry showed equivalent nuclear to cytoplasmic ratios of phospho-SMAD2/3 and phospho-SMAD1/5/8 in loaded and unloaded HISMCs (Figure 5F, G). Collectively, these data indicate cyclic stretching rapidly alters mRNA levels for many TGFβ superfamily genes, but that signaling pathways these genes could activate or inhibit were not altered in HISMCs, at least at this early time point.

**Figure 5:**
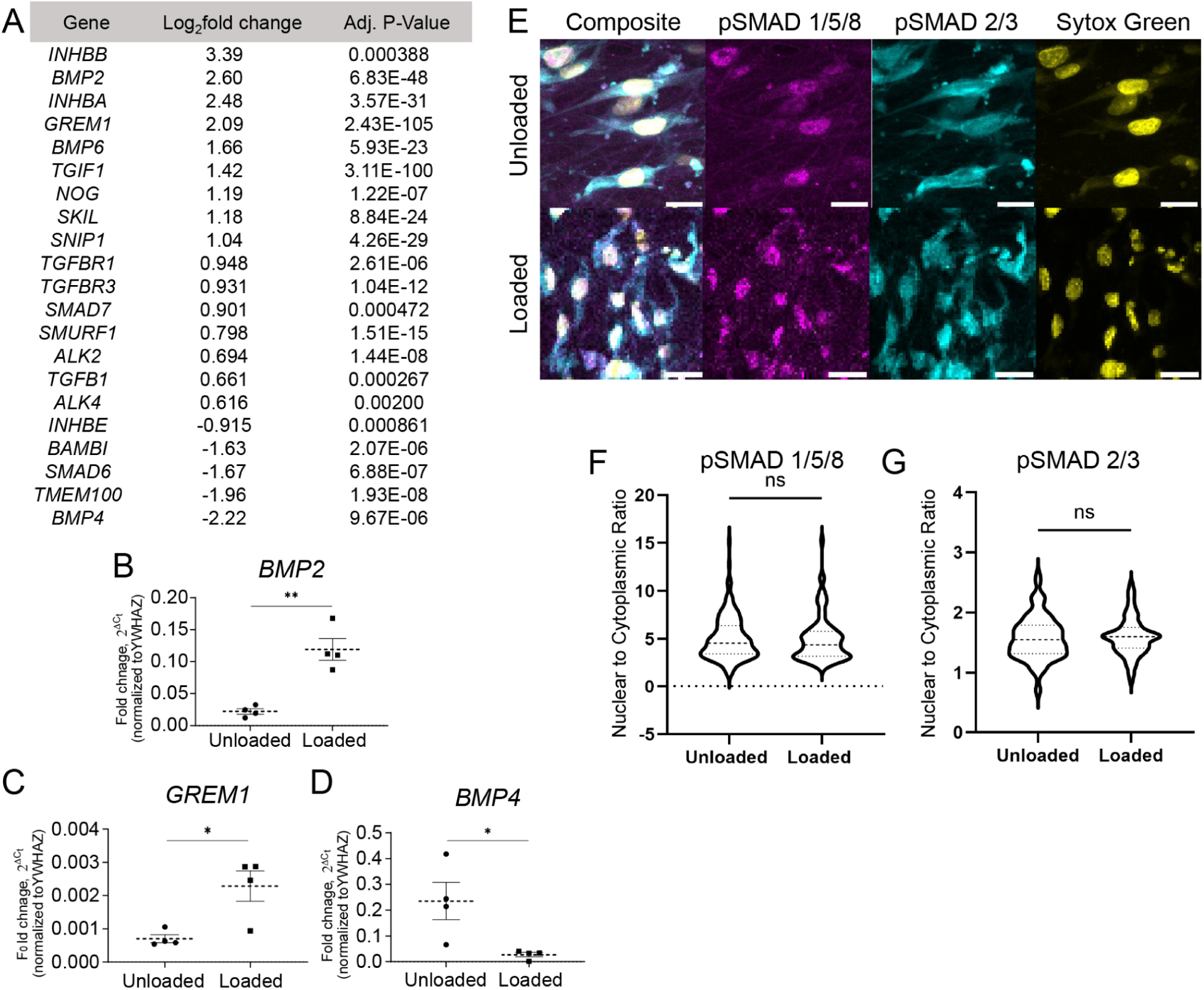
TGF-β superfamily genes are differentially expressed as a result of HISMC loading. (**A**) TGFβ superfamily genes differentially expressed between loaded and unloaded HISMCs identified by DESq2. (**B**) *BMP2* RTPCR results confirming 5.36x increased BMP2 expression in loaded HISMCs (mean +/- SEM unloaded: 0.02222 +/- 0.004256, loaded: 0.1191 +/- 0.01708, [0.01761], P=0.0015, n=4). (**C**) *GREM1* RTPCR results 3.16x increased expression in loaded HISMCs. (mean +/- SEM unloaded: 0.0007025 +/- 0.0001186, loaded: 0.00289 +/- 0.0004599, [0.0004749], P=0.0156, n=4). (**D**) *BMP4* RTPCR results confirming 8.63x increased BMP4 expression in unloaded HISMCs. (mean+/-SEM unloaded: 0.2358 +/- 0.07209, loaded: 0.02733 +/- 0.008379, [0.07260], P=0.0284, n=4). (**E**) Representative images taken with confocal microscope 63X oil objective of PSMAD 1/5/8 (left) and PSMAD 2/3 (right) in unloaded and loaded HISMCs. (**F**) IHC quantification of PSMAD 1/5/8 showed no differences in nuclear to cytoplasmic staining between loaded and unloaded HISMCs (median [interquartile range], unloaded: 4.549 [2.937], n=150; loaded: 4.391 [2.584], n=88; P=0.5339, Mann-Whitney). (**G**) IHC quantification of PSMAD 2/3 showed no differences in nuclear to cytoplasmic ratio between loaded and unloaded HISMCs (median [interquartile range], unloaded: 1.547 [0.471], n= 150; loaded: 1.598 [0.344], n=88; P=0.5099, Mann-Whitney).

### Pathologic loading induces differential expression of guidance molecules and of genes needed for cell-cell and cell-extracellular matrix (ECM) interactions

Cyclic HISMC loading rapidly altered mRNA levels for many ephrins, semaphorins, netrins, and slits (Table 1). In addition to central roles in neurobiology, these axon guidance molecules play key roles in vascular smooth muscle cell migration, cell proliferation, and inflammation in the context of cardiovascular disease (58). Several mRNAs involved in cell-ECM or cell-cell interactions, with possible roles in mechanosensation, were differentially expressed between loaded and unloaded HISMCs. These mRNA included integrins, cadherins, catenins and catenin antagonists, claudins, a tight junction protein, talins, syndecans, an actinin, an adherens junction protein, cell adhesion molecules, and focal adhesion genes (Table 2). Finally, there were differential changes in mRNA levels for many cytoskeletal proteins (Table 3). These changes in gene expression suggest that in response to cyclic stretching, HISMCs alter cell-cell and cell-ECM interactions, possibly consistent with a transition away from the contractile SMC cell fate.

**Table 1.** Pathologic loading leads to differential expression of guidance molecule genes.

| Gene | Log<sub>2</sub>fold change | Adj. P-Value | Type of Protein |
| --- | --- | --- | --- |
| *EFNB2* | 1.15 | 4.20E-05 | Ephrins |
| *EFNA4* | -0.665 | 0.000108 | Ephrins |
| *EFNA3* | -1.05 | 0.0108 | Ephrins |
| *EFNA1* | -1.47 | 0.0152 | Ephrins |
| *EPHA2* | 0.66 | 7.93E-07 | Ephrin Receptors |
| *EPHB2* | 0.40 | 0.000487 | Ephrin Receptors |
| *EPHB3* | -2.24 | 0.000554 | Ephrin Receptors |
| *SEMA7A* | 1.21 | 5.50E-07 | Semaphorins |
| *SEMA4C* | 0.55 | 1.61E-07 | Semaphorins |
| *SEMA5A* | -0.34 | 0.00793 | Semaphorins |
| *SRGAP2B* | -0.24 | 0.0252 | Slits |
| *SLITRK5* | -0.48 | 0.0368 | Slits |
| *SRGAP2* | -0.51 | 1.40E-13 | Slits |

Positive log<sub>2</sub>fold change denotes upregulated in loaded HSMCs.
Negative log<sub>2</sub>fold change denotes upregulated in unloaded HSMCs.
Adjusted P-value <0.05 was considered to be significant.

**Table 2.**
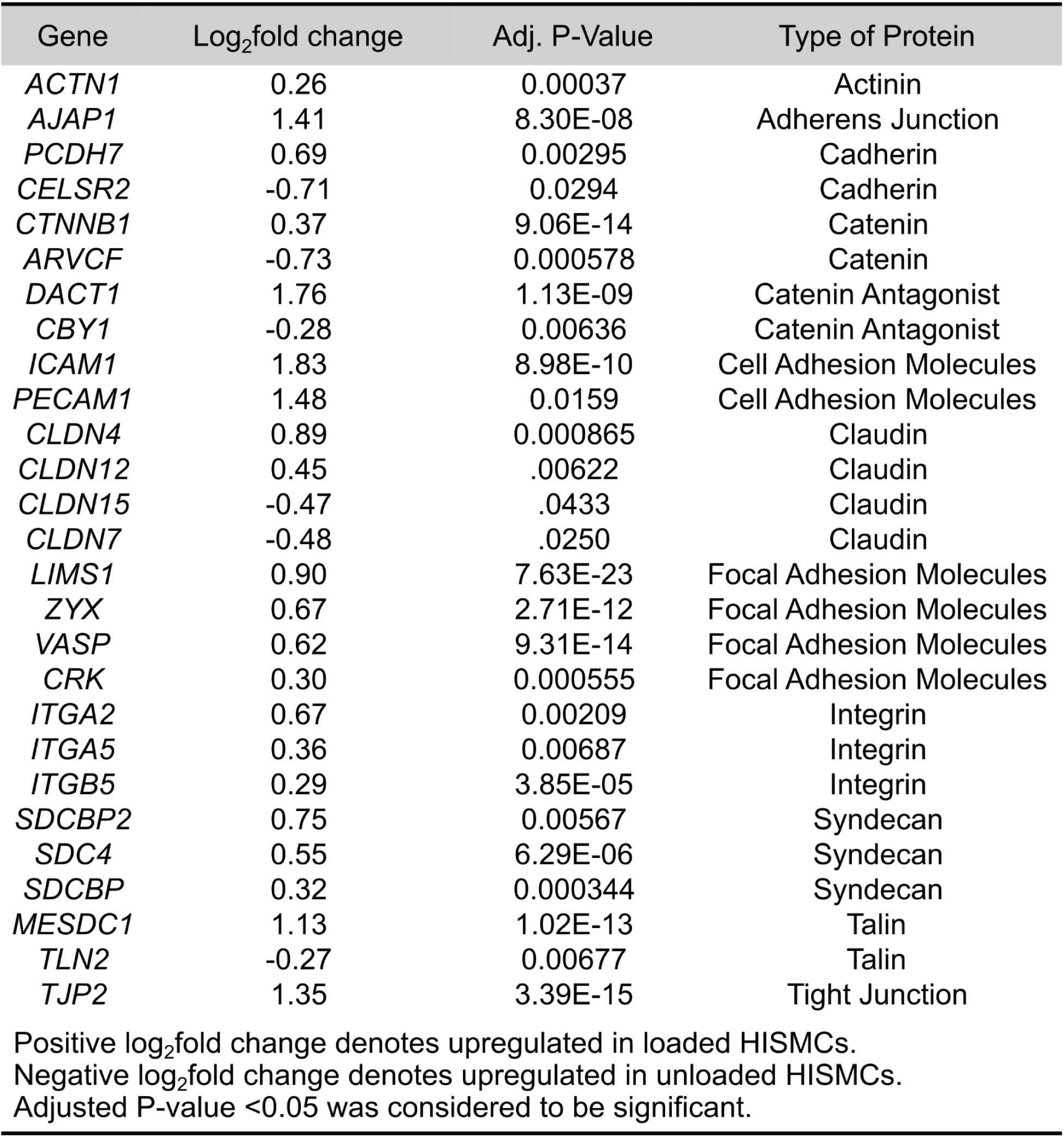
Pathologic loading leads to differential expression of cell-cell junction, and cell-ECM junction mechanosensor genes.

**Table 3.** Pathologic loading leads to differential cytoskeleton-associated genes.

| Gene | Log<sub>2</sub>fold change | Adj. P-Value |
| --- | --- | --- |
| *FMN1* | 3.18 | 6.12E-34 |
| *TUBA8* | 0.817 | 0.0170 |
| *MYO1E* | 0.629 | 2.88E-09 |
| *CAMSAP1* | 0.576 | 6.32E-16 |
| *FERMT2* | 0.542 | 0.000554 |
| *MYO9B* | 0.431 | 2.24E-05 |
| *TUBB6* | 0.425 | 1.02E-04 |
| *MYO10* | 0.417 | 2.94E-05 |
| *CAMSAP2* | 0.321 | 0.0231 |
| *ACTG1* | 0.319 | 0.000481 |
| *TTLL5* | 0.310 | 0.000253 |
| *MYL12B* | 0.289 | 3.24E-05 |
| *TUBB2A* | 0.256 | 0.00516 |
| *SYNM* | 0.249 | 0.0487 |
| *MYL12A* | 0.243 | 0.0336 |
| *CTTN* | 0.160 | 0.00332 |
| *TTLL4* | -0.308 | 0.0196 |
| *TUBG2* | -0.320 | 0.0401 |
| *TUBE1* | -0.345 | 0.00250 |
| *UTRN* | -0.393 | 0.00573 |
| *MYO9A* | -0.509 | 2.80E-07 |
| *TTLL11* | -0.825 | 3.14E-05 |
| *VILL* | -0.901 | 5.49E-06 |
| *VMAC* | -1.04 | 0.00163 |

Positive log<sub>2</sub>fold change denotes upregulated in loaded HSMCs. Negative log<sub>2</sub>fold change denotes upregulated in unloaded HSMCs. Adjusted P-value <0.05 was considered to be significant.

### HISMC loading induced expression of diverse ligands that could signal to nearby cells

Remarkably, many genes differentially regulated in HISMCs in response to cyclic loading encode secreted or cell surface ligands that could impact the biology of nearby cells by binding cell surface receptors. To identify possible cellular targets for differentially expressed HISMC ligands, we used NicheNet (30) and human bowel single nucleus RNAseq data from Drokhlyansky *et al.* (32). An overview of the analysis is presented in Supplemental Figure 1. NicheNet evaluates potential ligand-receptor interactions, ranking interactions based on ligand-target regulatory potential (incorporating intracellular signaling into regulatory potential scoring). These potential ligand-receptor interactions are represented in Sankey plots (Figures 6 and 7). On the left of each Sankey plot is a ligand whose mRNA is more abundant in loaded HISMCs than in **<u>un</u>**loaded HISMCs (Figure 6), or conversely, a ligand more abundant in **<u>un</u>**loaded HISMCs than in loaded HISMCs (Figure 7), based on our data. In the middle column are receptors (from Drokhlyansky et al.’s data (32)) for ligands differentially expressed from our data. On the right are genes whose activity or expression is regulated by receptor signaling, according to the NicheNet model. The color of each line indicates the receptor-bearing cell type, based on Drokhlyansky et al.’s data. Figures 6 and 7 show the top 10 prioritized ligands from HISMCs (based on log_2_FC) for each cell type in a re-annotated subset of Drokhlyansky et al.’s data, with additional data in Supplemental Figure 4 showing genes more abundant in loaded HISMCs for the next 10 prioritized ligands (Sankey figures show more than 10 ligands because the top prioritized ligands were not identical for all cell types). For example, *BMP4* mRNA is increased in unloaded compared to loaded HISMCs (as we confirmed in Figure 5A, B). The Sankey plot (Figure 7) shows that receptors for *BMP4* (i.e., *BMPR1A, BMPR1B*, and *BMPR2*) are expressed in visceral smooth muscle (VisceralSMC_1). However, *BMPR1A* is also expressed in enteric neurons, macrophage, Fibroblast_1, and epithelial cells, while *BMPR1B* is expressed in neurons, Interstitial cells of Cajal (ICC), and Fibroblast_1. The co-receptor *BMPR2* is expressed in VisceralSMC_1, vascular endothelial cells, enteric neurons, ICC, Fibroblast_1, and epithelial cells, but was not detected in macrophages in the Drokhlyansky et al. dataset. While some differentially-expressed HISMC ligands could signal to many adjacent cell types (e.g., *FGF18, IL6, IL11, AREG, EREG, BMP2, HBEGF* in loaded HISMCs; *GDF5, BMP4, EFNA1, EFNA3, EFNA4* in unloaded HISMCs), other differentially-expressed HISMC ligands were predicted to signal to only to neurons (*TNFSF15, IL16, INHBB* in loaded HISMCs; *ADM, APLN* in unloaded HISMCs) or to neurons, vascular endothelial cells, and macrophages (*CSF3* in loaded HISMCs) (for example). Notably, differentially-expressed HISMC ligands from the loaded cells have the largest number of targets in neurons, leading to the intriguing hypothesis that neurons may play an active role in how the bowel responds to pathologic mechanical stress. For differentially-expressed HISMC ligands in the unloaded cells, there is a lower number of targets identified across all the examined bowel cell types compared to the number of targets identified across all of the bowel cells types for differentially-expressed HISMC ligands in the loaded cells. Nevertheless, neuronal targets again feature most prominently. We note that some possible interactions indicated in the Sankey plots may not be biologically relevant (e.g., smooth muscle ICAM might never contact bowel epithelial cells). Nonetheless, these NicheNet analyses suggest that altered mechanical stress induces broad changes in HISMC gene expression, and that many differentially expressed genes are likely to bind to receptors, and influence function, of other cell types in the bowel wall.

**Figure 6.**
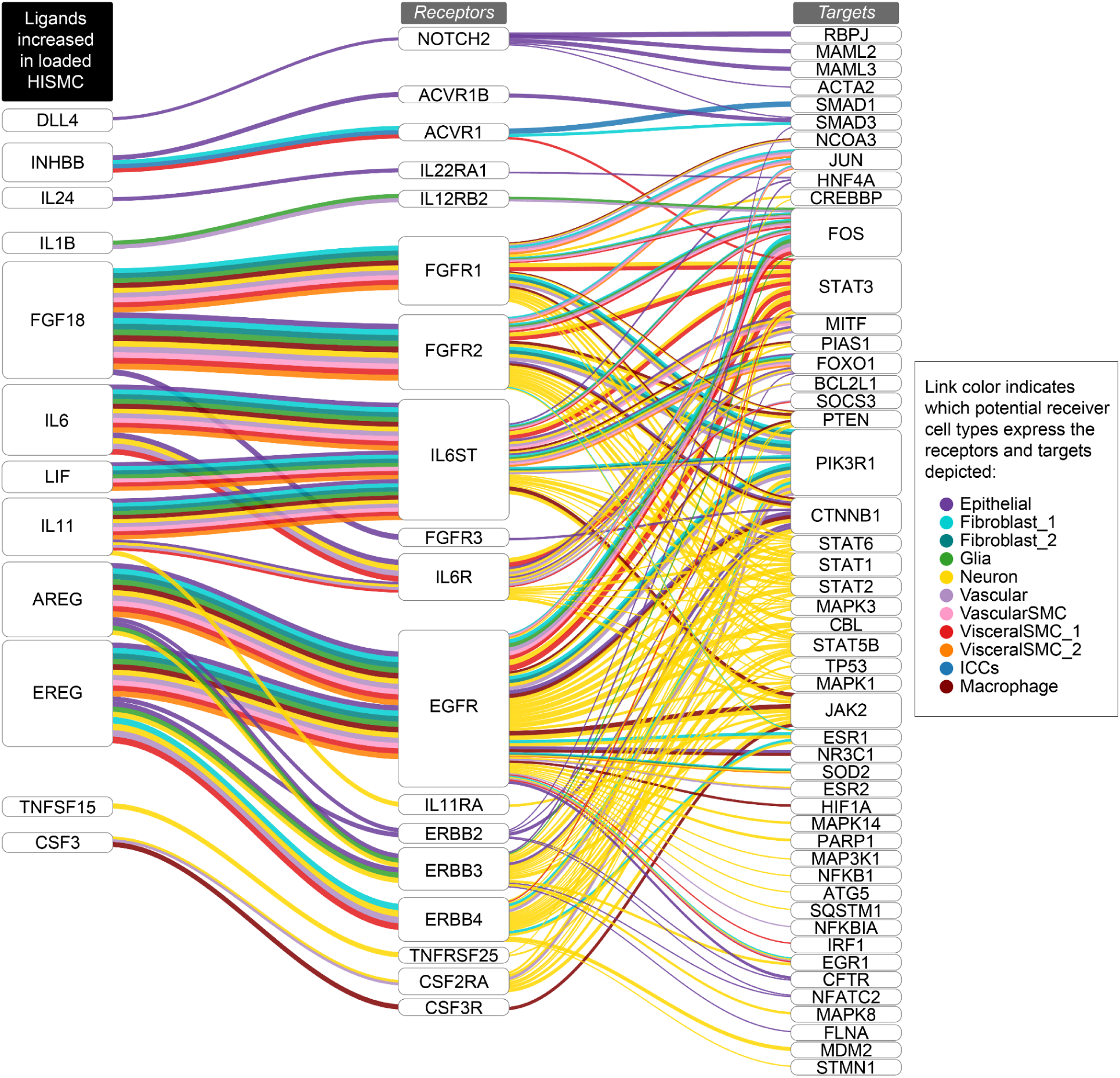
Sankey plot showing the potential ligand-receptor-target links based on NicheNet’s inferred signaling paths from “top 10” ligands upregulated in the loaded HISMC to Drokhlyansky *et al.* receiver cell targets. The NicheNet prioritized ligand analysis between secreted ligands that are more abundant in loaded compared to **<u>un</u>**loaded HISMCs (left column) and receptors (middle column) and target genes (right column) in re-annotated Drokhlyansky *et al.* receiver cell types was used to infer signaling paths from each ligand to target. Potential ligand-receptor-target links were determined based on the inferred signaling paths from NicheNet. See Methods and Supplemental Figure 1 for additional details on the NicheNet analysis and process for inferring ligand-receptor-targets paths.

**Figure 7.**
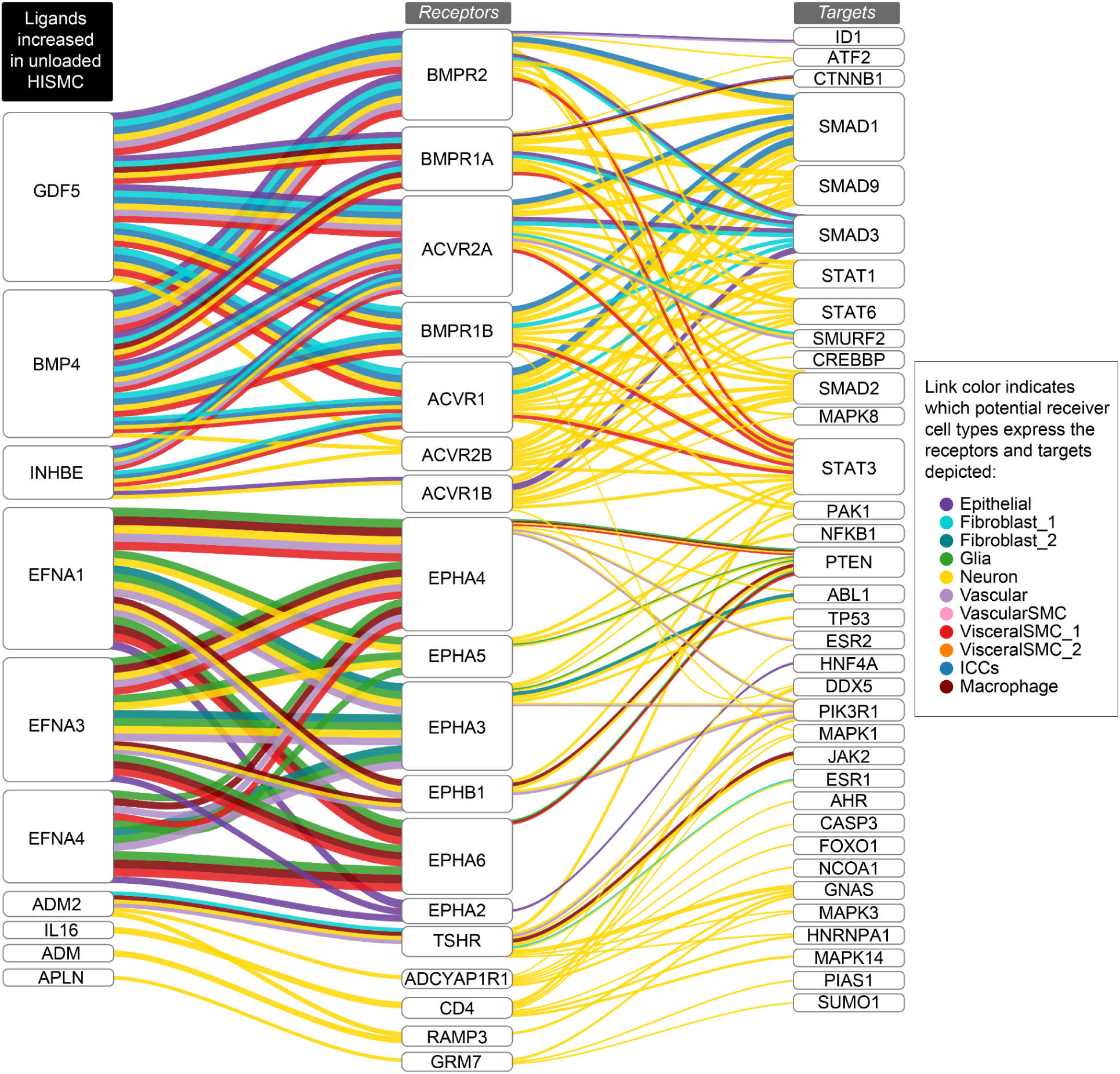
Sankey plot showing the potential ligand-receptor-target links based on NicheNet’s inferred signaling paths from “top 10” ligands upregulated in the unloaded HISMC to Drokhlyansky *et al.* receiver cell targets. The NicheNet prioritized ligand analysis between secreted ligands that are more abundant in **<u>un</u>**loaded compared to loaded HISMCs (left column) and receptors (middle column) and target genes (right column) in re-annotated Drokhlyansky *et al.* receiver cell types was used to infer signaling paths from each ligand to target. Potential ligand-receptor-target links were determined based on the inferred signaling paths from NicheNet. See Methods and Supplemental Figure 1 for additional details on the NicheNet analysis and process for inferring ligand-receptor-targets paths.

## Discussion

Mechanotransduction describes the ability of cells to actively sense, integrate, and convert mechanical stimuli into biochemical signals, including changes in transcription (59). Mechanotransduction is critical for normal bowel physiology, and also impacts disease pathophysiology in the context of pathologic mechanical stressors. Such pathologic force occurs in functional bowel obstruction (visceral myopathy, chronic intestinal pseudo-obstruction, Hirschsprung disease, ileus), mechanical bowel obstruction (volvulus, adhesions, malignancy), and in the context of transmural inflammatory infiltrates or fibrosis (Inflammatory bowel disease, scleroderma). Mechanical stress sensing also impacts symptoms in irritable bowel syndrome, functional dyspepsia, bowel diverticula, functional nausea, centrally mediated abdominal pain syndrome, dyssynergic defecation, achalasia, functional dysphagia, and visceral hypersensitivity (5). Although most bowel cell types appear capable of mechanotransduction (5), the impact of mechanical force on visceral smooth muscle phenotype remains under-explored.

Here, we tested the hypothesis that bowel smooth muscle phenotype might change in response to abnormal mechanical stress. Using cultured human intestinal smooth muscle, we show that even modest pathological stretching (3% uniaxial cyclic stretch at 1 Hz) can rapidly (within 6 hours) alter expression of 4537 genes in HISMCs, of which 2500 met our minimum cut-offs for fold change. The gene expression changes suggest mechanical loading induces HISMCs to transition to a synthetic, pro-inflammatory state. Predictive modeling with NicheNet further suggests that many of the genes induced by pathologic mechanical stress could act on a wide array of nearby cell types, causing complex and long-lasting changes in bowel physiology. The gene expression changes induced in HISMCs by low amplitude, high frequency cyclical stretch are consistent with “phenotypic class switching”, a well-described phenomena in vascular smooth muscle but also reported in visceral smooth muscle.

Unlike all other muscle types, smooth muscle is not terminally differentiated. SMC phenotypic class switching is an unusual attribute, describing the ability of SMCs to reversibly modulate cell fate in response to various mechanical, chemical, and cytoskeletal triggers (8, 60). There are two recognized phenotypes for smooth muscle (contractile and synthetic) (7, 8, 61, 62), but in vascular SMCs there may be as many as 9 identified SMC cell fates (63). These changes in SMC fate may be protective (e.g. forming a fibrous cap in damaged vasculature), but are also implicated in pathophysiology in various gastrointestinal, cardiovascular, and pulmonary diseases (7, 54, 62, 64, 65). In some cases, prevention or reversal of SMC phenotypic class switching provides a therapeutic target (e.g., in atherosclerosis) (66–68). While under-explored, SMC phenotypic class switching may be an important mechanism, and potential therapeutic target, in some types of bowel disease. Understanding mechanisms that underlie visceral SMC phenotypic class switching may, therefore, provide new avenues to prevent progression or reverse damage in human bowel disease.

### Mechanical stress induces phenotypic class switching from contractile to synthetic, proinflammatory HISMC

Contractile SMCs generate the force needed for normal bowel motility. These SMCs may experience pathologic mechanical stress in many settings, and appear to adapt over extended time periods. For example, after surgical manipulation, the bowel stops moving, a problem called “ileus” that typically lasts for many days. In contrast, mechanical obstruction leads to the early occurrence of high frequency clustered contractions (3-10 regular contractions, occurring 1 contraction per 5 seconds, lasting < 1 minute, repeating every 1-3 minutes) called “minute rhythm” and “prolonged simultaneous contractions” (> 8 seconds duration) (69, 70). These patterns also occur in intestinal neuropathy (2). The initial increase in motor activity after bowel obstruction is followed by suppression of motor activity and then by bowel muscle layer hypertrophy (2, 71). These observations reflect complex interactions between many cell types and provide context for our HISMC data.

One striking observation is that loaded HISMC had 30% less *MKL2* and 76% less *CARMN* mRNA after only 6 hours of cyclic stretching (Figure 3G, Supplemental Table 3). MKL2 and *CARMN* are crucial for expression of contractile apparatus genes and are abundant in contractile phenotype SMCs (50, 72). At the same time, loaded HISMCs had more mRNA encoding proteins that block contractile apparatus gene expression or that induce the synthetic/proliferative SMC cell fate (e.g., *AREG* (increased 49.9-fold), *EREG* (increased 46.2-fold), and *KLF4* (increased 4.3-fold) (Figure 3D, Supplemental Table 3) - identified in vascular SMC literature) (73–75). Loaded HISMCs had much higher levels of proinflammatory cytokines, including *IL8* (increased 46.9-fold), *CXCL3* (increased 44.9-fold), *IL11* (increased 43.1-fold), *IL1β* (increased 40.8-fold), *PTGS2* (increased 32.0-fold), *IL6* (increased 13.4-fold), *ICAM1* (increased 3.6-fold), and *CCL2* (increased 2.1-fold), amongst other genes (Figure 4B, Supplemental Table 3). Some of these observations fit with known signaling pathways. For example, IL11 is produced by vascular smooth muscle in response to TGFβ1, and then acts cell-autonomously to induce phenotypic switching from contractile to synthetic, proinflammatory SMCs. The IL11-treated vascular SMCs increased expression of ECM genes and increased mRNA for *IL6* and *CCL2* (among other inflammatory mediators) (76). Similarly, IL1β activates IL1 receptors (which are expressed in HISMCs, Figure 4B, Supplemental Table 3), triggering nuclear localization of NFκB (as we show in Figure 3D, E). Nuclear NFκB characteristically induces transcription of *ICAM1*, *CCL2* (also called *MCP1*), and *IL6*. This suggests cell-autonomous effects of IL1β, produced in response to pathologic mechanical stress, could trigger many of the loading-induced changes in HISMC gene expression (77). NFkB also mediates SMC phenotypic switching to a synthetic state (8, 40) by sequestering myocardin and preventing SRF-dependent expression of SMC contractile genes (78, 79). Consistent with our data, prior studies show static stretch (18%) increases SMC expression of *iNOS*, *IL6*, and *MCP1* within 3 hours, and that bowel proximal to obstruction markedly elevates PTGS2 (COX-2) after 24-48 hours (4, 11, 55). In addition, colon manipulation *in vivo* increases IL1β within 24 hours (11). Collectively, these studies strongly support the hypothesis that “mechano-transcription” powerfully modulates gene expression in bowel smooth muscle and highlights the complex self-reinforcing networks that induce SMC phenotypic switching (2). Our data demonstrates rapid changes in gene expression in human visceral SMCs towards a synthetic phenotype in response to pathologic mechanical stress, strengthening evidence for the role of mechanotransduction in bowel function that was previously identified in rodent models. These observations may have clinical implications for ileus, as well as for mechanical and functional bowel obstruction.

### Mechanical stress alters expression of many TGFβ family members in HISMC

Many differentially expressed genes in loaded HISMCs encode TGFβ superfamily proteins or components of their signaling pathways (including *BMP2, BMP4, BMP6, GREM1, Noggin, BAMBI, INHBA, TGFB1, TGFBR1, ALK2 (*reported as *ACVR1* in Supplemental Table 3), and *ALK4 (*reported as *ACVR1B* in Supplemental Table 3)). *TGFβ1* mRNA was increased 1.58-fold in loaded HISMCs, and receptors *TGFBR1* and *TGFBR3* mRNA increased ∼1.9 fold in response to loading. Elevated TGFβ1 signaling also serves as one possible explanation for the increase in IL11 noted above. However, our analysis showed equivalent levels of SMAD2/3 in the nucleus of loaded and unloaded HISMCs, indicating no increase in TGF receptor signaling at this early (6 hour) timepoint.

We were also intrigued by the changes in BMP family mRNA because bowel smooth muscle patterning depends on the interplay of BMP2, BMP4, and BMP7, as elegantly shown by Huycke *et al* (80). Furthermore, BMP2 increases vascular SMC migration and expression of synthetic markers (81, 82), and has anti-proliferative effects in pulmonary artery SMC (83). In addition, BMP2 counteracts many effects of TGFβ1 in SMCs by inducing PPARγ (84). While these observations are intriguing, our gene expression data (Figure 5A, Supplemental Table 3) showed increased *BMP2* and reduced *BAMBI* (BMP antagonist) mRNA in loaded HISMC. Based on prior data, these changes are expected to increase BMP receptor signaling. However, our analyses also showed reduced BMP4, and elevated *GREM1* and *NOG* (BMP antagonists) in loaded HISMCs, which would be expected to reduce BMP receptor signaling. To make sense of these observations, we looked for evidence of BMP signaling in HISMCs and found equivalent nuclear localization of SMAD1/5/8 in loaded and unloaded cells (Figure 5F), indicating no change in BMP receptor signaling. Collectively, these data suggest that TGFβ superfamily signaling changes rapidly in response to mechanical stress in HISMCs, but that these signaling systems were not (at least at this time point) affecting SMC cell phenotype, or that there is additional complexity to TGFβ signaling involving other bowel cell types that is not captured by our simplified system.

### Stress alters expression of axon guidance molecules and genes needed for cell-cell and cell-extracellular matrix (ECM) interactions

Many differentially expressed genes in loaded HISMCs encode axon guidance molecules (ephrins, netrins, semaphorins, and slits) that could influence bowel muscle innervation. Some encoded proteins also directly impact SMC biology, at least in the vasculature. However, whether they are specifically related to a contractile or synthetic phenotype is not well-understood. For example, ephrin B2 (*EFNB2*, increased 2.22-fold in loaded HISMCs) enhances vascular SMC contraction strength (85), while ephrin A1 (*EFNA1*, reduced 2.77-fold in loaded HISMCs) reduces integrin-induced vascular SMC spreading and inhibits SMC proliferation (86, 87). As another example, *SEMA7A* (increased 2.31-fold in loaded HISMCs) expression in vascular SMC is increased by PDGF (increased 2.04-fold in loaded HISMCs), and appears to be required for PDGF-induced vascular SMC proliferation and migration (88). In addition to these guidance molecules, which are not well studied in SMCs, loading changed expression of many cytoskeletal proteins (or regulators like *FMN1*), integrins, cadherins, focal adhesions, potentially altering SMC interactions with nearby cells and with the ECM.

### Pathologic stress-induced changes in HISMC gene expression could broadly affect the biology of many bowel cell types

Although our studies employed purified human intestinal smooth muscle in culture, HISMCs *in vivo* closely interact with many other cell types including enteric neurons, glia, muscularis macrophages, fibroblasts, and vascular SMCs. HISMCs also interact very closely with interstitial cells of Cajal (ICC) and PDGFRα+ cells to form the “SIP syncytium”, a network connected to SMCs by gap junctions (89–91). Recognizing that many differentially expressed genes induced by loading in HISMCs encode secreted or extracellular ligands, we employed NicheNet to try to unravel potential SMC-“niche” interactions that might occur in response to pathologic stretching. The resulting analyses (Figure 6-7) suggest that mechano-transcription responses to pathologic mechanical stress in HISMCs induce production of many growth factors (*EREG, AREG, HBEGF, FGF5, FGF7, FGF18, NRG1, PDGF, LIF*), cytokines (*IL6, IL11, CSF3, CLCF1*), and differentiation regulators (*WNT5A, BMP2, BMP6, DKK1, TGFB1, JAG1, DLL4, INHBB, INHBA*) that are likely to act on adjacent cells. For simplicity, our presented analyses include only the “Top 20” differentially expressed ligands in loaded versus unloaded HISMCs, based on the NicheNet model. Thus, these analyses show only a subset of the mechanotransduction-induced changes in HISMC gene expression. The interactions emphasize how physical stress experienced by HISMCs in a variety of disease contexts could remodel not only smooth muscle, but influence many other bowel cell types. These complex interactions may critically underlie some aspects of bowel dysfunction, especially for people with dysmotility or partial obstruction. In particular, the broad array of cellular changes predicted to occur in response to HISMC differential gene expression after mechanical stress might explain why recovery after bowel injury may be gradual (over days or months).

## Conclusions

We have presented some of the first and most detailed analyses of gene expression data in HISMCs showing that pathological mechanical stress, even over a short time scale, leads to a switch towards a synthetic, pro-inflammatory HISMC phenotype. Our novel NicheNet analysis has generated new, testable hypotheses regarding the interplay between visceral smooth muscle and other bowel cell types that may occur in response to pathologic mechanical stress. These interactions may govern how bowel function is altered over long periods of time in human bowel diseases in which such stresses are a significant part of disease pathophysiology.

## Abbreviations

HISMC: human intestinal smooth muscle cell
SMC: smooth muscle cell
PCL: Poly(ε-calprolactone)
ECM: extracellular matrix
GSEA: Gene Set Enrichment Analysis
LR: ligand-receptor
STRING: Search Tool for the Retrieval of Interacting Genes/Proteins
KEGG: Kyoto Encyclopedia of Genes and Genomes
GO: Gene Ontology,

