## Supplemental Figures 1-4 and Supplemental Tables 1-2 for "Rapid cyclic stretching induces a synthetic, proinflammatory phenotype in cultured human intestinal smooth muscle, with the potential to alter signaling to adjacent bowel cells"

### Supplemental Materials

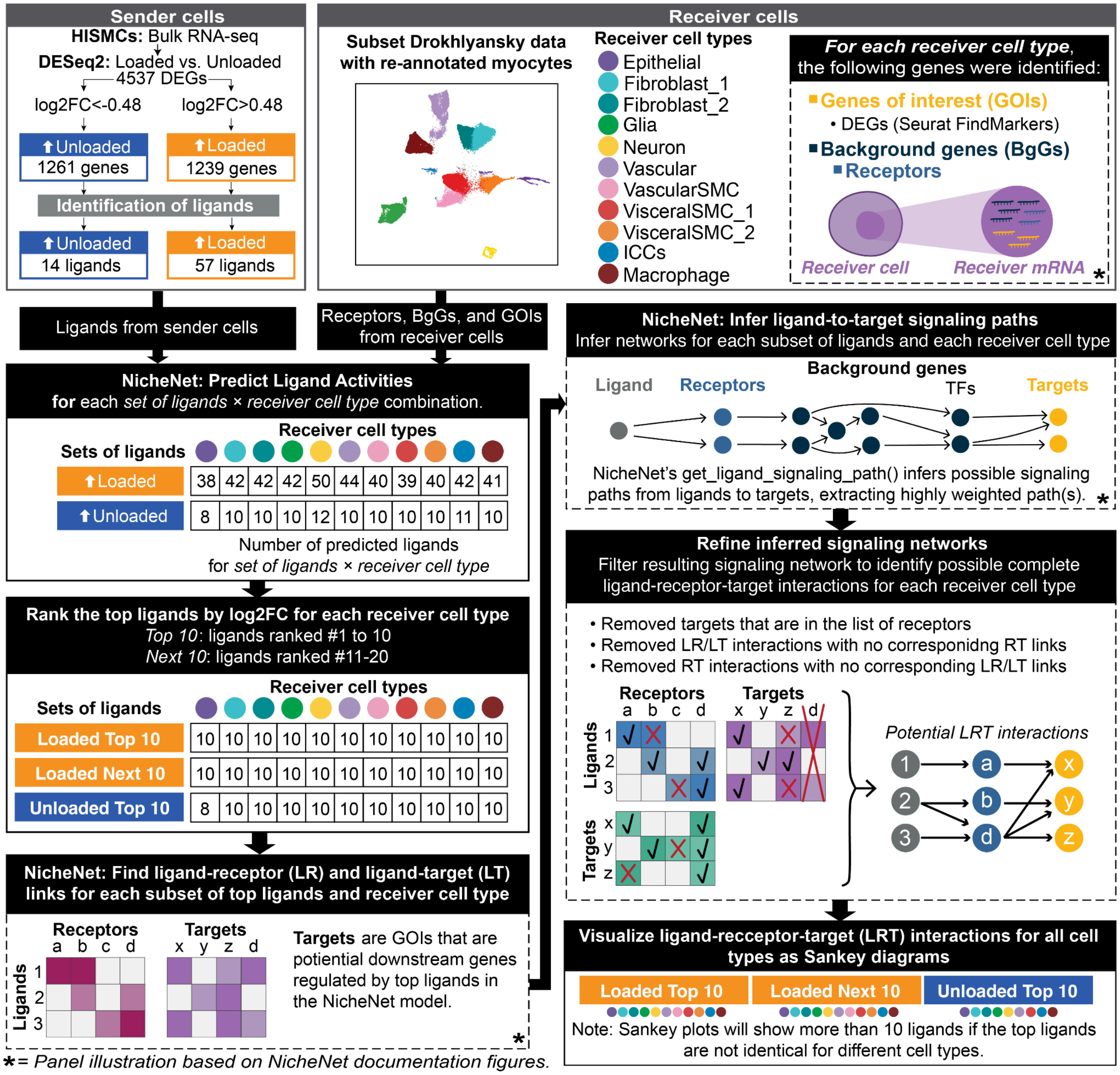

**Supplemental Figure 1.**
Schematic overview of the NicheNet analysis methods, including the process for filtering the inferred signaling network to keep only the potential interactions with ligand-receptor-target paths.

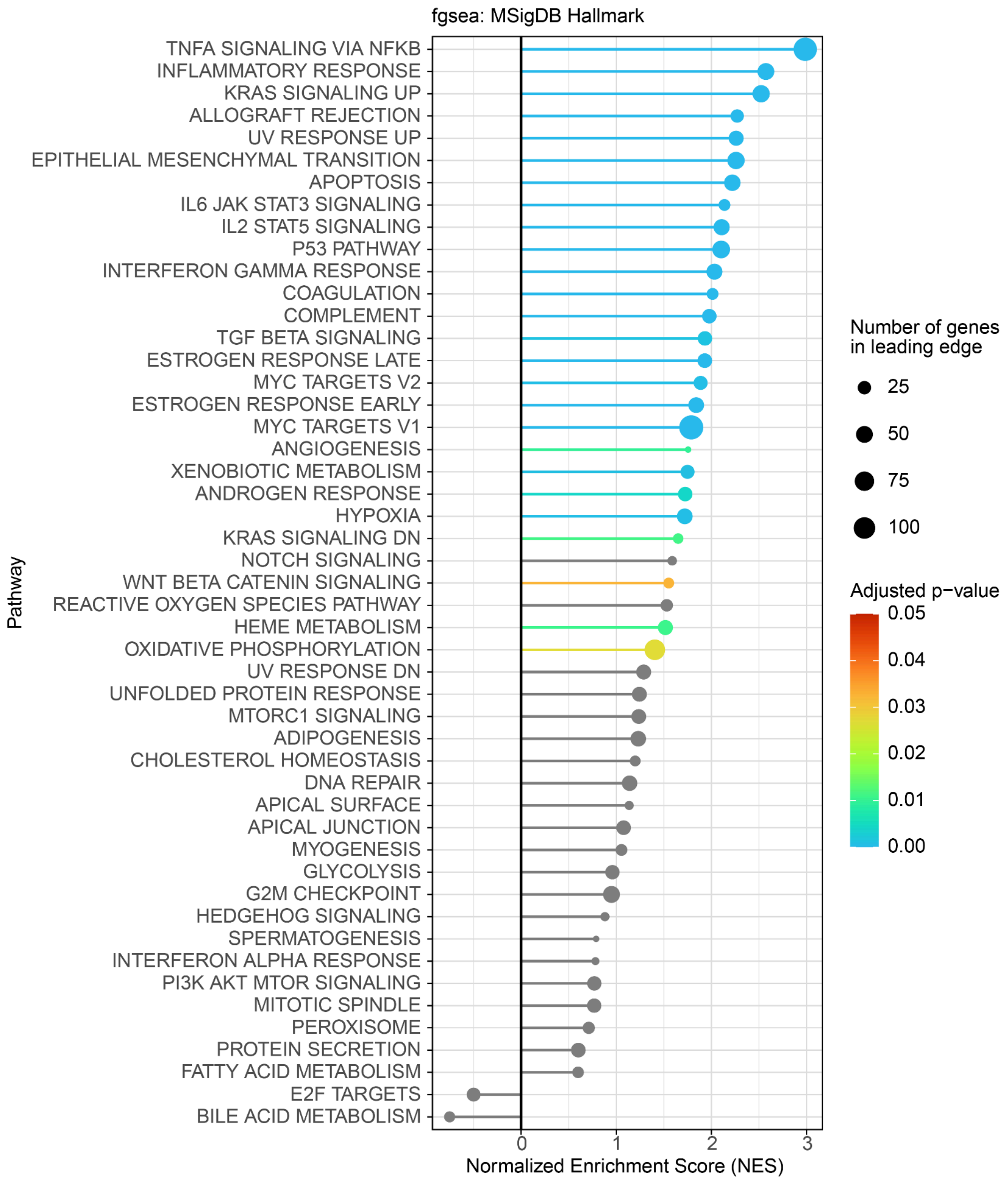

**Supplemental Figure 2.**

Gene set enrichment (fgsea) results for MSigDB Hallmark Pathways, using the differentially expressed genes between loaded and unloaded HISMCs.

PSTAT3 positive control

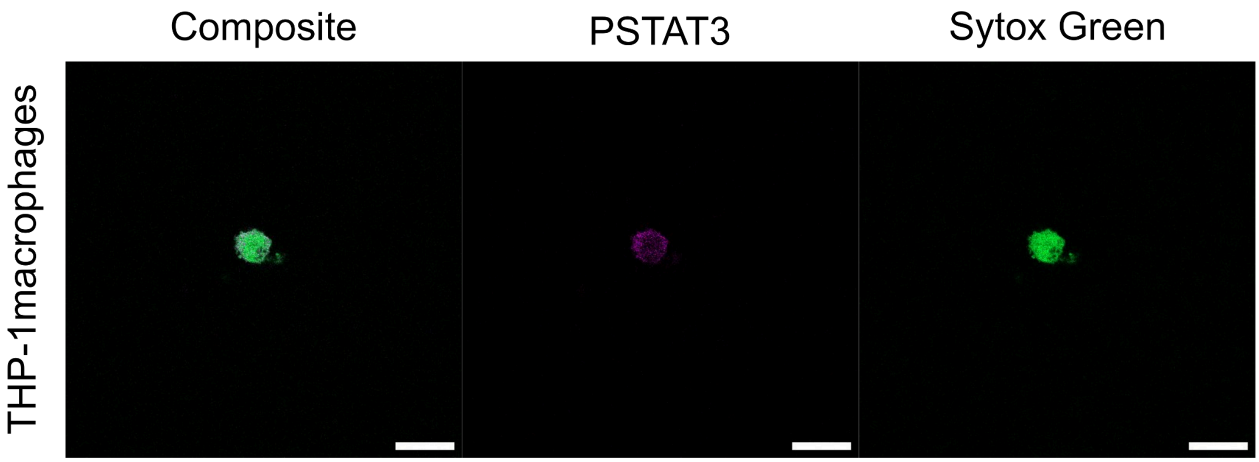

**Supplemental Figure 3.**
The presence of phospho-STAT3 (PSTAT3) staining was confirmed in THP-1 macrophages.

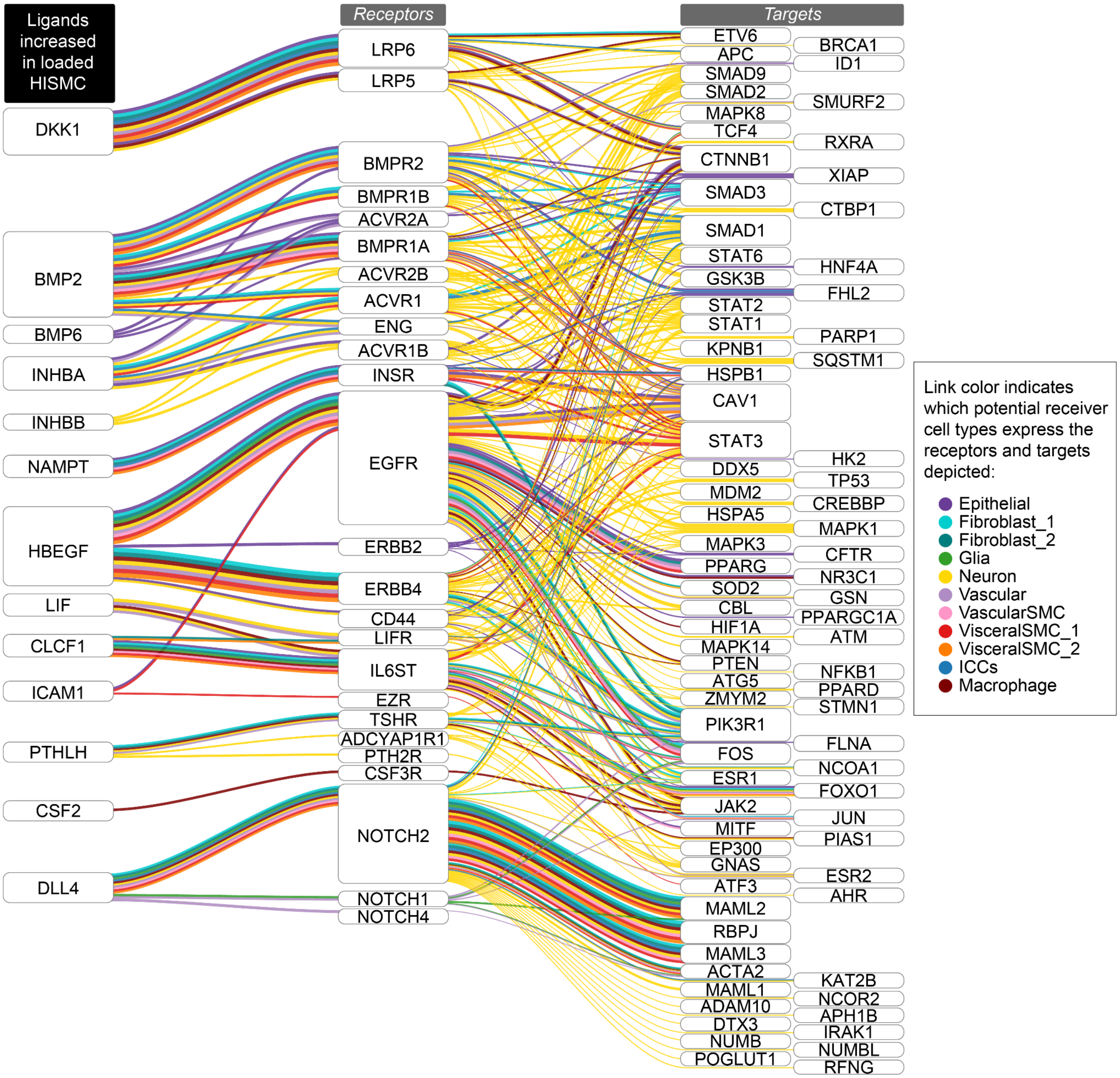

**Supplemental Figure 4.**

Sankey plot showing potential ligand-receptor-target links based on NicheNet's inferred signaling paths from the "next 10" ligands upregulated in the loaded HISMC to Drokhlyansky *et al.* receiver cell targets. The NicheNet prioritized ligand analysis between secreted ligands that are more abundant in loaded compared to unloaded HISMCs (left column) and receptors (middle column) and target genes (right column) in re-annotated Drokhlyansky *et al.* receiver cell types was used to infer signaling paths from each ligand to target. Potential ligand-receptor-target links were determined based on the inferred signaling paths from NicheNet. See Methods and Supplemental Figure 1 for additional details on the NicheNet analysis and process for inferring ligand-receptor-targets paths.

**Supplemental Table 1. Primers**

| Primer Name | Sequence (5' to 3') | Tm (˚C) | GC (%) |
|---|---|---|---|
| *YWHAZ* Fwd | TGCTTGCATCCCACAGACTA | 57 | 50 |
| *YWHAZ* Rev | AGGCAGACAATGACAGACCA | 57 | 50 |
| *ACTG2* Fwd | CATGTACGTCGCCATTCAAGC | 58 | 52 |
| *ACTG2* Rev | TTGATGTCTCGCACAATTTCTCT | 56 | 39 |
| *ACTA2* Fwd | CACTGTCAGGAATCCTGTGA | 55 | 50 |
| *ACTA2* Rev | CAAAGCCGGCCTTACAGA | 56 | 56 |
| *MYH11* Fwd | AGATGGTTCTGAGGAGGAAACG | 58 | 50 |
| *MYH11* Rev | AAAACTGTAGAAAGTTGCTTATTCACT | 55 | 33 |
| *IL6* Fwd | GGAGACTTGCCTGGTGAAA | 62 | 53 |
| *IL6* Rev | CTGGCTTGTTCCTCACTACTC | 62 | 50 |
| *IL11* Fwd | GAGAGGCTTGCTTGGGATATAG | 62 | 50 |
| *IL11* Rev | TCCCAAAGTGCCAGGATTAC | 62 | 50 |
| *GREM1* Fwd | GCAGGATAGTGGAGTGAGAAAG | 62 | 50 |
| *GREM1* Rev | TCAGCCTGTGTTCTGGTATTG | 62 | 48 |
| *BMP2* Fwd | GAGAAGGAGGAGGCAAAGAAA | 62 | 48 |
| *BMP2* Rev | GGGACACGTCCATTGAAAGA | 62 | 50 |
| *BMP4* Fwd | GGAGATGGTAGTAGAGGGATGT | 62 | 50 |
| *BMP4* Rev | CGTGTGTGTGTGGTGTATGT | 62 | 50 |
| *COL1A1* Fwd | GAGGGCCAAGACGAAGACATC | 62 | 57 |
| *COL1A1* Rev | CAGATCACGTCATCGCACAAC | 62 | 52 |
| *FN1* Fwd | CGGTGGCTGTCAGTCAAAG | 61 | 58 |
| *FN1* Rev | AAACCTCGGCTTCCTCCATAA | 61 | 48 |
| *VIM* Fwd | GCCCTAGACGAACTGGGTC | 61 | 63 |
| *VIM* Rev | GGCTGCAACTGCCTAATGAG | 61 | 55 |
| *MMP14* Fwd | GGCTACAGCAATATGGCTACC | 60 | 52 |
| *MMP14* Rev | GATGGCCGCTGAGAGTGAC | 62 | 63 |

**Supplemental Table 2. Antibodies and staining reagents**

| **Antibodies (species)** | **Application (Concentration)** | **Manufacturer** | **Catalog#, RRID** |
|---|---|---|---|
| Anti-phosphoSmad1/Smad5/Smad8, phospho-specific (Ser463/465) (rabbit) | Immunofluorescence (1:500) | Millipore | AB3848, RRID:AB_177439 |
| Smad2 (pS465/pS467)/Smad3 (pS423/pS425) (mouse) | Immunofluorescence (1:500) | BD Phosflow | 562586, RRID:AB_11151915 |
| IL11 (rabbit) | Immunofluorescence (1:200) | Invitrogen | PA5-36544, RRID:AB_2553579 |
| NF- B p65 (L8F6) (mouse) | Immunofluorescence (1:1000) | Cell Signaling | 6956, RRID:AB_10828935 |
| Smooth Muscle Myosin Heavy Chain (SMMS-1) (mouse) | Immunofluorescence (1:100) | Abcam | ab106919, RRID:AB_10866244 |
| Phospho-Stat3 (Tyr705) (D3A7) (rabbit) | Immunofluorescence (1:200) | Cell Signaling | 9145, RRID:AB_2491009 |
| Megakaryoblastic Leukemia 2 (MKL2) (rabbit) | Immunofluorescence (1:500) | Novus Bio | NBP1-46209, NP_054767.3 |
| **Secondary Antibodies** | | | |
| Donkey anti-mouse 594 | Immunofluorescence (1:400) | Invitrogen | A21203, RRID AB_2535789 |
| Donkey anti-rabbit 647 | Immunofluorescence (1:400) | Invitrogen | A31573, RRID AB_2536183 |
| Donkey anti-mouse 647 | Immunofluorescence (1:400) | Invitrogen | A3157, RRID AB_162542 |
| Donkey anti-rabbit 594 | Immunofluorescence (1:400) | Invitrogen | A21207, RRID AB_141637 |
| Donkey anti-chicken 594 | Immunofluorescence (1:400) | Invitrogen | A11042, RRID AB_2534099 |
| **Other staining reagents** | | | |
| Sytox Green nucleic acid stain | Immunofluorescence (1:30000) | ThermoFisher | S7020, RRID: not available |
| Alex Fluor 647 Phalloidin | Immunofluorescence (1:1000) | Invitrogen | A22287, RRID:AB_2620155 |

**Supplemental Table 3.**

This is a separate file with all DESeq2 differentially expressed genes.
