## Supplemental Table 3 for "Rapid cyclic stretching induces a synthetic, proinflammatory phenotype in cultured human intestinal smooth muscle, with the potential to alter signaling to adjacent bowel cells"

| | baseMean | log2FoldChange | lfcSE | pvalue | padj |
|---|---|---|---|---|---|
| XIRP1 | 41.48479737 | 7.971767312 | 1.338386856 | 1.20E-13 | 3.49E-12 |
| C2CD4A | 7.128882743 | 7.06324326 | 2.628677061 | 2.52E-07 | 2.92E-06 |
| LINC00473 | 46.65275601 | 6.796660305 | 0.8868883571 | 1.89E-14 | 5.91E-13 |
| SOX8 | 31.63272367 | 6.200020713 | 0.9421351667 | 2.51E-11 | 5.44E-10 |
| CSF3 | 284.7162758 | 6.004553315 | 0.2841582633 | 4.79E-99 | 5.48E-96 |
| NR4A3 | 294.4054965 | 5.680510132 | 0.2725819045 | 1.57E-97 | 1.67E-94 |
| AREG | 3230.195865 | 5.641058786 | 0.1610612407 | 1.21E-269 | 6.00E-266 |
| IL8 | 25512.68222 | 5.546525006 | 0.1242335815 | 0.00E+00 | 0.00E+00 |
| EREG | 671.071877 | 5.525773217 | 0.2472393183 | 2.36E-111 | 3.91E-108 |
| CXCL3 | 657.5317827 | 5.4904313 | 0.1896554355 | 2.03E-185 | 6.04E-182 |
| IL11 | 14286.52736 | 5.431536417 | 0.1750618849 | 2.46E-212 | 9.16E-209 |
| IL1B | 692.3808087 | 5.351605145 | 0.5307469958 | 3.31E-25 | 2.55E-23 |
| CCL20 | 525.6090953 | 5.326816237 | 0.320475329 | 2.64E-63 | 1.01E-60 |
| NR4A1 | 2414.44978 | 5.278665538 | 0.1361471068 | 0.00E+00 | 0.00E+00 |
| NR4A2 | 196.2775435 | 5.063926871 | 0.2511841226 | 6.52E-91 | 5.11E-88 |
| PTGS2 | 1011.817229 | 4.99705239 | 0.304467319 | 7.40E-62 | 2.69E-59 |
| FGF18 | 12.8805114 | 4.825885181 | 0.9004086245 | 3.48E-08 | 4.75E-07 |
| EGR2 | 40.335613 | 4.698014399 | 0.6998538662 | 1.27E-12 | 3.26E-11 |
| DUSP2 | 258.3851235 | 4.625151665 | 0.3528547582 | 2.14E-40 | 3.62E-38 |
| TNFSF15 | 265.7682792 | 4.374989351 | 0.8096709901 | 2.71E-09 | 4.42E-08 |
| NFATC2 | 388.0092761 | 4.259624575 | 0.2066164749 | 2.30E-95 | 2.28E-92 |
| SERPINB2 | 101.1269639 | 4.199987005 | 0.4014039671 | 1.04E-26 | 9.03E-25 |
| CXCL2 | 242.0841677 | 4.124661381 | 0.194766193 | 1.73E-100 | 2.14E-97 |
| BCL2A1 | 21.3249923 | 4.077732184 | 0.6498041557 | 4.25E-11 | 8.88E-10 |
| FOSB | 294.8434439 | 4.062138187 | 0.2042254461 | 2.82E-89 | 2.10E-86 |
| SYT12 | 11.34413284 | 3.84060202 | 0.8711082553 | 1.07E-06 | 1.09E-05 |
| TREM1 | 23.38811735 | 3.812485685 | 0.5842764732 | 7.38E-12 | 1.72E-10 |
| IL6 | 433.590388 | 3.743381552 | 0.1844311722 | 6.38E-93 | 5.28E-90 |
| STC1 | 10381.16188 | 3.533296299 | 0.2349351538 | 2.30E-52 | 6.84E-50 |
| RRAD | 706.6766194 | 3.519314221 | 0.2277497326 | 5.07E-55 | 1.61E-52 |
| MICALCL | 63.05058428 | 3.482878138 | 0.3208474992 | 2.00E-28 | 1.97E-26 |
| NIPAL1 | 33.78523101 | 3.44662779 | 0.4040365365 | 1.65E-18 | 7.48E-17 |
| INHBB | 7.344517148 | 3.391198297 | 0.9859788667 | 5.42E-05 | 0.000387742754 |
| KCNF1 | 36.27605921 | 3.304510428 | 0.392649656 | 3.65E-18 | 1.59E-16 |
| CCDC173 | 16.8918931 | 3.295256954 | 0.5360115109 | 8.89E-11 | 1.78E-09 |
| C11orf96 | 268.2803271 | 3.248799651 | 0.8037222623 | 2.07E-06 | 1.98E-05 |
| FMN1 | 139.621821 | 3.175965815 | 0.2571258616 | 4.15E-36 | 6.12E-34 |
| C8orf4 | 894.4131107 | 3.161624568 | 0.1971109256 | 3.80E-59 | 1.32E-56 |
| KBTBD8 | 699.6390491 | 3.151030929 | 0.1857286906 | 1.01E-65 | 4.31E-63 |
| MAP3K8 | 179.3225894 | 3.146495089 | 0.1817282872 | 3.18E-68 | 1.43E-65 |

| | baseMean | log2FoldChange | lfcSE | pvalue | padj |
|---|---|---|---|---|---|
| LIF | 2298.861771 | 3.121975892 | 0.1667765477 | 2.04E-79 | 1.17E-76 |
| IL33 | 22.04901559 | 2.971009165 | 0.6072437054 | 6.42E-08 | 8.37E-07 |
| JARID2 | 1934.340971 | 2.967116527 | 0.1095568384 | 8.32E-163 | 2.06E-159 |
| GCH1 | 177.8404711 | 2.95884923 | 0.3943363768 | 3.31E-15 | 1.10E-13 |
| LIMS3 | 142.0533158 | 2.956394272 | 1.24696725 | 0.000448008331 | 0.00250108588 |
| SPHK1 | 2694.914285 | 2.916558893 | 0.226292203 | 3.06E-39 | 4.80E-37 |
| MUM1L1 | 6.785288932 | 2.915152051 | 0.9450424314 | 1.47E-04 | 0.000939438427 |
| TNFAIP3 | 1480.278851 | 2.885504418 | 0.1993423343 | 1.01E-48 | 2.60E-46 |
| FAM83F | 8.572984949 | 2.870449087 | 0.7949647669 | 2.26E-05 | 0.000176386095 |
| GPRIN3 | 41.74476647 | 2.862731181 | 0.4917569293 | 3.21E-10 | 5.84E-09 |
| IL24 | 117.1674514 | 2.861343158 | 0.2677818679 | 9.53E-28 | 9.04E-26 |
| GEM | 2358.615323 | 2.820474238 | 0.1421388974 | 8.37E-89 | 5.66E-86 |
| MAK | 10.40610924 | 2.811190428 | 0.7040310719 | 4.48E-06 | 4.03E-05 |
| RNA18S5 | 91028.00999 | 2.80877284 | 0.2577643131 | 6.99E-29 | 7.23E-27 |
| GPR183 | 17.88446516 | 2.781580272 | 0.5463935407 | 2.32E-08 | 3.27E-07 |
| MAFF | 2860.095565 | 2.777076261 | 0.1117343346 | 1.62E-137 | 3.45E-134 |
| PRDM1 | 377.5820772 | 2.760862673 | 0.1959381925 | 2.69E-46 | 5.89E-44 |
| ANGPTL4 | 4310.117277 | 2.758839141 | 0.260901172 | 2.23E-27 | 2.02E-25 |
| LOC102724428 | 181.5103335 | 2.727098022 | 0.3012828986 | 9.50E-21 | 4.96E-19 |
| EGR1 | 345.200878 | 2.706792651 | 0.2742855841 | 3.17E-24 | 2.31E-22 |
| LONRF2 | 23.29090745 | 2.699608583 | 0.504381057 | 5.63E-09 | 8.73E-08 |
| DLL4 | 2272.600948 | 2.648488535 | 0.1779860249 | 2.73E-51 | 7.97E-49 |
| CHMP1B | 6171.174095 | 2.648084973 | 0.1146614225 | 3.07E-119 | 5.71E-116 |
| G0S2 | 3406.609109 | 2.636485707 | 0.1961917422 | 1.54E-42 | 2.87E-40 |
| SLC16A6 | 605.4298321 | 2.622135516 | 0.2570349307 | 1.18E-25 | 9.18E-24 |
| SH3RF3-AS1 | 89.55942277 | 2.615596546 | 0.3016799642 | 2.81E-19 | 1.33E-17 |
| BMP2 | 688.3709832 | 2.595977191 | 0.1762616005 | 2.43E-50 | 6.83E-48 |
| AMZ1 | 8.157870679 | 2.586707515 | 0.96395719 | 0.000343303023 | 0.001970485244 |
| SLC19A2 | 1664.027814 | 2.581330664 | 0.1456898431 | 2.00E-71 | 9.61E-69 |
| SRSF12 | 49.99280896 | 2.578678563 | 0.3827190317 | 9.29E-13 | 2.43E-11 |
| TMEM217 | 44.81663554 | 2.572681066 | 0.313961899 | 1.92E-17 | 7.83E-16 |
| RAMP1 | 333.4205007 | 2.543965277 | 0.2623373105 | 1.73E-23 | 1.15E-21 |
| BACH2 | 43.65654354 | 2.525531616 | 0.4110021271 | 4.64E-11 | 9.62E-10 |
| LFNG | 85.50459766 | 2.510697964 | 0.346898785 | 2.73E-14 | 8.48E-13 |
| INHBA | 3559.395838 | 2.482852869 | 0.2103625471 | 2.71E-33 | 3.57E-31 |
| PTHLH | 14.96621546 | 2.477382497 | 0.5402549164 | 2.97E-07 | 3.40E-06 |
| CXCL1 | 1823.532654 | 2.423032187 | 0.2885424722 | 2.15E-18 | 9.61E-17 |
| ITPRIP | 4060.894052 | 2.417615749 | 0.167654204 | 2.41E-48 | 5.89E-46 |
| HBEGF | 561.0605132 | 2.414440782 | 0.2659284032 | 6.55E-21 | 3.52E-19 |
| CSF2 | 30.6040662 | 2.411115039 | 0.6711459631 | 1.56E-05 | 1.27E-04 |

| | baseMean | log2FoldChange | lfcSE | pvalue | padj |
|---|---|---|---|---|---|
| LOC101928172 | 104.8020325 | 2.394197459 | 0.245868668 | 1.33E-23 | 9.09E-22 |
| DKK1 | 2040.7415 | 2.392392199 | 0.2503267214 | 7.28E-23 | 4.53E-21 |
| DENND2C | 127.4079473 | 2.382400667 | 0.2399619931 | 2.04E-24 | 1.51E-22 |
| HEYL | 7.460546729 | 2.368985875 | 1.007411603 | 0.000792807427 | 0.004117233966 |
| LRRC4 | 92.09546394 | 2.337389165 | 0.2460732198 | 1.58E-22 | 9.58E-21 |
| CBLN2 | 13.23918276 | 2.333904724 | 0.6436315193 | 1.70E-05 | 0.000136715976 |
| SIK1 | 189.9374267 | 2.328354202 | 0.2505123659 | 7.99E-22 | 4.56E-20 |
| MFSD2A | 448.3082838 | 2.311231032 | 0.2748873548 | 2.57E-18 | 1.13E-16 |
| LOC100996761 | 12.61636798 | 2.308954158 | 0.66602946 | 2.89E-05 | 2.21E-04 |
| SPSB1 | 1899.845342 | 2.288396334 | 0.1419360738 | 1.15E-59 | 4.08E-57 |
| SOWAHC | 794.2759879 | 2.263446876 | 0.113882201 | 4.80E-89 | 3.40E-86 |
| RGCC | 359.7382237 | 2.255861061 | 0.2180220078 | 2.79E-26 | 2.32E-24 |
| RASD1 | 52.76531821 | 2.255667814 | 0.2885003937 | 3.63E-16 | 1.32E-14 |
| CCNA1 | 22.33257701 | 2.23908863 | 0.4128932724 | 4.22E-09 | 6.69E-08 |
| IRAK2 | 1285.707447 | 2.237083089 | 0.1290458635 | 1.79E-68 | 8.32E-66 |
| SPRY4 | 2681.243535 | 2.235591965 | 0.1879213952 | 7.40E-34 | 1.00E-31 |
| LONRF3 | 16.31510236 | 2.232652328 | 0.4890038643 | 3.40E-07 | 3.87E-06 |
| SPRY2 | 3969.827656 | 2.232511422 | 0.2451613532 | 5.25E-21 | 2.85E-19 |
| ATF3 | 717.1998536 | 2.226189044 | 0.1543779091 | 2.45E-48 | 5.89E-46 |
| MEDAG | 189.4581466 | 2.217229154 | 0.2325742858 | 8.88E-23 | 5.46E-21 |
| FLJ31104 | 15.92969804 | 2.211884801 | 0.4923191579 | 5.04E-07 | 5.52E-06 |
| RNF152 | 661.587991 | 2.207393599 | 0.2373156038 | 8.69E-22 | 4.94E-20 |
| FKBP1A-SDCBP2 | 25.1836459 | 2.191340323 | 0.452418456 | 7.60E-08 | 9.76E-07 |
| TCF7 | 101.834316 | 2.182397607 | 0.2993058512 | 1.81E-14 | 5.68E-13 |
| NEXN-AS1 | 179.1267438 | 2.177749367 | 0.1614151658 | 1.35E-42 | 2.54E-40 |
| NIPAL4 | 83.26138426 | 2.177738291 | 0.4981996267 | 6.20E-07 | 6.66E-06 |
| IER3 | 3784.150809 | 2.161284418 | 0.1196738778 | 4.54E-74 | 2.33E-71 |
| FGL2 | 18.04098729 | 2.158832223 | 0.5174036589 | 1.82E-06 | 1.77E-05 |
| FZD8 | 608.1401055 | 2.144100827 | 0.135832194 | 3.05E-57 | 1.01E-54 |
| GPR3 | 152.7635592 | 2.140337867 | 0.1918422695 | 5.19E-30 | 5.77E-28 |
| MSC | 1328.932461 | 2.136021348 | 0.2040653832 | 7.51E-27 | 6.58E-25 |
| SPATA2L | 377.6897435 | 2.123740658 | 0.2165869107 | 6.62E-24 | 4.63E-22 |
| KLF4 | 211.1116113 | 2.114386924 | 0.2373262371 | 3.26E-20 | 1.65E-18 |
| CAMKK1 | 776.3288933 | 2.093101187 | 0.1019504349 | 8.83E-95 | 8.22E-92 |
| GREM1 | 12869.82147 | 2.085273509 | 0.09478917743 | 1.63E-108 | 2.43E-105 |
| FOS | 262.3151096 | 2.083768249 | 0.2375125787 | 1.07E-19 | 5.16E-18 |
| C5orf49 | 11.21456717 | 2.077836025 | 0.6359928879 | 5.92E-05 | 0.000420065383 |
| PTPRE | 343.191738 | 2.07205939 | 0.2276293465 | 5.54E-21 | 2.99E-19 |
| TFPI2 | 3620.895622 | 2.059448606 | 0.1251150903 | 4.70E-62 | 1.75E-59 |
| CXCL5 | 846.6128496 | 2.053891675 | 0.1930061965 | 1.23E-27 | 1.14E-25 |

| | baseMean | log2FoldChange | lfcSE | pvalue | padj |
|---|---|---|---|---|---|
| CSRNP1 | 1159.804877 | 2.052053352 | 0.1229961944 | 1.23E-63 | 4.81E-61 |
| SHISA2 | 110.0143975 | 2.051790988 | 0.2304369918 | 3.43E-20 | 1.73E-18 |
| GJA3 | 19.67462629 | 2.010609445 | 0.4654089614 | 9.54E-07 | 9.77E-06 |
| C3orf52 | 724.4847283 | 2.006126627 | 0.2806209837 | 5.42E-14 | 1.64E-12 |
| SAT1 | 2880.342435 | 1.983242457 | 0.2257196854 | 9.14E-20 | 4.43E-18 |
| MAP2K3 | 1502.85979 | 1.978398022 | 0.143512474 | 2.15E-44 | 4.38E-42 |
| KDM7A | 814.8106805 | 1.972529509 | 0.1717806695 | 1.05E-31 | 1.26E-29 |
| CLCF1 | 463.2035522 | 1.968261168 | 0.1345395059 | 1.19E-49 | 3.17E-47 |
| RASD2 | 36.99654406 | 1.964974879 | 0.4237161903 | 2.05E-07 | 2.42E-06 |
| KLF5 | 239.5907914 | 1.947440753 | 0.1366490286 | 3.14E-47 | 7.20E-45 |
| HIC1 | 633.3207181 | 1.944834304 | 0.2040708961 | 1.02E-22 | 6.23E-21 |
| CTGF | 6773.809611 | 1.941102879 | 0.198784235 | 7.54E-24 | 5.22E-22 |
| HIVEP3 | 62.62242444 | 1.937708313 | 0.4378644548 | 5.45E-07 | 5.95E-06 |
| IL1RL1 | 17.53687909 | 1.931480921 | 0.5305668842 | 1.57E-05 | 0.000127484259 |
| AVPI1 | 812.922406 | 1.930946224 | 0.1333907551 | 1.16E-48 | 2.94E-46 |
| KIAA0247 | 852.9756413 | 1.918665292 | 0.210395316 | 4.93E-21 | 2.69E-19 |
| DUSP6 | 4922.892239 | 1.914889024 | 0.1430929871 | 5.27E-42 | 9.57E-40 |
| TPPP | 30.32899123 | 1.894637151 | 0.3344527001 | 1.05E-09 | 1.80E-08 |
| MAFK | 976.8957555 | 1.88955242 | 0.1462164238 | 2.40E-39 | 3.84E-37 |
| ENTPD7 | 2292.421431 | 1.888573751 | 0.1393927412 | 5.54E-43 | 1.07E-40 |
| MYC | 4878.774889 | 1.878270884 | 0.1037826659 | 2.34E-74 | 1.29E-71 |
| ADORA2A | 20.03322863 | 1.870107624 | 0.4927033725 | 8.86E-06 | 7.56E-05 |
| GATA2 | 106.8047263 | 1.866400396 | 0.217834945 | 7.05E-19 | 3.26E-17 |
| NAMPT | 11165.20824 | 1.863179828 | 0.207884969 | 1.56E-20 | 8.12E-19 |
| PTGER3 | 154.5300284 | 1.858401395 | 0.2954191096 | 1.97E-11 | 4.31E-10 |
| FAM83G | 881.649128 | 1.857806522 | 0.1446963848 | 7.19E-39 | 1.11E-36 |
| DIRAS3 | 32.26924094 | 1.850258753 | 0.3158203933 | 3.42E-10 | 6.19E-09 |
| NKX3-1 | 210.9332657 | 1.84615862 | 0.1875741682 | 5.37E-24 | 3.81E-22 |
| EMR2 | 51.66292384 | 1.834382275 | 0.2540395695 | 3.97E-14 | 1.22E-12 |
| LOC100288175 | 180.8118828 | 1.831325928 | 0.2929605302 | 2.67E-11 | 5.76E-10 |
| ICAM1 | 1808.730768 | 1.82632917 | 0.2961000024 | 4.31E-11 | 8.98E-10 |
| ZC3H12C | 1876.50252 | 1.818571575 | 0.1378646618 | 7.21E-41 | 1.25E-38 |
| PMAIP1 | 2815.841051 | 1.815817781 | 0.1637766922 | 1.04E-29 | 1.13E-27 |
| RELB | 906.2805955 | 1.814235316 | 0.09444991359 | 2.38E-83 | 1.48E-80 |
| CCDC71L | 1939.80148 | 1.793983518 | 0.1767685194 | 2.31E-25 | 1.79E-23 |
| CD274 | 572.7238392 | 1.790072962 | 0.5213221178 | 3.00E-05 | 0.000227714845 |
| REL | 194.4610927 | 1.790039429 | 0.1968080093 | 6.71E-21 | 3.58E-19 |
| GJB2 | 40.39348964 | 1.76615392 | 0.3552399218 | 4.51E-08 | 6.07E-07 |
| FAM167A | 1976.167562 | 1.761606494 | 0.2139676542 | 1.24E-17 | 5.21E-16 |
| SMOX | 1546.441367 | 1.761370579 | 0.1504580515 | 8.24E-33 | 1.06E-30 |

| | baseMean | log2FoldChange | lfcSE | pvalue | padj |
|---|---|---|---|---|---|
| FOXO1 | 49.85955624 | 1.760322966 | 0.3634395329 | 7.85E-08 | 1.01E-06 |
| DACT1 | 918.4054002 | 1.759297501 | 0.2868424629 | 5.48E-11 | 1.13E-09 |
| FAM196A | 69.85888808 | 1.758123636 | 0.2667817296 | 3.03E-12 | 7.39E-11 |
| LOC101928841 | 157.0616838 | 1.746771074 | 0.2904062734 | 1.19E-10 | 2.33E-09 |
| ZC3H12A | 828.3120053 | 1.744227246 | 0.1542194337 | 8.23E-31 | 9.50E-29 |
| KCNJ15 | 176.7960319 | 1.740736382 | 0.1914367968 | 7.40E-21 | 3.92E-19 |
| HOTAIRM1 | 115.5913032 | 1.726925076 | 0.1946620914 | 5.78E-20 | 2.83E-18 |
| RND3 | 10261.68163 | 1.725072085 | 0.1073602607 | 7.16E-59 | 2.42E-56 |
| TSC22D1 | 5491.69044 | 1.722071633 | 0.08942737548 | 9.38E-84 | 6.07E-81 |
| BDKRB1 | 1339.502649 | 1.721325494 | 0.1481332029 | 2.33E-32 | 2.94E-30 |
| HCAR3 | 10.77567074 | 1.71996669 | 0.8962326588 | 0.001972501043 | 0.009068823473 |
| PITX3 | 7.215329314 | 1.711149281 | 0.7757538211 | 0.001340138481 | 0.006520693414 |
| MED12L | 34.71676924 | 1.710497624 | 0.3823129191 | 4.84E-07 | 5.32E-06 |
| ID4 | 18.25798587 | 1.710404725 | 0.4675356262 | 1.50E-05 | 0.000122681936 |
| SLC22A4 | 97.67393533 | 1.696368895 | 0.2063178741 | 1.55E-17 | 6.43E-16 |
| RGS16 | 12.91161925 | 1.693306443 | 0.5130752555 | 6.39E-05 | 0.000449156793 |
| NAB1 | 1715.610102 | 1.687886534 | 0.0933335325 | 2.98E-74 | 1.58E-71 |
| RLTPR | 95.33913833 | 1.687422999 | 0.1901624097 | 5.35E-20 | 2.64E-18 |
| TM4SF1 | 8339.035269 | 1.680812291 | 0.1316833975 | 1.94E-38 | 2.98E-36 |
| TBX2 | 3208.205113 | 1.680697975 | 0.1779387269 | 2.48E-22 | 1.48E-20 |
| AC105053.3 | 8.964833756 | 1.680069586 | 0.7922740283 | 0.001517701183 | 0.00726126379 |
| FOXL1 | 2787.658809 | 1.665595289 | 0.1267994715 | 1.52E-40 | 2.61E-38 |
| SLC30A1 | 728.1555217 | 1.665587055 | 0.1080888126 | 1.08E-54 | 3.35E-52 |
| BMP6 | 404.3133271 | 1.660376178 | 0.1654418081 | 7.92E-25 | 5.93E-23 |
| CMKLR1 | 136.8138551 | 1.66028512 | 0.4366623207 | 8.00E-06 | 6.88E-05 |
| ABHD17B | 810.7278073 | 1.654224031 | 0.09234910975 | 7.61E-73 | 3.78E-70 |
| CREB5 | 272.8839415 | 1.64999975 | 0.2322018101 | 8.43E-14 | 2.48E-12 |
| MMP1 | 1526.244455 | 1.64992204 | 0.2484793713 | 2.15E-12 | 5.34E-11 |
| SCXB | 30.15950963 | 1.639655756 | 0.4558001115 | 1.87E-05 | 1.49E-04 |
| ISG20 | 152.4721982 | 1.633706279 | 0.2331434174 | 1.66E-13 | 4.75E-12 |
| NFKB1 | 3291.740203 | 1.627912986 | 0.1590212507 | 9.69E-26 | 7.63E-24 |
| FAM196B | 981.0750629 | 1.617431428 | 0.2650862265 | 7.28E-11 | 1.48E-09 |
| RGS1 | 24.38421142 | 1.612987609 | 0.3722530813 | 1.04E-06 | 1.06E-05 |
| RELT | 671.9766656 | 1.599157273 | 0.153664074 | 1.80E-26 | 1.54E-24 |
| NFATC1 | 138.4409028 | 1.593040019 | 0.2486010821 | 1.06E-11 | 2.40E-10 |
| PIM3 | 926.2162293 | 1.588506121 | 0.1220105063 | 6.28E-40 | 1.05E-37 |
| PMEPA1 | 448.4795453 | 1.584839684 | 0.1543403916 | 7.59E-26 | 6.01E-24 |
| TAGLN3 | 208.1396073 | 1.5757728 | 0.2463190102 | 1.13E-11 | 2.55E-10 |
| MCL1 | 16801.29572 | 1.571351275 | 0.09405706382 | 9.19E-64 | 3.70E-61 |
| GPR68 | 919.4497003 | 1.571108213 | 0.2265079574 | 2.86E-13 | 8.02E-12 |

| | baseMean | log2FoldChange | lfcSE | pvalue | padj |
|---|---|---|---|---|---|
| CITED4 | 152.9689287 | 1.566896438 | 0.1607191097 | 1.53E-23 | 1.03E-21 |
| ARID3B | 264.3707713 | 1.563215751 | 0.1663399065 | 4.42E-22 | 2.60E-20 |
| PDE3A | 65.05377928 | 1.554585666 | 0.4312878083 | 1.79E-05 | 0.000143759969 |
| JUNB | 1332.143853 | 1.542534709 | 0.3155755714 | 5.91E-08 | 7.77E-07 |
| LOC101929475 | 20.96447101 | 1.540472785 | 0.4276790042 | 2.10E-05 | 0.000165588482 |
| MMP3 | 2201.210486 | 1.539529101 | 0.2297507919 | 1.61E-12 | 4.10E-11 |
| JAG1 | 888.0131531 | 1.531904748 | 0.1306016507 | 6.94E-33 | 8.99E-31 |
| TBX3 | 1372.399969 | 1.515194768 | 0.1241396873 | 2.52E-35 | 3.64E-33 |
| ZNF697 | 1221.08729 | 1.513292938 | 0.1568578223 | 3.89E-23 | 2.49E-21 |
| MARS2 | 916.3014092 | 1.50958472 | 0.1834869059 | 1.47E-17 | 6.11E-16 |
| PELI1 | 405.2620287 | 1.509000023 | 0.1848069825 | 2.41E-17 | 9.70E-16 |
| CHAC1 | 7856.109889 | 1.505593895 | 0.1298018073 | 3.22E-32 | 4.00E-30 |
| WNT16 | 88.19553196 | 1.499310812 | 0.2589392362 | 5.12E-10 | 9.09E-09 |
| PER2 | 275.6154262 | 1.498946815 | 0.1245841259 | 1.88E-34 | 2.64E-32 |
| PDE4D | 2114.480501 | 1.493658685 | 0.08835815528 | 3.29E-65 | 1.36E-62 |
| ZBTB46 | 306.5848733 | 1.492041635 | 0.2427855315 | 5.82E-11 | 1.19E-09 |
| NFKBIA | 4872.245293 | 1.49153635 | 0.09505708042 | 1.50E-56 | 4.85E-54 |
| CASS4 | 22.69034999 | 1.487640297 | 0.4040833203 | 1.59E-05 | 0.000129143991 |
| GPR137C | 74.61962388 | 1.48702025 | 0.2550997299 | 4.03E-10 | 7.23E-09 |
| EMP1 | 8334.040697 | 1.486507328 | 0.1826304241 | 2.68E-17 | 1.08E-15 |
| PECAM1 | 8.921078935 | 1.480881548 | 0.9203095337 | 0.003821596587 | 0.0159202491 |
| SCG2 | 20.65622405 | 1.474889874 | 0.4200865926 | 3.05E-05 | 0.000231020707 |
| YRDC | 1576.977252 | 1.468143558 | 0.1574692764 | 9.30E-22 | 5.26E-20 |
| HOXD1 | 10.15651083 | 1.466796368 | 0.7503434549 | 0.002275365449 | 0.01022266632 |
| CIRBP-AS1 | 16.39353898 | 1.465136794 | 0.564386192 | 0.000498243069 | 0.002742317176 |
| LINC00673 | 71.39959959 | 1.460969756 | 0.3360335863 | 9.37E-07 | 9.65E-06 |
| RHOB | 3200.715756 | 1.452116814 | 0.1025780823 | 1.41E-46 | 3.14E-44 |
| MEF2D | 806.5450452 | 1.445381136 | 0.1185468509 | 2.59E-35 | 3.70E-33 |
| RIPK2 | 1967.907295 | 1.437089513 | 0.1451327221 | 3.50E-24 | 2.54E-22 |
| TAF4B | 358.9037182 | 1.436285969 | 0.200021607 | 5.47E-14 | 1.65E-12 |
| ZSWIM6 | 1242.676193 | 1.434972811 | 0.1263010148 | 5.63E-31 | 6.65E-29 |
| RNA28S5 | 848535.6969 | 1.424371375 | 0.1856887423 | 2.26E-15 | 7.69E-14 |
| SRXN1 | 7027.826382 | 1.420606512 | 0.1091821935 | 1.41E-39 | 2.30E-37 |
| PPARGC1B | 113.5153846 | 1.419136877 | 0.169848573 | 5.60E-18 | 2.41E-16 |
| F2RL3 | 36.03203569 | 1.418864593 | 0.452037884 | 1.02E-04 | 6.84E-04 |
| BCOR | 948.4050493 | 1.417633071 | 0.125609662 | 1.25E-30 | 1.44E-28 |
| TGIF1 | 3089.795249 | 1.41731142 | 0.06600116373 | 2.30E-103 | 3.11E-100 |
| AJAP1 | 294.6862529 | 1.409591447 | 0.2615458427 | 5.32E-09 | 8.30E-08 |
| NR3C1 | 4437.283902 | 1.408172275 | 0.104947147 | 4.00E-42 | 7.35E-40 |
| PALM2 | 188.4948865 | 1.40492043 | 0.2008178299 | 2.16E-13 | 6.08E-12 |

| | baseMean | log2FoldChange | lfcSE | pvalue | padj |
|---|---|---|---|---|---|
| ELL | 694.6519894 | 1.403358403 | 0.1418646627 | 3.57E-24 | 2.58E-22 |
| C10orf67 | 18.67811973 | 1.402283134 | 0.4854654898 | 0.000240206507 | 0.001445456879 |
| KLHL21 | 1546.299719 | 1.399306486 | 0.1077423802 | 1.25E-39 | 2.07E-37 |
| GAB2 | 574.1552198 | 1.391852438 | 0.1001375032 | 5.68E-45 | 1.17E-42 |
| TMEM200A | 776.3773961 | 1.39113863 | 0.2065258745 | 1.27E-12 | 3.26E-11 |
| CXCL6 | 141.0060231 | 1.385617545 | 0.2944594792 | 1.80E-07 | 2.15E-06 |
| RNVU1-6 | 87.74737038 | 1.382683493 | 0.2499417796 | 2.38E-09 | 3.92E-08 |
| ABHD17C | 827.7700218 | 1.366143941 | 0.1503713792 | 8.63E-21 | 4.54E-19 |
| ARHGAP31 | 1748.399097 | 1.365345601 | 0.1317528057 | 3.30E-26 | 2.71E-24 |
| FOXC2 | 63.01187612 | 1.364609625 | 0.3099252744 | 8.00E-07 | 8.38E-06 |
| FST | 2303.267052 | 1.362162324 | 0.2052768601 | 2.47E-12 | 6.10E-11 |
| HTR7 | 262.494486 | 1.359396675 | 0.2384798707 | 9.46E-10 | 1.63E-08 |
| NFKBIZ | 1051.319454 | 1.357968405 | 0.1765555779 | 1.20E-15 | 4.16E-14 |
| HIVEP2 | 2620.066418 | 1.354649932 | 0.1613515716 | 3.84E-18 | 1.67E-16 |
| TJP2 | 2859.429371 | 1.354127895 | 0.1688930826 | 8.91E-17 | 3.39E-15 |
| KBTBD2 | 2181.358919 | 1.352826238 | 0.1128722732 | 3.53E-34 | 4.82E-32 |
| ZNF474 | 20.28050917 | 1.352553775 | 0.3759884425 | 2.40E-05 | 0.000186668998 |
| TMEM158 | 1885.03758 | 1.35082122 | 0.1850678225 | 2.37E-14 | 7.38E-13 |
| ZNF281 | 2654.378476 | 1.344140129 | 0.1427521505 | 4.15E-22 | 2.45E-20 |
| SOD2 | 5590.163097 | 1.339523599 | 0.2321046075 | 6.19E-10 | 1.09E-08 |
| ETS1 | 5880.660819 | 1.339204994 | 0.1386740156 | 4.01E-23 | 2.55E-21 |
| TDG | 1331.166586 | 1.33815008 | 0.1112063438 | 2.38E-34 | 3.29E-32 |
| PDE4B | 350.64886 | 1.335332087 | 0.2361518254 | 1.22E-09 | 2.07E-08 |
| PHLDA1 | 9189.910243 | 1.334538191 | 0.09075098996 | 6.33E-50 | 1.71E-47 |
| CA2 | 10.43345907 | 1.330115119 | 1.250282075 | 0.006980786208 | 0.02611702826 |
| NAB2 | 2673.977679 | 1.326884057 | 0.1741052957 | 2.10E-15 | 7.16E-14 |
| SNAI1 | 184.0432219 | 1.326291355 | 0.1533553639 | 4.52E-19 | 2.10E-17 |
| AKAP12 | 16672.32973 | 1.325135571 | 0.1145151418 | 3.71E-32 | 4.57E-30 |
| IER2 | 669.7334829 | 1.323134124 | 0.1201591755 | 3.08E-29 | 3.23E-27 |
| JUND | 888.046238 | 1.321755347 | 0.1179702465 | 3.26E-30 | 3.70E-28 |
| CHIC2 | 943.2565958 | 1.319243381 | 0.1180219032 | 4.65E-30 | 5.20E-28 |
| MUC12 | 87.7123027 | 1.31820287 | 0.2426485124 | 4.39E-09 | 6.94E-08 |
| DYRK3 | 911.2322296 | 1.31513527 | 0.09094796021 | 1.99E-48 | 4.95E-46 |
| SYTL3 | 211.9341325 | 1.311995503 | 0.1620912342 | 5.16E-17 | 2.02E-15 |
| TMEM155 | 15.05125371 | 1.310531943 | 0.4852681648 | 0.000459347075 | 0.002548144043 |
| TRMT10C | 1415.892533 | 1.30564646 | 0.1220008955 | 6.87E-28 | 6.55E-26 |
| SHC4 | 327.6691293 | 1.305318738 | 0.1354726014 | 4.84E-23 | 3.05E-21 |
| TNIP3 | 20.59906501 | 1.304543409 | 0.4270083883 | 0.000149907084 | 0.000958336873 |
| FGF2 | 2218.244428 | 1.3037576 | 0.1716259117 | 2.59E-15 | 8.77E-14 |
| IL13RA2 | 64.93436322 | 1.300146868 | 0.4239138484 | 0.000139501958 | 0.000899153532 |

| | baseMean | log2FoldChange | lfcSE | pvalue | padj |
|---|---|---|---|---|---|
| RCAN1 | 3928.342015 | 1.295919447 | 0.124550916 | 2.09E-26 | 1.76E-24 |
| SNRK | 1672.582867 | 1.295875572 | 0.08793635293 | 3.46E-50 | 9.54E-48 |
| OTUD4 | 2205.955328 | 1.293384009 | 0.166494494 | 6.89E-16 | 2.45E-14 |
| C12orf44 | 3602.458831 | 1.293129299 | 0.0635053181 | 3.38E-93 | 2.96E-90 |
| ST7-AS1 | 67.04706824 | 1.289673619 | 0.2126593844 | 1.12E-10 | 2.19E-09 |
| TMC6 | 11.27739444 | 1.286861322 | 0.5576490647 | 0.001289409127 | 0.006304142851 |
| AHR | 5677.67847 | 1.286459128 | 0.08823630712 | 2.33E-49 | 6.08E-47 |
| KCNK2 | 39.60286518 | 1.283467897 | 0.3249828881 | 6.06E-06 | 5.31E-05 |
| CD83 | 122.4614146 | 1.282361418 | 0.1730492521 | 1.16E-14 | 3.71E-13 |
| CEBPB | 1245.662903 | 1.28049309 | 0.1853596254 | 4.17E-13 | 1.13E-11 |
| SERPINB8 | 986.7203209 | 1.279975142 | 0.1640025467 | 5.23E-16 | 1.88E-14 |
| DUSP5 | 2695.356026 | 1.278875016 | 0.2825676442 | 4.66E-07 | 5.13E-06 |
| HMOX1 | 1260.412146 | 1.278436416 | 0.1985995159 | 1.03E-11 | 2.35E-10 |
| ADAMTS4 | 130.0073825 | 1.277155978 | 0.2017032906 | 2.07E-11 | 4.53E-10 |
| PITPNM3 | 46.39064913 | 1.274600954 | 0.3151097759 | 4.13E-06 | 3.74E-05 |
| PLIN2 | 11789.74528 | 1.270419128 | 0.1197131288 | 2.35E-27 | 2.12E-25 |
| LRRC8C | 2401.849033 | 1.268657988 | 0.2009159147 | 2.25E-11 | 4.90E-10 |
| STARD8 | 128.331757 | 1.267320483 | 0.218716883 | 5.92E-10 | 1.04E-08 |
| RPSAP52 | 318.218253 | 1.265623109 | 0.3139255534 | 4.20E-06 | 3.80E-05 |
| SH3TC1 | 129.1321624 | 1.263084377 | 0.3054779968 | 2.66E-06 | 2.50E-05 |
| OSGIN2 | 1131.475119 | 1.261308511 | 0.1452865064 | 3.52E-19 | 1.65E-17 |
| UBALD1 | 247.0312599 | 1.258381424 | 0.1874665412 | 1.67E-12 | 4.24E-11 |
| NFKBIE | 837.6754298 | 1.258193102 | 0.1496664796 | 3.89E-18 | 1.69E-16 |
| ETS2 | 9015.417075 | 1.257276983 | 0.09206306599 | 1.76E-43 | 3.49E-41 |
| MAMLD1 | 98.07996698 | 1.255506889 | 0.2429699673 | 1.99E-08 | 2.83E-07 |
| PXDC1 | 1738.920031 | 1.251086778 | 0.1281266264 | 1.47E-23 | 9.93E-22 |
| COQ10B | 1453.298897 | 1.247132611 | 0.08779948474 | 4.60E-47 | 1.04E-44 |
| DGKD | 1300.704902 | 1.240143154 | 0.09314531384 | 1.87E-41 | 3.31E-39 |
| ZNF597 | 147.5837287 | 1.235814578 | 0.2054707452 | 1.60E-10 | 3.08E-09 |
| STK38L | 1145.260175 | 1.235746056 | 0.1793958093 | 5.08E-13 | 1.37E-11 |
| PER1 | 979.7950853 | 1.232938793 | 0.1296345585 | 1.71E-22 | 1.03E-20 |
| PLAUR | 6150.625858 | 1.223812705 | 0.1926710842 | 1.78E-11 | 3.92E-10 |
| KLF3 | 1515.214711 | 1.220258429 | 0.08531297341 | 2.05E-47 | 4.76E-45 |
| IL27RA | 311.8042815 | 1.218068069 | 0.1762766845 | 4.33E-13 | 1.18E-11 |
| ARID5A | 336.6386316 | 1.216389089 | 0.1233326329 | 5.86E-24 | 4.12E-22 |
| DYRK2 | 1020.554067 | 1.215909762 | 0.1652793048 | 1.74E-14 | 5.48E-13 |
| ZSWIM4 | 285.6051599 | 1.21400784 | 0.1883110658 | 1.04E-11 | 2.36E-10 |
| SEMA7A | 953.3322717 | 1.211159912 | 0.240526987 | 4.07E-08 | 5.50E-07 |
| STK17A | 2601.948998 | 1.210338403 | 0.1916458207 | 2.41E-11 | 5.24E-10 |
| TACSTD2 | 25.3104498 | 1.208645719 | 0.413299951 | 0.000250280265 | 0.001497758389 |

| | baseMean | log2FoldChange | lfcSE | pvalue | padj |
|---|---|---|---|---|---|
| CDK17 | 2787.703517 | 1.208209858 | 0.1785343175 | 1.19E-12 | 3.07E-11 |
| RASSF8 | 1842.970772 | 1.205200319 | 0.08602987591 | 1.32E-45 | 2.81E-43 |
| MAML3 | 81.56729839 | 1.203467734 | 0.2765584913 | 1.13E-06 | 1.14E-05 |
| VEGFC | 1466.992745 | 1.20294892 | 0.1979587375 | 1.09E-10 | 2.14E-09 |
| EHD1 | 3014.090854 | 1.202549894 | 0.1059906711 | 7.58E-31 | 8.89E-29 |
| ZNF35 | 728.5352616 | 1.194573275 | 0.1067789288 | 4.03E-30 | 4.54E-28 |
| TNFAIP6 | 96.95332587 | 1.186232441 | 0.2370876614 | 5.06E-08 | 6.73E-07 |
| NOG | 144.6665989 | 1.185043859 | 0.2214858948 | 8.00E-09 | 1.22E-07 |
| FOXD1 | 435.225198 | 1.184099703 | 0.232575262 | 3.13E-08 | 4.30E-07 |
| KLF9 | 2930.057127 | 1.180246984 | 0.1932362545 | 9.41E-11 | 1.87E-09 |
| CRY1 | 1173.64258 | 1.177921321 | 0.09773672043 | 1.98E-34 | 2.75E-32 |
| SKIL | 1638.893949 | 1.177754744 | 0.1148443949 | 1.13E-25 | 8.84E-24 |
| CD55 | 1219.346208 | 1.174985255 | 0.2302072087 | 2.93E-08 | 4.04E-07 |
| GNAL | 119.4525446 | 1.174046409 | 0.2447560034 | 1.39E-07 | 1.71E-06 |
| PTPN1 | 5646.084667 | 1.170561637 | 0.1253277143 | 9.69E-22 | 5.46E-20 |
| METRNL | 543.3374405 | 1.169235054 | 0.1436769438 | 3.89E-17 | 1.55E-15 |
| KBTBD11 | 55.20871432 | 1.166459094 | 0.2873248621 | 4.15E-06 | 3.76E-05 |
| TTC32 | 82.36195427 | 1.157921144 | 0.1896444694 | 9.92E-11 | 1.96E-09 |
| FRAT2 | 245.5848466 | 1.15677189 | 0.1254379988 | 2.87E-21 | 1.58E-19 |
| CPEB4 | 825.4690398 | 1.155207876 | 0.1866278889 | 5.69E-11 | 1.17E-09 |
| NRG1 | 735.1283654 | 1.153154872 | 0.3100268452 | 1.61E-05 | 0.000130151829 |
| MMP10 | 68.46534847 | 1.153149117 | 0.2505762381 | 3.89E-07 | 4.35E-06 |
| KLF13 | 2981.345551 | 1.152962249 | 0.1039136192 | 1.31E-29 | 1.41E-27 |
| EFNB2 | 330.0921037 | 1.15251764 | 0.2859260699 | 4.69E-06 | 4.20E-05 |
| UCN2 | 81.7729422 | 1.1481407 | 0.2591140865 | 8.43E-07 | 8.78E-06 |
| CDC42SE1 | 5560.569852 | 1.146589288 | 0.07470883945 | 3.79E-54 | 1.15E-51 |
| CCRN4L | 617.8390923 | 1.146558585 | 0.1681609617 | 8.95E-13 | 2.34E-11 |
| WWTR1 | 3072.562021 | 1.144929177 | 0.1148702628 | 2.19E-24 | 1.61E-22 |
| IGF1R | 931.9704471 | 1.140855348 | 0.08929944129 | 2.29E-38 | 3.48E-36 |
| CD69 | 128.2207915 | 1.139588339 | 0.3278151012 | 4.07E-05 | 0.000299364799 |
| CSGALNACT2 | 1893.847782 | 1.138054465 | 0.1420028728 | 1.10E-16 | 4.13E-15 |
| FEM1B | 3308.419065 | 1.137511347 | 0.1068347185 | 2.02E-27 | 1.85E-25 |
| HRH1 | 1732.779071 | 1.134876781 | 0.08326137161 | 2.78E-43 | 5.45E-41 |
| B3GNT5 | 2275.054821 | 1.134468828 | 0.2126329002 | 7.32E-09 | 1.12E-07 |
| SDE2 | 1589.211168 | 1.134312576 | 0.08779089211 | 2.38E-39 | 3.84E-37 |
| ABCG1 | 139.0867683 | 1.132554114 | 0.2154161204 | 1.34E-08 | 1.97E-07 |
| MESDC1 | 521.4387972 | 1.132106621 | 0.1489963804 | 3.05E-15 | 1.02E-13 |
| C5orf30 | 474.457615 | 1.131678669 | 0.1232587705 | 4.37E-21 | 2.40E-19 |
| ZHX2 | 130.5193751 | 1.131058712 | 0.2129664498 | 1.03E-08 | 1.54E-07 |
| SOX4 | 688.9254554 | 1.129772212 | 0.2175441709 | 1.91E-08 | 2.73E-07 |

| | baseMean | log2FoldChange | lfcSE | pvalue | padj |
|---|---|---|---|---|---|
| ZNRF3 | 301.0381032 | 1.128335858 | 0.1633393681 | 4.82E-13 | 1.30E-11 |
| TLE3 | 715.1249183 | 1.123574357 | 0.1387716241 | 5.66E-17 | 2.21E-15 |
| SRSF5 | 6532.574177 | 1.123022134 | 0.08154751237 | 4.00E-44 | 8.05E-42 |
| NPAS1 | 24.34967878 | 1.121750543 | 0.3959385019 | 0.000351275624 | 0.002013917126 |
| FBXO33 | 861.1361457 | 1.116274851 | 0.08792471245 | 6.69E-38 | 1.01E-35 |
| HSPB8 | 121.5779991 | 1.114564286 | 0.1967339985 | 1.40E-09 | 2.37E-08 |
| RBM24 | 138.4670157 | 1.112711666 | 0.2211763336 | 4.67E-08 | 6.26E-07 |
| SLC35G2 | 569.9450505 | 1.112093144 | 0.1293360247 | 8.44E-19 | 3.88E-17 |
| SLC2A14 | 97.32208554 | 1.110687242 | 0.1976854567 | 1.88E-09 | 3.13E-08 |
| PURB | 1337.59366 | 1.110475679 | 0.1656367555 | 2.08E-12 | 5.19E-11 |
| LOC101927086 | 72.42980039 | 1.110326508 | 0.2807785305 | 6.62E-06 | 5.77E-05 |
| PNP | 3948.964231 | 1.109562402 | 0.2320628622 | 1.62E-07 | 1.96E-06 |
| B3GNT2 | 550.7841912 | 1.108471772 | 0.1620825772 | 8.15E-13 | 2.14E-11 |
| MT2A | 22954.26516 | 1.108424326 | 0.1535593939 | 5.07E-14 | 1.54E-12 |
| FAM107B | 1667.132974 | 1.104182212 | 0.1668178744 | 3.57E-12 | 8.67E-11 |
| CCL2 | 148.1270982 | 1.103084851 | 0.4868053218 | 0.001492241739 | 0.007154902511 |
| AGPAT9 | 447.0195552 | 1.100832407 | 0.1300415134 | 2.66E-18 | 1.17E-16 |
| NIP7 | 3028.90357 | 1.100478358 | 0.1465860576 | 6.13E-15 | 1.99E-13 |
| HTR1B | 34.92050831 | 1.100404739 | 0.3099695241 | 3.36E-05 | 0.000251689365 |
| ZNF267 | 795.0303029 | 1.099513239 | 0.1434089563 | 1.84E-15 | 6.31E-14 |
| FASTKD5 | 1167.334213 | 1.099325951 | 0.101781416 | 3.74E-28 | 3.59E-26 |
| RGS3 | 2139.071021 | 1.09872531 | 0.08112830047 | 9.49E-43 | 1.81E-40 |
| VPS37B | 1083.484915 | 1.098372919 | 0.1086465122 | 5.41E-25 | 4.11E-23 |
| STX11 | 12.82482864 | 1.098171732 | 0.5521580044 | 0.002908367678 | 0.01262837164 |
| EVA1A | 1296.557226 | 1.0974154 | 0.2129262138 | 2.46E-08 | 3.44E-07 |
| ETV3 | 1595.599857 | 1.097118006 | 0.1027072621 | 1.35E-27 | 1.24E-25 |
| TRPC4 | 55.38925383 | 1.090266037 | 0.4107095562 | 0.000579161015 | 0.003125454278 |
| BTG3 | 1540.176275 | 1.085501634 | 0.1215307459 | 4.46E-20 | 2.23E-18 |
| AQP3 | 75.39349931 | 1.084670252 | 0.4610801536 | 0.00126103867 | 0.00618839972 |
| LRRC8A | 3536.833461 | 1.079631627 | 0.1394771349 | 1.35E-15 | 4.65E-14 |
| HAS2 | 2060.848336 | 1.078378174 | 0.1399864005 | 1.40E-15 | 4.82E-14 |
| LHFPL2 | 1115.369635 | 1.078158959 | 0.141553875 | 2.77E-15 | 9.28E-14 |
| SRF | 2016.664549 | 1.078074977 | 0.109708585 | 9.50E-24 | 6.52E-22 |
| CLP1 | 1182.754322 | 1.076420413 | 0.1101559941 | 1.63E-23 | 1.09E-21 |
| RARA | 493.1236481 | 1.074250286 | 0.1630089435 | 4.55E-12 | 1.09E-10 |
| BEND3 | 252.6416996 | 1.073820027 | 0.1787493913 | 1.96E-10 | 3.70E-09 |
| HIST2H3C | 10831.29051 | 1.071682434 | 0.2919132694 | 2.15E-05 | 0.000169049742 |
| POLR1C | 2005.673833 | 1.071682129 | 0.1331318249 | 8.99E-17 | 3.41E-15 |
| PLCL2 | 187.8289304 | 1.071636925 | 0.1405953511 | 2.68E-15 | 9.05E-14 |
| ABL2 | 5290.334107 | 1.068276895 | 0.1928092263 | 3.02E-09 | 4.89E-08 |

| | baseMean | log2FoldChange | lfcSE | pvalue | padj |
|---|---|---|---|---|---|
| KIF21B | 152.9676319 | 1.066729312 | 0.2511198282 | 2.07E-06 | 1.98E-05 |
| SH2B2 | 48.07282433 | 1.066603965 | 0.3597860542 | 0.000249589071 | 0.001494222634 |
| NR5A2 | 86.68079798 | 1.06625692 | 0.2638950625 | 5.07E-06 | 4.50E-05 |
| IPPK | 857.6535323 | 1.066141409 | 0.1318841181 | 6.82E-17 | 2.64E-15 |
| POU2F2 | 200.9818424 | 1.066098737 | 0.2571771054 | 3.24E-06 | 2.99E-05 |
| YOD1 | 599.0821401 | 1.064191688 | 0.1204198204 | 1.12E-19 | 5.36E-18 |
| LRCH1 | 1284.893232 | 1.06209076 | 0.1030650256 | 7.54E-26 | 6.01E-24 |
| LOC101928820 | 29.05324579 | 1.060080662 | 0.5061990645 | 0.002384653073 | 0.01063982607 |
| GRAMD1B | 65.35265132 | 1.059602841 | 0.256598997 | 3.42E-06 | 3.14E-05 |
| NFKB2 | 1139.798448 | 1.059108164 | 0.1070623054 | 5.06E-24 | 3.62E-22 |
| RGS4 | 997.2782963 | 1.058346543 | 0.3133322408 | 6.37E-05 | 0.000448011173 |
| AXIN1 | 1015.592181 | 1.0582408 | 0.118489698 | 4.73E-20 | 2.34E-18 |
| ST3GAL1 | 3988.513109 | 1.057998958 | 0.2606478019 | 4.63E-06 | 4.16E-05 |
| CSNK1E | 4710.76182 | 1.057809752 | 0.09624522109 | 4.66E-29 | 4.86E-27 |
| MCTP1 | 388.6575484 | 1.055473464 | 0.1080649574 | 1.78E-23 | 1.17E-21 |
| SAMD4A | 2726.173184 | 1.055051237 | 0.1509184261 | 2.98E-13 | 8.33E-12 |
| KLHL15 | 627.8030033 | 1.052787323 | 0.1293071736 | 4.33E-17 | 1.71E-15 |
| C12orf79 | 15.5216731 | 1.052344061 | 0.5310268914 | 0.003138903424 | 0.01346445782 |
| LOC102724611 | 58.92727492 | 1.051805651 | 0.3204315398 | 9.12E-05 | 0.000620019396 |
| LOC100505555 | 10.39193736 | 1.048414488 | 0.6993601424 | 0.006715766138 | 0.02534026227 |
| SHB | 1399.15145 | 1.046076811 | 0.1239371075 | 3.63E-18 | 1.59E-16 |
| UBALD2 | 724.8942413 | 1.045784983 | 0.1473143629 | 1.36E-13 | 3.92E-12 |
| F2RL1 | 318.3276742 | 1.045238438 | 0.1310825941 | 1.75E-16 | 6.50E-15 |
| DAPK3 | 2610.196915 | 1.044811436 | 0.08073531436 | 3.03E-39 | 4.80E-37 |
| SNIP1 | 996.9959657 | 1.042685829 | 0.09125117755 | 3.58E-31 | 4.26E-29 |
| GPR157 | 158.0140706 | 1.039141008 | 0.1798360729 | 8.22E-10 | 1.43E-08 |
| RAB43 | 728.7160982 | 1.031849843 | 0.2286863148 | 6.45E-07 | 6.90E-06 |
| SH3BP1 | 492.953827 | 1.031230545 | 0.1407834193 | 2.65E-14 | 8.24E-13 |
| C22orf31 | 8.408404488 | 1.029841862 | 1.628699814 | 0.01045119766 | 0.03663085733 |
| TRIB1 | 719.7127307 | 1.027732466 | 0.1237026401 | 1.12E-17 | 4.69E-16 |
| PDGFA | 198.707721 | 1.027168589 | 0.2097557169 | 1.01E-07 | 1.28E-06 |
| PFKFB3 | 2304.276151 | 1.026080897 | 0.1443787821 | 1.34E-13 | 3.87E-12 |
| SPATA13 | 1424.47409 | 1.025166984 | 0.1758554634 | 5.99E-10 | 1.06E-08 |
| SH2B3 | 2339.78915 | 1.021014038 | 0.1558249881 | 6.29E-12 | 1.49E-10 |
| CHD7 | 377.7066873 | 1.014103164 | 0.1775498947 | 1.24E-09 | 2.10E-08 |
| PPIF | 4577.376918 | 1.013114305 | 0.1482543095 | 9.33E-13 | 2.43E-11 |
| LOC102724169 | 49.25002041 | 1.011629586 | 0.2737365494 | 2.18E-05 | 1.71E-04 |
| IFFO2 | 745.2182856 | 1.008120219 | 0.1220837366 | 1.77E-17 | 7.22E-16 |
| BAIAP2 | 551.6797587 | 1.006081402 | 0.1364309288 | 1.89E-14 | 5.91E-13 |
| SLC2A3 | 5817.682648 | 1.004512188 | 0.1455151628 | 6.20E-13 | 1.65E-11 |

| | baseMean | log2FoldChange | lfcSE | pvalue | padj |
|---|---|---|---|---|---|
| SERTAD1 | 1122.186674 | 1.002932104 | 0.1243364121 | 8.43E-17 | 3.21E-15 |
| NCR3LG1 | 968.777943 | 1.002468208 | 0.1795028313 | 2.58E-09 | 4.24E-08 |
| SPRED2 | 608.8294455 | 1.000823664 | 0.1435647972 | 3.65E-13 | 1.01E-11 |
| BPGM | 1120.601056 | 0.9986518299 | 0.09556981958 | 1.79E-26 | 1.53E-24 |
| PRKCH | 68.45644848 | 0.9979682351 | 0.2404743097 | 3.54E-06 | 3.24E-05 |
| ADAP1 | 158.8748948 | 0.996314128 | 0.1814975443 | 4.55E-09 | 7.18E-08 |
| UTP3 | 1994.372721 | 0.996159505 | 0.1159133793 | 1.07E-18 | 4.87E-17 |
| SLC25A25 | 1214.145918 | 0.995926999 | 0.08966691304 | 1.38E-29 | 1.47E-27 |
| AKAP8 | 1450.94566 | 0.994029969 | 0.1004016151 | 5.15E-24 | 3.67E-22 |
| AMPD3 | 1410.994941 | 0.992582615 | 0.1426504879 | 4.01E-13 | 1.10E-11 |
| ERRFI1 | 10859.22845 | 0.9911159065 | 0.2103281448 | 2.90E-07 | 3.33E-06 |
| SH3RF1 | 1420.77938 | 0.9882823973 | 0.1527930962 | 1.15E-11 | 2.59E-10 |
| TMEM156 | 468.8054479 | 0.9875724073 | 0.1646609646 | 2.31E-10 | 4.32E-09 |
| REST | 3255.097252 | 0.9874145782 | 0.1658856627 | 3.00E-10 | 5.48E-09 |
| FRMD8 | 1018.324303 | 0.9863671619 | 0.1133766472 | 4.02E-19 | 1.88E-17 |
| CDCA4 | 1789.615088 | 0.9855756104 | 0.1487091622 | 4.03E-12 | 9.70E-11 |
| NR1D2 | 902.6528662 | 0.9811964608 | 0.1043262567 | 6.39E-22 | 3.72E-20 |
| NAF1 | 222.4081576 | 0.9810571845 | 0.1718269315 | 1.33E-09 | 2.24E-08 |
| PPAPDC1A | 143.7784428 | 0.9798021757 | 0.2188304378 | 8.25E-07 | 8.62E-06 |
| TXNRD1 | 57642.27951 | 0.9796224432 | 0.1020788425 | 1.55E-22 | 9.39E-21 |
| MIR155HG | 123.0889091 | 0.9793617162 | 0.3015771187 | 1.13E-04 | 7.52E-04 |
| GRASP | 161.387977 | 0.9785367622 | 0.3586590814 | 0.000553989835 | 0.003003770814 |
| ZNF584 | 506.5658266 | 0.9756919693 | 0.1241364122 | 4.75E-16 | 1.71E-14 |
| ELF1 | 1136.217788 | 0.9755084303 | 0.09643074128 | 5.69E-25 | 4.28E-23 |
| BAK1 | 1870.795011 | 0.9746972395 | 0.08228635305 | 2.92E-33 | 3.81E-31 |
| IRS2 | 297.952644 | 0.9734105335 | 0.1416358762 | 7.66E-13 | 2.02E-11 |
| PRRX2 | 156.3907211 | 0.972079364 | 0.2974910057 | 0.000105102320 | 0.000704895697 |
| ANKRD42 | 520.985039 | 0.9718584662 | 0.1153506543 | 4.36E-18 | 1.88E-16 |
| IFNAR2 | 331.6896003 | 0.9704386248 | 0.2098809257 | 4.18E-07 | 4.62E-06 |
| IL6R | 368.9407529 | 0.9691965657 | 0.2511760088 | 1.19E-05 | 9.82E-05 |
| TRAF4 | 1803.05038 | 0.9674257845 | 0.09111600599 | 3.11E-27 | 2.79E-25 |
| USP36 | 826.8604644 | 0.9668654739 | 0.1443035162 | 2.50E-12 | 6.16E-11 |
| GDF15 | 931.9495561 | 0.9646907764 | 0.1482535024 | 9.13E-12 | 2.10E-10 |
| MT1E | 5146.261418 | 0.9629220099 | 0.1239208258 | 9.11E-16 | 3.23E-14 |
| NXT1 | 1093.060914 | 0.9628930007 | 0.0998996858 | 6.93E-23 | 4.33E-21 |
| SLC35F2 | 1592.178013 | 0.9625327482 | 0.1995578419 | 1.60E-07 | 1.94E-06 |
| DUSP1 | 2485.263463 | 0.962251683 | 0.1100839001 | 2.88E-19 | 1.36E-17 |
| C10orf2 | 666.2307537 | 0.9621740087 | 0.1604505483 | 2.38E-10 | 4.44E-09 |
| CXXC5 | 1249.111683 | 0.9571195477 | 0.2452031334 | 1.01E-05 | 8.52E-05 |
| RCL1 | 1006.383218 | 0.9563484411 | 0.1387671861 | 6.72E-13 | 1.79E-11 |

| | baseMean | log2FoldChange | lfcSE | pvalue | padj |
|---|---|---|---|---|---|
| HES4 | 32.52085292 | 0.9548147758 | 0.3012436475 | 0.000156579253 | 0.000994585541 |
| ENC1 | 1942.871821 | 0.9521262633 | 0.1970546633 | 1.56E-07 | 1.89E-06 |
| SIX4 | 1200.480647 | 0.9508146745 | 0.1147063619 | 1.45E-17 | 6.04E-16 |
| IQCJ-SCHIP1 | 435.598181 | 0.949751965 | 0.1312690364 | 5.82E-14 | 1.74E-12 |
| ERF | 1410.654816 | 0.9496060592 | 0.1388802996 | 9.94E-13 | 2.57E-11 |
| TGFBR1 | 3083.343655 | 0.9484291012 | 0.1998822938 | 2.23E-07 | 2.61E-06 |
| KLF16 | 411.3640965 | 0.9479649906 | 0.1430018638 | 4.22E-12 | 1.01E-10 |
| CCNH | 1993.384398 | 0.9469131206 | 0.1327943258 | 1.24E-13 | 3.59E-12 |
| STK17B | 1764.365885 | 0.9438504546 | 0.1323946981 | 1.27E-13 | 3.66E-12 |
| HECA | 504.9919985 | 0.9431308897 | 0.1232179798 | 2.48E-15 | 8.42E-14 |
| CDR2 | 2006.592381 | 0.9429473804 | 0.1053903256 | 4.69E-20 | 2.33E-18 |
| C17orf96 | 75.34150963 | 0.941736589 | 0.2170231768 | 1.66E-06 | 1.63E-05 |
| F3 | 20558.35878 | 0.9401699969 | 0.187855836 | 6.81E-08 | 8.82E-07 |
| CH25H | 14.77410471 | 0.9389369554 | 0.5373450312 | 0.005605521552 | 0.02182547343 |
| KCNG1 | 1537.700237 | 0.9369470405 | 0.1650993654 | 1.73E-09 | 2.89E-08 |
| DDX28 | 690.6338991 | 0.9355742617 | 0.09040383753 | 5.44E-26 | 4.38E-24 |
| BIN3 | 1305.790936 | 0.9344380503 | 0.08019327132 | 3.01E-32 | 3.77E-30 |
| SMOC1 | 372.3348595 | 0.9342345212 | 0.3667822142 | 0.000975514578 | 0.004936926092 |
| AGO2 | 1457.492468 | 0.9335724494 | 0.1252923219 | 1.20E-14 | 3.82E-13 |
| BEND7 | 200.1801687 | 0.9332273172 | 0.1949118776 | 2.03E-07 | 2.40E-06 |
| TACC2 | 299.6807127 | 0.9327267936 | 0.1586300951 | 5.18E-10 | 9.20E-09 |
| TGFBR3 | 601.8558334 | 0.9313146893 | 0.12738848 | 3.37E-14 | 1.04E-12 |
| LOC101928319 | 7.528233801 | 0.9311145807 | 0.9292683636 | 0.01246510784 | 0.04235348941 |
| PAQR5 | 436.1726536 | 0.9290551017 | 0.1626261681 | 1.38E-09 | 2.33E-08 |
| PGBD5 | 13.15178568 | 0.9274940159 | 0.7296634881 | 0.010235095 | 0.03597505417 |
| RRP1 | 1603.841031 | 0.9268630374 | 0.1149767368 | 9.81E-17 | 3.68E-15 |
| OTUD1 | 133.0648481 | 0.9265629077 | 0.1682571657 | 4.46E-09 | 7.05E-08 |
| CCDC147-AS1 | 15.4643094 | 0.9260404705 | 0.4798016011 | 0.004266491988 | 0.01740855007 |
| ZNF212 | 320.0631538 | 0.9245233244 | 0.1166701125 | 3.04E-16 | 1.11E-14 |
| AKAP2 | 15248.75313 | 0.9232481188 | 0.1571395336 | 5.28E-10 | 9.37E-09 |
| SOX9 | 127.2930733 | 0.9214292918 | 0.2995922996 | 0.000219122665 | 0.001334908908 |
| LATS2 | 1448.222564 | 0.9204972369 | 0.1134392609 | 7.31E-17 | 2.81E-15 |
| THBS1 | 33392.96137 | 0.9194323948 | 0.271359519 | 4.37E-05 | 0.000318533353 |
| S1PR2 | 575.2443527 | 0.9186761739 | 0.2791371274 | 1.06E-04 | 0.000707551278 |
| LOC101929188 | 37.41794926 | 0.9170654965 | 0.3930651648 | 0.001749157046 | 0.008179396751 |
| RP11-132A1.4 | 93.1209634 | 0.9157205875 | 0.2000117971 | 5.79E-07 | 6.27E-06 |
| NEDD9 | 2027.514561 | 0.9139386659 | 0.1928104649 | 2.59E-07 | 2.99E-06 |
| NCOA7 | 823.0433511 | 0.913672472 | 0.1645300517 | 3.56E-09 | 5.69E-08 |
| FAM46A | 813.6726922 | 0.913053878 | 0.1009042973 | 1.94E-20 | 1.00E-18 |
| PLEKHO1 | 4519.007102 | 0.9127963511 | 0.09448767801 | 6.07E-23 | 3.81E-21 |

| | baseMean | log2FoldChange | lfcSE | pvalue | padj |
|---|---|---|---|---|---|
| CSGALNACT1 | 159.4967824 | 0.9102693463 | 0.1941801627 | 3.40E-07 | 3.87E-06 |
| ERN1 | 232.1136109 | 0.9061312863 | 0.1735821897 | 2.28E-08 | 3.22E-07 |
| EAF1 | 1071.943604 | 0.9049605899 | 0.0831776193 | 2.03E-28 | 1.98E-26 |
| ZNF800 | 901.4171818 | 0.9047348756 | 0.1479647686 | 1.27E-10 | 2.47E-09 |
| FRS2 | 884.6782272 | 0.9035012768 | 0.1110878073 | 5.75E-17 | 2.24E-15 |
| SMAD7 | 399.2074279 | 0.9007200642 | 0.2626270225 | 6.76E-05 | 0.000472302262 |
| PROB1 | 28.38958316 | 0.8999968909 | 0.321827973 | 0.000548256012 | 0.002978104259 |
| MAD2L1BP | 1501.319373 | 0.8986497763 | 0.08027319486 | 5.95E-30 | 6.57E-28 |
| UCK2 | 4484.896053 | 0.8978146649 | 0.1163215497 | 1.03E-15 | 3.61E-14 |
| ZNF229 | 1113.314184 | 0.8971953626 | 0.124675433 | 8.35E-14 | 2.46E-12 |
| BNC1 | 1416.209646 | 0.8964194141 | 0.1953292028 | 5.54E-07 | 6.03E-06 |
| MAP6D1 | 125.7413766 | 0.8962762592 | 0.182039391 | 1.08E-07 | 1.35E-06 |
| KTI12 | 685.9927814 | 0.8962204769 | 0.1028846465 | 4.23E-19 | 1.97E-17 |
| GNA13 | 3572.965903 | 0.8959947141 | 0.1065415798 | 5.64E-18 | 2.41E-16 |
| OGFRL1 | 5321.880159 | 0.8956628219 | 0.1680594609 | 1.27E-08 | 1.88E-07 |
| LIMS3L | 266.4687497 | 0.8947899233 | 0.2938736919 | 0.000249412586 | 0.001493766691 |
| USP12 | 1580.271647 | 0.8943805889 | 0.1188088173 | 7.24E-15 | 2.35E-13 |
| C10orf90 | 26.03953383 | 0.8937726199 | 0.3869391724 | 0.001955237398 | 0.009004494158 |
| MYADM | 1552.561168 | 0.8923792102 | 0.1089004105 | 3.94E-17 | 1.57E-15 |
| PPTC7 | 688.1103846 | 0.8909540559 | 0.1362547419 | 8.85E-12 | 2.04E-10 |
| CLDN4 | 42.98986319 | 0.8879893716 | 0.2732360155 | 0.000133171823 | 0.000865471531 |
| LUZP1 | 4983.738835 | 0.8865014475 | 0.124846908 | 1.80E-13 | 5.13E-12 |
| RAPH1 | 403.3040301 | 0.8864820517 | 0.2044167096 | 1.81E-06 | 1.76E-05 |
| BCAR3 | 2375.455197 | 0.8851501461 | 0.1436127426 | 9.58E-11 | 1.90E-09 |
| PPP1R15A | 2719.756409 | 0.882528824 | 0.1211869992 | 4.53E-14 | 1.38E-12 |
| RNF126 | 748.4887127 | 0.8822130187 | 0.1207350889 | 3.76E-14 | 1.15E-12 |
| TNFRSF12A | 8290.924991 | 0.8819175758 | 0.1527349969 | 1.05E-09 | 1.81E-08 |
| ZFP69B | 228.2071251 | 0.8811980129 | 0.1609687825 | 5.93E-09 | 9.17E-08 |
| BRPF1 | 1023.322507 | 0.8810748841 | 0.1063384433 | 1.67E-17 | 6.87E-16 |
| CSRP1 | 13112.5724 | 0.8748217263 | 0.1058398726 | 2.00E-17 | 8.10E-16 |
| PALM2-AKAP2 | 955.0028304 | 0.8745325787 | 0.2077654703 | 3.23E-06 | 2.98E-05 |
| FAM89A | 1155.193917 | 0.8744540637 | 0.1588510241 | 4.96E-09 | 7.77E-08 |
| PHC2 | 7167.042489 | 0.8742430529 | 0.1023428278 | 1.88E-18 | 8.42E-17 |
| SHROOM3 | 1303.197293 | 0.8737340844 | 0.20480249 | 2.52E-06 | 2.37E-05 |
| AMER1 | 515.127243 | 0.871725336 | 0.1213933153 | 9.78E-14 | 2.86E-12 |
| SPOCD1 | 1420.846372 | 0.8700152734 | 0.102074492 | 2.24E-18 | 1.00E-16 |
| YPEL5 | 2504.803234 | 0.8693341515 | 0.1456123261 | 3.26E-10 | 5.92E-09 |
| CBFB | 1836.216438 | 0.8687035793 | 0.09637030228 | 2.88E-20 | 1.46E-18 |
| RNF19B | 715.4792301 | 0.8684568894 | 0.09217704846 | 6.49E-22 | 3.76E-20 |
| FEM1A | 1864.207604 | 0.8679200166 | 0.08283316699 | 1.60E-26 | 1.38E-24 |

| | baseMean | log2FoldChange | lfcSE | pvalue | padj |
|---|---|---|---|---|---|
| ZNF850 | 300.8992469 | 0.8666133398 | 0.2074188602 | 3.78E-06 | 3.45E-05 |
| KCTD10 | 4382.521819 | 0.8660598621 | 0.08759695033 | 5.52E-24 | 3.89E-22 |
| UBAP1 | 2217.861108 | 0.8648929767 | 0.07891498114 | 8.86E-29 | 9.09E-27 |
| PTPRN | 117.8416129 | 0.8635804471 | 0.2159610266 | 8.03E-06 | 6.90E-05 |
| RNF25 | 1533.144823 | 0.863204053 | 0.07903974861 | 1.35E-28 | 1.36E-26 |
| NR1D1 | 32.81745552 | 0.8611387215 | 0.331518975 | 0.00101776906 | 0.005129845477 |
| GABPB1 | 1709.53375 | 0.8602443673 | 0.139381117 | 9.60E-11 | 1.90E-09 |
| ZSCAN5A | 133.1082853 | 0.8589416176 | 0.1836480433 | 3.89E-07 | 4.34E-06 |
| GTPBP4 | 6009.200293 | 0.8588959704 | 0.1560621075 | 5.15E-09 | 8.06E-08 |
| PTP4A1 | 2125.769013 | 0.855826334 | 0.1424131113 | 2.77E-10 | 5.11E-09 |
| DUSP7 | 564.2624373 | 0.8556609365 | 0.1433750139 | 3.39E-10 | 6.15E-09 |
| PPP1R15B | 3083.817196 | 0.8539309034 | 0.08658761432 | 8.97E-24 | 6.18E-22 |
| TNFAIP1 | 2399.307374 | 0.8525388332 | 0.1143875955 | 1.33E-14 | 4.21E-13 |
| TNFSF11 | 20.49379292 | 0.8523318745 | 0.5164634959 | 0.007688584205 | 0.02825156225 |
| LRFN5 | 78.6568672 | 0.8516595791 | 0.2637029522 | 0.000149687003 | 0.000957340977 |
| STX3 | 707.2587775 | 0.8503988061 | 0.1417520532 | 2.82E-10 | 5.18E-09 |
| TNFAIP8 | 864.7088677 | 0.8483525052 | 0.2222265834 | 1.73E-05 | 0.000138796968 |
| AEN | 7310.388652 | 0.8476382985 | 0.1728561951 | 1.30E-07 | 1.60E-06 |
| LPAR6 | 204.7395037 | 0.8473975868 | 0.1783018305 | 2.74E-07 | 3.16E-06 |
| IL4R | 5350.246448 | 0.8463612977 | 0.1770136825 | 2.38E-07 | 2.77E-06 |
| CFLAR | 2299.285887 | 0.8461495044 | 0.1453401759 | 8.34E-10 | 1.45E-08 |
| SFMBT1 | 443.3738127 | 0.8461315729 | 0.1271453308 | 4.16E-12 | 1.00E-10 |
| ARL5B | 1441.604216 | 0.844274991 | 0.1391242809 | 1.88E-10 | 3.56E-09 |
| RABGEF1 | 2015.641119 | 0.8437287534 | 0.1228838974 | 9.74E-13 | 2.53E-11 |
| LINC00152 | 6886.211103 | 0.8400213727 | 0.1064537114 | 4.52E-16 | 1.63E-14 |
| GADD45A | 1045.473618 | 0.8392058523 | 0.1358958988 | 7.66E-11 | 1.55E-09 |
| LOC102724584 | 17.68997469 | 0.8350143493 | 0.4332274967 | 0.00506978581 | 0.02011834779 |
| SH3RF3 | 890.8769124 | 0.8342913906 | 0.1179371053 | 2.32E-13 | 6.53E-12 |
| CHD1 | 2988.756734 | 0.8340051634 | 0.1283326665 | 1.17E-11 | 2.64E-10 |
| FBXO30 | 1659.163494 | 0.8322188269 | 0.1258498721 | 5.60E-12 | 1.33E-10 |
| RAP1GAP | 8.944439523 | 0.8314137326 | 0.9476341042 | 0.01492688986 | 0.04917729462 |
| AP5B1 | 676.9456286 | 0.8311824207 | 0.1087652878 | 3.26E-15 | 1.08E-13 |
| USP27X | 139.8515568 | 0.8296610386 | 0.1697642857 | 1.47E-07 | 1.80E-06 |
| LINC00936 | 40.52869422 | 0.8291715568 | 0.264627078 | 0.000217295471 | 0.001326491299 |
| JMJD6 | 1885.530193 | 0.8284839423 | 0.1130271212 | 3.43E-14 | 1.06E-12 |
| KDM6B | 111.4950741 | 0.8282748442 | 0.2367071359 | 6.08E-05 | 0.000429772510 |
| PHF13 | 770.6272486 | 0.828050747 | 0.1397821889 | 4.61E-10 | 8.25E-09 |
| DCP1A | 1187.553254 | 0.8270147042 | 0.07935361249 | 3.14E-26 | 2.60E-24 |
| RNVU1-15 | 70.4625942 | 0.8260187427 | 0.2979519527 | 0.000655654820 | 0.003479604585 |
| TUSC1 | 211.2394855 | 0.82549044 | 0.1505715881 | 6.14E-09 | 9.47E-08 |

| | baseMean | log2FoldChange | lfcSE | pvalue | padj |
|---|---|---|---|---|---|
| KITLG | 2705.078171 | 0.8253510992 | 0.1766690453 | 4.27E-07 | 4.72E-06 |
| KIAA0040 | 33.00500847 | 0.8230288312 | 0.3636913164 | 0.002532186553 | 0.01120740951 |
| USB1 | 354.6009597 | 0.8229140288 | 0.1409002543 | 7.87E-10 | 1.37E-08 |
| ZNF394 | 739.040695 | 0.8215411775 | 0.1103675908 | 1.51E-14 | 4.78E-13 |
| TRABD2A | 197.0071717 | 0.8195905173 | 0.2621624823 | 0.000223717927 | 0.001357177362 |
| FAM222A | 24.96188625 | 0.8192722086 | 0.4294155875 | 0.005291211993 | 0.02081946495 |
| CHST1 | 21.31905993 | 0.8191552258 | 0.4245827152 | 0.005168637236 | 0.02044522843 |
| SGK1 | 3095.257131 | 0.8186983393 | 0.1400200076 | 7.52E-10 | 1.32E-08 |
| KCTD5 | 1494.962007 | 0.8171543808 | 0.08926806402 | 8.79E-21 | 4.61E-19 |
| TUBA8 | 21.50886633 | 0.8171311942 | 0.3986451946 | 0.004137374282 | 0.01698614695 |
| TMCC3 | 124.1919167 | 0.8167824972 | 0.3450500523 | 0.001961544923 | 0.009027957451 |
| VGLL3 | 2277.182221 | 0.8162309744 | 0.1842605494 | 1.35E-06 | 1.34E-05 |
| DFNB31 | 55.39023856 | 0.8151210353 | 0.2385143448 | 8.43E-05 | 0.000577362956 |
| E2F6 | 769.322265 | 0.8137014778 | 0.1132995505 | 1.09E-13 | 3.16E-12 |
| VGF | 10.04852536 | 0.8133794093 | 0.6069209967 | 0.01272293464 | 0.04305345571 |
| TIMP3 | 3610.150685 | 0.8120387813 | 0.2735734933 | 0.000374886868 | 0.002125548584 |
| DOT1L | 477.3308538 | 0.8112845453 | 0.1194932153 | 1.79E-12 | 4.52E-11 |
| CDC14A | 149.2190158 | 0.8112514743 | 0.1607056242 | 6.77E-08 | 8.79E-07 |
| RNF182 | 19.72511821 | 0.8106351016 | 0.5948553841 | 0.01197950173 | 0.0410595767 |
| RGMB | 3423.340632 | 0.8097594862 | 0.277450595 | 0.000436584152 | 0.002445561115 |
| NANOS1 | 24.49139399 | 0.8087649888 | 0.4211554026 | 0.005272798665 | 0.02075798502 |
| DDX21 | 14775.52758 | 0.8086379003 | 0.2025990839 | 1.03E-05 | 8.62E-05 |
| TFB2M | 1081.678372 | 0.8079318559 | 0.1254348843 | 1.86E-11 | 4.10E-10 |
| PIGA | 592.1962885 | 0.8068918223 | 0.1525699538 | 1.86E-08 | 2.67E-07 |
| PGM2L1 | 377.5644305 | 0.8060048888 | 0.1669539536 | 2.08E-07 | 2.44E-06 |
| RFX2 | 105.9919609 | 0.8057161978 | 0.1964981945 | 5.87E-06 | 5.16E-05 |
| ZNF670 | 192.4531205 | 0.8054592973 | 0.1626108797 | 1.11E-07 | 1.40E-06 |
| SGMS2 | 778.7591151 | 0.8054279148 | 0.193099158 | 4.40E-06 | 3.96E-05 |
| LIMS1 | 3827.266502 | 0.8046516589 | 0.07976692602 | 1.03E-24 | 7.63E-23 |
| PANX1 | 4199.51525 | 0.8043966925 | 0.1594091144 | 6.84E-08 | 8.84E-07 |
| TAF13 | 842.083791 | 0.8002983871 | 0.1294714697 | 1.00E-10 | 1.98E-09 |
| EIF5A2 | 553.4247193 | 0.8002964683 | 0.1011180784 | 4.06E-16 | 1.47E-14 |
| BIRC2 | 3676.11745 | 0.8002505193 | 0.1283062275 | 6.88E-11 | 1.40E-09 |
| DUSP11 | 1087.072523 | 0.7989052495 | 0.09481996494 | 5.88E-18 | 2.50E-16 |
| LOC102723864 | 51.24385192 | 0.7988554195 | 0.2287239434 | 6.74E-05 | 0.000471262869 |
| SMURF1 | 1714.434499 | 0.7978087238 | 0.09727904451 | 3.78E-17 | 1.51E-15 |
| HIF1A | 23259.02636 | 0.7961692513 | 0.1396263358 | 2.50E-09 | 4.12E-08 |
| FAM60A | 2189.793969 | 0.7954164647 | 0.1143100032 | 5.57E-13 | 1.50E-11 |
| HDAC9 | 200.2028487 | 0.7951266809 | 0.2539340732 | 0.000232123007 | 0.001402629648 |
| AMIGO2 | 805.6731548 | 0.7945778415 | 0.2434782647 | 0.000149418744 | 0.000956435225 |

| | baseMean | log2FoldChange | lfcSE | pvalue | padj |
|---|---|---|---|---|---|
| DNTTIP2 | 4423.368623 | 0.7935284298 | 0.1222841175 | 1.40E-11 | 3.12E-10 |
| MAPKAPK2 | 3832.038511 | 0.793341271 | 0.07465504555 | 3.84E-27 | 3.40E-25 |
| HAPLN3 | 85.51883282 | 0.7927539401 | 0.2330937088 | 9.25E-05 | 0.000627334488 |
| HOMER1 | 776.5316438 | 0.7924044709 | 0.1919157319 | 5.41E-06 | 4.78E-05 |
| SLC25A32 | 1753.895711 | 0.7915919826 | 0.1311627931 | 2.58E-10 | 4.79E-09 |
| SLC35E4 | 341.8633734 | 0.7902951673 | 0.141998773 | 4.16E-09 | 6.61E-08 |
| NRARP | 13.46396542 | 0.7897700291 | 0.5488977201 | 0.0117349393 | 0.04035138827 |
| BCAR1 | 2775.170505 | 0.7889736859 | 0.125643301 | 5.79E-11 | 1.19E-09 |
| SPECC1 | 769.7326299 | 0.787711919 | 0.09457651819 | 1.33E-17 | 5.58E-16 |
| PMM2 | 2178.759623 | 0.7876920954 | 0.1093485306 | 9.65E-14 | 2.83E-12 |
| ASB6 | 1187.804712 | 0.786639413 | 0.09705059012 | 9.43E-17 | 3.55E-15 |
| ADNP2 | 1732.419738 | 0.7837915167 | 0.08313600237 | 7.21E-22 | 4.13E-20 |
| MOAP1 | 471.7620073 | 0.783518066 | 0.1337166371 | 7.54E-10 | 1.32E-08 |
| GLI2 | 375.7203945 | 0.7822057706 | 0.1943123862 | 8.51E-06 | 7.28E-05 |
| TRPC6 | 62.59009047 | 0.7808561342 | 0.2874269439 | 0.000851791697 | 0.004379256416 |
| DIXDC1 | 1177.523708 | 0.7806708764 | 0.122698806 | 3.57E-11 | 7.53E-10 |
| SLC7A5P1 | 35.41056759 | 0.7806079282 | 0.3171812937 | 0.001729641393 | 0.008105958673 |
| CHST2 | 568.6858869 | 0.7791935982 | 0.1683558922 | 5.79E-07 | 6.27E-06 |
| RNA5-8S5 | 69788.33821 | 0.7790122043 | 0.1716826784 | 8.89E-07 | 9.19E-06 |
| OSGIN1 | 1313.492142 | 0.7779305898 | 0.24830504 | 0.000240278445 | 0.001445456879 |
| LINC00843 | 594.7535758 | 0.7777623594 | 0.1220289761 | 3.09E-11 | 6.64E-10 |
| NCBP2-AS2 | 557.1950499 | 0.7765362794 | 0.09817764032 | 4.35E-16 | 1.57E-14 |
| TICAM1 | 287.734414 | 0.7752189594 | 0.1607367676 | 2.27E-07 | 2.65E-06 |
| PLAU | 27724.18917 | 0.7750949772 | 0.1293295346 | 3.45E-10 | 6.24E-09 |
| WTAP | 3758.587751 | 0.7740874268 | 0.08239895854 | 9.90E-22 | 5.56E-20 |
| TRIM36 | 36.86166289 | 0.7732455731 | 0.3304012567 | 0.002373692479 | 0.01060045211 |
| DIEXF | 1312.018027 | 0.773085421 | 0.1535304255 | 7.87E-08 | 1.01E-06 |
| SMIM13 | 1115.016164 | 0.771557565 | 0.1592614316 | 2.04E-07 | 2.41E-06 |
| NDEL1 | 2148.670263 | 0.7713485031 | 0.07597097807 | 5.62E-25 | 4.25E-23 |
| GAREM | 246.5185743 | 0.7709294873 | 0.1279575405 | 2.78E-10 | 5.12E-09 |
| RNVU1-14 | 96.5902399 | 0.7701640089 | 0.256354656 | 0.000367864408 | 0.002089711249 |
| LEMD3 | 1439.88427 | 0.7696565812 | 0.1013429662 | 5.35E-15 | 1.75E-13 |
| ZFP36 | 271.0926675 | 0.769158874 | 0.1790010427 | 2.69E-06 | 2.53E-05 |
| CCDC68 | 2093.445292 | 0.7674364818 | 0.09465454964 | 9.23E-17 | 3.49E-15 |
| RPGR | 390.190346 | 0.7672293923 | 0.1474544925 | 3.24E-08 | 4.44E-07 |
| LRRC8E | 407.9316061 | 0.7669748593 | 0.1146893949 | 3.83E-12 | 9.25E-11 |
| LOC101929690 | 37.59796087 | 0.7650753047 | 0.3467446105 | 0.003306351945 | 0.01410956552 |
| AKIRIN1 | 2958.068649 | 0.7623973524 | 0.08076851755 | 6.66E-22 | 3.85E-20 |
| HS3ST3B1 | 795.3105444 | 0.7623606242 | 0.1178515069 | 1.73E-11 | 3.82E-10 |
| BNIP2 | 2746.011927 | 0.760974798 | 0.09716834292 | 8.57E-16 | 3.05E-14 |

| | baseMean | log2FoldChange | lfcSE | pvalue | padj |
|---|---|---|---|---|---|
| TIMP1 | 18831.58644 | 0.7604318896 | 0.1062619614 | 1.85E-13 | 5.27E-12 |
| C16orf87 | 574.9508969 | 0.7600072569 | 0.1412341105 | 1.24E-08 | 1.83E-07 |
| TRAF1 | 227.9106577 | 0.7597977226 | 0.1272444104 | 4.06E-10 | 7.28E-09 |
| PDZD2 | 27.55149671 | 0.7594724997 | 0.3940500082 | 0.006010974201 | 0.02311996768 |
| TLR6 | 117.743358 | 0.7594595848 | 0.1885575925 | 9.05E-06 | 7.69E-05 |
| C20orf112 | 353.8493876 | 0.7591764814 | 0.166977852 | 8.92E-07 | 9.22E-06 |
| ZNF230 | 215.2458618 | 0.7591048578 | 0.1669406653 | 8.76E-07 | 9.09E-06 |
| UPP1 | 1053.488607 | 0.757491061 | 0.1216700919 | 8.58E-11 | 1.72E-09 |
| BLOC1S4 | 506.364527 | 0.7572349127 | 0.1249854294 | 2.37E-10 | 4.43E-09 |
| RIN2 | 2857.52542 | 0.7570857092 | 0.1191408041 | 3.92E-11 | 8.24E-10 |
| LRIG1 | 774.4334036 | 0.7569987274 | 0.140162138 | 1.14E-08 | 1.69E-07 |
| DIS3 | 3415.063826 | 0.7566205346 | 0.1147257526 | 7.41E-12 | 1.73E-10 |
| NRBF2 | 1268.381265 | 0.7559837019 | 0.09202064946 | 3.77E-17 | 1.51E-15 |
| PLK3 | 688.8991544 | 0.7558967345 | 0.1348891062 | 3.60E-09 | 5.74E-08 |
| FOXO3B | 389.9402384 | 0.7553533998 | 0.1337045934 | 2.78E-09 | 4.53E-08 |
| FMNL1 | 87.71384725 | 0.7544988238 | 0.3115729391 | 0.001992222409 | 0.009140893512 |
| RASA2 | 707.5459193 | 0.7527287595 | 0.1162808309 | 1.69E-11 | 3.74E-10 |
| CCNJ | 649.9998367 | 0.7516032785 | 0.1704259276 | 1.70E-06 | 1.66E-05 |
| ZNF487 | 45.59691864 | 0.7511964734 | 0.2717984757 | 0.000797087580 | 0.004136576152 |
| FCHSD2 | 603.5824487 | 0.7502059194 | 0.1229077666 | 1.82E-10 | 3.45E-09 |
| ZNF263 | 1398.667955 | 0.7499443711 | 0.08738245502 | 1.70E-18 | 7.67E-17 |
| SDCBP2 | 59.0421693 | 0.7497806551 | 0.2840798629 | 0.001139970249 | 0.005667337451 |
| ZNF136 | 310.4524022 | 0.7474361981 | 0.1596899857 | 4.86E-07 | 5.34E-06 |
| CDC25A | 1497.545842 | 0.7448163111 | 0.2213684234 | 0.000117228626 | 0.000776776598 |
| ZNRF2 | 95.2567895 | 0.7445876096 | 0.171193325 | 2.28E-06 | 2.16E-05 |
| ZNF430 | 352.1573649 | 0.7443705107 | 0.1094093887 | 1.83E-12 | 4.62E-11 |
| GABARAPL1 | 3188.950686 | 0.7443024001 | 0.2308254947 | 0.000191475555 | 0.001186383495 |
| PTRH2 | 1131.268008 | 0.744057266 | 0.1363012554 | 8.40E-09 | 1.27E-07 |
| RNU11 | 51.87519231 | 0.7422193723 | 0.4227612871 | 0.008280334889 | 0.03009407203 |
| SNHG17 | 231.4353928 | 0.7420532273 | 0.1560017293 | 3.41E-07 | 3.88E-06 |
| SLC25A33 | 567.7035637 | 0.7420146153 | 0.1592695448 | 5.40E-07 | 5.91E-06 |
| C16orf91 | 585.4971872 | 0.7418614846 | 0.09302821625 | 2.80E-16 | 1.03E-14 |
| GNG4 | 47.38266222 | 0.741199181 | 0.2616555185 | 0.000676083929 | 0.003569579298 |
| BCL7A | 422.4497226 | 0.7392646645 | 0.1512573282 | 1.77E-07 | 2.12E-06 |
| ADO | 1029.423484 | 0.7389964834 | 0.09740738217 | 6.01E-15 | 1.96E-13 |
| APCDD1L | 94.63163983 | 0.7382879584 | 0.1839050915 | 1.00E-05 | 8.42E-05 |
| ZNF256 | 241.763056 | 0.7373618064 | 0.1519629201 | 2.15E-07 | 2.52E-06 |
| SOCS3 | 2586.934244 | 0.7370432546 | 0.121943786 | 2.82E-10 | 5.18E-09 |
| CYTH1 | 662.0331579 | 0.7364057643 | 0.1140715494 | 1.78E-11 | 3.92E-10 |
| SEC14L2 | 912.0291014 | 0.7362760713 | 0.11978976 | 1.44E-10 | 2.78E-09 |

| | baseMean | log2FoldChange | lfcSE | pvalue | padj |
|---|---|---|---|---|---|
| LAMC2 | 32.61063854 | 0.7361736771 | 0.3454393248 | 0.004163013027 | 0.01706113431 |
| STAT5A | 284.5131208 | 0.7346506833 | 0.3068098416 | 0.002229338385 | 0.01004216898 |
| PDLIM4 | 2479.816416 | 0.7344505805 | 0.1157075293 | 4.04E-11 | 8.46E-10 |
| FRMD4A | 1935.378466 | 0.7343377651 | 0.1814246312 | 8.68E-06 | 7.41E-05 |
| ABT1 | 1569.784562 | 0.732048618 | 0.07670998841 | 2.62E-22 | 1.55E-20 |
| OAF | 1714.119162 | 0.7307528433 | 0.1003023441 | 6.10E-14 | 1.82E-12 |
| MLF1 | 440.5420314 | 0.7307382835 | 0.1149823963 | 3.85E-11 | 8.10E-10 |
| TRAF3 | 1478.488919 | 0.7303111658 | 0.08604378087 | 4.03E-18 | 1.75E-16 |
| GRAMD3 | 484.9293898 | 0.730080538 | 0.1498719611 | 1.95E-07 | 2.32E-06 |
| TBC1D15 | 1077.030007 | 0.7300205717 | 0.1137692058 | 2.60E-11 | 5.62E-10 |
| DLC1 | 6043.234623 | 0.7299958752 | 0.119244638 | 1.82E-10 | 3.45E-09 |
| ZBTB7A | 414.5261204 | 0.7292823194 | 0.1519393433 | 2.81E-07 | 3.23E-06 |
| BAG5 | 1986.370252 | 0.7289744925 | 0.103607576 | 3.70E-13 | 1.02E-11 |
| GTF3C4 | 1673.709085 | 0.7282725338 | 0.09191479053 | 4.39E-16 | 1.59E-14 |
| SLC7A2 | 9558.487703 | 0.7263740975 | 0.1736503259 | 3.75E-06 | 3.42E-05 |
| SLC51B | 17.72349764 | 0.724198603 | 0.5021269651 | 0.01372379381 | 0.0458147009 |
| KDSR | 1443.982895 | 0.7227959064 | 0.09285204217 | 1.35E-15 | 4.65E-14 |
| BCL7B | 1609.8393 | 0.7225211321 | 0.1086605759 | 5.58E-12 | 1.33E-10 |
| KLHL25 | 397.1407632 | 0.7217962314 | 0.1190998119 | 2.52E-10 | 4.69E-09 |
| CTDP1 | 900.3972049 | 0.720147812 | 0.08429661176 | 2.54E-18 | 1.13E-16 |
| PNPLA8 | 1523.066384 | 0.7196463722 | 0.104326734 | 1.02E-12 | 2.64E-11 |
| EXOC8 | 989.0174857 | 0.7195803214 | 0.1028709424 | 4.99E-13 | 1.35E-11 |
| RNF19A | 1717.845626 | 0.7190182892 | 0.07826525213 | 8.02E-21 | 4.23E-19 |
| LOC101929792 | 335.1057306 | 0.7179962901 | 0.1303286937 | 6.70E-09 | 1.03E-07 |
| ZNF275 | 523.6336722 | 0.716761615 | 0.1394253105 | 5.09E-08 | 6.77E-07 |
| LOC92249 | 72.42977816 | 0.7161803358 | 0.214197578 | 0.000135890422 | 0.000880832605 |
| ZNF749 | 207.7750599 | 0.7158327883 | 0.1399786801 | 5.89E-08 | 7.75E-07 |
| ZNF614 | 583.0194872 | 0.7151630735 | 0.1266369045 | 3.07E-09 | 4.96E-08 |
| SPRTN | 640.0091708 | 0.7139665525 | 0.09997604377 | 1.78E-13 | 5.08E-12 |
| LOC102724935 | 26.58996061 | 0.7132278681 | 0.4435445837 | 0.0114641675 | 0.03957569904 |
| TANK | 1696.391497 | 0.7129979152 | 0.08899754646 | 2.21E-16 | 8.15E-15 |
| RNU6ATAC | 93.50090232 | 0.7122590248 | 0.3407957575 | 0.004769132208 | 0.01913951737 |
| CTSS | 50.94852644 | 0.7110716153 | 0.2923497532 | 0.002185944514 | 0.009880548838 |
| NOX4 | 88.35665908 | 0.7084237455 | 0.2183076194 | 1.97E-04 | 1.21E-03 |
| SBNO2 | 1556.990356 | 0.7074978992 | 0.1801546935 | 1.52E-05 | 0.000123654652 |
| RASSF3 | 1293.264688 | 0.7070261337 | 0.1192304162 | 5.84E-10 | 1.03E-08 |
| GPATCH3 | 537.9054438 | 0.7040287745 | 0.1190231721 | 6.39E-10 | 1.12E-08 |
| RNF168 | 2113.321147 | 0.7038577391 | 0.07626218397 | 5.52E-21 | 2.99E-19 |
| TNFRSF11B | 1864.773539 | 0.7036490749 | 0.1538049755 | 8.87E-07 | 9.18E-06 |
| LOC100131496 | 60.11256903 | 0.7031695081 | 0.3156492172 | 0.003684085342 | 0.01544701398 |

| | baseMean | log2FoldChange | lfcSE | pvalue | padj |
|---|---|---|---|---|---|
| KLHL26 | 111.0566339 | 0.7023761885 | 0.2234102714 | 0.000276343790 | 0.001633379395 |
| RAPGEF2 | 2609.755472 | 0.7021576671 | 0.1135725478 | 1.24E-10 | 2.41E-09 |
| MYPN | 166.6949832 | 0.7012864829 | 0.316304805 | 0.003730030784 | 0.01560450361 |
| C1D | 749.8775554 | 0.7003204013 | 0.1113811208 | 6.34E-11 | 1.30E-09 |
| CCDC59 | 1115.55465 | 0.7000204467 | 0.1284477846 | 9.58E-09 | 1.44E-07 |
| RPUSD2 | 762.252729 | 0.6996135873 | 0.1062114869 | 9.54E-12 | 2.19E-10 |
| TOR1A | 2357.044493 | 0.6963874229 | 0.07369738278 | 7.19E-22 | 4.13E-20 |
| NUFIP2 | 4505.129201 | 0.6959309142 | 0.1345856667 | 4.57E-08 | 6.14E-07 |
| TBC1D10A | 955.1979913 | 0.6959120231 | 0.1405335736 | 1.40E-07 | 1.72E-06 |
| WDR43 | 6049.041747 | 0.6952260302 | 0.1518490349 | 9.26E-07 | 9.54E-06 |
| C19orf26 | 94.68509412 | 0.694942324 | 0.2547947403 | 0.001033722623 | 0.005195688611 |
| EIF1 | 9873.62585 | 0.6945150491 | 0.05895140286 | 1.05E-32 | 1.34E-30 |
| ACVR1 | 978.7756334 | 0.6943596944 | 0.118139326 | 8.28E-10 | 1.44E-08 |
| RAB40B | 240.5087319 | 0.6942262828 | 0.2651719919 | 0.001387902515 | 0.006720156275 |
| LOC101927841 | 248.1010248 | 0.6926126921 | 0.1359556433 | 6.80E-08 | 8.82E-07 |
| GPCPD1 | 1436.025948 | 0.6914020454 | 0.1257420494 | 9.21E-09 | 1.39E-07 |
| ZBTB34 | 694.490944 | 0.6907655697 | 0.1312446676 | 2.81E-08 | 3.89E-07 |
| LINC00869 | 24.45674798 | 0.6904687542 | 0.4452939942 | 0.01320104003 | 0.04437802776 |
| PITPNC1 | 259.9682863 | 0.6899825285 | 0.2139341984 | 0.000220060157 | 0.001340071854 |
| FAM131B | 96.53545458 | 0.6893812819 | 0.4095084408 | 0.01083632945 | 0.03776814703 |
| ZNF48 | 471.5272814 | 0.6893116506 | 0.1293667585 | 1.97E-08 | 2.81E-07 |
| RYBP | 1004.104182 | 0.6891197141 | 0.1377462988 | 1.11E-07 | 1.39E-06 |
| LYSMD2 | 198.6085492 | 0.6881812312 | 0.1494154112 | 7.96E-07 | 8.35E-06 |
| PCDH7 | 1680.625465 | 0.6877093631 | 0.2333014007 | 0.000542452565 | 0.002948797121 |
| MAPK6 | 5039.386105 | 0.6876998163 | 0.1211664087 | 2.69E-09 | 4.40E-08 |
| ATP5J2-PTCD1 | 125.8461487 | 0.687037404 | 0.3433888412 | 0.006150527519 | 0.02358969712 |
| ACSL4 | 9392.399622 | 0.6863932606 | 0.1704311726 | 1.04E-05 | 8.74E-05 |
| TPST2 | 735.8863204 | 0.6852861981 | 0.09859220474 | 7.59E-13 | 2.01E-11 |
| STAT4 | 22.84337071 | 0.6848967844 | 0.3856424653 | 0.009559496067 | 0.03396118753 |
| GTF2B | 1542.515198 | 0.6847400978 | 0.06950785079 | 1.43E-23 | 9.74E-22 |
| URGCP | 1832.78326 | 0.6835717605 | 0.07610047508 | 5.65E-20 | 2.77E-18 |
| RGS9 | 103.4196848 | 0.6830213916 | 0.2318459449 | 0.000559351124 | 0.003028428686 |
| PTGES | 442.7406056 | 0.6821246291 | 0.3664842548 | 0.00819515902 | 0.02982589163 |
| CDC42SE2 | 1207.034885 | 0.681334097 | 0.1218423402 | 4.59E-09 | 7.23E-08 |
| NOP16 | 2112.16933 | 0.6805523754 | 0.164829206 | 7.01E-06 | 6.08E-05 |
| ADAM17 | 2140.528387 | 0.6804678688 | 0.1361228723 | 1.16E-07 | 1.45E-06 |
| NKRF | 449.5125479 | 0.679279775 | 0.1431214985 | 4.12E-07 | 4.58E-06 |
| UBTD1 | 874.5301934 | 0.6790453135 | 0.1255262497 | 1.29E-08 | 1.91E-07 |
| MED26 | 248.2638864 | 0.678222364 | 0.1187257793 | 2.30E-09 | 3.80E-08 |
| ZNF649 | 481.8124655 | 0.6771912428 | 0.1376726248 | 1.76E-07 | 2.11E-06 |

| | baseMean | log2FoldChange | lfcSE | pvalue | padj |
|---|---|---|---|---|---|
| RPS6KA3 | 2206.796703 | 0.6771248795 | 0.1217206734 | 5.47E-09 | 8.51E-08 |
| SNHG15 | 702.6936372 | 0.6752860333 | 0.1357528955 | 1.33E-07 | 1.64E-06 |
| RBBP6 | 1681.667037 | 0.6750697277 | 0.08985586131 | 1.25E-14 | 3.98E-13 |
| ITGA2 | 5374.055756 | 0.6744907662 | 0.218439611 | 0.000367011722 | 0.002086459539 |
| HS3ST3A1 | 582.6092369 | 0.6737164695 | 0.2061172854 | 0.000198079207 | 0.00122221356 |
| MAK16 | 2605.330025 | 0.6733739986 | 0.1535921586 | 2.30E-06 | 2.18E-05 |
| ETV6 | 548.7845937 | 0.6731068585 | 0.1714835664 | 1.68E-05 | 0.000135582285 |
| DPM1 | 2006.971945 | 0.6726868424 | 0.08091442481 | 2.04E-17 | 8.23E-16 |
| RNF139 | 1005.3526 | 0.672028352 | 0.1275750645 | 2.89E-08 | 3.98E-07 |
| C21orf91 | 430.3253654 | 0.6714875819 | 0.1321350655 | 7.67E-08 | 9.84E-07 |
| HNRNPDL | 9330.859308 | 0.6714023988 | 0.1103797898 | 2.33E-10 | 4.36E-09 |
| SMIM3 | 4690.170214 | 0.6702369791 | 0.1102519472 | 2.74E-10 | 5.07E-09 |
| PQLC1 | 1537.940485 | 0.6701057926 | 0.1352542536 | 1.48E-07 | 1.81E-06 |
| MAPK8 | 1165.617829 | 0.6699963417 | 0.1104482448 | 2.85E-10 | 5.22E-09 |
| CREM | 706.9538754 | 0.6697704607 | 0.1106024368 | 2.92E-10 | 5.35E-09 |
| WDR37 | 979.169192 | 0.6687174593 | 0.08659369474 | 2.49E-15 | 8.45E-14 |
| ARHGEF3 | 320.5333338 | 0.6683959124 | 0.1943586834 | 0.000110381845 | 0.000735994308 |
| TNFAIP8L3 | 189.6997854 | 0.6681074901 | 0.1401643633 | 3.84E-07 | 4.32E-06 |
| ZYX | 4213.381663 | 0.6675217079 | 0.09242315283 | 9.22E-14 | 2.71E-12 |
| ARHGEF26 | 212.0759127 | 0.6672801652 | 0.1655620676 | 1.10E-05 | 9.16E-05 |
| SPRED1 | 2248.624514 | 0.6664453744 | 0.1269402664 | 3.32E-08 | 4.54E-07 |
| TNFRSF1B | 903.3363144 | 0.6661787363 | 0.1338566508 | 1.34E-07 | 1.65E-06 |
| MALT1 | 1207.701575 | 0.6661399082 | 0.1304363254 | 6.88E-08 | 8.89E-07 |
| ZNF778 | 318.5293343 | 0.6660061278 | 0.1764393648 | 3.13E-05 | 2.36E-04 |
| PPP3R1 | 1359.844701 | 0.6658391479 | 0.1008401607 | 8.71E-12 | 2.01E-10 |
| URB2 | 1411.03033 | 0.6655540282 | 0.1870238258 | 7.22E-05 | 0.000502704030 |
| TANC2 | 1627.237296 | 0.663208686 | 0.1299498045 | 6.98E-08 | 9.00E-07 |
| PAPD7 | 732.7536167 | 0.6629722556 | 0.1108544291 | 4.80E-10 | 8.56E-09 |
| PHEX | 20.70903037 | 0.662547636 | 0.410415967 | 0.01350570941 | 0.04520829753 |
| GSTO1 | 13166.91249 | 0.6623639432 | 0.1406841296 | 5.43E-07 | 5.93E-06 |
| LTV1 | 2052.535758 | 0.6617816404 | 0.1385496654 | 3.71E-07 | 4.19E-06 |
| TGFB1 | 590.8556599 | 0.6610669265 | 0.1766241702 | 3.59E-05 | 0.000267496956 |
| GRPEL1 | 3453.127209 | 0.6606325966 | 0.1175859693 | 4.22E-09 | 6.69E-08 |
| PELI2 | 642.5892608 | 0.6596717238 | 0.2339869254 | 0.000862996568 | 0.004427689835 |
| DDX3X | 18363.37039 | 0.6593042542 | 0.1093182298 | 5.48E-10 | 9.69E-09 |
| ZNF408 | 423.8105977 | 0.6591939159 | 0.1351247352 | 2.24E-07 | 2.61E-06 |
| LOC102724594 | 58.83638989 | 0.6580404174 | 0.335553422 | 0.007339970437 | 0.0272462777 |
| PTBP2 | 740.4190108 | 0.6571226956 | 0.1353480532 | 2.50E-07 | 2.89E-06 |
| KLF6 | 3703.984238 | 0.656626718 | 0.1430226128 | 9.24E-07 | 9.53E-06 |
| KCNK1 | 74.35796125 | 0.6562869604 | 0.195704136 | 0.000155525543 | 0.000988736044 |

| | baseMean | log2FoldChange | lfcSE | pvalue | padj |
|---|---|---|---|---|---|
| MICAL2 | 9264.229436 | 0.6559233372 | 0.1451694979 | 1.31E-06 | 1.30E-05 |
| DDX5 | 25403.40134 | 0.6554632457 | 0.1157718886 | 3.26E-09 | 5.24E-08 |
| EPHA2 | 1992.614092 | 0.655048596 | 0.1271015354 | 6.05E-08 | 7.93E-07 |
| MT1X | 378.0110429 | 0.6548535047 | 0.1214575163 | 1.48E-08 | 2.16E-07 |
| BIRC3 | 264.8518558 | 0.6545192851 | 0.2752769304 | 0.002897921449 | 0.01259403166 |
| GSKIP | 1040.237177 | 0.6541767308 | 0.09102090544 | 1.49E-13 | 4.28E-12 |
| ST3GAL5 | 1217.124293 | 0.6540680721 | 0.09943529063 | 1.06E-11 | 2.40E-10 |
| AFF4 | 4377.817723 | 0.6537003282 | 0.1168679624 | 4.85E-09 | 7.63E-08 |
| HES1 | 43.57561223 | 0.6532628074 | 0.3403175525 | 0.008126338485 | 0.0296188626 |
| ZNF222 | 212.7699683 | 0.6499683639 | 0.124260943 | 3.70E-08 | 5.04E-07 |
| RICTOR | 1665.579882 | 0.6490995158 | 0.1238684025 | 3.44E-08 | 4.69E-07 |
| C2CD2 | 1532.419213 | 0.6484596928 | 0.1413954665 | 9.67E-07 | 9.89E-06 |
| SNN | 506.6716311 | 0.6482567613 | 0.1467561018 | 2.12E-06 | 2.02E-05 |
| FNIP1 | 1812.249464 | 0.6480147426 | 0.08461322634 | 4.38E-15 | 1.44E-13 |
| MED30 | 317.2253976 | 0.6476477167 | 0.11206924 | 1.68E-09 | 2.81E-08 |
| FOXP1 | 2030.422292 | 0.645196729 | 0.1036061177 | 1.06E-10 | 2.08E-09 |
| GFPT2 | 3793.900501 | 0.6447433397 | 0.1591919496 | 1.11E-05 | 9.30E-05 |
| CARD10 | 528.7376831 | 0.6446774066 | 0.1158124882 | 5.80E-09 | 8.98E-08 |
| STX1A | 458.9380229 | 0.6437550972 | 0.1908583155 | 0.000149481145 | 0.000956435225 |
| ABHD13 | 486.8080733 | 0.6432917897 | 0.1490578128 | 3.42E-06 | 3.14E-05 |
| DUSP14 | 2105.572431 | 0.6431486559 | 0.1023412696 | 7.46E-11 | 1.51E-09 |
| CEBPZ | 4608.340322 | 0.6430531375 | 0.1213928796 | 2.78E-08 | 3.86E-07 |
| FGF7 | 2948.828058 | 0.6430383732 | 0.2058208729 | 0.000312347104 | 0.001819458545 |
| POPDC3 | 639.35152 | 0.6411670507 | 0.1538770873 | 6.63E-06 | 5.78E-05 |
| RAI1 | 779.2705037 | 0.6388872695 | 0.1271605663 | 1.13E-07 | 1.41E-06 |
| FGFRL1 | 875.9542897 | 0.638404136 | 0.1165703877 | 9.81E-09 | 1.47E-07 |
| SOCS5 | 1281.930115 | 0.6376795277 | 0.0910310526 | 5.79E-13 | 1.55E-11 |
| IFNGR2 | 2629.561066 | 0.6375100745 | 0.09335158508 | 2.00E-12 | 5.02E-11 |
| ARRDC2 | 203.3221016 | 0.6374874419 | 0.1894335916 | 0.000155947713 | 0.000990996803 |
| SLK | 4390.288507 | 0.6364363716 | 0.117769183 | 1.49E-08 | 2.16E-07 |
| YTHDF2 | 3672.87434 | 0.6357542261 | 0.100137852 | 4.90E-11 | 1.01E-09 |
| YIPF5 | 3863.446662 | 0.6349961761 | 0.09000966701 | 4.05E-13 | 1.11E-11 |
| SSTR1 | 224.1814883 | 0.6344185639 | 0.2186499495 | 0.000729887021 | 0.003826509809 |
| PLEKHA3 | 514.4968227 | 0.6339271926 | 0.1135989762 | 5.63E-09 | 8.73E-08 |
| RCE1 | 496.4978494 | 0.6333106614 | 0.1025704021 | 1.55E-10 | 2.98E-09 |
| TOPORS | 1778.008675 | 0.6324131635 | 0.09269025861 | 2.11E-12 | 5.26E-11 |
| FGFR1OP2 | 1348.413295 | 0.6320191612 | 0.08903766026 | 3.00E-13 | 8.37E-12 |
| SERTAD2 | 521.8400882 | 0.630769728 | 0.1339179117 | 5.62E-07 | 6.11E-06 |
| DCUN1D5 | 2049.690821 | 0.6304984836 | 0.1175374574 | 1.76E-08 | 2.54E-07 |
| CENPQ | 328.0696256 | 0.6303250791 | 0.2014500923 | 0.000358203305 | 0.002046542214 |

| | baseMean | log2FoldChange | lfcSE | pvalue | padj |
|---|---|---|---|---|---|
| ANKH | 408.0041502 | 0.6300852321 | 0.1378821011 | 1.11E-06 | 1.12E-05 |
| MYO1E | 3860.777176 | 0.6293822284 | 0.1019009192 | 1.49E-10 | 2.88E-09 |
| MXD1 | 207.8186242 | 0.627621563 | 0.1414514253 | 2.05E-06 | 1.96E-05 |
| SRGN | 244.5806609 | 0.6263709812 | 0.2461159808 | 0.002072137285 | 0.009458017179 |
| C16orf45 | 2381.148548 | 0.6259938719 | 0.1606866155 | 2.14E-05 | 0.000168290417 |
| LIMA1 | 4555.75672 | 0.6259094531 | 0.1157698675 | 1.47E-08 | 2.15E-07 |
| ARPC5L | 1587.776655 | 0.6258991122 | 0.1135383566 | 8.34E-09 | 1.27E-07 |
| MED10 | 682.7749424 | 0.6239633241 | 0.1256900192 | 1.40E-07 | 1.71E-06 |
| TNFRSF10B | 7353.925351 | 0.6236227831 | 0.103122207 | 3.54E-10 | 6.38E-09 |
| VPS18 | 2255.586266 | 0.6233653707 | 0.06509269648 | 2.47E-22 | 1.48E-20 |
| SNORA23 | 375.2578747 | 0.6224187519 | 0.204578535 | 0.000483018237 | 0.00266752913 |
| RPP38 | 575.2890381 | 0.6216865486 | 0.0967592733 | 3.18E-11 | 6.80E-10 |
| FAM109A | 331.2433154 | 0.6212805744 | 0.135740116 | 1.09E-06 | 1.11E-05 |
| ALG2 | 1387.298123 | 0.6212049956 | 0.09871566902 | 7.53E-11 | 1.52E-09 |
| VASP | 5748.429818 | 0.62056402 | 0.08036381134 | 2.78E-15 | 9.31E-14 |
| ZNF26 | 440.6450882 | 0.620534229 | 0.1232164612 | 1.12E-07 | 1.40E-06 |
| IRF2BPL | 1449.702154 | 0.6204028942 | 0.2107535611 | 0.000643022056 | 0.003421713868 |
| DNMBP | 2387.333325 | 0.6192610995 | 0.1386420838 | 1.84E-06 | 1.78E-05 |
| CHSY1 | 1989.266859 | 0.6185895879 | 0.09763491464 | 6.77E-11 | 1.38E-09 |
| FOXN2 | 1209.6309 | 0.618537834 | 0.1271510933 | 2.50E-07 | 2.90E-06 |
| SOCS6 | 1574.662541 | 0.6185089725 | 0.0889315762 | 8.81E-13 | 2.31E-11 |
| USP3 | 1380.467143 | 0.6178641093 | 0.08406604036 | 4.93E-14 | 1.50E-12 |
| GLIS1 | 142.1912962 | 0.6178003263 | 0.1922150331 | 0.000278775476 | 0.001646445091 |
| NR2F1 | 713.0178377 | 0.6176136843 | 0.1856925077 | 0.000192396211 | 0.001191096548 |
| SNORA3 | 58.87744691 | 0.6174980507 | 0.2858897832 | 0.005477901393 | 0.0213844976 |
| TMEM167B | 2156.074233 | 0.6158730514 | 0.08396714177 | 5.53E-14 | 1.66E-12 |
| ACVR1B | 623.9838154 | 0.6156221507 | 0.1596212247 | 2.60E-05 | 0.000200113929 |
| ECE2 | 262.8147612 | 0.6155948909 | 0.1462468379 | 5.89E-06 | 5.17E-05 |
| RRP12 | 2712.175094 | 0.6153753384 | 0.2268157829 | 0.001327256819 | 0.006470702938 |
| TRIM38 | 1220.788475 | 0.6153285579 | 0.09875133666 | 1.13E-10 | 2.21E-09 |
| CITED2 | 3169.855992 | 0.6146319159 | 0.09984694523 | 1.83E-10 | 3.46E-09 |
| NHS | 58.41103432 | 0.6145337244 | 0.2854614113 | 0.005659577015 | 0.02201866793 |
| ANKRD12 | 1887.96154 | 0.6143334935 | 0.1423568424 | 3.73E-06 | 3.40E-05 |
| GBP1 | 3723.9076 | 0.6134027988 | 0.2133984166 | 0.000841805237 | 0.004332401722 |
| SNORA71D | 334.0159623 | 0.6129271386 | 0.2450529972 | 0.002418868259 | 0.01077311681 |
| RSC1A1 | 263.2600598 | 0.6125571512 | 0.1442649196 | 5.10E-06 | 4.52E-05 |
| RNF24 | 1320.319868 | 0.6124076054 | 0.1409625155 | 3.27E-06 | 3.01E-05 |
| IRAK3 | 491.8338428 | 0.6121774349 | 0.1917209186 | 0.000305547742 | 0.001785439695 |
| MAFG | 1489.698181 | 0.6119208783 | 0.09116351706 | 4.74E-12 | 1.13E-10 |
| WDR53 | 376.6697119 | 0.6111289633 | 0.1489314397 | 9.61E-06 | 8.11E-05 |

| | baseMean | log2FoldChange | lfcSE | pvalue | padj |
|---|---|---|---|---|---|
| L3MBTL3 | 539.7927186 | 0.6109253381 | 0.1324450589 | 9.51E-07 | 9.76E-06 |
| LSM11 | 239.0387366 | 0.6106552869 | 0.1658514091 | 5.29E-05 | 0.000379022926 |
| SNX9 | 3117.503936 | 0.6103153547 | 0.07759479042 | 9.36E-16 | 3.29E-14 |
| RBM38 | 375.9409955 | 0.6102992874 | 0.1141347949 | 2.20E-08 | 3.12E-07 |
| GK | 672.5037699 | 0.6094418486 | 0.1953444258 | 0.000392452089 | 0.002217540479 |
| ZNF71 | 511.9639015 | 0.6093649231 | 0.08700736286 | 5.44E-13 | 1.46E-11 |
| SNX4 | 1173.240892 | 0.6086935244 | 0.08590952315 | 3.48E-13 | 9.66E-12 |
| ISCA1 | 1151.732167 | 0.6086728357 | 0.07552267679 | 1.95E-16 | 7.22E-15 |
| HIVEP1 | 1177.949871 | 0.6061686946 | 0.1642694549 | 5.15E-05 | 0.000370144279 |
| CDC42EP1 | 1753.479345 | 0.6061039009 | 0.1208415105 | 1.30E-07 | 1.60E-06 |
| EFHD2 | 2772.858755 | 0.6049962232 | 0.09049456035 | 5.86E-12 | 1.39E-10 |
| KIFC3 | 2964.649441 | 0.6048461148 | 0.09193573241 | 1.20E-11 | 2.70E-10 |
| ZBTB2 | 1221.580301 | 0.604234854 | 0.1082614508 | 5.99E-09 | 9.26E-08 |
| TRMT12 | 702.7936988 | 0.6030335917 | 0.1020990564 | 8.82E-10 | 1.52E-08 |
| PITPNB | 1640.958909 | 0.6028576704 | 0.1063195915 | 3.56E-09 | 5.69E-08 |
| RB1CC1 | 2564.012256 | 0.602080444 | 0.1322777807 | 1.31E-06 | 1.30E-05 |
| DNAJC2 | 1654.077676 | 0.6019730508 | 0.1282876279 | 6.70E-07 | 7.15E-06 |
| PHF10 | 627.1311925 | 0.601878334 | 0.1370940151 | 2.75E-06 | 2.58E-05 |
| FAM57A | 1030.191891 | 0.601412485 | 0.1125484788 | 2.29E-08 | 3.23E-07 |
| CCDC12 | 1279.016922 | 0.6013209795 | 0.1039076181 | 1.82E-09 | 3.04E-08 |
| BACH1 | 4316.013453 | 0.6011817478 | 0.09934940976 | 3.65E-10 | 6.58E-09 |
| TNFRSF10A | 491.4438521 | 0.6002251989 | 0.190597473 | 0.000366728956 | 0.002085648368 |
| ZNF155 | 364.3764698 | 0.6001669321 | 0.1410510594 | 5.06E-06 | 4.49E-05 |
| TMEM2 | 2042.767541 | 0.5979315173 | 0.1725898473 | 0.000124288825 | 0.000815933122 |
| HSD11B1 | 498.5644844 | 0.5978280778 | 0.1142343743 | 4.23E-08 | 5.72E-07 |
| SNHG12 | 392.4996978 | 0.5964799655 | 0.1526871577 | 2.24E-05 | 0.000174861490 |
| FDX1 | 938.6825469 | 0.5958167709 | 0.1124082751 | 2.72E-08 | 3.78E-07 |
| KLHL28 | 366.294091 | 0.5955138746 | 0.1264170644 | 6.20E-07 | 6.65E-06 |
| PELO | 6007.741241 | 0.5947894738 | 0.08431318963 | 4.53E-13 | 1.23E-11 |
| SLC25A37 | 1475.170465 | 0.5947066811 | 0.1386297497 | 4.43E-06 | 3.98E-05 |
| TET3 | 376.840582 | 0.5946881924 | 0.1934030334 | 0.000477867719 | 0.002642707801 |
| NINJ1 | 1888.274104 | 0.594496113 | 0.1380816442 | 4.10E-06 | 3.72E-05 |
| KIAA1804 | 61.36299871 | 0.5944702189 | 0.2143418899 | 0.00122014535 | 0.006014557359 |
| RELL1 | 1410.046294 | 0.5942673639 | 0.07110842805 | 1.72E-17 | 7.05E-16 |
| SNAPC4 | 362.7211803 | 0.5936889286 | 0.1549864671 | 3.09E-05 | 0.00023355771 |
| TBPL1 | 778.2627827 | 0.5930068063 | 0.09745452874 | 3.03E-10 | 5.53E-09 |
| OTUD7B | 1388.124986 | 0.5901660169 | 0.0869486296 | 3.01E-12 | 7.34E-11 |
| TRIM47 | 471.6483535 | 0.5900081063 | 0.1955623753 | 0.000581967381 | 0.003138323918 |
| SS18L2 | 664.5592054 | 0.5889251929 | 0.0933470508 | 7.43E-11 | 1.51E-09 |
| SERTAD3 | 1115.114717 | 0.5886441098 | 0.1256864979 | 7.15E-07 | 7.58E-06 |

| | baseMean | log2FoldChange | lfcSE | pvalue | padj |
|---|---|---|---|---|---|
| DENND5B | 912.1968273 | 0.5881997445 | 0.1005611056 | 1.18E-09 | 2.01E-08 |
| SH3PXD2B | 2612.411728 | 0.5880541644 | 0.1001622509 | 1.14E-09 | 1.95E-08 |
| PHLPP2 | 824.5715039 | 0.5875919299 | 0.130902158 | 1.77E-06 | 1.72E-05 |
| ETF1 | 10552.57231 | 0.5873249327 | 0.1181604444 | 1.27E-07 | 1.57E-06 |
| DUSP4 | 363.9277789 | 0.5869194633 | 0.1802632843 | 0.000267429760 | 0.001588891343 |
| NPC1 | 2933.333562 | 0.5865466168 | 0.1138647227 | 6.72E-08 | 8.74E-07 |
| RPP14 | 1089.509884 | 0.586027998 | 0.08916949547 | 1.33E-11 | 2.98E-10 |
| FJX1 | 729.2051932 | 0.5859892177 | 0.2278944198 | 0.002202435574 | 0.009946030714 |
| NEDD4L | 364.0611243 | 0.5858528133 | 0.1237696831 | 5.67E-07 | 6.16E-06 |
| PTPN12 | 1322.538056 | 0.5856567374 | 0.1657203917 | 9.90E-05 | 0.000667820494 |
| CABLES1 | 770.8735197 | 0.5856251647 | 0.1435894819 | 1.15E-05 | 9.59E-05 |
| SPTY2D1 | 1416.942959 | 0.5853666508 | 0.08171732034 | 2.15E-13 | 6.06E-12 |
| PHLDA2 | 641.5068635 | 0.5846165659 | 0.1744738065 | 0.000193344040 | 0.001196466924 |
| FGF5 | 709.1501974 | 0.5843811244 | 0.281480499 | 0.007397999121 | 0.02740701889 |
| RBMS1 | 3957.571152 | 0.5842702728 | 0.1285404098 | 1.41E-06 | 1.39E-05 |
| PAIP2 | 1695.700552 | 0.5836960817 | 0.09811146078 | 7.16E-10 | 1.26E-08 |
| ATXN1L | 1240.975907 | 0.5836943741 | 0.1092943319 | 2.45E-08 | 3.43E-07 |
| RNF138 | 589.0019702 | 0.583668772 | 0.1457815981 | 1.57E-05 | 0.000127379838 |
| CBLB | 578.5852593 | 0.5835717291 | 0.1544663436 | 3.85E-05 | 0.000283642245 |
| AUTS2 | 66.78740098 | 0.5818984008 | 0.253818332 | 0.004601977593 | 0.01857382607 |
| CD82 | 2295.663231 | 0.5790549302 | 0.09895788958 | 1.31E-09 | 2.22E-08 |
| POLR3F | 573.3327658 | 0.5789473362 | 0.1050411554 | 9.54E-09 | 1.44E-07 |
| ZNF100 | 283.2669512 | 0.5785049646 | 0.1566855343 | 5.55E-05 | 0.000396061849 |
| NOV | 67.1352402 | 0.5772836047 | 0.2500703218 | 0.004466978671 | 0.01811243067 |
| CLMP | 4763.84437 | 0.5767638464 | 0.1538221854 | 4.54E-05 | 0.000329631916 |
| CAMSAP1 | 2618.246627 | 0.5764083449 | 0.06877716219 | 1.52E-17 | 6.32E-16 |
| C6orf141 | 180.3120761 | 0.5761910218 | 0.3068321854 | 0.01139643607 | 0.03940178623 |
| LARP1B | 1226.265923 | 0.5755530791 | 0.1217127891 | 5.92E-07 | 6.38E-06 |
| ATOH8 | 881.9918466 | 0.5748178383 | 0.1869670454 | 0.000506186304 | 0.002776937319 |
| MAP1S | 352.2557403 | 0.5747032774 | 0.1729974911 | 0.000221273412 | 0.001345103304 |
| PHLDB1 | 4017.304968 | 0.5746054243 | 0.1044693465 | 1.03E-08 | 1.54E-07 |
| QPCT | 197.4002308 | 0.5744819055 | 0.1486577492 | 2.88E-05 | 0.000219601501 |
| ZNF410 | 3137.592136 | 0.5735417858 | 0.08146048841 | 5.39E-13 | 1.45E-11 |
| SASH1 | 1017.427033 | 0.5725683482 | 0.08513881154 | 4.87E-12 | 1.16E-10 |
| PLEKHG3 | 347.1879674 | 0.572450126 | 0.1418920201 | 1.43E-05 | 0.000116715189 |
| PNO1 | 1983.077652 | 0.5723396302 | 0.1839569861 | 0.000454026392 | 0.002526354523 |
| ZBTB6 | 501.2689157 | 0.5719635506 | 0.1031172588 | 7.97E-09 | 1.22E-07 |
| RUNX1 | 701.1567415 | 0.571772387 | 0.1461230137 | 2.37E-05 | 0.000184319674 |
| TRAPPC2P1 | 280.8645385 | 0.5715312223 | 0.1184979753 | 3.79E-07 | 4.27E-06 |
| HIST1H2AJ | 7625.395635 | 0.5713316009 | 0.1997267525 | 0.000959589905 | 0.004859637451 |

| | baseMean | log2FoldChange | lfcSE | pvalue | padj |
|---|---|---|---|---|---|
| NOLC1 | 5594.953301 | 0.5710441516 | 0.1310796009 | 3.39E-06 | 3.12E-05 |
| SNORA61 | 186.9472882 | 0.5700410138 | 0.2646054417 | 0.006529227531 | 0.02476781368 |
| UBL3 | 762.0691491 | 0.569309171 | 0.1063101977 | 2.44E-08 | 3.42E-07 |
| ITPKB | 326.6268435 | 0.5689568682 | 0.1183260077 | 4.09E-07 | 4.54E-06 |
| TMEM87A | 2931.165589 | 0.5688243033 | 0.06493315492 | 5.59E-19 | 2.59E-17 |
| ELK1 | 1153.044575 | 0.5685046491 | 0.08526926215 | 7.37E-12 | 1.72E-10 |
| HMGXB4 | 1615.465925 | 0.5679785755 | 0.1069692964 | 2.87E-08 | 3.96E-07 |
| PVRL2 | 3675.270428 | 0.5676026627 | 0.07519476949 | 1.26E-14 | 4.01E-13 |
| RNF145 | 2913.339775 | 0.5672448823 | 0.1125372264 | 1.23E-07 | 1.53E-06 |
| GNPNAT1 | 3031.470557 | 0.5672378969 | 0.2009507209 | 0.001140014485 | 0.005667337451 |
| ZFAND5 | 3910.040851 | 0.5662296365 | 0.1334571486 | 5.91E-06 | 5.19E-05 |
| ISG20L2 | 790.2392191 | 0.5657637699 | 0.08311632426 | 2.83E-12 | 6.93E-11 |
| CNST | 1169.417387 | 0.565720988 | 0.1088970638 | 5.68E-08 | 7.49E-07 |
| RIT1 | 1666.535185 | 0.5655578823 | 0.1203806098 | 7.01E-07 | 7.46E-06 |
| KLHL29 | 500.2979095 | 0.5654337585 | 0.2787047271 | 0.008767782299 | 0.0315932988 |
| JOSD1 | 2689.435667 | 0.5652074956 | 0.08890596013 | 4.62E-11 | 9.60E-10 |
| PLEKHF2 | 698.2729016 | 0.564756949 | 0.1294598248 | 3.26E-06 | 3.01E-05 |
| GATAD2B | 282.3070528 | 0.5637708585 | 0.1969644398 | 0.001025837045 | 0.005163284206 |
| SLC20A1 | 6728.952729 | 0.5635224309 | 0.2035960893 | 0.001413819991 | 0.006821246224 |
| PPM1A | 1443.411209 | 0.5632648774 | 0.09020528588 | 1.22E-10 | 2.37E-09 |
| LYAR | 1358.800178 | 0.5630830215 | 0.1813762824 | 0.000477991276 | 0.002642707801 |
| AGAP3 | 1828.922346 | 0.5629266611 | 0.101117296 | 7.77E-09 | 1.19E-07 |
| AAR2 | 1944.892786 | 0.5625333513 | 0.0671407702 | 1.56E-17 | 6.44E-16 |
| RRN3 | 3008.790927 | 0.5622066601 | 0.1037271988 | 1.67E-08 | 2.43E-07 |
| AMMECR1L | 1216.810017 | 0.5620830793 | 0.09758269444 | 2.39E-09 | 3.94E-08 |
| TUFT1 | 409.8901019 | 0.5613733727 | 0.137735314 | 1.24E-05 | 0.000102280746 |
| FHOD1 | 1106.322884 | 0.5578110263 | 0.1236095468 | 1.82E-06 | 1.77E-05 |
| ZBTB21 | 1164.603037 | 0.5574557038 | 0.1716231266 | 0.000301126238 | 0.001764450439 |
| ZNF787 | 670.5041251 | 0.5571413069 | 0.1345873391 | 9.59E-06 | 8.10E-05 |
| NSUN2 | 2712.274038 | 0.5569457867 | 0.1414695704 | 2.23E-05 | 0.000174546926 |
| RNVU1-7 | 16846.18162 | 0.55633431 | 0.2358236958 | 0.004615147722 | 0.01860176893 |
| CNOT6L | 748.2887707 | 0.5561338384 | 0.150265727 | 5.78E-05 | 0.000411022151 |
| SPESP1 | 81.93322775 | 0.556131046 | 0.1975528048 | 0.001216511777 | 0.005999550796 |
| GOLPH3 | 2326.669984 | 0.5556617459 | 0.06783230327 | 7.66E-17 | 2.93E-15 |
| FAM65A | 302.3847822 | 0.5553922773 | 0.1553712622 | 9.39E-05 | 0.000636328318 |
| AMD1 | 5038.063336 | 0.5552029737 | 0.1527179873 | 7.37E-05 | 0.000512102233 |
| LITAF | 1613.961919 | 0.5549196254 | 0.187121594 | 0.000766636995 | 0.003999459784 |
| LOC100996763 | 383.3841116 | 0.5547027534 | 0.09356730169 | 8.82E-10 | 1.52E-08 |
| SEMA4C | 1352.305955 | 0.5537605028 | 0.1006126544 | 1.08E-08 | 1.61E-07 |
| RNU4ATAC | 32.64217456 | 0.5532994308 | 0.2758965661 | 0.009714602694 | 0.03444646809 |

| | baseMean | log2FoldChange | lfcSE | pvalue | padj |
|---|---|---|---|---|---|
| ZBTB10 | 619.3873241 | 0.5531190012 | 0.134829303 | 1.14E-05 | 9.48E-05 |
| ZDHHC18 | 1028.030802 | 0.5521306466 | 0.1077303928 | 8.59E-08 | 1.09E-06 |
| YAF2 | 381.1751197 | 0.5520051763 | 0.09589978384 | 2.54E-09 | 4.18E-08 |
| MOCS3 | 453.143241 | 0.5517309015 | 0.1255439492 | 3.09E-06 | 2.87E-05 |
| SDC4 | 6265.714947 | 0.5514894439 | 0.1161101109 | 5.82E-07 | 6.29E-06 |
| UGDH | 3756.257351 | 0.5510439436 | 0.1036816866 | 3.15E-08 | 4.32E-07 |
| GATAD2A | 2303.209461 | 0.5509589244 | 0.1172178846 | 7.43E-07 | 7.84E-06 |
| JMJD1C | 3049.806471 | 0.5507528205 | 0.1421053256 | 2.94E-05 | 0.000223435144 |
| ANXA2R | 157.7277813 | 0.5501604662 | 0.196405325 | 0.001283164689 | 0.006280420465 |
| GNL2 | 6093.675323 | 0.5496604792 | 0.1541833251 | 0.000102674739 | 0.000690792676 |
| CRY2 | 853.5112843 | 0.5495831104 | 0.119239666 | 1.15E-06 | 1.16E-05 |
| PIM2 | 369.9880177 | 0.5495747674 | 0.1491495264 | 6.28E-05 | 0.000442764300 |
| UBASH3B | 3266.042244 | 0.5495681266 | 0.131425648 | 8.60E-06 | 7.35E-05 |
| FAIM3 | 45.04860191 | 0.549008347 | 0.2933924861 | 0.01278288317 | 0.04320765643 |
| WDR26 | 1770.774562 | 0.5485154833 | 0.1449324468 | 4.27E-05 | 3.12E-04 |
| UTP15 | 1200.030519 | 0.547072444 | 0.165033696 | 0.000246945743 | 0.001481375974 |
| TRMT61A | 836.9966163 | 0.5469949035 | 0.1129851397 | 3.76E-07 | 4.24E-06 |
| LANCL2 | 946.1248837 | 0.5468919609 | 0.09862749369 | 8.43E-09 | 1.28E-07 |
| RSL24D1 | 4627.907115 | 0.5464318919 | 0.0852448065 | 4.33E-11 | 9.03E-10 |
| LOC440354 | 453.1843728 | 0.5463124176 | 0.1353358015 | 1.54E-05 | 0.000125100138 |
| DYRK1A | 1720.878867 | 0.5461348074 | 0.0912499075 | 6.85E-10 | 1.20E-08 |
| SRFBP1 | 1146.693652 | 0.5456368174 | 0.1864303253 | 0.000892090938 | 0.004559677989 |
| HIST2H2AC | 5779.406469 | 0.5449056327 | 0.1758977038 | 0.000518766465 | 0.002839674229 |
| NUP153 | 4444.234637 | 0.5447321365 | 0.1331871202 | 1.20E-05 | 9.90E-05 |
| DDI2 | 319.4207613 | 0.5445780377 | 0.1635719776 | 0.000238188851 | 0.00143520591 |
| FLVCR2 | 134.7066164 | 0.544506834 | 0.1723726025 | 0.000425324736 | 0.002389683021 |
| FBXO36 | 237.1233032 | 0.5444060354 | 0.1360025025 | 1.77E-05 | 0.000141915661 |
| MRPL50 | 1283.879626 | 0.5442400429 | 0.1102667542 | 2.34E-07 | 2.72E-06 |
| SNX30 | 1376.659992 | 0.543394683 | 0.2215767022 | 0.003511943284 | 0.01483806003 |
| RPPH1 | 236583.3928 | 0.5433348377 | 0.2247136651 | 0.003837717574 | 0.0159697532 |
| PPAN | 1182.4458 | 0.5430441029 | 0.1253334628 | 4.26E-06 | 3.85E-05 |
| CCNL1 | 1762.329381 | 0.5430095029 | 0.1151471083 | 7.04E-07 | 7.48E-06 |
| CCNJL | 97.74550526 | 0.5429739347 | 0.2046874289 | 0.002038623702 | 0.009333661839 |
| FERMT2 | 7982.073148 | 0.5423051954 | 0.1495525704 | 8.05E-05 | 0.000554040622 |
| WDR66 | 198.8611344 | 0.5416062022 | 0.1891235518 | 0.001096881779 | 0.005469347891 |
| ZNF20 | 316.1422514 | 0.5414092616 | 0.126412787 | 5.32E-06 | 4.70E-05 |
| ANKRD13C | 1519.701904 | 0.5413175035 | 0.09084608304 | 7.76E-10 | 1.35E-08 |
| DCAF12 | 1046.984197 | 0.5401982156 | 0.08808075618 | 2.64E-10 | 4.88E-09 |
| REXO1 | 423.0764742 | 0.5400494873 | 0.1441540816 | 5.25E-05 | 0.000376325297 |
| LRP12 | 1039.227277 | 0.5398548042 | 0.09930696856 | 1.72E-08 | 2.48E-07 |

| | baseMean | log2FoldChange | lfcSE | pvalue | padj |
|---|---|---|---|---|---|
| TSHZ1 | 365.9003171 | 0.5394704218 | 0.1151418154 | 8.20E-07 | 8.58E-06 |
| SLC39A14 | 4632.227436 | 0.5387353915 | 0.1543189473 | 0.000136340261 | 0.000882595716 |
| NNMT | 20257.06275 | 0.5378265224 | 0.1766636224 | 0.000613840337 | 0.003289945565 |
| CCDC19 | 49.8253907 | 0.5373192606 | 0.2704571769 | 0.01071247207 | 0.03742327467 |
| FOSL1 | 1733.59576 | 0.537187016 | 0.2347669656 | 0.005395657149 | 0.02112991565 |
| ATP2B1 | 4966.55824 | 0.5367199841 | 0.1718169551 | 0.000490672439 | 0.002704784137 |
| GLA | 1387.255394 | 0.5363790636 | 0.08495339927 | 8.34E-11 | 1.68E-09 |
| SIX1 | 443.3491622 | 0.5358525579 | 0.125837184 | 6.07E-06 | 5.32E-05 |
| VCPKMT | 213.2228018 | 0.5352328476 | 0.1531176354 | 1.35E-04 | 8.78E-04 |
| TMEM189 | 1518.804464 | 0.5344742534 | 0.1232584533 | 4.32E-06 | 3.90E-05 |
| PARP8 | 600.1925839 | 0.534440483 | 0.1168376848 | 1.43E-06 | 1.41E-05 |
| TRMT6 | 941.8574983 | 0.5344133903 | 0.1778559161 | 0.000725904374 | 0.003809654646 |
| ZNF669 | 303.6695668 | 0.5342492046 | 0.1213165857 | 3.18E-06 | 2.94E-05 |
| TLE4 | 837.1827685 | 0.5340177452 | 0.1266561369 | 7.36E-06 | 6.37E-05 |
| SCAF4 | 329.56601 | 0.5339740169 | 0.1458357082 | 7.28E-05 | 0.000506211556 |
| RNF103 | 468.0009398 | 0.5339478961 | 0.1100264508 | 3.64E-07 | 4.12E-06 |
| MAP1LC3B | 2488.042329 | 0.5333943906 | 0.08261234292 | 3.34E-11 | 7.11E-10 |
| ZNF791 | 537.572156 | 0.5333775184 | 0.1140635849 | 8.80E-07 | 9.12E-06 |
| TMEM243 | 430.1845092 | 0.533309488 | 0.1186696604 | 2.09E-06 | 2.00E-05 |
| LOC101929758 | 96.93484447 | 0.5330334827 | 0.2209626532 | 0.004016241228 | 0.0165874662 |
| SIRT1 | 771.5765308 | 0.5329127904 | 0.1117683351 | 5.65E-07 | 6.13E-06 |
| SCARNA6 | 976.291974 | 0.5326478492 | 0.1411861745 | 4.54E-05 | 0.000329815317 |
| ZNF398 | 761.8279969 | 0.5326094515 | 0.09755453424 | 1.48E-08 | 2.16E-07 |
| CDC42EP4 | 1177.281106 | 0.5325898631 | 0.07784006865 | 2.45E-12 | 6.04E-11 |
| C16orf52 | 830.0909291 | 0.5324640774 | 0.1593831352 | 0.000235078975 | 0.001419339362 |
| IRF2BP2 | 2119.057028 | 0.5321367668 | 0.09244619164 | 2.68E-09 | 4.39E-08 |
| IL17D | 60.67725984 | 0.5317784126 | 0.2314800802 | 0.005442705258 | 0.02126940645 |
| CXorf40B | 978.3882838 | 0.5314125764 | 0.08284679843 | 4.44E-11 | 9.23E-10 |
| TMEM115 | 2080.364645 | 0.5312314437 | 0.1010189628 | 4.44E-08 | 5.99E-07 |
| 7-Mar | 3867.855929 | 0.5309616305 | 0.09256382504 | 3.04E-09 | 4.92E-08 |
| RELA | 1251.571017 | 0.5308074428 | 0.07618313539 | 1.03E-12 | 2.65E-11 |
| MAP7D1 | 3107.569253 | 0.5298242611 | 0.1037309128 | 1.02E-07 | 1.29E-06 |
| DCUN1D3 | 783.805579 | 0.5297900254 | 0.07979258154 | 9.98E-12 | 2.29E-10 |
| NSRP1 | 821.6941233 | 0.52947639 | 0.1060742362 | 1.84E-07 | 2.20E-06 |
| MTHFD2L | 97.11233502 | 0.5294529103 | 0.2020862639 | 0.002329539523 | 0.01041576996 |
| NT5C3A | 524.432915 | 0.5292740469 | 0.1558046592 | 0.000195494160 | 0.001208767672 |
| CRTC2 | 416.5277156 | 0.5286144434 | 0.1092544158 | 4.01E-07 | 4.46E-06 |
| TMEM206 | 624.8996255 | 0.526603402 | 0.1396063318 | 4.80E-05 | 0.000345787474 |
| GPC1 | 4262.319204 | 0.5263708118 | 0.1638977808 | 0.000377619806 | 0.002139414497 |
| EXOSC4 | 785.1545198 | 0.5258145115 | 0.104517988 | 1.52E-07 | 1.85E-06 |

| | baseMean | log2FoldChange | lfcSE | pvalue | padj |
|---|---|---|---|---|---|
| N4BP2 | 578.7649782 | 0.5249296394 | 0.1538566203 | 0.000186671694 | 0.001161452096 |
| DNAJB4 | 5160.181122 | 0.5248728083 | 0.1892088861 | 0.001513978091 | 0.007245779428 |
| PPP3CC | 914.2337913 | 0.5247122136 | 0.1071979863 | 3.22E-07 | 3.67E-06 |
| RMND5A | 1385.826447 | 0.5246170099 | 0.1150345308 | 1.58E-06 | 1.55E-05 |
| PRTG | 172.566764 | 0.5239416403 | 0.1710001768 | 0.000618208071 | 0.003305899859 |
| HPS6 | 1717.557538 | 0.5232655328 | 0.07764490569 | 5.18E-12 | 1.23E-10 |
| GLIS3 | 866.1777158 | 0.5231977831 | 0.1149915914 | 1.66E-06 | 1.63E-05 |
| AHCTF1P1 | 43.81143475 | 0.5223215549 | 0.2566913692 | 0.01021131324 | 0.03592538818 |
| FAM174A | 240.1959363 | 0.5221422311 | 0.1766693691 | 0.000881112138 | 0.004508205713 |
| BCL10 | 1158.408209 | 0.5220277368 | 0.09227856371 | 4.93E-09 | 7.75E-08 |
| LINC00984 | 836.3712093 | 0.5217347792 | 0.1778660197 | 0.000950370309 | 0.00482278921 |
| ZNF804A | 399.2688224 | 0.521723477 | 0.1834441292 | 0.001252038785 | 0.006152345042 |
| RAE1 | 2121.126741 | 0.5210396045 | 0.1049641465 | 2.18E-07 | 2.55E-06 |
| CASP3 | 2066.393534 | 0.5209578341 | 0.09543355801 | 1.50E-08 | 2.19E-07 |
| LAMB3 | 114.1021223 | 0.5208465688 | 0.2008568054 | 0.002596181027 | 0.0114498043 |
| PIK3CA | 1487.212105 | 0.5202467687 | 0.1075563539 | 4.17E-07 | 4.61E-06 |
| DCAF4 | 695.0967969 | 0.5201537294 | 0.106994195 | 3.70E-07 | 4.18E-06 |
| CHMP4B | 2135.21022 | 0.519125876 | 0.09651099712 | 2.60E-08 | 3.62E-07 |
| LOC100507507 | 586.7437761 | 0.518875627 | 0.1014017797 | 9.86E-08 | 1.25E-06 |
| RNU1-2 | 12888.87207 | 0.5185116394 | 0.2296957129 | 0.00618994506 | 0.02366340532 |
| RNU1-27P | 12888.87207 | 0.5185116394 | 0.2296957129 | 0.00618994506 | 0.02366340532 |
| RNU1-4 | 12888.87207 | 0.5185116394 | 0.2296957129 | 0.00618994506 | 0.02366340532 |
| ARL8A | 900.3607104 | 0.5181356103 | 0.09940353982 | 5.98E-08 | 7.85E-07 |
| MAP3K2 | 1881.264271 | 0.5177132166 | 0.1301281931 | 2.14E-05 | 0.000168379752 |
| GNG11 | 4431.785714 | 0.5173482912 | 0.1150280457 | 2.19E-06 | 2.08E-05 |
| EIF1AD | 921.0500633 | 0.5159208636 | 0.08789374125 | 1.43E-09 | 2.41E-08 |
| ATP6V1G1 | 3211.465329 | 0.5158389819 | 0.07853742421 | 1.68E-11 | 3.72E-10 |
| NPAS2 | 330.4121924 | 0.5156923338 | 0.1387027745 | 6.17E-05 | 0.000435578839 |
| TRIM62 | 168.2862771 | 0.5155679594 | 0.2311656837 | 0.006717577753 | 0.02534026227 |
| BTN2A2 | 386.2748746 | 0.5132265544 | 0.1063705794 | 4.52E-07 | 4.99E-06 |
| ALAS1 | 2980.569812 | 0.5129656021 | 0.06726790796 | 8.12E-15 | 2.63E-13 |
| CARD6 | 287.3160729 | 0.5129395681 | 0.1367281632 | 5.43E-05 | 0.000387809846 |
| VDR | 180.8939686 | 0.5124236328 | 0.156055698 | 0.000311878458 | 0.001817439674 |
| MED6 | 1128.688004 | 0.511910337 | 0.105210464 | 3.86E-07 | 4.32E-06 |
| ABTB2 | 99.68185082 | 0.51142591 | 0.224855549 | 0.006178626243 | 0.02364873165 |
| TNFRSF10D | 8758.285307 | 0.5112970021 | 0.1660459295 | 0.000627868244 | 0.003350656019 |
| CYLD | 1440.488034 | 0.511197604 | 0.1005446241 | 1.21E-07 | 1.50E-06 |
| TEX10 | 2301.980141 | 0.5103806081 | 0.1270218095 | 1.86E-05 | 0.000148188699 |
| ZNF200 | 463.95719 | 0.5103070765 | 0.1190523034 | 5.82E-06 | 5.12E-05 |
| SNHG10 | 333.1868543 | 0.5102635713 | 0.1520308846 | 0.000240066388 | 0.001445349156 |

| | baseMean | log2FoldChange | lfcSE | pvalue | padj |
|---|---|---|---|---|---|
| WNT5A | 4169.86959 | 0.5100662473 | 0.1016587706 | 1.71E-07 | 2.07E-06 |
| YES1 | 2696.624723 | 0.5093778386 | 0.08699497471 | 1.59E-09 | 2.67E-08 |
| SKI | 615.5826992 | 0.508510861 | 0.1540498934 | 0.000295952276 | 0.001738097097 |
| GAN | 480.014737 | 0.5084643088 | 0.1387330703 | 7.78E-05 | 0.000536969126 |
| SLC6A6 | 2206.272975 | 0.5082850402 | 0.1684045878 | 0.000762010656 | 0.003978112435 |
| CREB3L2 | 2334.494076 | 0.5075223023 | 0.1252448347 | 1.70E-05 | 0.000136926177 |
| RAB35 | 2874.399591 | 0.507038148 | 0.06299022779 | 2.87E-16 | 1.05E-14 |
| NCOA3 | 2641.599169 | 0.50700331 | 0.07420900679 | 2.85E-12 | 6.96E-11 |
| SOCS2 | 942.9466412 | 0.5069975169 | 0.2330630467 | 0.007892483153 | 0.02889382387 |
| CMC2 | 1203.900376 | 0.5063720472 | 0.1112067458 | 1.72E-06 | 1.67E-05 |
| ELL2 | 4161.806373 | 0.506015962 | 0.09671522666 | 4.38E-08 | 5.92E-07 |
| BICD1 | 243.051092 | 0.5055524011 | 0.1630767307 | 0.000591208036 | 0.003182392067 |
| FASTKD3 | 461.9608113 | 0.5039806285 | 0.1521302648 | 0.000288196301 | 0.001697371333 |
| MTF1 | 607.59339 | 0.5039491797 | 0.1235125319 | 1.43E-05 | 0.000116853402 |
| ZBTB49 | 218.4686457 | 0.5036640728 | 0.1681331368 | 0.000831253377 | 0.004288472466 |
| ARHGEF40 | 2881.162012 | 0.5033785163 | 0.1631073382 | 0.000634057287 | 0.003380049751 |
| SPIRE1 | 2105.418758 | 0.5032325372 | 0.116068297 | 4.73E-06 | 4.24E-05 |
| ZRANB1 | 2860.583021 | 0.5026588687 | 0.08325583001 | 5.35E-10 | 9.47E-09 |
| OTUD3 | 536.1623868 | 0.5021974499 | 0.1101832695 | 1.71E-06 | 1.67E-05 |
| SCARF2 | 582.2886255 | 0.5020168683 | 0.1755239616 | 0.001275107622 | 0.006245091244 |
| CDYL2 | 64.27021692 | 0.5007356882 | 0.2205259107 | 0.006468361851 | 0.02456195858 |
| SMIM12 | 1887.687646 | 0.5004192714 | 0.1000772479 | 1.77E-07 | 2.12E-06 |
| ZNF282 | 1088.143527 | 0.5003251613 | 0.08955861597 | 7.89E-09 | 1.21E-07 |
| SLC19A1 | 380.1533827 | 0.4994372279 | 0.2282515986 | 0.007857602957 | 0.02879445002 |
| ASPHD1 | 146.6618418 | 0.4993256803 | 0.182665182 | 0.001867239066 | 0.008666247648 |
| HIST1H2AH | 6593.166551 | 0.4992410164 | 0.2040132616 | 0.004057916229 | 0.01672710818 |
| YAE1D1 | 474.5136041 | 0.4991558849 | 0.1032594524 | 4.46E-07 | 4.92E-06 |
| PDCD1LG2 | 484.3847796 | 0.4983956123 | 0.1991579453 | 0.003599610938 | 0.01514399753 |
| USP38 | 1437.838806 | 0.4976809625 | 0.1046860141 | 7.46E-07 | 7.86E-06 |
| NFKBIB | 656.4426662 | 0.4970850797 | 0.1320214117 | 5.67E-05 | 4.03E-04 |
| ZNF777 | 397.6228698 | 0.4968706045 | 0.1038311451 | 5.85E-07 | 6.31E-06 |
| NFKBID | 76.82845197 | 0.4960404765 | 0.2218883246 | 0.007099703094 | 0.02648646439 |
| ZBTB43 | 787.6412654 | 0.4953698071 | 0.09124458454 | 1.96E-08 | 2.80E-07 |
| PNRC2 | 4116.89698 | 0.4952728838 | 0.08930669747 | 1.01E-08 | 1.52E-07 |
| SDHAF2 | 1347.717977 | 0.4948666471 | 0.07455669325 | 1.13E-11 | 2.55E-10 |
| SNORD3B-2 | 29588.28754 | 0.4943889996 | 0.1334932541 | 7.37E-05 | 0.000512102233 |
| FAM118A | 515.1303984 | 0.493909703 | 0.1355907918 | 8.86E-05 | 0.000603846063 |
| NGDN | 1217.942225 | 0.4936176882 | 0.1059355979 | 1.08E-06 | 1.10E-05 |
| RBM3 | 9372.150164 | 0.493597896 | 0.0760583408 | 3.03E-11 | 6.53E-10 |
| CLDND1 | 2027.941501 | 0.4934844661 | 0.07470457761 | 1.40E-11 | 3.12E-10 |

| | baseMean | log2FoldChange | lfcSE | pvalue | padj |
|---|---|---|---|---|---|
| TTC39C | 461.3158407 | 0.4933980257 | 0.2081006381 | 0.005156874164 | 0.02041496927 |
| ZNF707 | 141.0205645 | 0.4932526095 | 0.1889597314 | 0.002725015657 | 0.01192965543 |
| MIR4435-1HG | 2468.504956 | 0.4927741353 | 0.1078033482 | 1.67E-06 | 1.63E-05 |
| RPUSD1 | 929.4190344 | 0.4922834396 | 0.1177156074 | 9.80E-06 | 8.24E-05 |
| ABI3 | 65.02511685 | 0.4920047887 | 0.2216226238 | 0.007532875344 | 0.02778225935 |
| GRK5 | 2512.995351 | 0.4917083186 | 0.2123812001 | 0.005829195686 | 0.02254894637 |
| VPS37C | 684.1407719 | 0.4913346168 | 0.1072523086 | 1.59E-06 | 1.56E-05 |
| MFHAS1 | 911.514081 | 0.4911923921 | 0.09123848758 | 2.56E-08 | 3.56E-07 |
| RN7SK | 1330173.882 | 0.4911898879 | 0.1854363428 | 0.00248386614 | 0.01103290064 |
| SH3GL1 | 4127.06844 | 0.491113062 | 0.09027697903 | 1.87E-08 | 2.68E-07 |
| NEDD4 | 7874.247198 | 0.4911049101 | 0.1285082139 | 4.43E-05 | 0.000322392047 |
| FCHO2 | 534.8737565 | 0.4910308552 | 0.1378262165 | 0.000121656671 | 0.000800771960 |
| CTSL | 5128.181541 | 0.4906180422 | 0.05345083299 | 1.59E-20 | 8.24E-19 |
| FOXJ3 | 1051.688756 | 0.4904050007 | 0.09472976883 | 7.92E-08 | 1.01E-06 |
| ZNF746 | 926.7317352 | 0.489335743 | 0.1274974994 | 4.17E-05 | 0.000305510602 |
| VIMP | 2160.266169 | 0.4892747694 | 0.07739596355 | 9.41E-11 | 1.87E-09 |
| TNIP1 | 4737.050142 | 0.4888412583 | 0.1005835242 | 4.08E-07 | 4.54E-06 |
| NET1 | 1878.731174 | 0.487674372 | 0.1474982651 | 0.000313177611 | 0.001821445884 |
| PSTK | 115.0294219 | 0.4873708362 | 0.1960389491 | 0.003905164631 | 0.01620061192 |
| KIAA1217 | 105.1847473 | 0.4873106812 | 0.2235311996 | 0.008400297672 | 0.03047564134 |
| AFAP1 | 2827.683667 | 0.4867890241 | 0.1290228788 | 5.52E-05 | 0.000393970543 |
| TAF5L | 1021.655995 | 0.4865358286 | 0.1103999499 | 3.64E-06 | 3.33E-05 |
| SPAG1 | 119.1045901 | 0.4862319869 | 0.1758397764 | 0.00179212492 | 0.00835408514 |
| GLRX | 2848.456523 | 0.485139323 | 0.07761905714 | 1.49E-10 | 2.88E-09 |
| PMPCA | 3665.744925 | 0.4846186608 | 0.06705358022 | 1.81E-13 | 5.14E-12 |
| SGK223 | 142.89403 | 0.4838066366 | 0.1620120667 | 0.000927129689 | 0.004712882878 |
| ZFAND2A | 644.8459442 | 0.4837992492 | 0.07779120863 | 1.82E-10 | 3.45E-09 |
| SNORA74A | 292.7852858 | 0.483557948 | 0.2004121049 | 0.004790616505 | 0.01919986249 |
| MUL1 | 1341.653026 | 0.4828331113 | 0.07746177861 | 1.66E-10 | 3.18E-09 |
| UBE2D1 | 333.9930218 | 0.4828081982 | 0.1432163675 | 0.000251134280 | 0.001501059136 |
| PHRF1 | 2169.036168 | 0.4823917344 | 0.09846770091 | 3.41E-07 | 3.88E-06 |
| SCARNA16 | 118.5102226 | 0.4816853182 | 0.1906406563 | 0.003569094278 | 0.01504536939 |
| CKMT2-AS1 | 184.3220613 | 0.4811237075 | 0.1820571731 | 0.002602432173 | 0.01146718337 |
| PPRC1 | 1276.122101 | 0.480910361 | 0.1318094563 | 9.06E-05 | 0.000616680194 |
| RAB18 | 2207.800429 | 0.4808275605 | 0.08410990214 | 4.05E-09 | 6.43E-08 |
| SREK1IP1 | 1039.310734 | 0.4805915521 | 0.1139779012 | 8.76E-06 | 7.47E-05 |
| C7orf26 | 601.5600401 | 0.4803666529 | 0.08866873624 | 2.19E-08 | 3.11E-07 |
| TOP1 | 2707.537549 | 0.4802742214 | 0.1619043951 | 0.000993907562 | 0.005021475974 |
| MAPK1IP1L | 2985.734806 | 0.4800952349 | 0.09806379544 | 3.85E-07 | 4.32E-06 |
| ZNFX1 | 2936.268583 | 0.4790304311 | 0.1153313673 | 1.16E-05 | 9.59E-05 |

| | baseMean | log2FoldChange | lfcSE | pvalue | padj |
|---|---|---|---|---|---|
| SLC7A5 | 2631.814655 | 0.4789528396 | 0.1130752508 | 8.07E-06 | 6.93E-05 |
| C9orf72 | 327.9365109 | 0.4788282798 | 0.1090762757 | 4.06E-06 | 3.68E-05 |
| STK10 | 2020.458125 | 0.4782820735 | 0.09151415112 | 6.31E-08 | 8.23E-07 |
| PRPSAP1 | 872.062917 | 0.4781132355 | 0.1010676361 | 8.00E-07 | 8.38E-06 |
| ZNF845 | 639.2163882 | 0.4771541947 | 0.1456958872 | 0.000359563225 | 0.00205273653 |
| EHD4 | 2698.727247 | 0.4769681562 | 0.1822310036 | 0.002851928253 | 0.01243043318 |
| GEMIN7 | 931.3271134 | 0.4767715197 | 0.09520448348 | 2.01E-07 | 2.38E-06 |
| HIST1H4E | 17290.04804 | 0.4766929333 | 0.1570571867 | 0.000811662989 | 0.004203426175 |
| ZNF579 | 141.4173503 | 0.4762187234 | 0.186850828 | 0.00344499308 | 0.01460249436 |
| TMEM201 | 292.8501076 | 0.4759240345 | 0.1491356019 | 0.000483375127 | 0.002668510297 |
| MIEF1 | 1660.979634 | 0.475500859 | 0.07792721656 | 3.95E-10 | 7.10E-09 |
| BTG1 | 3008.908327 | 0.4754314215 | 0.1118747999 | 8.26E-06 | 7.08E-05 |
| BZW1 | 17866.79252 | 0.474946055 | 0.1484526727 | 0.000470659918 | 0.002606045192 |
| RRS1 | 866.54837 | 0.4747217199 | 0.1598962811 | 0.001003545586 | 0.005063297264 |
| SH3BP5 | 486.9499214 | 0.4747163592 | 0.1528958418 | 0.000644774273 | 0.003427363143 |
| DGKH | 359.0833696 | 0.4745163247 | 0.1684048551 | 0.001603984883 | 0.007610494241 |
| FAM131A | 449.5410524 | 0.4744752324 | 0.1203093887 | 2.86E-05 | 0.000218209629 |
| ZNF316 | 382.9431976 | 0.4742728976 | 0.1309375587 | 0.000102894047 | 0.000691955499 |
| PLEKHH3 | 327.0719644 | 0.4742681622 | 0.2061520887 | 0.006613777446 | 0.02502484008 |
| ZBTB1 | 1696.76698 | 0.4731657148 | 0.1028054692 | 1.52E-06 | 1.50E-05 |
| RCC2 | 4640.995479 | 0.4727453133 | 0.07608162066 | 1.97E-10 | 3.71E-09 |
| ARF6 | 3965.352839 | 0.4726215606 | 0.117541456 | 2.08E-05 | 0.000164491455 |
| SCARNA13 | 4443.335064 | 0.4725964837 | 0.1582514148 | 0.000950780484 | 0.004823226792 |
| PDP2 | 405.1404004 | 0.4725606521 | 0.1382248417 | 0.000220377423 | 0.001341455218 |
| MIR22HG | 1628.735109 | 0.4722691698 | 0.138479944 | 0.000225807567 | 0.001367242321 |
| RN7SL1 | 3866187.732 | 0.4719909212 | 0.1681428955 | 0.001676708363 | 0.007902662493 |
| HDAC5 | 1714.532287 | 0.470957669 | 0.1846645672 | 0.003497240101 | 0.01478852822 |
| MIR17HG | 340.2065759 | 0.4709433186 | 0.2303756263 | 0.01218853115 | 0.0416609367 |
| USP16 | 3178.897594 | 0.4700413753 | 0.1171864231 | 2.15E-05 | 0.000169049742 |
| SAMD8 | 1200.027073 | 0.4697987217 | 0.1302660083 | 0.000110278777 | 0.000735636522 |
| ATXN1 | 554.1460375 | 0.4692362459 | 0.2383813748 | 0.01448653874 | 0.04798444389 |
| GPR176 | 7404.914286 | 0.4691504663 | 0.1526212993 | 0.000745440168 | 0.003897071159 |
| SNX18 | 2683.601837 | 0.4689128943 | 0.116388775 | 2.05E-05 | 0.000162017382 |
| RP9 | 616.7065773 | 0.4686320391 | 0.1013931696 | 1.41E-06 | 1.39E-05 |
| C1orf52 | 677.1884029 | 0.4683205725 | 0.1186159765 | 2.86E-05 | 0.000218546458 |
| RNF4 | 3663.952302 | 0.468289189 | 0.09568979691 | 3.70E-07 | 4.18E-06 |
| HIST1H2AK | 2120.569336 | 0.4676263293 | 0.1952931215 | 0.005391477137 | 0.02111910105 |
| KYNU | 228.8508121 | 0.4674416352 | 0.166642314 | 0.001706680231 | 0.008018542744 |
| SCARNA7 | 4661.277134 | 0.4668854333 | 0.1264568185 | 6.75E-05 | 0.000472166558 |
| B4GALT5 | 1954.79461 | 0.4661321558 | 0.1158681999 | 2.04E-05 | 0.000161617949 |

| | baseMean | log2FoldChange | lfcSE | pvalue | padj |
|---|---|---|---|---|---|
| MBIP | 520.9359509 | 0.4658970013 | 0.08742656642 | 3.74E-08 | 5.08E-07 |
| HAUS6 | 2512.787067 | 0.4657194966 | 0.1756176404 | 0.002682167504 | 0.01178017462 |
| ID3 | 5431.369978 | 0.4653312251 | 0.1714127775 | 0.002456853513 | 0.01092921779 |
| PWP1 | 4151.035926 | 0.4650391257 | 0.11789489 | 2.93E-05 | 0.000222925775 |
| SMN1 | 1570.467888 | 0.464371991 | 0.1153644011 | 2.11E-05 | 1.66E-04 |
| ZNF486 | 446.7137033 | 0.4643137312 | 0.1039730205 | 2.98E-06 | 2.78E-05 |
| ZNF876P | 112.6239027 | 0.4639761834 | 0.1648687599 | 0.001682130379 | 0.00792320127 |
| RUSC2 | 1625.314976 | 0.4636320173 | 0.09772132392 | 7.23E-07 | 7.64E-06 |
| TNC | 2863.298297 | 0.4635344785 | 0.2321014282 | 0.01412709051 | 0.04697147178 |
| HIST1H3B | 16434.85827 | 0.4620328425 | 0.188971833 | 0.004932932113 | 0.01967490657 |
| SLC7A6 | 1234.855472 | 0.4619106066 | 0.1626427915 | 0.001568480135 | 0.007472992233 |
| TP53RK | 1179.879308 | 0.4609336348 | 0.1010259303 | 1.92E-06 | 1.86E-05 |
| ZNF304 | 605.7477601 | 0.4608503152 | 0.1154413632 | 2.45E-05 | 0.000189828759 |
| PALLD | 3552.914361 | 0.4605729062 | 0.1609812882 | 0.001479647727 | 0.007108898034 |
| SAV1 | 788.7329315 | 0.459972742 | 0.09934350233 | 1.43E-06 | 1.42E-05 |
| SGPP1 | 569.1738645 | 0.4598167478 | 0.1062518286 | 5.74E-06 | 5.05E-05 |
| NDFIP2 | 1199.505269 | 0.4594779122 | 0.1019330221 | 2.52E-06 | 2.37E-05 |
| ATP13A3 | 4083.376069 | 0.4592422995 | 0.1817964117 | 0.003909260747 | 0.01621308726 |
| A4GALT | 191.4143772 | 0.4590700709 | 0.1656734648 | 0.001953599868 | 0.008999736521 |
| RNF149 | 862.9412523 | 0.4586147113 | 0.08355053337 | 1.58E-08 | 2.29E-07 |
| ST3GAL2 | 1252.324858 | 0.4584780935 | 0.1287413761 | 0.000139797268 | 0.000900667037 |
| HIST1H2AL | 6388.717532 | 0.4582939527 | 0.2031473634 | 0.007610915232 | 0.0280214928 |
| ZSCAN25 | 572.7712758 | 0.4577844614 | 0.1086093242 | 9.48E-06 | 8.01E-05 |
| AHCTF1 | 4120.826045 | 0.4576339812 | 0.1517797189 | 0.000894430969 | 0.004570069561 |
| ATP1B1 | 1197.333421 | 0.4575556988 | 0.1408642487 | 0.000422806116 | 0.002377326387 |
| TWIST2 | 464.1690579 | 0.4573694415 | 0.1025655909 | 3.14E-06 | 2.90E-05 |
| FBXL14 | 205.2257674 | 0.456616379 | 0.1663472242 | 0.002116240103 | 0.009617354053 |
| H1FX | 1476.703361 | 0.4554921106 | 0.1598852061 | 0.001562088173 | 0.007447304131 |
| IER5 | 1976.719982 | 0.455264959 | 0.1063585607 | 7.17E-06 | 6.21E-05 |
| PRKCD | 694.5382482 | 0.455042803 | 0.08917861108 | 1.31E-07 | 1.62E-06 |
| H3F3B | 22083.30112 | 0.4549455084 | 0.07879976446 | 2.54E-09 | 4.18E-08 |
| PIP5K1A | 3489.338806 | 0.4543707839 | 0.08347971241 | 2.08E-08 | 2.96E-07 |
| SDF2 | 1987.498695 | 0.4542864987 | 0.06279261038 | 1.88E-13 | 5.33E-12 |
| CHST3 | 976.0543373 | 0.4537352769 | 0.1201535261 | 6.04E-05 | 0.000427387078 |
| EMC3-AS1 | 184.7247474 | 0.4534233046 | 0.2076758162 | 0.009533989231 | 0.03387865529 |
| ZMYND19 | 999.0900436 | 0.4532325474 | 0.1007119253 | 2.69E-06 | 2.53E-05 |
| KCTD13 | 111.1064493 | 0.4530871127 | 0.1764404965 | 0.00357675731 | 0.0150654489 |
| CASP7 | 1482.247493 | 0.4529280685 | 0.07831163004 | 2.93E-09 | 4.74E-08 |
| GNL3 | 6829.020213 | 0.4521364786 | 0.16050244 | 0.001770996713 | 0.00826655261 |
| UHRF1BP1L | 1501.16261 | 0.4521056528 | 0.1213555755 | 7.43E-05 | 0.000515630797 |

| | baseMean | log2FoldChange | lfcSE | pvalue | padj |
|---|---|---|---|---|---|
| MCM9 | 335.6202748 | 0.4519046534 | 0.121563469 | 7.64E-05 | 0.000527974051 |
| KLHL42 | 1431.862007 | 0.4516439625 | 0.1297458307 | 0.000187593725 | 0.001165727455 |
| CDR2L | 897.2507203 | 0.4514708016 | 0.1027300938 | 4.38E-06 | 3.95E-05 |
| LOC493754 | 589.0055178 | 0.4506874368 | 0.09877318103 | 1.98E-06 | 1.91E-05 |
| ATG4D | 486.5809595 | 0.4506171537 | 0.1094115835 | 1.47E-05 | 0.000120224258 |
| SRSF6 | 6040.758479 | 0.4504062965 | 0.08569992902 | 5.83E-08 | 7.68E-07 |
| CCDC9 | 362.533727 | 0.4501516525 | 0.131968244 | 0.000243903656 | 0.001465488916 |
| CLDN12 | 1002.643451 | 0.4497563759 | 0.1538980221 | 0.001269228437 | 0.006222437341 |
| C16orf72 | 2115.120466 | 0.4496754338 | 0.06767620932 | 1.24E-11 | 2.78E-10 |
| ACBD3 | 3166.545543 | 0.4495797627 | 0.07555108152 | 1.08E-09 | 1.85E-08 |
| ZNF468 | 604.1839426 | 0.4491475511 | 0.1127744909 | 2.60E-05 | 0.000200311141 |
| CEP135 | 505.2383593 | 0.4488397594 | 0.1647428178 | 0.002320728769 | 0.01037949253 |
| SHROOM2 | 81.25160133 | 0.4488368357 | 0.2252828729 | 0.01489782172 | 0.04909554397 |
| HMCES | 1289.894836 | 0.4487261898 | 0.08581328616 | 6.82E-08 | 8.83E-07 |
| AMOTL1 | 4054.41615 | 0.4486945596 | 0.1054552036 | 7.51E-06 | 6.48E-05 |
| HGS | 4520.399017 | 0.4480296624 | 0.1040895766 | 6.60E-06 | 5.75E-05 |
| ASAP1 | 2593.330739 | 0.4476181908 | 0.1348208857 | 3.17E-04 | 0.00184130458 |
| SNCA | 287.7498974 | 0.4470084196 | 0.1685127025 | 0.002867633248 | 0.01248175651 |
| GPRIN1 | 679.3861044 | 0.4469045743 | 0.1756085983 | 0.003886625179 | 0.01613718971 |
| LLPH | 1437.217617 | 0.4468044039 | 0.1123052966 | 2.71E-05 | 0.000208138800 |
| POM121 | 385.1684815 | 0.4461642311 | 0.1216091773 | 9.47E-05 | 0.000641271444 |
| INF2 | 1309.323281 | 0.4459604082 | 0.1146141554 | 3.88E-05 | 0.000285849137 |
| C12orf29 | 1017.991863 | 0.4456860413 | 0.1470607484 | 0.000921673919 | 0.004691556575 |
| RLIM | 3828.951915 | 0.4451979015 | 0.1214384627 | 0.000103584813 | 0.000695658227 |
| PYGO1 | 419.2087514 | 0.4449998684 | 0.1439844423 | 0.000764369609 | 0.003989028782 |
| SYNJ2 | 2122.746187 | 0.4449460855 | 0.143190287 | 0.000705920868 | 0.003716568533 |
| PISD | 1618.490928 | 0.4448345663 | 0.09001785224 | 3.15E-07 | 3.60E-06 |
| RNPS1 | 4666.463018 | 0.4445793294 | 0.1125535067 | 3.09E-05 | 0.000233811908 |
| SLU7 | 1807.829209 | 0.4443833739 | 0.09349467599 | 8.21E-07 | 8.58E-06 |
| SIAH2 | 1331.935154 | 0.4441874501 | 0.07977998979 | 1.06E-08 | 1.58E-07 |
| RANBP9 | 1166.274701 | 0.4439026944 | 0.07981575168 | 1.10E-08 | 1.64E-07 |
| ZNF469 | 379.2313518 | 0.4437761264 | 0.1702565822 | 0.003289098376 | 0.01404801656 |
| RBM15 | 1736.862267 | 0.443761928 | 0.08759277757 | 1.66E-07 | 2.01E-06 |
| GPN2 | 882.7121678 | 0.443677315 | 0.08364515085 | 4.60E-08 | 6.17E-07 |
| C8orf33 | 1859.270483 | 0.4419528863 | 0.09757875719 | 2.47E-06 | 2.33E-05 |
| GGNBP2 | 2489.042271 | 0.4418814879 | 0.06835339077 | 4.21E-11 | 8.80E-10 |
| PLEKHM1 | 618.3429165 | 0.4414039661 | 0.1229533258 | 0.000128792075 | 0.000841046147 |
| ACSL1 | 2505.947188 | 0.4412070287 | 0.09175036125 | 6.21E-07 | 6.66E-06 |
| C9orf85 | 225.5920587 | 0.4410702712 | 0.1228197201 | 0.000128768547 | 0.000841046147 |
| FBXO28 | 1429.128815 | 0.4410356597 | 0.1223806823 | 0.000123797025 | 0.000813421851 |

| | baseMean | log2FoldChange | lfcSE | pvalue | padj |
|---|---|---|---|---|---|
| SERPINE2 | 1630.951855 | 0.4408876708 | 0.1100787021 | 2.48E-05 | 0.000191865613 |
| ANKRD49 | 322.1585408 | 0.4408817272 | 0.1241342301 | 0.000150837530 | 0.000963457740 |
| NUMBL | 659.7458679 | 0.4408521334 | 0.1520908073 | 0.001412394053 | 0.006816575382 |
| SLC6A15 | 1104.422471 | 0.4407310337 | 0.158704989 | 0.002046138944 | 0.009359435558 |
| CAB39 | 2941.325833 | 0.4404149749 | 0.125350272 | 0.000177952341 | 0.001115592598 |
| WDR44 | 1009.382679 | 0.4402336343 | 0.1107307946 | 2.81E-05 | 0.000215348663 |
| PRR24 | 103.0832581 | 0.4402228957 | 0.2141663885 | 0.01350484913 | 0.04520829753 |
| BCL2L2 | 1435.60279 | 0.4401501995 | 0.09001082984 | 4.14E-07 | 4.59E-06 |
| ARL16 | 977.0454582 | 0.4388576781 | 0.09491237925 | 1.53E-06 | 1.51E-05 |
| CORO7 | 718.012143 | 0.4386563649 | 0.1376809803 | 0.000562369379 | 0.003043663284 |
| HIST2H2AA3 | 12906.093 | 0.4381449551 | 0.1877020266 | 0.00678373501 | 0.02553806082 |
| KLF7 | 1513.950772 | 0.4380530063 | 0.1257771589 | 0.000196209570 | 0.001212687544 |
| C3orf33 | 85.01887984 | 0.4377327831 | 0.20552813 | 0.01146155004 | 0.03957569904 |
| C8orf88 | 422.6961407 | 0.4375201314 | 0.1120600783 | 3.81E-05 | 0.000281503527 |
| SMIM15 | 1813.53943 | 0.4367326416 | 0.1071385783 | 1.82E-05 | 0.000145755074 |
| ZNF595 | 341.7877158 | 0.4367134885 | 0.1343681749 | 0.000453158138 | 0.002524156947 |
| MASTL | 1515.164871 | 0.4362939642 | 0.1664754895 | 0.003271903342 | 0.01399062862 |
| ASB1 | 1986.450519 | 0.4357572092 | 0.1296702313 | 0.000302086330 | 0.001769379771 |
| SQSTM1 | 25743.54491 | 0.4355807315 | 0.1366245812 | 0.000562805691 | 0.003044917857 |
| ZNF567 | 301.8142664 | 0.4354408615 | 0.1274204479 | 0.000250874215 | 0.001500557483 |
| IST1 | 4934.300389 | 0.4353301099 | 0.08282424129 | 6.16E-08 | 8.06E-07 |
| TMF1 | 3218.32075 | 0.4352344131 | 0.09043635773 | 6.25E-07 | 6.70E-06 |
| MBD4 | 1383.051913 | 0.4351408432 | 0.08110611258 | 3.40E-08 | 4.64E-07 |
| HACE1 | 1233.725868 | 0.4348412833 | 0.1848042767 | 0.006751528307 | 0.02542838967 |
| DAXX | 2432.927086 | 0.4340011697 | 0.0826041166 | 6.27E-08 | 8.20E-07 |
| EYA4 | 181.879081 | 0.4339898584 | 0.1618626527 | 0.002783003371 | 0.01215849096 |
| TGS1 | 1500.609258 | 0.4337738192 | 0.1087173591 | 2.74E-05 | 0.000209815999 |
| CLN8 | 843.9246948 | 0.4336928466 | 0.09934088147 | 5.26E-06 | 4.65E-05 |
| CENPJ | 1285.741454 | 0.4334196164 | 0.1767950855 | 0.00529441422 | 0.02082106004 |
| DCLK2 | 404.0275369 | 0.4333958747 | 0.1693696063 | 0.003936060032 | 0.01631514416 |
| KDM6A | 1113.719977 | 0.433125887 | 0.08551672346 | 1.72E-07 | 2.07E-06 |
| MAGOH | 740.2394961 | 0.4329551773 | 0.1066163255 | 2.01E-05 | 0.000159134290 |
| UBE2J2 | 1587.975835 | 0.4328267052 | 0.08542214327 | 1.71E-07 | 2.06E-06 |
| ARHGAP17 | 1163.380705 | 0.4322720115 | 0.07596729726 | 5.44E-09 | 8.49E-08 |
| SLAIN2 | 1540.502092 | 0.4312800294 | 0.1076367778 | 2.54E-05 | 0.000196099307 |
| TSSC4 | 1870.918575 | 0.4308348255 | 0.08106131715 | 4.55E-08 | 6.11E-07 |
| MYO9B | 3059.933718 | 0.4306381752 | 0.09488236247 | 2.37E-06 | 2.24E-05 |
| CDKN2B | 225.6312035 | 0.4300796169 | 0.2004523122 | 0.01143355352 | 0.03949748916 |
| GMEB2 | 765.5589825 | 0.4298862891 | 0.1286636748 | 0.000337213496 | 0.001942271471 |
| MAP3K5 | 2078.123621 | 0.4298202163 | 0.1369564296 | 0.000678167767 | 0.003579312259 |

| | baseMean | log2FoldChange | lfcSE | pvalue | padj |
|---|---|---|---|---|---|
| GOLIM4 | 2624.99549 | 0.429529772 | 0.1179645408 | 0.000111287773 | 0.000741702625 |
| ITPRIPL2 | 2626.906732 | 0.4294175702 | 0.1567578743 | 0.002388451657 | 0.01065200785 |
| CSNK1G3 | 1141.349861 | 0.4293704086 | 0.09342090274 | 1.82E-06 | 1.77E-05 |
| CNOT2 | 2557.243492 | 0.4291597827 | 0.09182991976 | 1.26E-06 | 1.27E-05 |
| PID1 | 492.0250131 | 0.4289671417 | 0.1119414218 | 5.22E-05 | 0.000374864578 |
| MLLT11 | 847.9926719 | 0.4287299908 | 0.1339506762 | 0.000551559752 | 0.002993865534 |
| COPRS | 1910.070997 | 0.4287136462 | 0.1334065595 | 0.000528474115 | 0.00288538728 |
| TDP2 | 1158.967253 | 0.4282687122 | 0.08889123306 | 6.18E-07 | 6.64E-06 |
| ZNF574 | 684.2637888 | 0.4282422038 | 0.1260066858 | 0.000276273057 | 0.001633379395 |
| PIK3C3 | 2183.384708 | 0.4273878311 | 0.08579597834 | 2.77E-07 | 3.19E-06 |
| ANKLE2 | 3551.530929 | 0.4273237815 | 0.1008890223 | 9.39E-06 | 7.95E-05 |
| HSPA14 | 1942.438257 | 0.4267991409 | 0.09444300757 | 2.64E-06 | 2.49E-05 |
| KLHL18 | 1159.23066 | 0.4265667471 | 0.1232302619 | 0.000220699982 | 0.001342321094 |
| PSMD12 | 2218.688405 | 0.4260484742 | 0.1693215033 | 0.004578700973 | 0.01851501325 |
| RNMT | 1601.924871 | 0.4259498795 | 0.1013300415 | 1.12E-05 | 9.33E-05 |
| CHSY3 | 445.6299337 | 0.4258627889 | 0.1485383812 | 0.001642468859 | 0.00776586816 |
| SETD1A | 172.720932 | 0.4256534552 | 0.1800088551 | 0.006810685843 | 0.02560362571 |
| LOC101928527 | 339.7710794 | 0.4254509128 | 0.1214627093 | 0.000189696682 | 0.001175850921 |
| TUBB6 | 20660.32549 | 0.4249326191 | 0.1014344819 | 1.24E-05 | 1.02E-04 |
| ZBTB24 | 633.1785489 | 0.4248921225 | 0.1205233911 | 1.75E-04 | 0.001102124659 |
| ELOVL4 | 220.052892 | 0.4241592638 | 0.1268374998 | 0.000341449334 | 0.001963630414 |
| NUS1 | 1707.081608 | 0.4238926967 | 0.1346840191 | 0.000672136087 | 0.003552514807 |
| CCDC94 | 673.5181568 | 0.4235981464 | 0.1427174256 | 0.001206049789 | 0.005957821936 |
| MRPS12 | 1818.665425 | 0.4232675701 | 0.1125025777 | 7.08E-05 | 0.000493479219 |
| TMEM204 | 516.4494083 | 0.4231105807 | 0.1606060798 | 0.003311297947 | 0.01412662325 |
| DNAJB5 | 876.1169373 | 0.4229724317 | 0.09402458915 | 2.95E-06 | 2.76E-05 |
| CMTM3 | 1441.638073 | 0.4226619438 | 0.09394058098 | 2.97E-06 | 2.77E-05 |
| RNF6 | 2341.272171 | 0.4223120334 | 0.08453327904 | 2.56E-07 | 2.96E-06 |
| CNBP | 12181.67808 | 0.4222056232 | 0.07831225653 | 2.74E-08 | 3.81E-07 |
| RN7SL2 | 4304220.894 | 0.4220535556 | 0.1573511751 | 0.002915321291 | 0.01265118587 |
| TRIM8 | 1256.739947 | 0.4217308324 | 0.08970714407 | 1.12E-06 | 1.13E-05 |
| CHKA | 752.4257369 | 0.4217152606 | 0.08863606971 | 8.54E-07 | 8.88E-06 |
| EFR3B | 77.05891186 | 0.4216910271 | 0.1763584198 | 0.006446075639 | 0.02448982398 |
| RAB12 | 947.5332408 | 0.4216852799 | 0.08294695273 | 1.62E-07 | 1.96E-06 |
| SF1 | 1840.748896 | 0.4212517851 | 0.106974624 | 3.51E-05 | 0.000262079152 |
| THEMIS2 | 368.9454319 | 0.4212314318 | 0.1801268408 | 0.007364721084 | 0.02731772103 |
| MTMR6 | 2477.94125 | 0.4210108667 | 0.1350109289 | 0.000748022609 | 0.003906456906 |
| PPP2R1B | 2306.863878 | 0.420192996 | 0.1581315429 | 0.003153021368 | 0.01351722866 |
| BID | 1749.790447 | 0.4196195134 | 0.08292791171 | 1.84E-07 | 2.20E-06 |
| IFNGR1 | 1630.000696 | 0.418148716 | 0.1585078107 | 0.003346516126 | 0.01424421915 |

| | baseMean | log2FoldChange | lfcSE | pvalue | padj |
|---|---|---|---|---|---|
| WDR35 | 1177.824721 | 0.4181104036 | 0.1545991398 | 0.002752002626 | 0.01203718187 |
| NAA50 | 6332.179656 | 0.4175368095 | 0.1313298114 | 0.000608832659 | 0.003264281407 |
| MYO10 | 7957.914277 | 0.4171996385 | 0.09375186725 | 3.19E-06 | 2.94E-05 |
| TMEM70 | 688.6015114 | 0.4171013084 | 0.1074581408 | 4.49E-05 | 0.000326237661 |
| TMED5 | 2399.921519 | 0.4170083847 | 0.09709216463 | 7.68E-06 | 6.62E-05 |
| ZNF330 | 1617.736898 | 0.4165050242 | 0.09052502409 | 1.86E-06 | 1.80E-05 |
| ZNF668 | 436.2971422 | 0.4164888231 | 0.1304782083 | 0.000591053612 | 0.003182392067 |
| EMC6 | 892.7582992 | 0.4163536488 | 0.1033919546 | 2.46E-05 | 0.000190785636 |
| RRP1B | 2622.084418 | 0.4162334044 | 0.1271541755 | 0.000439112819 | 0.002457876226 |
| ARHGAP5 | 2986.19712 | 0.4158293271 | 0.1516769849 | 0.002494049479 | 0.01106822733 |
| RNF44 | 140.8349563 | 0.4157434602 | 0.156687596 | 0.003229962362 | 0.01382319908 |
| RNR2 | 66992.93974 | 0.4156890566 | 0.1647689103 | 0.00466018599 | 0.01877821629 |
| MAML1 | 721.8663101 | 0.4156829004 | 0.10713312 | 4.55E-05 | 0.000330090661 |
| ATAD3B | 1910.970018 | 0.4155606915 | 0.1833549952 | 0.00905883817 | 0.03250049193 |
| ALG11 | 428.3615999 | 0.4153826305 | 0.1621012804 | 0.004161234854 | 0.01705854233 |
| C2orf69 | 514.5337219 | 0.4149268654 | 0.1695557373 | 0.005715652239 | 0.0222021543 |
| SLC6A8 | 613.8731195 | 0.4147752248 | 0.140721645 | 0.00134910003 | 0.006560009909 |
| PCGF6 | 502.8865041 | 0.414673534 | 0.09861538027 | 1.15E-05 | 9.56E-05 |
| ABL1 | 4414.313363 | 0.4143038132 | 0.1046728888 | 3.30E-05 | 0.000247320852 |
| IP6K2 | 2645.258687 | 0.4142681017 | 0.08974339114 | 1.74E-06 | 1.69E-05 |
| MPP6 | 871.650267 | 0.4141396237 | 0.100472352 | 1.65E-05 | 0.000133531699 |
| TIMM23 | 3632.015431 | 0.4139673136 | 0.1191668273 | 0.000211543234 | 0.00129296684 |
| PDLIM5 | 3661.966986 | 0.4138760118 | 0.1279712583 | 0.000518686590 | 0.002839674229 |
| PGRMC2 | 1416.353003 | 0.4133581281 | 0.1273752989 | 0.000498471617 | 0.002742317176 |
| ATP6V0B | 2918.765199 | 0.4132264378 | 0.1126853464 | 0.000100745059 | 0.000678729952 |
| SYAP1 | 4084.514042 | 0.4129598737 | 0.0791612565 | 8.42E-08 | 1.07E-06 |
| SNRNP27 | 1009.193535 | 0.4126916151 | 0.1126496156 | 0.000108223828 | 0.000724451164 |
| MTPAP | 1150.243639 | 0.4126468591 | 0.10546462 | 4.01E-05 | 0.000295163585 |
| MAP3K11 | 1348.598215 | 0.4120717067 | 0.1287153318 | 0.000582602823 | 0.003140613119 |
| ZNF805 | 416.9634094 | 0.4115462493 | 0.1497005942 | 0.002475083368 | 0.01100045262 |
| TSPYL2 | 569.9125676 | 0.4113362459 | 0.1266083232 | 0.000496006765 | 0.002731155596 |
| SLC41A2 | 182.0504136 | 0.4103565686 | 0.1323726106 | 0.000828027395 | 0.004277758464 |
| USP42 | 253.6734543 | 0.4102003496 | 0.1533835861 | 0.003103597062 | 0.01332836938 |
| EPB41L4A-AS1 | 622.7677057 | 0.409910753 | 0.192557683 | 0.01285786072 | 0.04340074547 |
| LOC100652736 | 98.63770628 | 0.4096352309 | 0.1813043329 | 0.009438810414 | 0.0336126401 |
| EMC3 | 2710.116308 | 0.4093822901 | 0.07773717337 | 6.34E-08 | 8.27E-07 |
| ZFP91 | 3503.635106 | 0.4093241467 | 0.1071129671 | 5.88E-05 | 0.000417351805 |
| THUMPD1 | 1866.258561 | 0.4093094886 | 0.09546898794 | 8.63E-06 | 7.37E-05 |
| CERK | 5143.701798 | 0.409131826 | 0.09716768234 | 1.13E-05 | 9.43E-05 |
| HIST1H2AI | 10665.97496 | 0.4090852386 | 0.1917021539 | 0.01236763812 | 0.04211842727 |

| | baseMean | log2FoldChange | lfcSE | pvalue | padj |
|---|---|---|---|---|---|
| RAB5A | 1753.756877 | 0.4088655958 | 0.08622707564 | 9.67E-07 | 9.89E-06 |
| SCARNA5 | 2169.240956 | 0.4086413465 | 0.1435235255 | 0.00185706342 | 0.008624397148 |
| SYDE1 | 2376.498669 | 0.4085019376 | 0.1029208053 | 3.17E-05 | 0.000238200706 |
| ZMYND8 | 1731.666105 | 0.4084051094 | 0.1327217177 | 0.000911201108 | 0.004643009345 |
| ZFAND6 | 1751.753257 | 0.4078574467 | 0.08913904553 | 2.17E-06 | 2.07E-05 |
| EIF1B | 958.7048706 | 0.407844377 | 0.08167658237 | 2.70E-07 | 3.11E-06 |
| PTBP1 | 6924.655784 | 0.4077996114 | 0.1184667529 | 0.000306601737 | 0.001790895748 |
| PPP4R2 | 2014.757777 | 0.4076795945 | 0.09097494361 | 3.36E-06 | 3.09E-05 |
| SLC52A2 | 1651.952787 | 0.4071997687 | 0.1066803609 | 6.02E-05 | 0.000426600806 |
| EXOSC6 | 617.4449818 | 0.4071379884 | 0.1083575108 | 7.63E-05 | 0.000527593734 |
| SNORA73A | 13462.37732 | 0.4070273304 | 0.1783967038 | 0.007239540223 | 0.02692360799 |
| FXR2 | 3010.503141 | 0.4068216771 | 0.1062936475 | 5.75E-05 | 0.000408803304 |
| CD276 | 3014.065365 | 0.4066113132 | 0.09247423552 | 4.95E-06 | 4.42E-05 |
| AKAP8L | 1743.387613 | 0.4063696457 | 0.1047592907 | 4.69E-05 | 0.000339514821 |
| POLR2C | 4308.248975 | 0.4063295785 | 0.08389784506 | 5.84E-07 | 6.31E-06 |
| RAPGEF1 | 1313.529501 | 0.4061887926 | 0.1021993597 | 3.15E-05 | 0.000237230689 |
| INPP1 | 702.1524739 | 0.4059753232 | 0.08688171923 | 1.36E-06 | 1.35E-05 |
| NPAT | 1052.639819 | 0.4057363746 | 0.1090960234 | 9.06E-05 | 0.000616680194 |
| CLK3 | 1553.619006 | 0.4053034821 | 0.1186078988 | 0.000279257091 | 0.001648635539 |
| SF3B4 | 1757.823845 | 0.4052146486 | 0.1306559224 | 0.000836000337 | 0.004305502947 |
| UXS1 | 1347.071177 | 0.4052116902 | 0.09225352739 | 5.11E-06 | 4.53E-05 |
| GPATCH2L | 2331.265531 | 0.4051965251 | 0.09607865872 | 1.15E-05 | 9.56E-05 |
| CLIP2 | 2383.359115 | 0.4048971655 | 0.1394598246 | 0.001586208826 | 0.00753816253 |
| MTHFD1L | 4407.587383 | 0.4048051235 | 0.121751174 | 0.000399181995 | 0.002253002552 |
| ING1 | 335.9669012 | 0.4045558302 | 0.1016527249 | 3.11E-05 | 0.000234380497 |
| MOB4 | 1806.146418 | 0.4045428625 | 0.1041658495 | 4.61E-05 | 0.000334347181 |
| ZNF551 | 383.6946374 | 0.4044873981 | 0.111931019 | 0.000135250494 | 0.000877448635 |
| SCARNA10 | 4393.266996 | 0.4038387854 | 0.1469188189 | 0.002553189859 | 0.01128359864 |
| GLS | 3373.074856 | 0.4032857393 | 0.1400078242 | 0.001714186993 | 0.008044135148 |
| NUP54 | 2356.440181 | 0.4030694633 | 0.1140834176 | 0.000184183921 | 0.001148853963 |
| DNAAF2 | 387.0328549 | 0.4030162511 | 0.1489768601 | 0.002908341957 | 0.01262837164 |
| TCEB3 | 3329.492428 | 0.4029151613 | 0.1124150136 | 0.000152934542 | 0.000974344205 |
| CBR3 | 348.2419201 | 0.4027695514 | 0.1278419346 | 0.000716512664 | 0.003764515669 |
| SFSWAP | 1151.777646 | 0.4023752264 | 0.08734163752 | 1.89E-06 | 1.83E-05 |
| C18orf21 | 522.7830373 | 0.4018559601 | 0.09317702337 | 7.42E-06 | 6.42E-05 |
| SNORD3A | 427060.6402 | 0.4017981323 | 0.1056918061 | 6.55E-05 | 0.000458800877 |
| ZNF765 | 547.0670691 | 0.4011345535 | 0.1318052887 | 0.001029782609 | 0.005179876104 |
| DENND6A | 479.8670215 | 0.4007191751 | 0.08993020823 | 3.89E-06 | 3.54E-05 |
| NUFIP1 | 528.4771755 | 0.4006207676 | 0.1074203255 | 8.75E-05 | 0.000597212098 |
| DUS3L | 910.696935 | 0.4004437888 | 0.1662353602 | 0.006679153016 | 0.02522727277 |

| | baseMean | log2FoldChange | lfcSE | pvalue | padj |
|---|---|---|---|---|---|
| EID2 | 463.7607538 | 0.4003241254 | 0.1117517429 | 1.55E-04 | 9.84E-04 |
| ARL13B | 511.7485715 | 0.4001658987 | 0.1227586294 | 0.000498586378 | 0.002742317176 |
| CUL3 | 2770.791798 | 0.3999633194 | 0.1125691433 | 0.000172295941 | 0.001084699482 |
| OSBPL10 | 1127.863161 | 0.3998509478 | 0.1510235867 | 0.003473135237 | 0.01469912182 |
| UGCG | 3305.770832 | 0.3998052602 | 0.189582994 | 0.01329553669 | 0.04460602008 |
| MAPKAPK5-AS1 | 542.6763081 | 0.3997557008 | 0.09356712214 | 8.94E-06 | 7.62E-05 |
| CPSF6 | 3571.068526 | 0.3996553117 | 0.07987356325 | 2.66E-07 | 3.06E-06 |
| MIER3 | 959.7905046 | 0.3993019683 | 0.08404704577 | 9.52E-07 | 9.76E-06 |
| RAI14 | 11151.92761 | 0.3992342982 | 0.1367614209 | 0.001542835992 | 0.007367314012 |
| GPATCH2 | 841.8815398 | 0.3992061389 | 0.08403370008 | 9.52E-07 | 9.76E-06 |
| HSF2 | 843.6006749 | 0.3986516318 | 0.0915921067 | 6.24E-06 | 5.46E-05 |
| EML1 | 1007.097531 | 0.3984537412 | 0.1009129527 | 3.59E-05 | 0.000267603198 |
| MOB1B | 911.2885728 | 0.397764748 | 0.122409104 | 0.000520948009 | 0.002848474077 |
| PFDN2 | 2569.909168 | 0.3977166868 | 0.1643918201 | 0.006221279173 | 0.02375092964 |
| RFK | 668.8350167 | 0.3970706786 | 0.1643862965 | 0.0066542242 | 0.02515864503 |
| SIPA1L1 | 2090.664009 | 0.3969573647 | 0.09901511822 | 2.83E-05 | 2.17E-04 |
| ELK3 | 2154.862172 | 0.3966682282 | 0.1271830413 | 0.000816581213 | 0.004223021079 |
| SLC25A28 | 818.4301137 | 0.3965676779 | 0.09125271809 | 6.51E-06 | 5.67E-05 |
| EXOSC9 | 1744.528322 | 0.3964635396 | 0.1819881002 | 0.01208275716 | 0.04138490255 |
| VGLL4 | 1634.04707 | 0.3964506756 | 0.1530395998 | 0.004140957558 | 0.01699238024 |
| SIN3A | 1631.535074 | 0.3962074031 | 0.09908448661 | 3.00E-05 | 2.28E-04 |
| DNAJA2 | 4135.58706 | 0.3958484026 | 0.06836742133 | 3.39E-09 | 5.44E-08 |
| FBRS | 310.7003591 | 0.3957045743 | 0.1541907736 | 0.004446558503 | 0.01804929377 |
| ANKRD28 | 8449.195352 | 0.3955421066 | 0.1241358699 | 0.000643699987 | 0.003422874684 |
| UTP23 | 750.4817231 | 0.3954252566 | 0.1016918044 | 4.69E-05 | 0.000339514821 |
| RNF217 | 674.4419821 | 0.395209448 | 0.152485341 | 0.004151819532 | 0.01702932259 |
| EPHB2 | 2920.30548 | 0.3951950487 | 0.1042658259 | 6.98E-05 | 0.000486836205 |
| CRTC3 | 430.2651854 | 0.3949810275 | 0.1379500619 | 0.001866413665 | 0.008665117887 |
| ITPKC | 1338.139735 | 0.394587298 | 0.1086036868 | 0.000128655864 | 0.000840894275 |
| TWISTNB | 1593.524844 | 0.3943498586 | 0.1338099404 | 0.00143935415 | 0.00692648479 |
| KCTD15 | 1036.549907 | 0.3942968535 | 0.1651006458 | 0.00721852399 | 0.02686243531 |
| FOXK1 | 1277.928788 | 0.3942484813 | 0.1201239483 | 0.000454676104 | 0.002528827986 |
| MRPL36 | 1974.740872 | 0.3940090865 | 0.1239744181 | 0.000674285240 | 0.003561345495 |
| ZNF655 | 2537.489493 | 0.3937915688 | 0.1020118215 | 5.29E-05 | 0.000378725584 |
| TP53BP2 | 2103.832684 | 0.3936343133 | 0.1120369345 | 2.04E-04 | 0.001252309455 |
| FAM132B | 247.229744 | 0.3933089277 | 0.1420882151 | 0.002498265173 | 0.01108363235 |
| DSTYK | 1550.234461 | 0.3924988512 | 0.08325474358 | 1.16E-06 | 1.17E-05 |
| TATDN2 | 1822.762936 | 0.3920429431 | 0.08292020837 | 1.09E-06 | 1.10E-05 |
| TRIM52-AS1 | 340.9890471 | 0.3920057321 | 0.1244530696 | 0.000745058640 | 0.003896444715 |
| METAP2 | 4651.961542 | 0.3910051931 | 0.09595068225 | 1.92E-05 | 0.000152763497 |

| | baseMean | log2FoldChange | lfcSE | pvalue | padj |
|---|---|---|---|---|---|
| ATF1 | 1100.546483 | 0.3909891135 | 0.07922886797 | 3.88E-07 | 4.34E-06 |
| ZNF565 | 123.8113276 | 0.390275849 | 0.1502332244 | 0.004160259238 | 0.01705854233 |
| TAGLN | 22266.89196 | 0.3902618646 | 0.1790948241 | 0.01231053762 | 0.04195345885 |
| ENAH | 5477.556396 | 0.3898548963 | 0.1134539993 | 0.000275673716 | 0.001631361671 |
| WASL | 1097.361528 | 0.3892724676 | 0.09397264902 | 1.65E-05 | 0.000133089429 |
| TNFAIP2 | 1280.286569 | 0.3892089893 | 0.1353861672 | 0.001831368571 | 0.008521014581 |
| SYNCRIP | 9618.580578 | 0.389053609 | 0.1551318976 | 0.005361106615 | 0.02103333765 |
| PTS | 1137.38258 | 0.3890525042 | 0.1001383008 | 4.87E-05 | 0.000350447899 |
| ZNF530 | 154.9823464 | 0.3889562513 | 0.1751612689 | 0.01127754807 | 0.0390855245 |
| CBLL1 | 564.0123921 | 0.3889024376 | 0.1094539859 | 0.000179357529 | 0.001122039606 |
| DEGS1 | 4103.239749 | 0.3884321173 | 0.07670156947 | 2.02E-07 | 2.40E-06 |
| GFOD1 | 103.2002298 | 0.3883201533 | 0.172044227 | 0.01033166109 | 0.03628877141 |
| MYBL2 | 2084.367027 | 0.388236865 | 0.1755110862 | 0.01151635479 | 0.03972822209 |
| C12orf45 | 591.0640399 | 0.3878168978 | 0.1354493789 | 0.00191010845 | 0.008835701409 |
| PET117 | 344.7886146 | 0.3876844308 | 0.1658840829 | 0.008472663438 | 0.03066346279 |
| NUAK1 | 724.2035091 | 0.3874143721 | 0.1377189288 | 0.002247234709 | 0.0101115375 |
| SAR1A | 7502.859605 | 0.3871319212 | 0.08059081531 | 7.53E-07 | 7.92E-06 |
| TMEM11 | 1083.750299 | 0.3871098165 | 0.1037972438 | 9.13E-05 | 0.000620308282 |
| MRPS2 | 2079.967553 | 0.3869981852 | 0.1004685571 | 5.60E-05 | 0.000398643800 |
| BAHD1 | 497.5249717 | 0.3869760796 | 0.1054124796 | 0.000114949249 | 0.000763712349 |
| SNRPB | 12106.33876 | 0.3869434076 | 0.1345061737 | 0.0018225894 | 0.008482817624 |
| AAED1 | 514.111904 | 0.3868835185 | 0.09461540593 | 2.09E-05 | 0.000164881603 |
| CDK7 | 1811.710453 | 0.3865639028 | 0.09353371483 | 1.73E-05 | 0.000139037941 |
| SCARNA12 | 5386.642244 | 0.386508913 | 0.154051297 | 0.004708254104 | 0.01893091962 |
| CIRH1A | 1246.161659 | 0.3860021025 | 0.1538801452 | 0.005407460199 | 0.0211594415 |
| LRRFIP1 | 4425.074179 | 0.3854885793 | 0.1118807724 | 0.000270218852 | 0.001602265427 |
| GLIPR1 | 2894.158508 | 0.3853720927 | 0.1564029205 | 0.006075008464 | 0.02333010086 |
| BAZ1A | 3566.353874 | 0.3846860189 | 0.155738571 | 0.006025082909 | 0.02316226683 |
| WBP11 | 2924.549171 | 0.3846236001 | 0.1220451614 | 0.000769752757 | 0.004010429287 |
| GTPBP1 | 1471.04745 | 0.3844970231 | 0.123200774 | 0.000846905218 | 0.004357142987 |
| ERICH1 | 397.3507745 | 0.3843510314 | 0.1123545904 | 0.000296161752 | 0.001738097097 |
| LOC100996717 | 485.11873 | 0.38407443 | 0.126198236 | 0.001088692095 | 0.005433971373 |
| SECISBP2L | 1877.682371 | 0.384023039 | 0.08896722557 | 7.84E-06 | 6.75E-05 |
| LMCD1 | 808.0204715 | 0.3839943172 | 0.1447784514 | 0.003647553889 | 0.0153110882 |
| C15orf39 | 278.2220183 | 0.3834879443 | 0.1308006471 | 0.001572847216 | 0.007489006138 |
| NUDT4P1 | 172.8007754 | 0.3832826274 | 0.1455207156 | 0.003835142486 | 0.01596349915 |
| TFAM | 1597.68317 | 0.383070783 | 0.1453295332 | 0.003840587355 | 0.0159727668 |
| RFWD3 | 3184.808768 | 0.3825915716 | 0.1690226659 | 0.01038673503 | 0.03644781945 |
| FUS | 3025.858962 | 0.3818467899 | 0.1110579302 | 0.000281080353 | 0.001658084542 |
| DCAF16 | 1208.104747 | 0.3813517011 | 0.1038445038 | 0.000116765284 | 0.000774237445 |

| | baseMean | log2FoldChange | lfcSE | pvalue | padj |
|---|---|---|---|---|---|
| SFPQ | 10153.30993 | 0.3810319872 | 0.1125175851 | 0.000358482472 | 0.002047351568 |
| WWP1 | 719.6686986 | 0.3803085224 | 0.1196372471 | 0.000705893702 | 0.003716568533 |
| PTPN9 | 2418.059639 | 0.380184665 | 0.0654346306 | 3.16E-09 | 5.10E-08 |
| FAS | 1155.659773 | 0.3801769962 | 0.1077913709 | 0.000203837509 | 0.001251520278 |
| FUNDC2 | 2195.923213 | 0.3801127939 | 0.07669134034 | 3.59E-07 | 4.07E-06 |
| STAT3 | 5535.912162 | 0.3800771511 | 0.1727196828 | 0.01208950781 | 0.04139850087 |
| TBC1D23 | 2667.310831 | 0.3799746581 | 0.1149060182 | 0.000453462178 | 0.002524905899 |
| GMPS | 3881.904342 | 0.3796528341 | 0.1035551057 | 0.000120076427 | 0.000792472487 |
| SNRNP35 | 501.8353505 | 0.37959314 | 0.1011564269 | 8.51E-05 | 0.000581783920 |
| CTH | 262.6504317 | 0.3791172045 | 0.1569433145 | 0.007110609898 | 0.02651386696 |
| KRAS | 1096.391732 | 0.3787435223 | 0.1031235895 | 1.18E-04 | 0.000778089602 |
| TXN | 13330.68647 | 0.3787354391 | 0.07127054061 | 5.43E-08 | 7.18E-07 |
| ATG16L1 | 1580.675456 | 0.3786690467 | 0.07851712838 | 7.17E-07 | 7.58E-06 |
| AFAP1L2 | 583.2955139 | 0.3785090294 | 0.1734668178 | 0.01285561005 | 0.04340074547 |
| PRDM4 | 1651.538623 | 0.3784636549 | 0.1007459621 | 8.41E-05 | 0.000576118911 |
| NUMB | 2095.801043 | 0.3779576223 | 0.07663324589 | 4.11E-07 | 4.56E-06 |
| EXT1 | 5341.99148 | 0.3777689632 | 0.1040829687 | 0.000130236580 | 0.000849733764 |
| UBE2K | 1420.011959 | 0.3777637286 | 0.1039476985 | 0.000127032379 | 0.000832109591 |
| QKI | 3437.587752 | 0.3769101609 | 0.1043082303 | 0.000153939432 | 0.000979907745 |
| SNRPG | 2362.463284 | 0.3767222641 | 0.1472353486 | 0.004878431381 | 0.0194888556 |
| PTBP3 | 1728.164684 | 0.376047143 | 0.112699521 | 0.000414215177 | 0.002331663432 |
| UTP11L | 2093.003411 | 0.3759573486 | 0.1024032132 | 0.000118258827 | 0.000782210430 |
| EMC8 | 1721.29425 | 0.375871208 | 0.100228854 | 9.03E-05 | 0.000615278695 |
| COMMD5 | 1033.522579 | 0.375805418 | 0.08360109199 | 3.51E-06 | 3.21E-05 |
| CCNC | 1886.075622 | 0.3754073058 | 0.1086129396 | 0.000265016503 | 0.001575181927 |
| DUSP12 | 997.1265761 | 0.3753044783 | 0.1155487045 | 0.000564747308 | 0.003052095344 |
| UTP18 | 2350.707591 | 0.3747812598 | 0.1478751722 | 0.005333596635 | 0.02094748623 |
| SNRPA1 | 1241.215565 | 0.374369922 | 0.1726987457 | 0.0135262093 | 0.04523623769 |
| KRR1 | 1924.659056 | 0.3742943203 | 0.1317339738 | 0.001928472871 | 0.008906908832 |
| SIK2 | 1155.837739 | 0.3734390094 | 0.08541850507 | 6.25E-06 | 5.46E-05 |
| CTNNB1 | 23929.4513 | 0.3731736658 | 0.04769809874 | 2.70E-15 | 9.06E-14 |
| MAP2K7 | 385.1016049 | 0.3724610375 | 0.123574838 | 0.001252775567 | 0.006153934482 |
| MEAF6 | 1462.756905 | 0.372245845 | 0.07152651965 | 1.01E-07 | 1.28E-06 |
| RNMTL1 | 740.4394454 | 0.3721084116 | 0.1383435441 | 0.003466911648 | 0.01468112842 |
| TMEM39A | 1974.653897 | 0.3721027315 | 0.09278054098 | 3.07E-05 | 0.000232350252 |
| DBN1 | 10298.55833 | 0.3720795051 | 0.08788816138 | 1.18E-05 | 9.73E-05 |
| DHRS7 | 1899.414445 | 0.3717532874 | 0.1015940041 | 0.000125760284 | 0.000824502364 |
| GNL1 | 3284.908789 | 0.3716279126 | 0.1044121617 | 0.000148291286 | 0.000949638261 |
| KIF9 | 141.0606787 | 0.3714623045 | 0.1612621217 | 0.009777651017 | 0.03462879305 |
| RNR1 | 13515.64641 | 0.3708866689 | 0.1505339976 | 0.006582791291 | 0.02493288719 |

| | baseMean | log2FoldChange | lfcSE | pvalue | padj |
|---|---|---|---|---|---|
| AP5Z1 | 1124.057008 | 0.3707788685 | 0.1421947711 | 0.004380444093 | 0.01779956933 |
| FAM103A1 | 1182.768902 | 0.3706761467 | 0.0731183145 | 2.07E-07 | 2.44E-06 |
| C11orf95 | 1153.051679 | 0.3703278046 | 0.09115930102 | 2.47E-05 | 1.91E-04 |
| ZNF586 | 496.57615 | 0.3702310935 | 0.090275893 | 2.10E-05 | 0.000165493206 |
| TBC1D25 | 291.5409228 | 0.3699933879 | 0.1234148725 | 0.001329874754 | 0.006479222909 |
| CYR61 | 4128.917955 | 0.3699702146 | 0.142991341 | 0.004583900364 | 0.01852597517 |
| RC3H1 | 1357.068084 | 0.369776972 | 0.08884456433 | 1.61E-05 | 1.30E-04 |
| SIVA1 | 2120.60228 | 0.3696567854 | 0.1441832452 | 0.004924917662 | 0.01964820447 |
| DEXI | 1073.335967 | 0.3695963063 | 0.08332746602 | 4.72E-06 | 4.23E-05 |
| E2F3 | 751.3913467 | 0.3692692597 | 0.1263253914 | 0.001699011078 | 0.007987551605 |
| HIPK1 | 2034.703009 | 0.3689986346 | 0.08011909307 | 2.14E-06 | 2.04E-05 |
| SRC | 563.1380899 | 0.3688675509 | 0.1219605739 | 0.001227366382 | 0.006047074141 |
| ORAOV1 | 465.7464943 | 0.3686688413 | 0.1185088735 | 0.000923979356 | 0.004700078115 |
| WIPI2 | 2603.590778 | 0.3684723733 | 0.08236071053 | 3.97E-06 | 3.60E-05 |
| COPS2 | 3341.732304 | 0.368374677 | 0.1089813454 | 0.000357118535 | 0.002041911629 |
| MTF2 | 1063.725329 | 0.3682727361 | 0.1275144876 | 0.001922609964 | 0.00888667456 |
| SELK | 1120.934949 | 0.3681344175 | 0.1276817394 | 0.001993354017 | 0.009143268008 |
| THAP4 | 2266.915303 | 0.3680446376 | 0.1005548187 | 1.28E-04 | 8.39E-04 |
| TIMM8A | 461.4765602 | 0.3680124166 | 0.1573956777 | 0.009087257217 | 0.03257105747 |
| GMEB1 | 525.1485953 | 0.3679873372 | 0.1301086325 | 0.002282298208 | 0.01024762908 |
| TMEM57 | 848.1110911 | 0.367946125 | 0.1074157788 | 0.000308468696 | 0.001799682767 |
| KCTD12 | 525.4893698 | 0.367935043 | 0.1408854083 | 0.004361474658 | 0.01774262191 |
| TAX1BP1 | 4764.82094 | 0.3679299547 | 0.1004966129 | 1.27E-04 | 0.000833673764 |
| FAM160B1 | 1293.026051 | 0.3678028565 | 0.1293162326 | 0.002160508249 | 0.009789350978 |
| AKIRIN2 | 419.149065 | 0.3674724861 | 0.1263162938 | 0.001779305138 | 0.008299521995 |
| PAK1IP1 | 1775.718449 | 0.3673988693 | 0.1651710079 | 0.0121056212 | 0.04143461932 |
| NRF1 | 674.7056002 | 0.3669945424 | 0.1381954299 | 0.003893822699 | 0.01616256653 |
| JUN | 3402.616294 | 0.3669282539 | 0.1542767738 | 0.008239012162 | 0.02997084097 |
| CRIP2 | 776.9997188 | 0.3665747411 | 0.1359821997 | 0.003620815086 | 0.01522030373 |
| GFPT1 | 4083.437326 | 0.3659910766 | 0.09539948047 | 6.30E-05 | 0.000443689268 |
| NUDT11 | 906.8169127 | 0.3656634381 | 0.08307542375 | 5.57E-06 | 4.91E-05 |
| C2CD2L | 240.3610669 | 0.3655022767 | 0.1181421029 | 0.000988056942 | 0.004995307237 |
| FHL3 | 596.7174177 | 0.3648047728 | 0.1585607114 | 0.01009816177 | 0.03556932353 |
| KIAA0930 | 2578.774081 | 0.3646329412 | 0.1124776135 | 0.000614163752 | 0.00329049446 |
| CENPC | 684.8428026 | 0.3645025867 | 0.1321570685 | 0.003011584837 | 0.01299695265 |
| RWDD2A | 295.3772292 | 0.3643141876 | 0.1357642334 | 0.003575386282 | 0.0150654489 |
| PJA2 | 5566.804 | 0.3642648222 | 0.09010853774 | 2.60E-05 | 0.000200372819 |
| PUS7 | 1264.895524 | 0.3640813161 | 0.1457221087 | 0.00595804299 | 0.02296383693 |
| ZNF317 | 951.6926628 | 0.3639959244 | 0.07509968771 | 6.62E-07 | 7.07E-06 |
| HNRNPAB | 7522.949167 | 0.3639857554 | 0.1589895286 | 0.01062374617 | 0.03717437291 |

| | baseMean | log2FoldChange | lfcSE | pvalue | padj |
|---|---|---|---|---|---|
| MSANTD3 | 2819.54804 | 0.3638587779 | 0.1438912004 | 0.005559722725 | 0.02167549402 |
| RASA3 | 3823.316797 | 0.3638521752 | 0.09198929173 | 3.97E-05 | 0.000291829353 |
| FMNL3 | 1659.33436 | 0.363455608 | 0.09140716726 | 3.66E-05 | 2.71E-04 |
| DDX20 | 1277.515215 | 0.3630895527 | 0.1468353372 | 0.006525754275 | 0.02476094684 |
| SMIM4 | 321.8050247 | 0.3629244742 | 0.1491775338 | 0.007208794824 | 0.02683293653 |
| TNKS1BP1 | 3647.612613 | 0.3628791331 | 0.1172792114 | 0.000995972107 | 0.005028493965 |
| PRKD3 | 2658.518192 | 0.3626896729 | 0.134041059 | 0.003365090521 | 0.01431918627 |
| KLHL17 | 275.6850734 | 0.3624270277 | 0.1416910223 | 0.005144353941 | 0.02038166201 |
| RUSC1 | 1152.821323 | 0.3622067533 | 0.07442218488 | 6.00E-07 | 6.47E-06 |
| ARFIP2 | 2178.670188 | 0.362042735 | 0.07853459035 | 2.13E-06 | 2.03E-05 |
| NANP | 567.0232586 | 0.3619581202 | 0.1567612262 | 0.01004611064 | 0.035411113 |
| SGTB | 804.2425383 | 0.3617668769 | 0.1180733929 | 0.0010905123 | 0.005441232452 |
| YTHDC1 | 1941.526123 | 0.3617396092 | 0.08831455023 | 2.22E-05 | 0.000174207082 |
| SAMD4B | 965.2493229 | 0.3613727179 | 0.08906521107 | 2.61E-05 | 2.01E-04 |
| VMP1 | 3271.778116 | 0.3611659408 | 0.09142582121 | 4.08E-05 | 0.000299915881 |
| CYTH2 | 1935.303253 | 0.3609596703 | 0.09488301103 | 7.41E-05 | 0.000514548771 |
| ITGA5 | 22682.62413 | 0.3609459566 | 0.1209993977 | 0.001426495592 | 0.006871269124 |
| WDR20 | 794.2673792 | 0.3606283596 | 0.08615455111 | 1.50E-05 | 0.000122659865 |
| ZNF207 | 8759.240503 | 0.3605695767 | 0.0774589556 | 1.80E-06 | 1.75E-05 |
| SMG8 | 1635.920737 | 0.3604151296 | 0.07260288731 | 3.69E-07 | 4.18E-06 |
| PVT1 | 214.1989703 | 0.3589748054 | 0.1536664597 | 0.009485408124 | 0.03374629428 |
| LEF1 | 378.345694 | 0.3589100189 | 0.1412582541 | 0.005454981602 | 0.02130060872 |
| MOB2 | 573.3156896 | 0.3588285356 | 0.1421596035 | 0.005701355244 | 0.02216383243 |
| COLEC10 | 3912.096976 | 0.3587704691 | 0.09819692727 | 0.000136517055 | 0.000883356125 |
| DNAL1 | 448.6659115 | 0.3583460617 | 0.1383943079 | 0.004781492804 | 0.01917361873 |
| NPLOC4 | 4264.536141 | 0.3581717173 | 0.0829530498 | 8.42E-06 | 7.21E-05 |
| JKAMP | 1904.836382 | 0.3580516022 | 0.1101074593 | 0.000580761495 | 0.003132955764 |
| MBNL1-AS1 | 406.0235499 | 0.3579086801 | 0.1578091905 | 0.01128495277 | 0.0391020856 |
| ELF2 | 1503.195392 | 0.3578219557 | 0.1333490679 | 0.003582603796 | 0.01508523414 |
| RAB9A | 790.4946479 | 0.3577925546 | 0.0877429037 | 2.39E-05 | 0.000185678772 |
| MRGBP | 1216.263437 | 0.3575775858 | 0.08953852905 | 3.45E-05 | 0.000257515030 |
| THEM4 | 605.24382 | 0.3572689481 | 0.08656137736 | 1.95E-05 | 0.000154987673 |
| ZNF350 | 325.9296504 | 0.3571167957 | 0.1312104369 | 0.003272916717 | 0.01399094372 |
| SLC25A15 | 1420.627447 | 0.3570705302 | 0.09768572898 | 0.000134418572 | 0.000872431615 |
| RAB23 | 1818.042669 | 0.3570327373 | 0.08261707191 | 8.30E-06 | 7.11E-05 |
| KRT18 | 4954.329174 | 0.3569484828 | 0.1186380245 | 0.001358089778 | 0.006599412109 |
| PPP1R18 | 2986.902615 | 0.3569280781 | 0.08547266969 | 1.58E-05 | 0.000128334181 |
| GYPC | 4552.161599 | 0.3567326284 | 0.08876306422 | 3.10E-05 | 0.000234380497 |
| MDFIC | 1414.958686 | 0.3565030677 | 0.07554557041 | 1.28E-06 | 1.28E-05 |
| SNHG9 | 182.090614 | 0.3564449619 | 0.1519761495 | 0.00927474063 | 0.03314729075 |

| | baseMean | log2FoldChange | lfcSE | pvalue | padj |
|---|---|---|---|---|---|
| NDUFV2 | 5628.638413 | 0.3563717208 | 0.0931769311 | 5.89E-05 | 0.000417910742 |
| MCC | 1065.375861 | 0.3562468659 | 0.1647819366 | 0.01472551008 | 0.04861377374 |
| SCML1 | 597.9285241 | 0.3556910055 | 0.106836368 | 0.000456348744 | 0.002536236078 |
| CLEC2B | 767.7298114 | 0.3554412397 | 0.143889043 | 0.006694273752 | 0.02527156235 |
| XBP1 | 7501.229915 | 0.3552387541 | 0.095895985 | 9.90E-05 | 0.000667820494 |
| RBM25 | 3625.792856 | 0.3551245713 | 0.104512798 | 0.000356397899 | 0.00203857408 |
| HIST1H3A | 3439.372673 | 0.3550243823 | 0.163009259 | 0.01432366751 | 0.04747664417 |
| RPF1 | 1922.431953 | 0.3549712123 | 0.1219272047 | 0.001848630763 | 0.008598645246 |
| JAK1 | 12156.28107 | 0.3549515593 | 0.1029046354 | 0.000295421473 | 0.001735805177 |
| DR1 | 2192.55594 | 0.3549093593 | 0.08613643567 | 2.06E-05 | 0.000162738114 |
| ZNF124 | 1178.833399 | 0.3546994146 | 0.08917975949 | 3.72E-05 | 0.000275277709 |
| ABRACL | 1307.515853 | 0.3544166228 | 0.1222343697 | 0.001941617432 | 0.00895562018 |
| MTMR10 | 476.6836705 | 0.3538618839 | 0.1356302251 | 0.00467111073 | 0.018806968 |
| ATP6V0A2 | 1048.153914 | 0.3537578923 | 0.1199336302 | 0.001649128904 | 0.007794882619 |
| LRFN4 | 740.2081399 | 0.3536561564 | 0.1210419087 | 0.001801259902 | 0.008388789078 |
| WAPAL | 3117.608201 | 0.3535292551 | 0.1092710643 | 0.000637487532 | 0.003395904069 |
| USP7 | 3898.998035 | 0.3535127237 | 0.1194874594 | 0.001611431709 | 0.007638524903 |
| WDR45B | 4016.140046 | 0.3533839488 | 0.06801929293 | 1.12E-07 | 1.40E-06 |
| TBK1 | 1183.330882 | 0.3533550942 | 0.1080765002 | 0.000567144766 | 0.003063939921 |
| RPS16 | 14468.13341 | 0.3531821055 | 0.08694530429 | 2.46E-05 | 0.000190785244 |
| CERS5 | 1233.693412 | 0.3530408521 | 0.08113263799 | 7.34E-06 | 6.35E-05 |
| RBMXL1 | 794.8487479 | 0.3528622756 | 0.1163347636 | 0.001298223292 | 0.006343697601 |
| U2AF1 | 1924.363055 | 0.3528326449 | 0.1144977321 | 0.001074862275 | 0.005373950441 |
| SLC25A44 | 809.9896212 | 0.3527209252 | 0.1145455911 | 0.001086134703 | 0.00542392095 |
| EMP2 | 756.1375739 | 0.3523980865 | 0.09456161204 | 1.04E-04 | 6.97E-04 |
| FOXF1 | 3904.792608 | 0.352396311 | 0.1571297921 | 0.01254569074 | 0.04258841529 |
| GMFB | 2240.847454 | 0.352028864 | 0.1223898334 | 0.002089027677 | 0.009520518237 |
| RIN3 | 144.5108563 | 0.351549069 | 0.1583758782 | 0.0130069394 | 0.04383438678 |
| UTP6 | 2832.828045 | 0.3513808608 | 0.1195915535 | 0.001851171965 | 0.008605088788 |
| POLR1D | 2839.618697 | 0.3511762149 | 0.07732045535 | 3.05E-06 | 2.83E-05 |
| MALSU1 | 1036.35669 | 0.3510583298 | 0.08036082314 | 6.81E-06 | 5.91E-05 |
| PPP2CA | 5792.899014 | 0.3509495136 | 0.1123858025 | 0.001034597348 | 0.005197071498 |
| DNAJB6 | 4184.660588 | 0.3509028162 | 0.08743343083 | 3.24E-05 | 0.000243264651 |
| SNORD17 | 3720.951775 | 0.3507439625 | 0.1541011201 | 0.01126370341 | 0.0390570875 |
| MCMBP | 4768.777727 | 0.3503993344 | 0.1132738455 | 0.001044964985 | 0.005242076704 |
| UFM1 | 2867.209132 | 0.3500884243 | 0.125018797 | 0.002708134895 | 0.01186622144 |
| DAZAP2 | 4916.458655 | 0.3500478454 | 0.06852163553 | 1.82E-07 | 2.18E-06 |
| SUPV3L1 | 1054.805258 | 0.3499501075 | 0.1039487857 | 0.000406885641 | 0.00229213784 |
| IER3IP1 | 1899.369094 | 0.3498741231 | 0.07544885197 | 1.95E-06 | 1.87E-05 |
| UBA2 | 6728.231886 | 0.3494371312 | 0.1073564266 | 0.000616024674 | 0.003298091112 |

| | baseMean | log2FoldChange | lfcSE | pvalue | padj |
|---|---|---|---|---|---|
| MRPL47 | 2275.088413 | 0.348883625 | 0.1279236097 | 0.003326876825 | 0.01418902007 |
| BTN2A1 | 1146.389155 | 0.348715548 | 0.08119916072 | 9.62E-06 | 8.12E-05 |
| ZBTB33 | 780.4238983 | 0.3481084651 | 0.1053112258 | 0.000510887886 | 0.002798605499 |
| PLAGL2 | 301.6304553 | 0.3479350378 | 0.140516331 | 0.006828441978 | 0.0256468232 |
| PPP4R1L | 176.5719269 | 0.3478247014 | 0.1549664171 | 0.01245161616 | 0.04231730497 |
| TSHZ3 | 890.2518346 | 0.3477743365 | 0.1546532195 | 0.01230916821 | 0.04195345885 |
| GRWD1 | 2717.981564 | 0.3477216601 | 0.1177358743 | 0.001661363756 | 0.007845241029 |
| RNF220 | 2276.62975 | 0.347498343 | 0.06827997892 | 2.00E-07 | 2.38E-06 |
| CDK20 | 231.1187484 | 0.3473484027 | 0.1587771165 | 0.01429037741 | 0.04738740073 |
| ADRM1 | 4475.417921 | 0.3473204505 | 0.1161895485 | 0.001426349885 | 0.006871269124 |
| ERBB2IP | 6062.792187 | 0.347180806 | 0.1275428245 | 0.003395414077 | 0.01443584243 |
| HSPA9 | 19860.30838 | 0.346920536 | 0.1338974076 | 0.005237223691 | 0.02065069479 |
| TTL | 3427.390712 | 0.3467698169 | 0.1205925639 | 0.002133899105 | 0.009680567877 |
| LRWD1 | 821.9381163 | 0.3467514211 | 0.1258078292 | 0.003066903797 | 0.0131974366 |
| DUSP10 | 1697.89432 | 0.3462119713 | 0.1321437261 | 0.004544921748 | 0.01839340579 |
| SEC62 | 3445.805379 | 0.3460516965 | 0.08603554145 | 3.17E-05 | 0.000238200706 |
| NOP58 | 4023.192007 | 0.3457828348 | 0.1585929251 | 0.01468357123 | 0.04851835153 |
| FKBP1A | 7785.791669 | 0.3453885134 | 0.0715324318 | 7.64E-07 | 8.04E-06 |
| AGFG1 | 2708.1318 | 0.344796999 | 0.1296975969 | 0.004131339358 | 0.01697506472 |
| CENPB | 3507.577107 | 0.3443577302 | 0.09375749339 | 0.000130702741 | 0.000851655634 |
| ARHGAP21 | 5123.186331 | 0.3440856536 | 0.1104066193 | 0.000987947833 | 0.004995307237 |
| TOM1 | 1692.520115 | 0.3440290594 | 0.1585384199 | 0.01522090483 | 0.04996121076 |
| DDX51 | 557.2985627 | 0.343929909 | 0.1075304153 | 0.000747956555 | 0.003906456906 |
| RAP1B | 4097.111933 | 0.3439119617 | 0.1169355625 | 0.001692670589 | 0.00796528837 |
| RREB1 | 603.5120513 | 0.3434385539 | 0.1158910514 | 0.001572389754 | 0.007489006138 |
| PLEKHG2 | 814.6464594 | 0.3431451736 | 0.1493214743 | 0.01105883042 | 0.03840796037 |
| KCMF1 | 2966.612291 | 0.3430445427 | 0.09243416522 | 0.000113901646 | 0.000757428143 |
| ZBED4 | 1108.802332 | 0.3429853067 | 0.1236497297 | 0.002961620643 | 0.01282220697 |
| LRIG3 | 1673.880191 | 0.3426394027 | 0.1189441557 | 0.002125743312 | 0.009655336233 |
| LINC01128 | 317.6730794 | 0.3425868159 | 0.114621582 | 0.001509620479 | 0.007227247368 |
| ZBTB39 | 598.2689416 | 0.3425153106 | 0.0897807067 | 7.57E-05 | 0.000523666791 |
| DKC1 | 4399.150394 | 0.3425052365 | 0.155424996 | 0.01410121488 | 0.04691686891 |
| HMGXB3 | 3137.752228 | 0.3419249075 | 0.09551039485 | 0.000180374656 | 0.001127455186 |
| TOMM40 | 3991.341471 | 0.3419019371 | 0.1565307637 | 0.01482263935 | 0.04888023859 |
| ZNF324 | 466.4316277 | 0.341704005 | 0.1232999786 | 0.002980233243 | 0.01288405713 |
| NKIRAS1 | 292.7900249 | 0.341464874 | 0.1562682337 | 0.01471565706 | 0.04859201997 |
| ST7 | 480.083364 | 0.3411110939 | 0.09440513615 | 0.000167702364 | 0.00105891455 |
| PVR | 3054.93444 | 0.3410841913 | 0.1385759469 | 0.00741128931 | 0.02744942451 |
| DDX46 | 3398.319036 | 0.3405655371 | 0.1100317581 | 0.001072715166 | 0.005368130236 |
| FAM101B | 2858.944655 | 0.3401740945 | 0.07956143459 | 1.08E-05 | 9.07E-05 |

| | baseMean | log2FoldChange | lfcSE | pvalue | padj |
|---|---|---|---|---|---|
| ALDH1B1 | 3255.128978 | 0.3400602504 | 0.09919684013 | 0.000343066466 | 0.001970485244 |
| LYPLA2 | 2000.571625 | 0.3397703112 | 0.1077746683 | 0.000887531277 | 0.004537930356 |
| VPS4B | 2376.73711 | 0.3396075101 | 0.06808537051 | 3.50E-07 | 3.97E-06 |
| C6orf47 | 901.8676677 | 0.3395989217 | 0.09273652186 | 0.000139281645 | 0.000898122310 |
| CHERP | 210.3143925 | 0.3392550354 | 0.1305663704 | 0.005009812126 | 0.0199228346 |
| FRMD6 | 9933.48602 | 0.3388231863 | 0.1463776359 | 0.01088774716 | 0.03791105415 |
| FNDC3A | 4270.220497 | 0.3385603657 | 0.08540091791 | 4.15E-05 | 0.000304409917 |
| EDNRA | 617.8975026 | 0.3384412868 | 0.1367112128 | 0.007086426209 | 0.02644362462 |
| NEK3 | 635.1204929 | 0.3380247689 | 0.1253069304 | 0.003778373032 | 0.01577571399 |
| RIOK2 | 1535.682225 | 0.3380183279 | 0.1244666843 | 0.003552656942 | 0.01498880964 |
| BTBD3 | 1008.460822 | 0.3377789469 | 0.1197704674 | 0.00261138433 | 0.01150322523 |
| KPNA4 | 4856.695047 | 0.3370362255 | 0.130742494 | 0.005343276696 | 0.02097997013 |
| PINX1 | 666.3515919 | 0.336953788 | 0.1350385777 | 0.006731475435 | 0.02537982724 |
| SCO2 | 902.1241106 | 0.3369456172 | 0.09397278127 | 0.000188659233 | 0.001170394717 |
| UBE2B | 684.4610621 | 0.3367395927 | 0.1015663373 | 0.000510017118 | 0.002794863772 |
| B4GALT1 | 989.7521027 | 0.3366169026 | 0.1317833598 | 0.005715686576 | 0.0222021543 |
| TMEM55A | 644.9057822 | 0.3364538909 | 0.08157225823 | 2.11E-05 | 0.000166008795 |
| ZNF134 | 670.7852423 | 0.3360079506 | 0.08528490917 | 4.63E-05 | 0.000335362072 |
| SETD3 | 2693.247842 | 0.3359089276 | 0.08303543603 | 2.98E-05 | 0.000226397405 |
| PSPC1 | 2151.159169 | 0.3357421535 | 0.07658021108 | 6.67E-06 | 5.81E-05 |
| TNPO1 | 12458.81628 | 0.335081527 | 0.128420996 | 0.004953083351 | 0.01973941596 |
| BAG1 | 906.1068718 | 0.3349906127 | 0.09232899754 | 0.000160973778 | 0.001020757489 |
| KIF1B | 2967.004418 | 0.334933059 | 0.1042695256 | 0.000735942525 | 0.00385282991 |
| TESK1 | 370.0156779 | 0.3342535287 | 0.1330173832 | 0.006457854259 | 0.0245283143 |
| SLC31A1 | 3156.686504 | 0.3341629499 | 0.09667256052 | 0.000308000211 | 0.001797653918 |
| CNNM4 | 258.8795109 | 0.3337530918 | 0.1246992316 | 0.004060612993 | 0.01673359171 |
| MIER1 | 1396.031244 | 0.3335089794 | 0.07743445301 | 9.56E-06 | 8.08E-05 |
| PRKAB1 | 1059.728946 | 0.3334631322 | 0.08922249087 | 0.000105693783 | 0.000708224456 |
| RBM7 | 1194.866059 | 0.3332850157 | 0.1010894195 | 0.000552826275 | 0.002999646655 |
| ARIH2 | 3444.244789 | 0.3331249929 | 0.07841708524 | 1.25E-05 | 0.000103567048 |
| DBR1 | 698.7720427 | 0.3330092169 | 0.1096893247 | 0.001340636953 | 0.00652098778 |
| CLUH | 1357.883265 | 0.3328640514 | 0.1323945001 | 0.00647180092 | 0.02456875163 |
| ANKRD40 | 2158.986325 | 0.3327462285 | 0.1123522974 | 0.001708159088 | 0.008022959199 |
| SIRPA | 1096.569375 | 0.3327419546 | 0.1129781386 | 0.001795869837 | 0.008366303505 |
| ACTR6 | 604.2476475 | 0.3326048253 | 0.1150821609 | 0.002149007467 | 0.009740204622 |
| TMEM161B | 365.0369701 | 0.3325269347 | 0.135451247 | 0.007630853229 | 0.0280810118 |
| DHX15 | 9882.295723 | 0.3323528913 | 0.1219796822 | 0.003546723104 | 0.01497650604 |
| SUPT4H1 | 2170.293517 | 0.3322515216 | 0.07023135074 | 1.30E-06 | 1.30E-05 |
| NXT2 | 237.599252 | 0.3317887037 | 0.1211060602 | 0.003399300725 | 0.01444824108 |
| DHX33 | 790.1665133 | 0.3316230179 | 0.1325718551 | 0.006742917739 | 0.02541653221 |

| | baseMean | log2FoldChange | lfcSE | pvalue | padj |
|---|---|---|---|---|---|
| SPCS3 | 5166.692468 | 0.3315564573 | 0.1375235447 | 0.008626181054 | 0.03116603002 |
| CAPN15 | 962.6192384 | 0.3314819954 | 0.1113297022 | 0.001592177288 | 0.007561699408 |
| RAB21 | 1132.388481 | 0.3313815655 | 0.1098647174 | 0.001436065577 | 0.006915129489 |
| IPO11-LRRC70 | 288.5291227 | 0.3311056009 | 0.1312370387 | 0.006343957654 | 0.02413264832 |
| MEX3C | 1833.05724 | 0.3307855744 | 0.1007847541 | 0.000542464592 | 0.002948797121 |
| RBM39 | 8432.002905 | 0.3302156842 | 0.06237031002 | 7.05E-08 | 9.08E-07 |
| ISY1-RAB43 | 1702.258575 | 0.3301207073 | 0.1208890992 | 0.00342014353 | 0.01452025008 |
| CCND3 | 3460.752774 | 0.3299690115 | 0.07985894836 | 2.10E-05 | 0.000165450462 |
| NUTM2A-AS1 | 249.2243211 | 0.3299284678 | 0.1231603699 | 0.004087143632 | 0.0168242968 |
| MPLKIP | 778.3424037 | 0.3294524476 | 0.1160803315 | 0.002539105761 | 0.01122802069 |
| ZNF639 | 988.9909627 | 0.3290621169 | 0.09586817541 | 0.000343137185 | 0.001970485244 |
| SCO1 | 1741.985136 | 0.3288666016 | 0.1058571989 | 0.00107305908 | 0.005368130236 |
| PAQR4 | 377.280928 | 0.328635973 | 0.1460770975 | 0.01316126449 | 0.04425430599 |
| IFT20 | 899.0374284 | 0.3285946923 | 0.09933592451 | 0.000537150015 | 0.002926317814 |
| PSMD7 | 5908.080633 | 0.3285104806 | 0.09878318957 | 0.000505868022 | 0.002776863608 |
| ZNF511 | 1151.960529 | 0.328261286 | 0.1132917187 | 0.00210443955 | 0.009579027959 |
| DDX10 | 2829.893215 | 0.3282390817 | 0.1188834966 | 0.003445400141 | 0.01460249436 |
| BRMS1 | 2865.918435 | 0.3281894464 | 0.1252791641 | 0.004914173482 | 0.01961585227 |
| EIF4E | 3853.405315 | 0.3276856544 | 0.1457440152 | 0.01361917993 | 0.04550627692 |
| VAPA | 6203.018848 | 0.3274350903 | 0.06961278301 | 1.52E-06 | 1.49E-05 |
| POM121C | 292.1369764 | 0.3273670173 | 0.1460776037 | 0.01356038124 | 0.04534033603 |
| SOCS7 | 704.9357576 | 0.3266037563 | 0.1005938344 | 0.000670984038 | 0.003548945414 |
| SDCBP | 5174.913668 | 0.3262678881 | 0.08271875257 | 4.77E-05 | 0.000344135928 |
| RNF219 | 1197.073623 | 0.3262008732 | 0.1267007821 | 0.005682349983 | 0.02209571922 |
| CCDC107 | 455.6055861 | 0.3259354177 | 0.1257158624 | 0.005309101577 | 0.02086779656 |
| STARD3NL | 2672.457366 | 0.3257734023 | 0.08127986055 | 3.60E-05 | 0.000268167250 |
| BNIP1 | 654.1992807 | 0.3250970224 | 0.1228608302 | 0.004612685662 | 0.01859687973 |
| ELF4 | 931.7365326 | 0.3250845407 | 0.08709883316 | 0.000111410422 | 0.000742187822 |
| TPBG | 1759.918031 | 0.3247865551 | 0.1210283238 | 0.004125825536 | 0.01696945205 |
| RIPK1 | 1672.732277 | 0.3244471156 | 0.1222759567 | 0.004492353476 | 0.01820044923 |
| SEC22B | 5430.130548 | 0.3244171497 | 0.08507379977 | 8.03E-05 | 0.000553532449 |
| GTF2IRD1 | 1312.337152 | 0.3241518833 | 0.1221600683 | 0.004530543354 | 0.01834020119 |
| CTDSPL2 | 1434.806115 | 0.3239915029 | 0.09003726708 | 0.000187775397 | 0.00116590672 |
| PHF5A | 1844.853428 | 0.3237354077 | 0.1072399737 | 0.001451928165 | 0.006982480119 |
| PVRL3 | 3262.113497 | 0.3236155198 | 0.1288219825 | 0.006665338775 | 0.02518787539 |
| SCARA3 | 3128.715419 | 0.3235380223 | 0.116637947 | 0.003285735768 | 0.01403768145 |
| RIOK3 | 2342.010506 | 0.3230886491 | 0.09306900479 | 0.000309024164 | 0.001802217306 |
| SLC31A2 | 244.2358499 | 0.3230409283 | 0.1318522723 | 0.007988155155 | 0.02917234292 |
| ATF5 | 392.4262234 | 0.3227824648 | 0.1071416609 | 0.001489171561 | 0.00714543196 |
| NMD3 | 2678.822712 | 0.3227176332 | 0.1097522474 | 0.001885397386 | 0.008742348701 |

| | baseMean | log2FoldChange | lfcSE | pvalue | padj |
|---|---|---|---|---|---|
| PPP1R2 | 1197.900476 | 0.3226395203 | 0.0767436585 | 1.56E-05 | 0.000127108469 |
| SUB1 | 5184.493134 | 0.3226292539 | 0.08090573181 | 4.00E-05 | 0.000294298119 |
| HNRNPF | 11068.7271 | 0.3224109681 | 0.07875559817 | 2.53E-05 | 0.000194833451 |
| UQCRHL | 509.9098132 | 0.3220510267 | 0.134031627 | 0.009091896977 | 0.03257984455 |
| ZNF513 | 688.1314699 | 0.3219002352 | 0.09630680033 | 4.84E-04 | 0.002673153828 |
| MOSPD1 | 1258.283005 | 0.3218777669 | 0.09823550217 | 0.000616446614 | 0.003299163782 |
| CAMSAP2 | 2354.785284 | 0.3212265729 | 0.1257134161 | 0.006018049564 | 0.0231412035 |
| MEF2A | 1009.123238 | 0.3209928041 | 0.1179898709 | 0.003753579105 | 0.01568137757 |
| FAM133B | 820.8054358 | 0.3208658779 | 0.1243141096 | 0.005600567613 | 0.02181188888 |
| RAB22A | 1437.831782 | 0.3205888455 | 0.1147543692 | 0.002992681907 | 0.0129303659 |
| PDE8A | 1364.874179 | 0.3201523091 | 0.09703012084 | 0.000570231066 | 0.003079495958 |
| FAM175B | 1102.35358 | 0.3199048222 | 0.0993000988 | 0.000747241310 | 0.003905116135 |
| OSER1 | 1497.228348 | 0.3197958617 | 0.09789692544 | 0.000619945098 | 0.003311603778 |
| PMF1 | 1777.638974 | 0.3190254766 | 0.09758290333 | 0.000634807632 | 0.003382838524 |
| AUP1 | 4074.10949 | 0.3189887062 | 0.1105843378 | 0.002273394374 | 0.0102168937 |
| ACTG1 | 178521.0475 | 0.3188162795 | 0.08078245125 | 6.90E-05 | 0.000480931485 |
| SPEN | 657.487427 | 0.3181187212 | 0.1333405625 | 0.009973890473 | 0.03522325789 |
| DHX30 | 3811.314166 | 0.3179115454 | 0.1077056759 | 0.001850479698 | 0.008604557223 |
| CARHSP1 | 4844.739425 | 0.3176862744 | 0.07748688511 | 2.49E-05 | 0.000192663460 |
| CGRRF1 | 366.4737725 | 0.3176124667 | 0.09993901353 | 0.000876346961 | 0.004486908496 |
| STAMBPL1 | 1177.83865 | 0.3174439653 | 0.1252249999 | 0.006599840005 | 0.02499110321 |
| RPIA | 487.6496749 | 0.3173825556 | 0.1257106697 | 0.006631967957 | 0.02508086637 |
| DNLZ | 414.662755 | 0.3168952249 | 0.143125202 | 0.01505584533 | 0.04949580064 |
| EIF2AK3 | 814.6656207 | 0.3167496143 | 0.1015763346 | 0.001078859109 | 0.005391840305 |
| COX11 | 886.5080247 | 0.3162160184 | 0.09955043116 | 0.000882762517 | 0.004515098291 |
| EPN2 | 1493.60846 | 0.3158265315 | 0.08809055975 | 0.000202781546 | 0.001247609277 |
| BAG2 | 1987.754894 | 0.3155988784 | 0.1341572125 | 0.01063240128 | 0.03719591697 |
| FOPNL | 2089.883459 | 0.3154768165 | 0.0850240148 | 0.000124773862 | 0.000818395613 |
| CD9 | 3327.1189 | 0.3153367346 | 0.08810533606 | 0.000207625128 | 0.001271106308 |
| KIF7 | 770.7225237 | 0.3149171792 | 0.09469034547 | 0.000526253958 | 0.002874319582 |
| PITHD1 | 1381.482046 | 0.3147250506 | 0.1385409437 | 0.01309229772 | 0.0440721729 |
| STX5 | 2066.673987 | 0.3140624654 | 0.07836347944 | 3.73E-05 | 0.000276127530 |
| TAB2 | 1431.487476 | 0.3135380123 | 0.1064172853 | 0.001910277576 | 0.008835701409 |
| DDX27 | 2496.289558 | 0.3133440229 | 0.09797585542 | 0.000830517147 | 0.004286801031 |
| SBDS | 3742.25729 | 0.3132099838 | 0.09374289281 | 0.000502669322 | 0.002761713484 |
| MTMR2 | 3115.939759 | 0.312169856 | 0.09596955951 | 0.000671957611 | 0.003552514807 |
| SNX8 | 1250.598448 | 0.3121022189 | 0.1066670157 | 0.002043426038 | 0.009349898673 |
| SIK3 | 660.1490996 | 0.3119895371 | 0.1177637187 | 0.004755824973 | 0.01909640723 |
| RAP2C | 947.8985448 | 0.3119562456 | 0.0805789654 | 6.69E-05 | 0.000467788440 |
| TMEM184B | 1238.768462 | 0.3119528588 | 0.08712424992 | 0.000209182028 | 0.001280111474 |

| | baseMean | log2FoldChange | lfcSE | pvalue | padj |
|---|---|---|---|---|---|
| ZNF598 | 1118.913312 | 0.3117504027 | 0.1142575966 | 0.003772829142 | 0.01575698544 |
| FHL2 | 6823.9702 | 0.311695292 | 0.1085813679 | 0.002662274835 | 0.0116997078 |
| RPP40 | 409.5118964 | 0.3114515898 | 0.1290569276 | 0.009184115623 | 0.03287074459 |
| CCSER2 | 1504.495377 | 0.3114447422 | 0.1098439159 | 0.002641555308 | 0.01161551004 |
| NRAS | 5779.038328 | 0.3113686006 | 0.1197213672 | 0.005488521245 | 0.02141472558 |
| CSNK1G1 | 1271.213604 | 0.3113480026 | 0.08798369097 | 0.000245285520 | 0.001473197305 |
| CSRNP2 | 1144.550701 | 0.3113178798 | 0.08676800237 | 0.000203530294 | 0.001250665517 |
| EOGT | 846.0541031 | 0.3113116225 | 0.111165066 | 0.003033136156 | 0.01306723502 |
| WASF3 | 396.9176857 | 0.3111091212 | 0.1337811799 | 0.01161001686 | 0.04000498519 |
| ZNF195 | 804.7426159 | 0.3108662751 | 0.1181415926 | 0.00502245869 | 0.01996779371 |
| RBM12 | 5226.026455 | 0.3105885449 | 0.1107379807 | 0.002965388198 | 0.01283105635 |
| RCOR1 | 976.1787248 | 0.310418317 | 0.09495940592 | 0.000654728550 | 0.003476552566 |
| PDE12 | 1131.001965 | 0.3103862817 | 0.1069203586 | 0.002219368394 | 0.01001338667 |
| PPP2R2D | 1139.765866 | 0.3101256473 | 0.09897287643 | 0.001043667308 | 0.005237331495 |
| NSFL1C | 2330.713846 | 0.3099680422 | 0.09370360561 | 0.000595708675 | 0.003201987895 |
| TCEB1 | 2261.208849 | 0.3099260745 | 0.124341761 | 0.007449410432 | 0.02754265506 |
| SPPL3 | 426.2636576 | 0.309815499 | 0.1183890053 | 0.005247982769 | 0.02067122102 |
| MAPKAPK5 | 1374.355777 | 0.3096940236 | 0.1085828241 | 0.002598706843 | 0.0114573904 |
| ZNF286A | 1644.155771 | 0.3094932175 | 0.07952654624 | 6.14E-05 | 0.000433339220 |
| BRAP | 1030.298716 | 0.3092881931 | 0.07761962835 | 4.18E-05 | 0.000305777927 |
| MRPS23 | 2704.463656 | 0.3088768183 | 0.1257921976 | 0.00828097341 | 0.03009407203 |
| FURIN | 1515.733783 | 0.3088602721 | 0.08992589905 | 0.000364076325 | 0.002072937822 |
| GNA12 | 3379.689415 | 0.3087657167 | 0.07018241291 | 6.78E-06 | 5.89E-05 |
| SLC3A2 | 9370.656187 | 0.308586568 | 0.08997490231 | 0.000370548178 | 0.002102550239 |
| C9orf89 | 1084.164987 | 0.3083375459 | 0.1026966124 | 0.001623710636 | 0.00768449703 |
| PI4KB | 3779.10006 | 0.3081744196 | 0.06132745161 | 2.57E-07 | 2.97E-06 |
| FAM219A | 958.0451382 | 0.3080533119 | 0.07690039754 | 3.84E-05 | 0.000283642245 |
| ORMDL3 | 1583.669593 | 0.3079429984 | 0.09871131661 | 0.001100529411 | 0.005485698826 |
| ZNF622 | 1909.714698 | 0.3078736908 | 0.07488471098 | 2.44E-05 | 0.000189346691 |
| PUS1 | 659.8145579 | 0.3076717893 | 0.1133680348 | 0.00397568365 | 0.0164381988 |
| ABCF1 | 3913.668654 | 0.3073027418 | 0.1087116301 | 0.002757353403 | 0.01205350406 |
| TMEM132A | 1350.470059 | 0.3069723199 | 0.1269952472 | 0.009217031019 | 0.03296477897 |
| MRPS35 | 2667.115517 | 0.30681564 | 0.08514116627 | 0.000194344014 | 0.001202155391 |
| PRCC | 1047.786731 | 0.3066167867 | 0.09185030985 | 0.000519080327 | 0.002840348031 |
| ASNSD1 | 2107.419182 | 0.3065586119 | 0.1019541575 | 0.001595929165 | 0.007577101191 |
| CMTM7 | 670.2825196 | 0.3065085893 | 0.09375683718 | 0.000660382215 | 0.003500331367 |
| SDF4 | 3014.21525 | 0.3064350364 | 0.09329804317 | 0.000629273289 | 0.003356950914 |
| UBE3A | 2690.818304 | 0.3064111097 | 0.1073853455 | 0.002618845454 | 0.01153267967 |
| FAM3C | 1426.761153 | 0.3063692037 | 0.1273796068 | 0.009617318612 | 0.03415031166 |
| FUBP1 | 8407.172523 | 0.3057894129 | 0.1223612505 | 0.007350269567 | 0.02727470714 |

| | baseMean | log2FoldChange | lfcSE | pvalue | padj |
|---|---|---|---|---|---|
| TSPYL1 | 4068.094491 | 0.3051750261 | 0.05364164106 | 8.13E-09 | 1.24E-07 |
| C8orf76 | 718.3731932 | 0.3049423188 | 0.07973024316 | 8.19E-05 | 0.000563114463 |
| KRBOX4 | 488.6945302 | 0.3048670442 | 0.09785023511 | 0.001126962336 | 0.005613697633 |
| STAT5B | 2000.464815 | 0.3047008393 | 0.1144730176 | 0.004691741735 | 0.01886962255 |
| BOD1L1 | 2982.706362 | 0.3044741374 | 0.11537038 | 0.005001250455 | 0.0198994169 |
| HPCAL1 | 656.4434149 | 0.3042131457 | 0.1269002677 | 0.009821668953 | 0.03475162287 |
| KLHDC3 | 2785.048471 | 0.303634363 | 0.09060381879 | 0.000499907895 | 0.002747555797 |
| TMEM165 | 2244.09654 | 0.3034436928 | 0.07549670898 | 3.67E-05 | 0.000272194591 |
| TBC1D12 | 449.9356715 | 0.3022884538 | 0.1172037319 | 0.006002718586 | 0.02310014914 |
| PHLDA3 | 2338.853974 | 0.3020187811 | 0.1197263399 | 0.007036368103 | 0.02630951398 |
| MAPRE1 | 6250.414878 | 0.3019937609 | 0.1208993937 | 0.007900413984 | 0.02891574823 |
| ZBTB9 | 444.085608 | 0.3019262817 | 0.114815535 | 0.005204104822 | 0.02055276305 |
| GIT1 | 983.9779175 | 0.3015971134 | 0.1001385063 | 0.00160795145 | 0.00762445514 |
| SF3B14 | 3423.902517 | 0.3014128792 | 0.08260677433 | 0.000166199443 | 0.001050761579 |
| TVP23B | 1421.011451 | 0.3012704916 | 0.1163517766 | 0.00582082712 | 0.02252242593 |
| FBXW7 | 881.7351121 | 0.3011612315 | 0.07571663925 | 4.42E-05 | 0.000321901895 |
| TEAD4 | 682.5663546 | 0.299983577 | 0.09838190723 | 0.001422180396 | 0.006854918718 |
| ARHGAP42 | 546.5739114 | 0.2995216102 | 0.1298850544 | 0.0127231651 | 0.04305345571 |
| CEP170B | 1041.203477 | 0.2995195298 | 0.1124887538 | 0.004757582234 | 0.01909831272 |
| PSMG3 | 1157.892118 | 0.2994528076 | 0.09063943106 | 0.000597792161 | 0.003207397295 |
| CRK | 4101.779467 | 0.2992845723 | 0.07806086282 | 8.07E-05 | 0.000555463587 |
| CDV3 | 11260.01463 | 0.299142223 | 0.1216318665 | 0.008467157589 | 0.03065844099 |
| PSMG2 | 2068.869697 | 0.2990612107 | 0.06344141736 | 1.56E-06 | 1.54E-05 |
| FOXJ2 | 354.5708905 | 0.2987936071 | 0.1183034775 | 0.007067420405 | 0.02639965513 |
| TMEM199 | 823.0848373 | 0.2987565076 | 0.1128186291 | 0.004974979249 | 0.01980017804 |
| TRAF3IP2 | 718.166996 | 0.2985297888 | 0.08476936201 | 0.000269858826 | 0.001600768156 |
| ZDHHC5 | 3111.610184 | 0.2982173546 | 0.07385934833 | 3.47E-05 | 0.000258879979 |
| PLEKHM2 | 2594.247253 | 0.2976195306 | 0.08583309228 | 0.000333616555 | 0.001924531926 |
| PPP6R3 | 5181.379869 | 0.2975690262 | 0.08110642356 | 0.000155392252 | 0.000988310658 |
| NACC1 | 2591.052409 | 0.2975500914 | 0.1015608026 | 0.002116735122 | 0.009617354053 |
| MGAT1 | 5400.476351 | 0.2971046407 | 0.07314615607 | 3.11E-05 | 0.000234598893 |
| U2SURP | 4396.010006 | 0.2969193774 | 0.122129358 | 0.009489543611 | 0.03375294191 |
| ALG5 | 1083.225059 | 0.2969050335 | 0.08036376329 | 0.000140943705 | 0.000906875899 |
| BICD2 | 2447.602593 | 0.2968816352 | 0.1129053426 | 0.005293077264 | 0.02082106004 |
| RP2 | 1041.055463 | 0.2967092891 | 0.103710948 | 0.002637503496 | 0.01160111951 |
| ZFYVE27 | 742.9410734 | 0.2961471387 | 0.0870406688 | 0.000423366328 | 0.002379577678 |
| RAB7A | 4006.751228 | 0.296065959 | 0.05657631175 | 1.09E-07 | 1.37E-06 |
| BAG3 | 1761.636383 | 0.2957039146 | 0.1063355366 | 0.003389293633 | 0.01441393685 |
| ASB7 | 604.1453935 | 0.29549836 | 0.07961303711 | 0.000132112001 | 0.000860461761 |
| DCUN1D1 | 1146.141805 | 0.2954561475 | 0.1115823348 | 0.005049700542 | 0.02005467894 |

| | baseMean | log2FoldChange | lfcSE | pvalue | padj |
|---|---|---|---|---|---|
| CRBN | 926.5119271 | 0.2954394402 | 0.1177073875 | 0.007441686592 | 0.02752093186 |
| ZBTB8OS | 732.1647878 | 0.2952262157 | 0.08265340352 | 2.27E-04 | 0.001373036229 |
| NXF1 | 1743.426963 | 0.2950011674 | 0.08258163971 | 0.000227492148 | 0.001375763851 |
| SYF2 | 1266.535289 | 0.294801419 | 0.1123602992 | 0.005438556705 | 0.02125877416 |
| LIN7C | 1920.070677 | 0.294790712 | 0.1137334723 | 0.005946158136 | 0.02292990119 |
| KCTD20 | 4157.52599 | 0.2946830775 | 0.06739863375 | 7.99E-06 | 6.88E-05 |
| BMPR1A | 1909.014753 | 0.2945504564 | 0.1066712901 | 0.003609564676 | 0.01518158431 |
| LDLRAD3 | 538.4410511 | 0.294359279 | 0.1240956144 | 0.01091632828 | 0.03798392421 |
| CIR1 | 900.1198934 | 0.2938827057 | 0.1055134218 | 0.003428794671 | 0.01455282892 |
| USP10 | 4128.369408 | 0.2936882871 | 0.1160648202 | 0.007119265049 | 0.02653949357 |
| KDM2B | 966.5978526 | 0.2936538746 | 0.09089803821 | 0.000788730647 | 0.004100352867 |
| PRNP | 6655.931296 | 0.2936407204 | 0.1022047774 | 0.002458035579 | 0.0109312102 |
| PAFAH1B2 | 5327.136716 | 0.2935007209 | 0.0824152575 | 0.00023757791 | 0.001432104252 |
| OAT | 5963.969906 | 0.2934956133 | 0.06801654185 | 1.06E-05 | 8.87E-05 |
| SLC37A3 | 1554.009258 | 0.2930421631 | 0.1088626311 | 0.004456681684 | 0.01808052686 |
| NDUFS5 | 9870.231958 | 0.2929965892 | 0.07532341406 | 6.51E-05 | 0.000456816868 |
| MTMR12 | 1142.743385 | 0.292816307 | 0.1293266974 | 0.01460330634 | 0.0482745622 |
| TYW3 | 1783.491907 | 0.2919168279 | 0.1030927052 | 0.002962668455 | 0.01282301472 |
| ZNF484 | 482.8615756 | 0.2917150349 | 0.1261653983 | 0.0128322364 | 0.04334373134 |
| MRPS22 | 2142.31612 | 0.2915536523 | 0.07244352877 | 3.86E-05 | 0.000284542654 |
| POLR2K | 1457.983176 | 0.2912512671 | 0.09702786304 | 0.001714288456 | 0.008044135148 |
| TSFM | 1390.382922 | 0.290990706 | 0.08577018657 | 0.000446823367 | 0.002498217466 |
| WDFY1 | 3624.402685 | 0.2906639395 | 0.08118577395 | 0.000223718704 | 0.001357177362 |
| ITGB5 | 9502.351305 | 0.2904057656 | 0.06438610802 | 4.27E-06 | 3.85E-05 |
| ROBO3 | 413.459459 | 0.2903797135 | 0.1179520123 | 0.008636622563 | 0.0311961847 |
| UBE2G2 | 2077.208735 | 0.2901995889 | 0.1154852263 | 0.007519475616 | 0.02776034526 |
| PSMC1 | 9019.10151 | 0.2898131027 | 0.1160281103 | 0.007939998242 | 0.02901782863 |
| TMX4 | 1457.178007 | 0.2896121831 | 0.07544950513 | 8.12E-05 | 0.000558238593 |
| ZNF593 | 724.5984474 | 0.2893577386 | 0.08017125296 | 0.000200310581 | 0.001234958279 |
| CASP4 | 4622.224941 | 0.2892860648 | 0.08777661573 | 0.000620105536 | 0.003311603778 |
| SZRD1 | 7545.678317 | 0.288839751 | 0.09263761771 | 0.00107916476 | 0.005391840305 |
| MYL12B | 15195.46137 | 0.2886524617 | 0.06292760552 | 3.55E-06 | 3.24E-05 |
| ZNF557 | 629.8145101 | 0.2886352208 | 0.1145692653 | 0.007433169578 | 0.02750380439 |
| MGRN1 | 437.7101848 | 0.2883322029 | 0.1104584068 | 0.005748369248 | 0.02229420936 |
| ST3GAL4 | 1093.487824 | 0.2880660644 | 0.09907250377 | 0.002340297381 | 0.01046072882 |
| PUF60 | 6842.473621 | 0.287767214 | 0.09972363997 | 0.002512199354 | 0.01112887122 |
| TEN1 | 451.6124393 | 0.2876295553 | 0.1210333684 | 0.01095581794 | 0.03811242367 |
| SLBP | 2674.100187 | 0.2873552822 | 0.1102144044 | 0.005814910871 | 0.02250538289 |
| NOB1 | 2448.568199 | 0.2865402727 | 0.07057839083 | 3.25E-05 | 0.000243768167 |
| STRN4 | 2390.970356 | 0.2855936982 | 0.08881646019 | 0.000856678239 | 0.004401339649 |

| | baseMean | log2FoldChange | lfcSE | pvalue | padj |
|---|---|---|---|---|---|
| HNRNPA0 | 3610.64346 | 0.2854826092 | 0.07669471252 | 0.000130455491 | 0.000850789231 |
| 15-Sep | 3779.330591 | 0.2854492111 | 0.06239037803 | 3.18E-06 | 2.94E-05 |
| ISOC1 | 1350.709395 | 0.2852545354 | 0.09406167865 | 0.001576985292 | 0.00750630883 |
| SF3B5 | 4736.06779 | 0.2848408793 | 0.09993013862 | 0.002876475176 | 0.01251178466 |
| GOLGA7 | 1926.328513 | 0.2845675814 | 0.07558541808 | 0.000109771702 | 0.000732582196 |
| FNBP4 | 1811.209772 | 0.2845271149 | 0.1019491711 | 0.003401096713 | 0.01445146253 |
| GFOD2 | 766.8418259 | 0.2844376676 | 0.09751623869 | 0.002300303485 | 0.01031602969 |
| SMCR8 | 955.2746589 | 0.2843505805 | 0.0804042582 | 0.000268549014 | 0.001594904779 |
| LINC00667 | 783.1639304 | 0.2841070407 | 0.08251158979 | 0.000379254475 | 0.002147858458 |
| MMADHC | 5544.318177 | 0.2840418167 | 0.08368083685 | 0.000451455420 | 0.002517498037 |
| FARP2 | 644.732422 | 0.2839022829 | 0.09821662088 | 0.002506620496 | 0.0111135882 |
| USP11 | 1939.71096 | 0.2838558479 | 0.08802726439 | 0.000833156671 | 0.004295314987 |
| RPS7 | 52176.43109 | 0.2836413116 | 0.06977567394 | 3.24E-05 | 0.000243125511 |
| ARHGEF1 | 2004.23924 | 0.2835154896 | 0.0954945015 | 0.001952866823 | 0.00899914396 |
| CHUK | 1458.674126 | 0.2830024158 | 0.1225259151 | 0.01213371682 | 0.04151169802 |
| MRPL44 | 1341.840028 | 0.2828852451 | 0.08837667276 | 0.000903604480 | 0.004609032925 |
| EWSR1 | 13433.4843 | 0.2826708987 | 0.09406427639 | 0.001622869488 | 0.00768449703 |
| SPOPL | 556.2573007 | 0.2826282914 | 0.1195947426 | 0.01158091201 | 0.03991393494 |
| TRIM11 | 618.9754241 | 0.2826099622 | 0.1032615225 | 0.004026790602 | 0.01661720767 |
| PSMB7 | 8121.900384 | 0.282470938 | 0.1201082854 | 0.01189979272 | 0.04083337494 |
| CHMP1A | 3490.853189 | 0.2823463935 | 0.0811614419 | 0.000335206344 | 0.001932954015 |
| CLPX | 1723.916482 | 0.2819399998 | 0.06559271358 | 1.16E-05 | 9.61E-05 |
| UQCRFS1 | 4072.351326 | 0.2814329813 | 0.09704233865 | 0.002399823389 | 0.01069469334 |
| PPP6C | 1813.518187 | 0.2811546049 | 0.08185520801 | 0.000396303279 | 0.002237603161 |
| CANT1 | 2150.013489 | 0.2808376806 | 0.08612023422 | 0.000737915774 | 0.003861802448 |
| GNG5 | 2580.086821 | 0.280685968 | 0.07047292981 | 4.59E-05 | 0.000332733828 |
| IDS | 1589.652867 | 0.280184785 | 0.0826305159 | 0.000466555147 | 0.002584278123 |
| TRAF6 | 690.5024127 | 0.2801064501 | 0.1165883769 | 0.01051213782 | 0.03681844741 |
| UBLCP1 | 931.5693426 | 0.2800720138 | 0.1020760096 | 0.003984098993 | 0.01646384954 |
| FBXL3 | 1323.590959 | 0.2799962555 | 0.08926623702 | 0.001135890329 | 0.005652497027 |
| MORF4L1 | 8269.010454 | 0.2799811111 | 0.07300514254 | 8.43E-05 | 0.000577362956 |
| YAP1 | 4519.499684 | 0.279819325 | 0.06826898772 | 2.85E-05 | 0.000217779627 |
| PEX13 | 969.2408039 | 0.2797769923 | 0.09865572591 | 0.003009054065 | 0.01298979587 |
| TMEM222 | 1224.6848 | 0.2794685662 | 0.09251275954 | 0.001668269061 | 0.007870360597 |
| RERE | 576.6288579 | 0.2794441641 | 0.111923558 | 0.008131269413 | 0.02962958157 |
| IDH3B | 3368.42384 | 0.2793702304 | 0.093952868 | 0.001956219041 | 0.00900622922 |
| COA6 | 688.7316817 | 0.2792247466 | 0.1078788383 | 0.006288989143 | 0.02396027619 |
| PDCD10 | 1316.315062 | 0.2791855798 | 0.1100465353 | 0.007053663504 | 0.02636094275 |
| TRAF2 | 689.2444743 | 0.2791305049 | 0.1223837987 | 0.01449687407 | 0.04799732221 |
| SNRPB2 | 2308.212548 | 0.2790586008 | 0.1182841138 | 0.01182211287 | 0.04060425342 |

| | baseMean | log2FoldChange | lfcSE | pvalue | padj |
|---|---|---|---|---|---|
| KHSRP | 3209.194481 | 0.2786847058 | 0.1045844054 | 0.005062685882 | 0.02010040635 |
| PPP6R1 | 1786.423197 | 0.2779228823 | 0.09496360194 | 0.002294373846 | 0.01029253757 |
| ATP9A | 1414.03227 | 0.2778583297 | 0.0980592703 | 0.003022321064 | 0.01303572953 |
| MED17 | 1300.8932 | 0.2775906418 | 0.08796199318 | 0.001073337613 | 0.005368130236 |
| DYNC1LI1 | 2387.023941 | 0.2775863992 | 0.1031798904 | 0.004665474134 | 0.01878935471 |
| 5-Mar | 929.9477962 | 0.2769760781 | 0.09040032976 | 0.00146246625 | 0.007028618464 |
| POMP | 4373.428109 | 0.2764248653 | 0.1158868324 | 0.01112815262 | 0.03862169331 |
| COX7B | 3135.109426 | 0.2762701824 | 0.1015286541 | 0.004314038834 | 0.01757846858 |
| SWAP70 | 2293.212391 | 0.2762389369 | 0.08609035428 | 0.000896522622 | 0.004577614993 |
| SPTSSA | 912.9180893 | 0.2761399671 | 0.07876920497 | 0.000307806023 | 0.001797225051 |
| TOMM20 | 7750.363255 | 0.2760657542 | 0.06832656705 | 3.62E-05 | 0.000269292952 |
| CDYL | 1720.389526 | 0.2759965086 | 0.08955208254 | 0.001381982019 | 0.006695844544 |
| SMAD3 | 2925.51128 | 0.2751005173 | 0.1144041705 | 0.01004370658 | 0.03541102233 |
| PPP2CB | 2892.530996 | 0.275070772 | 0.09680921443 | 0.003014144842 | 0.0130042314 |
| ZNF496 | 1875.707591 | 0.2750618851 | 0.109129022 | 0.007738751451 | 0.02840785758 |
| C10orf32 | 397.9401129 | 0.2745697283 | 0.100484407 | 0.004181359489 | 0.01712805529 |
| LMNA | 29084.2798 | 0.2742900932 | 0.1171444615 | 0.01175997935 | 0.04041882099 |
| FBXO31 | 1065.480666 | 0.2738587499 | 0.09076926258 | 0.001718645813 | 0.008059501578 |
| DDX24 | 6578.771368 | 0.2737388815 | 0.08714927162 | 0.001146579933 | 0.005698073637 |
| GNA11 | 1969.853858 | 0.2736697066 | 0.1118903655 | 0.009532485731 | 0.03387865529 |
| WDR1 | 29822.97435 | 0.2728275257 | 0.07354382365 | 0.000142441828 | 0.000915723825 |
| POMK | 400.651215 | 0.2726965117 | 0.1106874807 | 0.009079954383 | 0.03255271871 |
| SEC61B | 4192.86845 | 0.2726042461 | 0.07856922133 | 0.000357924680 | 0.002045735344 |
| PAXIP1 | 685.5791546 | 0.2720915862 | 0.1082232025 | 0.00794390132 | 0.02902496853 |
| PSMA3 | 4590.810499 | 0.2719660183 | 0.1001574279 | 0.004018001755 | 0.01658991378 |
| MAP3K4 | 2196.489269 | 0.2717065173 | 0.0880677467 | 0.001400437675 | 0.006767891031 |
| WDR74 | 1681.30918 | 0.2714187283 | 0.1091679817 | 0.008596177793 | 0.03106516776 |
| PSMC2 | 6216.783388 | 0.2712050438 | 0.1059919289 | 0.007024470293 | 0.02627162477 |
| BCLAF1 | 9450.524525 | 0.2710349511 | 0.1174021041 | 0.01411872446 | 0.04695414082 |
| TMEM127 | 1141.932383 | 0.2709848743 | 0.1184114791 | 0.0146024971 | 0.0482745622 |
| SP3 | 3657.652793 | 0.2708814224 | 0.06252569292 | 1.04E-05 | 8.69E-05 |
| ENY2 | 2743.008206 | 0.2704843707 | 0.08682635633 | 0.001259677025 | 0.006183755763 |
| RAB31 | 2813.184417 | 0.2691701596 | 0.1091193351 | 0.009143296163 | 0.03274038878 |
| NHP2L1 | 6543.94974 | 0.2687748528 | 0.1020277392 | 0.005892797318 | 0.02275359421 |
| MED27 | 1063.423045 | 0.268505483 | 0.1038613113 | 0.006512512188 | 0.02471700076 |
| ZC3H14 | 3469.250474 | 0.2680813972 | 0.09020427215 | 0.002063064883 | 0.009421410088 |
| BRD1 | 881.2535869 | 0.2680384033 | 0.09897665713 | 0.004595806654 | 0.01855898163 |
| DLGAP4 | 808.0080602 | 0.2678800567 | 0.09088448027 | 0.002183997072 | 0.009874744124 |
| B3GAT3 | 1672.895745 | 0.2674947063 | 0.1001562506 | 0.005139081923 | 0.02036619397 |
| NCS1 | 4851.577092 | 0.2672034657 | 0.08563424564 | 0.001243444024 | 0.006116167188 |

| | baseMean | log2FoldChange | lfcSE | pvalue | padj |
|---|---|---|---|---|---|
| MAP2K4 | 2011.122484 | 0.2670808676 | 0.1026985064 | 0.006308046998 | 0.02402673619 |
| BUD31 | 4041.996765 | 0.2668863992 | 0.05835334882 | 3.37E-06 | 3.10E-05 |
| SMAP2 | 3045.744767 | 0.2668824883 | 0.0755008796 | 0.000283985655 | 0.001673896445 |
| MRPL14 | 2195.072128 | 0.266697168 | 0.1080121933 | 0.009125608751 | 0.03268491429 |
| FAM168B | 4678.889728 | 0.2665868501 | 0.05321025071 | 3.78E-07 | 4.26E-06 |
| QRICH1 | 2959.845218 | 0.2648068501 | 0.08953436919 | 0.0021470501 | 0.009734296267 |
| YTHDF3 | 2764.314172 | 0.2641843646 | 0.1124518069 | 0.01272202244 | 0.04305345571 |
| PSMD6 | 4040.234236 | 0.2640604257 | 0.1041065473 | 0.00753259216 | 0.02778225935 |
| EIF2A | 5220.157806 | 0.2639307966 | 0.08958204716 | 0.002229794523 | 0.01004216898 |
| KIAA1462 | 3187.49481 | 0.2637175958 | 0.09710182576 | 0.004488201096 | 0.01818857543 |
| SMU1 | 4744.494832 | 0.2635929855 | 0.07563356052 | 0.000344324528 | 0.001975586859 |
| STX6 | 1548.833513 | 0.2634292065 | 0.09869547752 | 0.005185904175 | 0.02049719333 |
| RPS15A | 30448.03604 | 0.2632360799 | 0.07675152007 | 0.000385970981 | 0.002184234871 |
| CAPZA2 | 4779.22926 | 0.263064408 | 0.08809904681 | 0.001904835476 | 0.008818748572 |
| MRPL32 | 1938.224849 | 0.2630511089 | 0.09450941588 | 0.003719065715 | 0.01556737965 |
| ATP5G3 | 7848.733631 | 0.2628915156 | 0.08748535249 | 0.001628521608 | 0.007702369192 |
| DRG1 | 4218.818378 | 0.2620327166 | 0.09432992804 | 0.003784151736 | 0.01578656072 |
| DCTN6 | 1245.697903 | 0.2616412754 | 0.06340663852 | 2.63E-05 | 0.000202010554 |
| GTF2A2 | 1614.242577 | 0.2613238986 | 0.111675231 | 0.01314934711 | 0.04423421354 |
| USP8 | 2516.386258 | 0.2613019189 | 0.083245192 | 0.00118992714 | 0.005887944563 |
| BLOC1S2 | 1478.885757 | 0.261011833 | 0.0789997483 | 0.000673502378 | 0.003558472998 |
| SLC9B2 | 514.9104826 | 0.2608094199 | 0.1043368693 | 0.008551735418 | 0.03093459394 |
| NCOA4 | 10992.27244 | 0.2607843479 | 0.06086281834 | 1.26E-05 | 0.000104243896 |
| GSK3A | 1005.566166 | 0.2606223852 | 0.1044404791 | 0.008664127097 | 0.03127277293 |
| RPL34 | 20616.48232 | 0.2605604803 | 0.07280517617 | 0.000245416401 | 0.001473389034 |
| CD63 | 24800.89728 | 0.2600540469 | 0.05558493849 | 2.12E-06 | 2.03E-05 |
| ZNF140 | 615.8773921 | 0.2597245194 | 0.08083758366 | 0.000923823404 | 0.004700078115 |
| EIF4A3 | 3936.525391 | 0.2590661996 | 0.07262981107 | 0.000256701814 | 0.001528202047 |
| RAB11FIP3 | 947.1940653 | 0.2589160028 | 0.1054373097 | 0.00978465959 | 0.03463713662 |
| TERF2 | 580.92053 | 0.2587584189 | 0.08061754942 | 0.000939185990 | 0.004770911023 |
| ACAA2 | 3689.777775 | 0.2584515216 | 0.103526658 | 0.008694529562 | 0.03135970219 |
| FAM102B | 1083.298211 | 0.2580262963 | 0.1123113225 | 0.01484905899 | 0.04895651888 |
| DYNLT1 | 2720.817051 | 0.2579403217 | 0.08047038679 | 0.000961654866 | 0.00486843907 |
| DIP2C | 1010.338969 | 0.2577902629 | 0.09752111086 | 0.005706588777 | 0.02217838692 |
| MTMR14 | 1565.49979 | 0.2576557693 | 0.07520594846 | 0.000436141656 | 0.002444922108 |
| LETM1 | 2942.558954 | 0.2571931769 | 0.1049861747 | 0.009964624602 | 0.03519888391 |
| MPZL1 | 3771.950771 | 0.257109125 | 0.1083308549 | 0.01215419255 | 0.04156264881 |
| PFDN1 | 3485.326551 | 0.2569451059 | 0.1098623029 | 0.01336669481 | 0.0447933196 |
| SRA1 | 2447.971179 | 0.2569404839 | 0.0873869156 | 0.002314361631 | 0.0103541257 |
| KIAA0947 | 3366.816273 | 0.2567894431 | 0.1095642154 | 0.01320558833 | 0.04438329678 |

| | baseMean | log2FoldChange | lfcSE | pvalue | padj |
|---|---|---|---|---|---|
| KCTD6 | 372.1611112 | 0.2567848769 | 0.1095738041 | 0.01321567772 | 0.04439716281 |
| ACTN1 | 25215.37475 | 0.2565386651 | 0.06472644686 | 5.15E-05 | 3.70E-04 |
| ADIPOR2 | 2982.859929 | 0.2564967735 | 0.07425221642 | 0.000391864463 | 0.002215060742 |
| TUBB2A | 5098.16614 | 0.2564263145 | 0.08048738135 | 0.00102558085 | 0.005163284206 |
| POLR2F | 1921.215217 | 0.2561775864 | 0.1110805535 | 0.0145816939 | 0.04823524561 |
| PARL | 1284.68893 | 0.2560968913 | 0.07373140525 | 0.000367621864 | 0.002089130513 |
| LYRM4 | 1418.620873 | 0.2554571011 | 0.09387890523 | 0.004582036984 | 0.01852347235 |
| HSPA13 | 3177.276468 | 0.2554294471 | 0.09961131635 | 0.00731791421 | 0.02717117822 |
| RPS29 | 18709.24112 | 0.2549553616 | 0.08694222085 | 0.002052930315 | 0.009384734254 |
| PPIG | 2259.682222 | 0.2549488335 | 0.09706988599 | 0.006070877935 | 0.02332025325 |
| YTHDF1 | 999.6183084 | 0.2541970414 | 0.0921845363 | 0.004129201373 | 0.01697506472 |
| NCLN | 2641.312565 | 0.2540605037 | 0.105972827 | 0.01155743718 | 0.03984225102 |
| CWC22 | 1110.660757 | 0.2539484176 | 0.103235679 | 0.00974830735 | 0.03454952348 |
| CDK13 | 1842.4804 | 0.2538674709 | 0.087979787 | 0.002792102983 | 0.01219109129 |
| GRSF1 | 4704.209281 | 0.2531546564 | 0.09349805674 | 0.004803660777 | 0.01922626487 |
| NAA20 | 2174.948459 | 0.2531454624 | 0.09860337876 | 0.007376184395 | 0.02735342701 |
| ZFAND1 | 1023.949714 | 0.2529884198 | 0.09426229579 | 0.005155696986 | 0.02041496927 |
| BLZF1 | 1228.502178 | 0.2525180792 | 0.102978225 | 0.01000097622 | 0.03530216569 |
| RPL24 | 26128.40726 | 0.2524591755 | 0.07751763507 | 0.000813959717 | 0.004212390071 |
| RHEB | 3616.135246 | 0.2523969155 | 0.1065506337 | 0.01239354038 | 0.04216805822 |
| IMPAD1 | 3111.791309 | 0.2522517567 | 0.09179580352 | 0.004246093886 | 0.01734909217 |
| MARCKSL1 | 3711.187641 | 0.2521505954 | 0.07397785005 | 0.000474117882 | 0.00262324086 |
| C15orf57 | 725.4532968 | 0.2519392579 | 0.1030522556 | 0.01020526564 | 0.03591259753 |
| PPP1R8 | 2642.499349 | 0.2515657921 | 0.08504967347 | 0.00222180531 | 0.01002134482 |
| HNRNPK | 29560.59232 | 0.2511799708 | 0.08830593549 | 0.003174730696 | 0.01360246485 |
| MYEOV2 | 1901.586239 | 0.251179356 | 0.09818352147 | 0.007457795958 | 0.02756681331 |
| ZBED1 | 1917.294413 | 0.2509534199 | 0.07148639436 | 0.000324315921 | 0.001878156262 |
| CDK12 | 2078.020395 | 0.2506894998 | 0.09327078207 | 0.005153934051 | 0.02041418571 |
| PRPS2 | 1729.830173 | 0.2502755402 | 0.09826387568 | 0.007772160057 | 0.02852346342 |
| PWWP2A | 849.479586 | 0.2501232745 | 0.09886967932 | 0.008278048953 | 0.03009407203 |
| WLS | 7370.486508 | 0.249991486 | 0.06678401015 | 0.000136252135 | 0.000882408892 |
| COX7A2 | 4725.817378 | 0.249974358 | 0.1056379047 | 0.01248154516 | 0.0423899922 |
| HIAT1 | 2346.659917 | 0.2497133206 | 0.07775009059 | 0.000956957411 | 0.004849604797 |
| MED4 | 994.2350249 | 0.2495701611 | 0.09258166834 | 0.005032894177 | 0.01999860192 |
| SYNM | 458.4746147 | 0.2491857439 | 0.1079051499 | 0.01474607533 | 0.04867087467 |
| IL17RA | 887.4844132 | 0.2489230313 | 0.09233762171 | 0.005039719784 | 0.02002038097 |
| TLK2 | 1792.848912 | 0.248763362 | 0.08331922473 | 0.002048439661 | 0.009367081729 |
| ANP32B | 1525.376043 | 0.2485242768 | 0.1022099795 | 0.01069883249 | 0.03738439732 |
| EPS8 | 6624.906183 | 0.2481118694 | 0.08760139306 | 0.003339791273 | 0.0142277975 |
| ATP6V1E1 | 5457.944633 | 0.2474855432 | 0.06253621266 | 5.57E-05 | 0.000397342273 |

| | baseMean | log2FoldChange | lfcSE | pvalue | padj |
|---|---|---|---|---|---|
| INTS12 | 932.6326132 | 0.2474174698 | 0.08818802015 | 0.003626678981 | 0.0152363497 |
| B4GALT4 | 1087.8966 | 0.2472844562 | 0.08456476739 | 0.002502914397 | 0.01110095098 |
| ADPRM | 392.0274742 | 0.2470499558 | 0.09718222488 | 0.007869001059 | 0.02882203119 |
| SUPT5H | 3261.733699 | 0.2467002733 | 0.07246832392 | 0.000490020786 | 0.002702192403 |
| SLC35C1 | 755.7312847 | 0.2463863608 | 0.07487050869 | 0.000730651444 | 0.003827821731 |
| FAF2 | 3234.474361 | 0.2463700617 | 0.08538693012 | 0.002843485416 | 0.01239726335 |
| RTF1 | 2228.087156 | 0.2461243304 | 0.06336116 | 7.59E-05 | 0.000524947452 |
| RNF216P1 | 751.1417673 | 0.2456210107 | 0.09648842268 | 0.007863262861 | 0.02880810057 |
| ALG13 | 679.5551625 | 0.2455661905 | 0.08290038932 | 0.002229192077 | 0.01004216898 |
| ECD | 1695.683215 | 0.2455619536 | 0.09253484136 | 0.005773300998 | 0.02236759786 |
| PEX5 | 1198.214942 | 0.2450694652 | 0.09943085959 | 0.009851598146 | 0.03484096076 |
| DNAJB12 | 1858.273816 | 0.2448383157 | 0.08655918476 | 0.003401999878 | 0.01445146253 |
| SENP3 | 3105.832698 | 0.2448068189 | 0.09435608266 | 0.006786269429 | 0.02554114396 |
| ZBTB8A | 418.051558 | 0.2447953759 | 0.08781697116 | 0.003857000419 | 0.01603654824 |
| FAU | 15024.70032 | 0.2444686577 | 0.1017255294 | 0.01099212693 | 0.03820167619 |
| SOD1 | 8620.057939 | 0.244420351 | 0.06974287075 | 0.000338285394 | 0.001946939016 |
| STAG2 | 4595.724296 | 0.2442682098 | 0.07868811575 | 0.001400804795 | 0.006767891031 |
| WHAMM | 411.9457618 | 0.2437190197 | 0.0940964659 | 0.007086027328 | 0.02644362462 |
| TROVE2 | 1984.115216 | 0.2431231005 | 0.08394955957 | 0.002773136852 | 0.01211894176 |
| TNIP2 | 1457.838227 | 0.2429856249 | 0.0917319246 | 0.005881905808 | 0.02272922283 |
| MYL12A | 17244.21459 | 0.2428518758 | 0.09798550677 | 0.009421506183 | 0.0335774808 |
| GNG10 | 2779.480076 | 0.2426662393 | 0.08438444944 | 0.002956992501 | 0.01280961925 |
| CXorf56 | 975.3281824 | 0.2425456406 | 0.08660940788 | 0.003734450043 | 0.015618603 |
| C6orf106 | 3837.675559 | 0.2424843833 | 0.06470064882 | 1.33E-04 | 0.000863811060 |
| RPS10 | 45665.337 | 0.2423285382 | 0.08690734431 | 0.003882682475 | 0.01612531642 |
| PSMA6 | 6545.149366 | 0.2422608114 | 0.09452913791 | 0.007539018671 | 0.02779803095 |
| LSM1 | 1809.178503 | 0.2422134881 | 0.08790197066 | 0.004303673098 | 0.01754103169 |
| RPL35A | 25034.13525 | 0.2421871423 | 0.0731858748 | 0.000660689309 | 0.003500712856 |
| SNX11 | 701.8216316 | 0.2417850483 | 0.08018561787 | 0.001891563705 | 0.008765481484 |
| SHFM1 | 2958.481801 | 0.2417718696 | 0.07627590459 | 0.001128640997 | 0.005620179197 |
| SMNDC1 | 498.6049318 | 0.2416730642 | 0.09989743206 | 0.01126671978 | 0.0390570875 |
| RPL9 | 53660.94918 | 0.2413357662 | 0.06736823876 | 0.000169354088 | 0.00106803776 |
| CXorf38 | 608.1442317 | 0.2412986471 | 0.07674807205 | 0.001231291636 | 0.006062401182 |
| POLR3C | 2006.827666 | 0.2411123792 | 0.09183677038 | 0.006333857527 | 0.02410654517 |
| ARL8B | 2317.712386 | 0.2409596871 | 0.08528204562 | 0.003475931972 | 0.01470259975 |
| UBE2J1 | 3388.092284 | 0.2403536763 | 0.1014440616 | 0.01283207019 | 0.04334373134 |
| MATR3 | 17153.45676 | 0.2398294687 | 0.07615390782 | 0.001289705093 | 0.006304142851 |
| PCGF3 | 1633.05451 | 0.23914807 | 0.07428191808 | 0.000957361404 | 0.004850001345 |
| SURF4 | 16357.82809 | 0.2390397438 | 0.08107588391 | 0.002370944703 | 0.01059135784 |
| UBE2Z | 5062.262696 | 0.2388939747 | 0.06884225346 | 0.000395405340 | 0.002233380163 |

| | baseMean | log2FoldChange | lfcSE | pvalue | padj |
|---|---|---|---|---|---|
| SH3BP4 | 7138.093716 | 0.2387561538 | 0.0714189126 | 0.000615620763 | 0.003297114223 |
| TMSB4X | 61588.38408 | 0.2385654675 | 0.07076885508 | 0.000488948576 | 0.002697278752 |
| MANBAL | 1400.083464 | 0.2383205811 | 0.07219636786 | 0.000719893432 | 0.003780773657 |
| ARHGAP12 | 1795.096632 | 0.2378819253 | 0.09000702131 | 0.006087129512 | 0.02336459688 |
| PPHLN1 | 2309.615072 | 0.2366808968 | 0.08850169776 | 0.005549626727 | 0.02164179998 |
| PGM3 | 5682.427063 | 0.2365705927 | 0.1006947337 | 0.01381848502 | 0.04606883643 |
| GPATCH8 | 825.4989502 | 0.2362955841 | 0.08535440984 | 0.0041914647 | 0.01716356378 |
| UQCRH | 6046.46744 | 0.2361852648 | 0.07969832662 | 0.002243997386 | 0.0101000233 |
| ZNF644 | 2153.546153 | 0.2360053831 | 0.09507925833 | 0.009695818579 | 0.03439624561 |
| GLUL | 8722.153942 | 0.2358812174 | 0.0565428802 | 2.31E-05 | 0.000180063635 |
| XRN2 | 7534.632907 | 0.2350487211 | 0.08523797493 | 0.00432545571 | 0.01760730616 |
| TGFBRAP1 | 864.0902197 | 0.2345805532 | 0.09531026422 | 0.0102515615 | 0.03602442745 |
| IREB2 | 1836.220904 | 0.2344688073 | 0.09343850548 | 0.009006966551 | 0.03236117881 |
| GSR | 4163.971994 | 0.2344395667 | 0.07791001905 | 0.001986282991 | 0.009119262242 |
| BSDC1 | 2841.714241 | 0.2342229229 | 0.08900292269 | 0.006331708826 | 0.02410452895 |
| RPS21 | 12628.12173 | 0.2339740286 | 0.086863111 | 0.005284739642 | 0.02079949472 |
| GALNT1 | 5592.320021 | 0.2338561961 | 0.08856714083 | 0.006184450182 | 0.02366340532 |
| PRPF18 | 1012.826105 | 0.233335041 | 0.09846948986 | 0.01294976438 | 0.043681251 |
| DDA1 | 1240.895596 | 0.2332492763 | 0.0863837806 | 0.005184917415 | 0.02049719333 |
| RPS20 | 35825.23793 | 0.2327600398 | 0.07663413964 | 0.001454532107 | 0.006992744119 |
| OTUD5 | 1705.22745 | 0.2326759557 | 0.08041479771 | 0.002869143505 | 0.0124835411 |
| RPL23A | 42439.03668 | 0.2326589926 | 0.07030061183 | 0.000712631531 | 0.003746599883 |
| RAB2A | 3180.906313 | 0.2325395615 | 0.06224529826 | 0.000152603066 | 0.000972648568 |
| ZNF830 | 643.0745212 | 0.2325239446 | 0.09423760131 | 0.01013719104 | 0.0356899119 |
| NRBP1 | 6691.899459 | 0.2324488701 | 0.06310476139 | 0.000175923126 | 0.001103855895 |
| PTRF | 32196.77871 | 0.2323933074 | 0.06472841162 | 0.000251535846 | 0.001502856025 |
| SS18 | 3708.78957 | 0.2320776034 | 0.08674995812 | 0.005621851771 | 0.0218833336 |
| SIAH1 | 661.6285419 | 0.2320639319 | 0.08488659638 | 0.004676983689 | 0.01882043517 |
| ZNF24 | 3059.716316 | 0.2315798949 | 0.09030266711 | 0.00782308762 | 0.02868915063 |
| WAC | 3210.486624 | 0.2311699408 | 0.07749263361 | 0.002161217807 | 0.009789586834 |
| RBM23 | 2101.081466 | 0.2311173096 | 0.06338288265 | 2.04E-04 | 0.001252253689 |
| RPL26 | 40453.89412 | 0.2300561421 | 0.06855037494 | 0.000662053029 | 0.003506690698 |
| AAMP | 4192.365048 | 0.2297397669 | 0.09699964704 | 0.0131369748 | 0.04421063557 |
| SLC39A7 | 6277.758652 | 0.2295901172 | 0.05818037935 | 6.14E-05 | 0.000433339220 |
| ANXA7 | 5527.128614 | 0.2295035637 | 0.08290834623 | 0.004254853549 | 0.01737534681 |
| DNAJC1 | 949.4721545 | 0.2291540004 | 0.09629093744 | 0.01296987937 | 0.04372928759 |
| SETD7 | 3861.593177 | 0.2288730155 | 0.09664521768 | 0.01352412129 | 0.04523623769 |
| SNRPD2 | 10157.50102 | 0.2286395926 | 0.09349557152 | 0.01042288587 | 0.03654022787 |
| RPL32 | 45171.10242 | 0.2264723012 | 0.06864044924 | 0.000594516687 | 0.00319673491 |
| RPL37 | 40826.75767 | 0.2263773535 | 0.08781458012 | 0.005593825282 | 0.02179133036 |

| | baseMean | log2FoldChange | lfcSE | pvalue | padj |
|---|---|---|---|---|---|
| MAP3K7 | 2488.870338 | 0.2261951145 | 0.08665055666 | 0.006887167077 | 0.02583693381 |
| SAP18 | 4357.752655 | 0.224880855 | 0.08344199789 | 0.0053765804 | 0.02107183616 |
| MORC3 | 1439.520804 | 0.2232330749 | 0.08722745011 | 0.008064990314 | 0.02941686447 |
| G3BP2 | 6878.405428 | 0.2231724023 | 0.0895649657 | 0.008905701841 | 0.0320592347 |
| PARP12 | 554.0854033 | 0.2231398729 | 0.09079847271 | 0.01064227557 | 0.03722171504 |
| NDUFB9 | 4177.982422 | 0.2231228799 | 0.09175713294 | 0.01126434501 | 0.0390570875 |
| CAMKK2 | 1312.224662 | 0.2230504045 | 0.08164071543 | 0.00484614245 | 0.01937546051 |
| CD2BP2 | 4352.759482 | 0.2228565246 | 0.06222267365 | 0.000252807500 | 0.001509242532 |
| NDUFA12 | 2637.559197 | 0.2223825495 | 0.09435909494 | 0.01426459742 | 0.04735464681 |
| RAP1A | 751.546841 | 0.2223603661 | 0.08474663809 | 0.006683127517 | 0.02523588273 |
| KAT6A | 1749.473231 | 0.2220402465 | 0.08206062773 | 0.005265948079 | 0.02073649853 |
| PLA2G15 | 1218.967828 | 0.2220079656 | 0.06744510048 | 0.000771834194 | 0.004019531067 |
| RPS5 | 39054.95563 | 0.2218361287 | 0.08910690244 | 0.008433147084 | 0.03057246821 |
| RPL27 | 28034.24883 | 0.2216451861 | 0.08339828935 | 0.006542314605 | 0.02479717408 |
| BTBD1 | 3666.566165 | 0.2216402657 | 0.07683025034 | 0.003029709445 | 0.01305625005 |
| GLUD1 | 10464.04591 | 0.2215726449 | 0.06237060673 | 0.000296155555 | 0.001738097097 |
| PSMB1 | 9092.293381 | 0.2212203403 | 0.08091055254 | 0.004801612391 | 0.0192232339 |
| MTFMT | 600.6723772 | 0.2211899643 | 0.09474817769 | 0.01494879262 | 0.04921994102 |
| LRIG2 | 596.1828206 | 0.2208356925 | 0.09083023918 | 0.01153567891 | 0.03977645283 |
| RNF11 | 2042.999437 | 0.2205391651 | 0.06706744881 | 0.000787353194 | 0.004097048039 |
| PIGY | 1244.266308 | 0.2202285095 | 0.08788893427 | 0.00942442691 | 0.0335774808 |
| PYURF | 1244.266308 | 0.2202285095 | 0.08788893427 | 0.00942442691 | 0.0335774808 |
| STK24 | 3380.587822 | 0.2195751626 | 0.06258357659 | 0.000296589903 | 0.001739923983 |
| TM2D2 | 2014.710397 | 0.2190887409 | 0.08262874583 | 0.006229900443 | 0.02377165241 |
| SLC25A26 | 656.2587658 | 0.2184148871 | 0.08091000483 | 0.005404535478 | 0.02115355645 |
| TES | 2497.735326 | 0.2180622368 | 0.06445254012 | 0.000563039804 | 0.003045077971 |
| RPL36AL | 7627.521911 | 0.2163314113 | 0.0815906667 | 0.006239529392 | 0.02379619701 |
| CSNK1A1 | 11229.53296 | 0.2160272773 | 0.06176733924 | 0.000363961582 | 0.002072937822 |
| PI4K2A | 1828.5895 | 0.2151676549 | 0.08425658774 | 0.008325015804 | 0.0302172502 |
| SLC35A2 | 934.6037856 | 0.2147955708 | 0.08197753006 | 0.006907495501 | 0.02589267385 |
| SH3GLB1 | 3212.940603 | 0.2145912184 | 0.08351978199 | 0.007986855221 | 0.02917234292 |
| ARF4 | 10756.06382 | 0.2140622644 | 0.09184574716 | 0.01459605126 | 0.04827201403 |
| UBXN2A | 622.4257733 | 0.2139937005 | 0.08463616523 | 0.008956663454 | 0.03220375807 |
| KLF10 | 2339.602106 | 0.2139177368 | 0.08241589299 | 0.007425080209 | 0.02748682726 |
| FKRP | 591.6559498 | 0.2137492464 | 0.09086026839 | 0.01454365752 | 0.04813081059 |
| FAM105B | 1118.483238 | 0.2133565138 | 0.08935688672 | 0.01325201694 | 0.04448912746 |
| CUL1 | 4228.729025 | 0.2132785736 | 0.08297228288 | 0.008048294087 | 0.02937035555 |
| SMARCE1 | 5636.966483 | 0.2132010594 | 0.0678414592 | 0.001328677073 | 0.00647550669 |
| NDUFA1 | 2862.135093 | 0.2122096249 | 0.08350415208 | 0.008686651919 | 0.03133888064 |
| VCP | 28154.5131 | 0.2120214193 | 0.0871789243 | 0.01177254502 | 0.04045267086 |

| | baseMean | log2FoldChange | lfcSE | pvalue | padj |
|---|---|---|---|---|---|
| RPL31 | 36534.3318 | 0.2119652181 | 0.07350546559 | 0.003035222836 | 0.01307244224 |
| SCNM1 | 1067.346718 | 0.2117896616 | 0.07788944707 | 0.005163426858 | 0.02043547647 |
| RPS23 | 45171.67335 | 0.2112844812 | 0.06546767242 | 0.001039900377 | 0.00522151676 |
| C14orf2 | 2865.70553 | 0.2111417023 | 0.07886116783 | 0.005866535214 | 0.02268160031 |
| PHAX | 2163.989613 | 0.2104504391 | 0.08920071678 | 0.01453668777 | 0.04811844023 |
| TSNAX | 1283.567708 | 0.210208995 | 0.08549383251 | 0.01099592508 | 0.03820167619 |
| GLRX5 | 1409.567744 | 0.2094612496 | 0.07148700086 | 0.002704335491 | 0.01185306185 |
| RPS14 | 49586.2768 | 0.2089494656 | 0.08745013398 | 0.01133712387 | 0.03925545053 |
| LARP7 | 1807.856119 | 0.2081459352 | 0.08066132294 | 0.00783849216 | 0.02873856433 |
| KDM5A | 4006.047251 | 0.2081156453 | 0.08389880145 | 0.01034400178 | 0.03631642491 |
| EPC2 | 732.9696168 | 0.2069826435 | 0.08677136768 | 0.01352386093 | 0.04523623769 |
| RNF114 | 2626.734287 | 0.2059788968 | 0.08503243666 | 0.01227936458 | 0.04190407043 |
| RBM15B | 2792.89757 | 0.2059020871 | 0.07813184169 | 0.006716384146 | 0.02534026227 |
| RPS18 | 72373.25371 | 0.2058955482 | 0.07668319043 | 0.005732677111 | 0.02225498797 |
| EEF1B2 | 23363.00075 | 0.2058932092 | 0.06645986042 | 0.001544331796 | 0.007372092371 |
| PPP3CA | 1786.724926 | 0.2054486274 | 0.07665435725 | 0.005906309157 | 0.02279985404 |
| RPS27A | 42406.04917 | 0.2054231773 | 0.07571459519 | 0.004069460828 | 0.01676541291 |
| CHPF2 | 2761.382109 | 0.2053338927 | 0.08319344088 | 0.01085103076 | 0.03780088839 |
| RPL41 | 62094.25009 | 0.2051256919 | 0.08287345491 | 0.01065095504 | 0.03723457843 |
| FAM32A | 3781.616129 | 0.2050919174 | 0.05566547995 | 0.000182557814 | 0.001140144003 |
| GAK | 1884.47294 | 0.2046302833 | 0.0737732006 | 0.004459730527 | 0.01808796726 |
| LMO4 | 1659.15732 | 0.2041511771 | 0.07214038228 | 0.003751142795 | 0.01567960839 |
| RAB5C | 5263.424428 | 0.20319455 | 0.07714481859 | 0.006791207064 | 0.02554177091 |
| AFTPH | 841.9869732 | 0.2031060257 | 0.08659957875 | 0.01521089715 | 0.04994506785 |
| MED28 | 1281.250019 | 0.2025914368 | 0.06724255692 | 0.002098701067 | 0.009558751968 |
| FAM118B | 1127.759139 | 0.2025709003 | 0.07981063836 | 0.008972566414 | 0.03225314856 |
| TMCO1 | 2932.742381 | 0.2020730446 | 0.06989642992 | 0.003116387749 | 0.01337943979 |
| MAX | 1029.790645 | 0.2009679847 | 0.07216805149 | 0.004356503797 | 0.01772724379 |
| RPS12 | 43727.86492 | 0.2006175235 | 0.08139624876 | 0.01080220999 | 0.03767705541 |
| RPL17 | 57667.28674 | 0.1998463489 | 0.06735348235 | 0.002102109049 | 0.009571346066 |
| ERLEC1 | 2500.93697 | 0.1993106441 | 0.07433301851 | 0.005984650579 | 0.02305445342 |
| ACP1 | 4279.089367 | 0.199196502 | 0.06001986196 | 0.000741195096 | 0.003877601473 |
| LRRC57 | 747.7536633 | 0.1986633336 | 0.08445815363 | 0.01506538486 | 0.04951622853 |
| IFT57 | 2101.22003 | 0.1975188749 | 0.06760437639 | 0.002853145479 | 0.01243209922 |
| PDCL | 1363.004983 | 0.1973797069 | 0.07686272092 | 0.008165751583 | 0.02973340066 |
| JAGN1 | 1267.893508 | 0.1970391283 | 0.07618377536 | 0.007907514807 | 0.02892751547 |
| ACO1 | 9426.301213 | 0.1965082346 | 0.07486748889 | 0.007133812881 | 0.02658041541 |
| RPL38 | 15807.13148 | 0.1951259608 | 0.07166727024 | 0.005368854371 | 0.02104709656 |
| C11orf58 | 5947.565954 | 0.1949452634 | 0.06407566913 | 0.00196685705 | 0.009046813289 |
| RPS3A | 66617.23608 | 0.194625871 | 0.06926832793 | 0.004070755873 | 0.01676610904 |

| | baseMean | log2FoldChange | lfcSE | pvalue | padj |
|---|---|---|---|---|---|
| RPL30 | 26483.08963 | 0.1940830222 | 0.0733107193 | 0.006153811054 | 0.02359092802 |
| HDDC2 | 1842.839043 | 0.1940070661 | 0.07638696092 | 0.009067484793 | 0.03251584323 |
| FUBP3 | 2312.351678 | 0.1937912129 | 0.06839708255 | 0.003798949843 | 0.01583498438 |
| ATG12 | 1824.344299 | 0.1937124596 | 0.06683285745 | 0.003098797222 | 0.01331222588 |
| MFF | 2714.043915 | 0.1936776852 | 0.06525779247 | 0.002485233895 | 0.0110356837 |
| RPS17 | 44065.25728 | 0.1935691203 | 0.08027632027 | 0.01164155673 | 0.04010438181 |
| CHMP5 | 2574.777473 | 0.193198724 | 0.0682217273 | 0.003817849917 | 0.01590931078 |
| PEX19 | 2805.179494 | 0.1931979523 | 0.07393781035 | 0.007417132629 | 0.0274642347 |
| MLF2 | 4287.000314 | 0.192157885 | 0.08057780487 | 0.01414904757 | 0.04703397393 |
| PNISR | 2846.62043 | 0.1921176228 | 0.07602304895 | 0.009465649866 | 0.03369210157 |
| DENND5A | 5969.407273 | 0.1920617874 | 0.0735991776 | 0.007490620573 | 0.027667539 |
| MTCH1 | 10674.1381 | 0.1918791245 | 0.07911108724 | 0.01248025638 | 0.0423899922 |
| YY1AP1 | 1431.20397 | 0.1916099293 | 0.06870004512 | 0.004381450843 | 0.01779956933 |
| WBP5 | 3686.673195 | 0.1912212272 | 0.06890405796 | 0.004573098429 | 0.01850240829 |
| USP22 | 10726.06371 | 0.1908954369 | 0.06468154491 | 0.002628328496 | 0.01156417937 |
| KLC1 | 4145.193958 | 0.1890760561 | 0.0571987793 | 0.000813654257 | 0.004212273378 |
| HYAL2 | 2663.359019 | 0.18900465 | 0.07861532004 | 0.01341940947 | 0.0449497385 |
| RNF7 | 3883.129533 | 0.188705607 | 0.07596655015 | 0.01034441252 | 0.03631642491 |
| AMFR | 4931.257082 | 0.1880190373 | 0.05321319465 | 0.000346514519 | 0.001987386242 |
| RPL23 | 43263.08025 | 0.1878900821 | 0.07731379193 | 0.01064741269 | 0.03723093648 |
| MKRN2 | 2535.250156 | 0.1878563667 | 0.06480985946 | 0.003143193645 | 0.01347897759 |
| GRB2 | 3480.989445 | 0.186900325 | 0.06170563934 | 0.002062599765 | 0.009421410088 |
| RPL21 | 39794.13317 | 0.1862936925 | 0.06773468612 | 0.004865526544 | 0.01944251871 |
| RAF1 | 3998.398733 | 0.1860008847 | 0.06186456615 | 0.002225870422 | 0.01003360119 |
| RRAGA | 3537.993315 | 0.1850748705 | 0.05859935729 | 0.001337831778 | 0.006511597691 |
| SNW1 | 3874.177576 | 0.1847784465 | 0.06752942915 | 0.005215095327 | 0.02057433872 |
| EMD | 1444.2504 | 0.1844514251 | 0.07541666419 | 0.01209481799 | 0.04140716142 |
| TFIP11 | 1823.573757 | 0.1833781409 | 0.06846361788 | 0.006219463138 | 0.02375008635 |
| MCAM | 5426.537703 | 0.1833068162 | 0.06227149107 | 0.002687547372 | 0.01180032227 |
| BLMH | 2059.892985 | 0.1814269898 | 0.06176122119 | 0.002803933815 | 0.01223557168 |
| SPG20 | 3794.555943 | 0.1810746349 | 0.07173544535 | 0.009775979695 | 0.03462879305 |
| RPL11 | 44891.41602 | 0.177910653 | 0.05977013075 | 0.002520376383 | 0.01116177393 |
| CCNY | 3084.662557 | 0.1778169768 | 0.06037278512 | 0.0027567169 | 0.01205350406 |
| RPL39 | 21439.39517 | 0.1760146979 | 0.06155206515 | 0.003636914768 | 0.01527504203 |
| TPM4 | 51960.3191 | 0.1734264004 | 0.06039871245 | 0.003510640454 | 0.01483676575 |
| GHITM | 8618.475622 | 0.1730453687 | 0.07084441118 | 0.01243885286 | 0.04228357995 |
| RPS13 | 21562.70796 | 0.1723492693 | 0.06595246417 | 0.007658330564 | 0.02816820251 |
| PARD3 | 3191.906158 | 0.1715664593 | 0.06030384211 | 0.003828981549 | 0.0159423116 |
| DDX19B | 1340.435078 | 0.1706806988 | 0.07172698466 | 0.01486531987 | 0.04899927997 |
| PDXK | 4112.725358 | 0.170582442 | 0.06245229126 | 0.005450288959 | 0.02128931427 |

| | baseMean | log2FoldChange | lfcSE | pvalue | padj |
|---|---|---|---|---|---|
| TMED3 | 4309.849045 | 0.1701045148 | 0.0681077963 | 0.01074352252 | 0.03751414326 |
| ZDHHC7 | 2829.512428 | 0.1690163165 | 0.06951873939 | 0.01296838863 | 0.04372928759 |
| RPS24 | 42712.38129 | 0.16899661 | 0.05935245116 | 0.003946198235 | 0.01634806498 |
| RBM4 | 4373.160899 | 0.1688157906 | 0.05683550133 | 0.002582069431 | 0.01140107703 |
| RPL7 | 61940.86205 | 0.168290245 | 0.06367447367 | 0.007130010528 | 0.02657289781 |
| ARF3 | 4693.026444 | 0.1668179454 | 0.06671289633 | 0.01078856666 | 0.03764494234 |
| RNF185 | 2027.167268 | 0.1656361683 | 0.06125405356 | 0.005964259808 | 0.02298184893 |
| FBXO7 | 4121.306803 | 0.1646008073 | 0.06308810362 | 0.007932150452 | 0.02900338607 |
| CTTN | 12008.13095 | 0.1601612751 | 0.04726636164 | 0.000621417511 | 0.003317420341 |
| COLGALT1 | 9610.167454 | 0.1591542193 | 0.06532958844 | 0.01292513264 | 0.0436080444 |
| GORASP1 | 1740.838273 | 0.1564624449 | 0.06552429023 | 0.01492923445 | 0.04917729462 |
| FIP1L1 | 2277.841047 | 0.1528293377 | 0.06267318995 | 0.01308752872 | 0.0440660821 |
| BFAR | 3226.602478 | 0.1511778596 | 0.05897121439 | 0.009224118626 | 0.03297705001 |
| NACA | 29768.47489 | 0.1505851427 | 0.06238259673 | 0.01378677311 | 0.04598370627 |
| UBA52 | 23268.69632 | 0.1479857722 | 0.05994868718 | 0.01217404494 | 0.04162097706 |
| PPIB | 26357.21413 | 0.1456814143 | 0.06083430103 | 0.01503308134 | 0.04943187898 |
| NEK6 | 3312.419405 | 0.1420995628 | 0.05262657862 | 0.006266981967 | 0.02388254275 |
| RPL4 | 124452.637 | 0.1289695782 | 0.04852325749 | 0.006916077315 | 0.02591831743 |
| TXLNA | 5419.076647 | -0.1443514935 | 0.0561771332 | 0.008953963071 | 0.03220182516 |
| TIA1 | 2501.987543 | -0.1513878071 | 0.06025787871 | 0.01052714453 | 0.03686233652 |
| FAM104A | 1512.976379 | -0.1563313375 | 0.06538605998 | 0.0147794254 | 0.04874853007 |
| UBFD1 | 3773.431953 | -0.1595765356 | 0.0604327795 | 0.007240387291 | 0.02692360799 |
| PRKCA | 2878.379814 | -0.1636798373 | 0.06650572228 | 0.011971625 | 0.04105147964 |
| KANSL3 | 1690.411834 | -0.1727555393 | 0.06546267025 | 0.00710235122 | 0.02648970624 |
| VEZF1 | 2224.221192 | -0.1738790643 | 0.06079485014 | 0.003620196109 | 0.01522030373 |
| C16orf58 | 2915.48933 | -0.1740927572 | 0.06875479947 | 0.009643973917 | 0.03423679725 |
| LONP2 | 4010.144177 | -0.1752315203 | 0.0505030304 | 0.000446481014 | 0.002497241104 |
| EIF2AK1 | 2376.395423 | -0.1762988899 | 0.06135395479 | 0.003456764292 | 0.01464648934 |
| SNX5 | 3797.730099 | -0.1833903614 | 0.07180892893 | 0.008855681211 | 0.0318945906 |
| LYRM2 | 1688.750419 | -0.1851834837 | 0.07848510486 | 0.01517926821 | 0.04985751696 |
| NSF | 2928.988831 | -0.1868355815 | 0.0699984953 | 0.006171535886 | 0.02362766722 |
| TRIM13 | 1189.91144 | -0.1887787894 | 0.07792887519 | 0.01272275976 | 0.04305345571 |
| SMARCA2 | 1977.482305 | -0.1893837545 | 0.07781721677 | 0.01228340348 | 0.04190426083 |
| TBC1D14 | 2896.382666 | -0.190287813 | 0.07019693994 | 0.005941964726 | 0.02291966653 |
| DDX42 | 3606.655404 | -0.1904314046 | 0.06889938796 | 0.004600273395 | 0.01857198226 |
| FBXO38 | 2109.553193 | -0.1925610567 | 0.07300821124 | 0.006820169633 | 0.02562986009 |
| CPNE1 | 1921.830636 | -0.192866102 | 0.07701834877 | 0.01002743155 | 0.03537877449 |
| DCTN4 | 2803.929426 | -0.1947396296 | 0.07421702884 | 0.007072551285 | 0.02641164186 |
| STAMBP | 1620.172794 | -0.1960224435 | 0.07259434956 | 0.00572484921 | 0.02223194572 |
| KCTD2 | 1001.855893 | -0.1964326615 | 0.07647570791 | 0.0082788722 | 0.03009407203 |

| | baseMean | log2FoldChange | lfcSE | pvalue | padj |
|---|---|---|---|---|---|
| SP1 | 3035.400129 | -0.1971201486 | 0.07047875946 | 0.004199698123 | 0.01717613706 |
| LRCH3 | 1179.606272 | -0.1971532533 | 0.06976282019 | 0.003839750384 | 0.0159727668 |
| FAM114A1 | 6244.765003 | -0.1988767753 | 0.05614517377 | 0.000321282932 | 0.001862040321 |
| ZZEF1 | 1622.412194 | -0.1991913175 | 0.0781123174 | 0.008686461403 | 0.03133888064 |
| VPS11 | 1753.85343 | -0.199867041 | 0.07982131069 | 0.009870148375 | 0.0348817088 |
| FAM114A2 | 1139.976135 | -0.2002222563 | 0.08528031739 | 0.01509489151 | 0.04960225992 |
| SUMF2 | 3068.392136 | -0.2024942861 | 0.0857221936 | 0.01446900755 | 0.04793703904 |
| RCBTB1 | 1170.161102 | -0.2025806709 | 0.08376664241 | 0.01238284777 | 0.04214130754 |
| DCUN1D4 | 1165.334323 | -0.2036896373 | 0.08261982676 | 0.0108987215 | 0.03793152511 |
| CDC26 | 865.8594424 | -0.2042180531 | 0.08335371415 | 0.01135951189 | 0.03930554789 |
| SRSF4 | 2074.890785 | -0.2053748887 | 0.0814948578 | 0.00931263476 | 0.03325079591 |
| 8-Sep | 2889.679622 | -0.2054447648 | 0.05270041702 | 7.83E-05 | 0.000540045708 |
| TLDC1 | 1576.893532 | -0.2071026155 | 0.07475959713 | 0.004452777815 | 0.01806961267 |
| DYM | 1255.804796 | -0.2077686823 | 0.08435282431 | 0.01086588347 | 0.03784377521 |
| SPG7 | 1927.423028 | -0.2082034979 | 0.0759012679 | 0.004798220304 | 0.01921482036 |
| METTL14 | 883.958646 | -0.2087867888 | 0.07844772698 | 0.006154017358 | 0.02359092802 |
| PPFIA1 | 1519.283069 | -0.2094014333 | 0.06263253241 | 0.000659324092 | 0.003495967385 |
| CDK5RAP3 | 2018.906935 | -0.2101285306 | 0.08229200068 | 0.008453219996 | 0.03062286923 |
| BDNF | 3231.739345 | -0.2117923685 | 0.07383319611 | 0.003245836791 | 0.01388714482 |
| NXPE3 | 1874.443313 | -0.2121171723 | 0.08572637342 | 0.01038093675 | 0.03643606018 |
| ATF2 | 1857.717451 | -0.2145063176 | 0.08504982956 | 0.008584448926 | 0.03103371178 |
| FNBP1L | 1462.876531 | -0.2160478128 | 0.0777812846 | 0.00426458692 | 0.01740855007 |
| NNT | 3772.731764 | -0.2163604826 | 0.08102430314 | 0.005800351023 | 0.02246027442 |
| DNTTIP1 | 1479.156725 | -0.2182531087 | 0.08021652 | 0.005030242276 | 0.01999340023 |
| PYCR2 | 1513.990166 | -0.2185981335 | 0.08769267792 | 0.009731281557 | 0.03449739312 |
| NDRG3 | 1006.710125 | -0.2186748953 | 0.07805805887 | 0.003902251917 | 0.01619304035 |
| GTPBP10 | 668.4379327 | -0.2188328867 | 0.08933508298 | 0.01095885216 | 0.03811407375 |
| TCFL5 | 682.0330695 | -0.2189061986 | 0.08240924252 | 0.006103210102 | 0.02342028227 |
| WDR54 | 1222.723435 | -0.2200377421 | 0.09029270689 | 0.01129854721 | 0.03913097685 |
| EXD2 | 842.0608627 | -0.2201262404 | 0.09131651299 | 0.0121500367 | 0.04155798218 |
| HLCS | 940.3008803 | -0.2212082295 | 0.09410420774 | 0.01423877002 | 0.04730054593 |
| CGGBP1 | 2243.14123 | -0.2214953271 | 0.07984339781 | 0.00424893753 | 0.01735594812 |
| FDFT1 | 12606.67806 | -0.2215047248 | 0.09435199128 | 0.01427296733 | 0.0473666493 |
| HAUS4 | 1334.792307 | -0.2220924993 | 0.06452664233 | 0.000448194793 | 0.00250118901 |
| MAP4K2 | 642.4720717 | -0.2227908445 | 0.08883658517 | 0.00921551447 | 0.03296477897 |
| VPS52 | 1397.25319 | -0.2229168397 | 0.07895518725 | 0.003656879335 | 0.01534590654 |
| SLC35D2 | 650.6576367 | -0.2234444408 | 0.09278346132 | 0.01213016018 | 0.041509068 |
| PARG | 1177.371228 | -0.2240057462 | 0.08075968886 | 0.00420750565 | 0.01720086535 |
| GMPR2 | 2358.606838 | -0.2264293315 | 0.06081613443 | 1.52E-04 | 0.000966862136 |
| VPS39 | 2491.359354 | -0.226822523 | 0.09379232252 | 0.01167886839 | 0.04018642743 |

| | baseMean | log2FoldChange | lfcSE | pvalue | padj |
|---|---|---|---|---|---|
| PSPH | 1960.19353 | -0.2269023339 | 0.09524234176 | 0.0128769856 | 0.04345544845 |
| MECP2 | 793.9213867 | -0.2273330952 | 0.08947268905 | 0.008319255427 | 0.03020370496 |
| PPWD1 | 1197.20113 | -0.2279158309 | 0.09451188992 | 0.011879588 | 0.04077344071 |
| FBXO8 | 660.9206706 | -0.2280414478 | 0.09004957274 | 0.008470678305 | 0.03066346279 |
| RHBDD1 | 949.4568001 | -0.2281906678 | 0.08322140908 | 0.004605119562 | 0.01858147023 |
| MON1B | 629.8768245 | -0.2298059071 | 0.08710785289 | 0.006231727945 | 0.02377253328 |
| MMS19 | 4195.376929 | -0.2311478512 | 0.09890596873 | 0.01428387616 | 0.04737639389 |
| EXOC2 | 2459.01207 | -0.2315077584 | 0.07956175985 | 0.002712138951 | 0.01188026974 |
| TBC1D13 | 663.2918612 | -0.2323625155 | 0.08976415999 | 0.007142786968 | 0.02660719419 |
| MARCKS | 1682.196589 | -0.2325176911 | 0.06743173744 | 0.000426489743 | 0.002394421489 |
| IFT81 | 620.1545056 | -0.2325402624 | 0.09421856107 | 0.01004099174 | 0.03540983564 |
| FYCO1 | 1617.832849 | -0.2331170917 | 0.06881487484 | 0.000531498511 | 0.002898711115 |
| ATG14 | 733.6957322 | -0.2337460804 | 0.0936144751 | 0.009205122945 | 0.03293801383 |
| MTR | 2380.131497 | -0.2339854208 | 0.0972019923 | 0.01164926141 | 0.04012164081 |
| NFE2L1 | 6764.200483 | -0.2343843975 | 0.05958030005 | 6.34E-05 | 0.000446018797 |
| MOV10 | 2400.309553 | -0.234577805 | 0.09193722764 | 0.00788818183 | 0.02888517936 |
| ZBTB7B | 743.2706203 | -0.2364027963 | 0.1003612457 | 0.01349177892 | 0.045181983 |
| DCAF8 | 2044.350884 | -0.2369953553 | 0.09501726686 | 0.009224891753 | 0.03297705001 |
| FAM192A | 2302.392816 | -0.2370344877 | 0.09197612829 | 0.007268585847 | 0.02699837532 |
| ASCC1 | 1234.988466 | -0.2373196717 | 0.08988294109 | 0.006070168732 | 0.02332025325 |
| AHI1 | 821.290748 | -0.2374864049 | 0.09464373145 | 0.008766567332 | 0.0315932988 |
| ERMARD | 462.475386 | -0.237545186 | 0.1023765797 | 0.01471444302 | 0.04859201997 |
| SYNJ2BP | 1334.404733 | -0.2377633139 | 0.07906164147 | 0.001964869259 | 0.009040463039 |
| THAP11 | 965.5072986 | -0.2379441338 | 0.09031941451 | 0.006169695484 | 0.02362669652 |
| CHD8 | 3220.572088 | -0.2380908421 | 0.0844635457 | 0.003548051577 | 0.01497786785 |
| SUCLG2 | 4623.115651 | -0.23832115 | 0.07058630558 | 0.000544837111 | 0.002960613049 |
| KIAA1715 | 1222.024419 | -0.2384597864 | 0.09634792839 | 0.009708025718 | 0.03443134705 |
| NUB1 | 1545.165162 | -0.2403294236 | 0.08842407199 | 0.004788472 | 0.019196435 |
| ACBD5 | 995.2199556 | -0.2406535759 | 0.09763639291 | 0.009854773541 | 0.03484391433 |
| GBE1 | 5401.816107 | -0.2414592647 | 0.0854336827 | 0.003429931096 | 0.01455350359 |
| CXorf40A | 547.0631812 | -0.2418484961 | 0.1027678744 | 0.01334554831 | 0.04473252335 |
| SRGAP2B | 433.5086643 | -0.2426394102 | 0.09329683709 | 0.006679010627 | 0.02522727277 |
| RSBN1L | 555.6870561 | -0.2441661725 | 0.09203426528 | 0.005665766487 | 0.02203698987 |
| MAT2B | 2841.902583 | -0.2444712256 | 0.09804999103 | 0.009062277657 | 0.03250499929 |
| ZNF618 | 646.0169861 | -0.2448063635 | 0.1058597322 | 0.01475485779 | 0.04868906863 |
| SLC35B3 | 635.5394385 | -0.244874031 | 0.09149002578 | 0.005351948337 | 0.02100294117 |
| CETN3 | 474.2707287 | -0.2459905309 | 0.104084459 | 0.01283907645 | 0.04335699916 |
| HARS2 | 1550.190035 | -0.2461118934 | 0.08789573962 | 0.003663063367 | 0.01536319732 |
| PLEKHG4 | 405.6912252 | -0.2471926808 | 0.09497547319 | 0.006605624408 | 0.02500664679 |
| PARP4 | 5998.233227 | -0.2472974504 | 0.08512815678 | 0.002628177137 | 0.01156417937 |

| | baseMean | log2FoldChange | lfcSE | pvalue | padj |
|---|---|---|---|---|---|
| ACAD8 | 616.7093482 | -0.2474162235 | 0.084322344 | 0.002406782835 | 0.01072249839 |
| ST7L | 562.0871605 | -0.24742813 | 0.08854659715 | 0.003736615565 | 0.01562327131 |
| THUMPD3 | 1602.575621 | -0.2474813508 | 0.09393375399 | 0.005992910604 | 0.02307433307 |
| GBF1 | 2630.863354 | -0.2483886228 | 0.09431320181 | 0.005998541297 | 0.02309004172 |
| C9orf69 | 926.1088557 | -0.2488435808 | 0.09777510911 | 0.007722703206 | 0.02835628418 |
| SMARCC2 | 2403.496741 | -0.2493366663 | 0.09943297519 | 0.008564268617 | 0.03097240598 |
| GTF2A1 | 845.2641049 | -0.2493786528 | 0.09457269823 | 0.005949623653 | 0.02293732433 |
| ZNF561 | 765.7142601 | -0.2529237414 | 0.09212611962 | 0.004237364365 | 0.01731817679 |
| CNOT1 | 7896.721309 | -0.2538907843 | 0.07491340169 | 0.000495937168 | 0.002731155596 |
| EIF4G3 | 2863.699646 | -0.2546275034 | 0.08540158948 | 0.002027365917 | 0.009293550225 |
| RSPRY1 | 1722.407748 | -0.2546739097 | 0.09136985617 | 0.003714392958 | 0.01555219256 |
| EHMT1 | 1617.93225 | -0.255101879 | 0.0686057892 | 0.000143387773 | 0.000921165080 |
| ZNF512B | 1020.813189 | -0.2557594195 | 0.1015497355 | 0.008171731493 | 0.0297478998 |
| FBXO42 | 941.0328887 | -0.2557761752 | 0.09798183588 | 0.006263730226 | 0.02387626199 |
| HERC1 | 3257.79891 | -0.257900558 | 0.1078624384 | 0.01151943133 | 0.03972963009 |
| MTO1 | 1299.026109 | -0.2579704034 | 0.1015001619 | 0.007583619997 | 0.0279486431 |
| TRPV2 | 2494.883901 | -0.2579896439 | 0.09394670518 | 0.004181642889 | 0.01712805529 |
| COQ6 | 585.5857763 | -0.258924839 | 0.08818103417 | 0.002310624422 | 0.01034362207 |
| KBTBD4 | 1003.390748 | -0.2596970352 | 0.08670434283 | 0.001906699342 | 0.008824633666 |
| PATZ1 | 681.9400898 | -0.2597382327 | 0.08776996186 | 0.002133847069 | 0.009680567877 |
| ARHGEF17 | 1285.189101 | -0.2598370445 | 0.1067634107 | 0.01022301748 | 0.03594108791 |
| ZNF217 | 1859.868748 | -0.2599959478 | 0.1098630448 | 0.01222599568 | 0.04176023163 |
| PKNOX1 | 777.5717661 | -0.2600548612 | 0.08987960671 | 0.002648236428 | 0.0116414503 |
| FAM175A | 595.0551975 | -0.2602855803 | 0.1048270872 | 0.008911039569 | 0.03207069571 |
| STX16 | 760.9384485 | -0.2609450818 | 0.09265152254 | 0.003328879297 | 0.0141934948 |
| ELP6 | 1263.045893 | -0.2611044887 | 0.08384100367 | 0.001229757465 | 0.006056850445 |
| SMEK2 | 2566.045594 | -0.2611076428 | 0.07134148915 | 0.000177087997 | 0.001110641613 |
| TTPAL | 882.7568472 | -0.2611175948 | 0.1088679337 | 0.01124551318 | 0.03901082147 |
| ZZZ3 | 2980.583415 | -0.2612396998 | 0.0865516517 | 0.00176306637 | 0.0082340951 |
| POGZ | 796.8232288 | -0.2612983968 | 0.0925882989 | 0.003298045505 | 0.01407815353 |
| ZXDC | 681.0645758 | -0.2615516442 | 0.0856927817 | 0.001577636997 | 0.007507010945 |
| KMT2C | 1477.777528 | -0.2615881333 | 0.1073134234 | 0.01005509026 | 0.03543437607 |
| GATAD1 | 756.2060337 | -0.2616562383 | 0.09319509483 | 0.003407207746 | 0.01446945697 |
| NBR1 | 5739.605745 | -0.2621607119 | 0.1032352084 | 0.007563834527 | 0.02788262745 |
| TTC21B | 1027.955031 | -0.2624932269 | 0.1138142929 | 0.0141125679 | 0.0469441518 |
| FAM212B | 448.7282349 | -0.2625598166 | 0.1120298694 | 0.01275494947 | 0.04314139997 |
| KDM4C | 848.3506262 | -0.2628830291 | 0.09572733558 | 0.004134203606 | 0.01698045724 |
| DIS3L2 | 782.7160722 | -0.2634231069 | 0.09851053901 | 0.005106048967 | 0.02024067174 |
| KIAA0195 | 447.4082656 | -0.2658686986 | 0.1148453827 | 0.01373847238 | 0.04585342194 |
| DYNC1LI2 | 3653.481237 | -0.2662167938 | 0.06470679206 | 2.52E-05 | 0.000194302993 |

| | baseMean | log2FoldChange | lfcSE | pvalue | padj |
|---|---|---|---|---|---|
| OSBPL1A | 1255.653512 | -0.2676638992 | 0.1058545932 | 0.007658231069 | 0.02816820251 |
| VWA8 | 822.7592212 | -0.2677654681 | 0.1063876329 | 0.007903040892 | 0.02891825408 |
| XPNPEP3 | 1032.953727 | -0.2682927105 | 0.1008316716 | 0.0052259482 | 0.02061169344 |
| ZKSCAN5 | 968.8264067 | -0.2684826968 | 0.08903369005 | 0.001738722437 | 0.008140829676 |
| ARHGAP35 | 2933.214442 | -0.2685591187 | 0.08090803756 | 0.000617041949 | 0.003301163342 |
| AP3S2 | 1412.152348 | -0.2687102387 | 0.09396000884 | 0.00285961903 | 0.01245666113 |
| NCOA5 | 1230.665699 | -0.2687629283 | 0.08601704474 | 0.00121464481 | 0.005994314411 |
| CCDC174 | 613.0041568 | -0.2689269088 | 0.1015676576 | 0.005364431627 | 0.02104083838 |
| THADA | 1819.80825 | -0.2690580259 | 0.09022495037 | 0.001939641321 | 0.008951874929 |
| CLASP1 | 2657.389561 | -0.2690990394 | 0.06031731427 | 5.66E-06 | 4.99E-05 |
| NBAS | 5973.806075 | -0.2693848534 | 0.07705371007 | 0.000321174477 | 0.001862040321 |
| C15orf41 | 462.6270073 | -0.2700295327 | 0.1141860291 | 0.01187298402 | 0.04076017042 |
| ZRANB3 | 427.7711712 | -0.2701843073 | 0.1005541616 | 0.004827245276 | 0.01931546759 |
| UQCC1 | 1449.322421 | -0.2702033352 | 0.116365711 | 0.01328905394 | 0.0446034094 |
| ITSN2 | 744.0638653 | -0.2703720721 | 0.1055771685 | 0.006948385212 | 0.02601320277 |
| DNAJC13 | 5143.507379 | -0.2705681057 | 0.09751270173 | 0.003700914073 | 0.01550447654 |
| MRPS27 | 3853.575156 | -0.2707258325 | 0.07813681615 | 0.000361292070 | 0.002061815883 |
| UBE4B | 1852.652306 | -0.2707411893 | 0.1065814868 | 0.007390323654 | 0.02738739584 |
| TLN2 | 1235.488904 | -0.2709510963 | 0.08799517601 | 0.001401397545 | 0.006767891031 |
| CDC25B | 2096.912109 | -0.271217244 | 0.1043948423 | 0.006222961485 | 0.02375126212 |
| USO1 | 5290.116851 | -0.2713434334 | 0.1084219967 | 0.008145381825 | 0.02967374358 |
| METTL13 | 1248.167078 | -0.2714340683 | 0.1133598794 | 0.01089756011 | 0.03793152511 |
| R3HDM1 | 608.1601374 | -0.2714566515 | 0.1015577097 | 0.004956161947 | 0.01974639958 |
| SIN3B | 1080.847675 | -0.2716488735 | 0.08395916833 | 0.000810183944 | 0.004197226428 |
| HUWE1 | 5550.647383 | -0.2726050627 | 0.08542597421 | 0.000952676619 | 0.004831199655 |
| DROSHA | 1115.171895 | -0.272829446 | 0.1001414679 | 0.004275501346 | 0.01744053138 |
| COQ9 | 1384.933073 | -0.2729062689 | 0.1076026588 | 0.00738254097 | 0.0273701824 |
| MSH3 | 773.6731414 | -0.2730134084 | 0.094326583 | 0.002507256548 | 0.0111135882 |
| GLG1 | 5186.287187 | -0.2732088867 | 0.09622528682 | 0.002999097658 | 0.01295432696 |
| PGM1 | 2716.792377 | -0.2732141749 | 0.10793815 | 0.007488273088 | 0.02766573152 |
| ICK | 1882.257287 | -0.2732570699 | 0.08462053075 | 8.31E-04 | 0.004286801031 |
| TAF8 | 762.9024975 | -0.2734888208 | 0.08178601882 | 0.000556677474 | 0.003016146624 |
| TRAPPC11 | 2108.547069 | -0.2736189994 | 0.06920378373 | 5.23E-05 | 0.000374864578 |
| C6orf62 | 3266.276595 | -0.2738355231 | 0.08190546985 | 0.000555891925 | 0.003012986852 |
| TXNRD3 | 441.2146199 | -0.2739851904 | 0.09595308753 | 0.002863399801 | 0.0124694822 |
| ZCCHC11 | 820.015118 | -0.2743937593 | 0.1186320355 | 0.01356913089 | 0.0453492233 |
| POLR3GL | 1005.504925 | -0.2755099651 | 0.07534229938 | 0.000172195269 | 0.001084524264 |
| EXOG | 457.8632544 | -0.2760165946 | 0.09892873186 | 0.003474953827 | 0.01470259975 |
| BRCC3 | 744.1834327 | -0.2768912261 | 0.1165198986 | 0.01134729731 | 0.03928154142 |
| APAF1 | 1261.018145 | -0.2771318791 | 0.1020491077 | 0.004325846326 | 0.01760730616 |

| | baseMean | log2FoldChange | lfcSE | pvalue | padj |
|---|---|---|---|---|---|
| PPCS | 996.1435678 | -0.2775481907 | 0.09951134182 | 0.003458775944 | 0.01465084353 |
| EEF2K | 1007.967718 | -0.2787302483 | 0.09043464468 | 0.001358535625 | 0.006599424772 |
| WIBG | 732.9047581 | -0.2791455076 | 0.101180706 | 0.003783448132 | 0.01578656072 |
| CCND2 | 6069.441452 | -0.2793055364 | 0.121015762 | 0.01330838141 | 0.04463809209 |
| HSD17B7 | 643.1112108 | -0.2795116217 | 0.1096040163 | 0.00694508306 | 0.0260073797 |
| SCRN3 | 437.4259145 | -0.2801836236 | 0.1221507412 | 0.01383288527 | 0.04609620161 |
| SIKE1 | 1204.070834 | -0.2802471558 | 0.0940856308 | 0.001896850319 | 0.008787244678 |
| TRIM27 | 2348.994185 | -0.2808224352 | 0.06646360898 | 1.59E-05 | 0.000129143991 |
| TRIM2 | 1220.400735 | -0.2813039703 | 0.1060801647 | 0.005187448025 | 0.02049785394 |
| NRDE2 | 552.5499216 | -0.281507277 | 0.1096360236 | 0.006566748302 | 0.02487845177 |
| LOC93622 | 250.0201345 | -0.2818777097 | 0.1212708012 | 0.01272043685 | 0.04305345571 |
| IPO13 | 1115.150653 | -0.2821123719 | 0.08960654952 | 0.001070736819 | 0.005360524713 |
| UFSP2 | 1096.065748 | -0.2828144887 | 0.08328261566 | 0.000449108838 | 0.002505350881 |
| DNAJC16 | 920.054112 | -0.2828407339 | 0.1081942819 | 0.005793645022 | 0.02244057771 |
| PPIP5K1 | 610.3637843 | -0.2834623819 | 0.1233730364 | 0.01356687107 | 0.0453492233 |
| INTS6 | 2311.944785 | -0.2834751121 | 0.07776901735 | 0.000175801462 | 0.001103855895 |
| ZNF407 | 973.4118888 | -0.2837303798 | 0.1220554572 | 0.0126328298 | 0.04283539122 |
| VAT1L | 2487.405985 | -0.2838244214 | 0.07224503076 | 6.31E-05 | 0.000444194172 |
| CBY1 | 765.5769757 | -0.2847999694 | 0.09220713933 | 0.001301648343 | 0.006356261783 |
| WDR7 | 753.6358814 | -0.2857493434 | 0.09388520359 | 0.001492586957 | 0.007154902511 |
| IGHMBP2 | 413.0966284 | -0.2858154004 | 0.1130021767 | 0.0071797231 | 0.02673809336 |
| ATM | 3690.860306 | -0.2859930496 | 0.1228014009 | 0.01225372141 | 0.04184533442 |
| LIPG | 664.8309869 | -0.2860297261 | 0.1047474944 | 0.004032138235 | 0.01663005711 |
| PRR14L | 1940.08912 | -0.2866589196 | 0.1049896017 | 0.004045604816 | 0.0166809776 |
| VAPB | 1142.226542 | -0.2866822895 | 0.06662906744 | 1.11E-05 | 9.29E-05 |
| BRD7 | 2154.886228 | -0.2867883725 | 0.08196409568 | 0.000303669424 | 0.001777253954 |
| FKTN | 1217.489155 | -0.2879237583 | 0.1042975785 | 0.003659099081 | 0.01535089496 |
| CYP2U1 | 498.0513206 | -0.2881179909 | 0.1061086521 | 0.004192968725 | 0.01716500175 |
| EFR3A | 2143.633664 | -0.288385299 | 0.1003059087 | 0.002567508497 | 0.01134014655 |
| GALNT11 | 1147.726233 | -0.2883991092 | 0.08987418047 | 0.000857891833 | 0.004406054334 |
| RPUSD3 | 924.3306091 | -0.2886524067 | 0.109236656 | 0.005183390504 | 0.02049719333 |
| PIP4K2C | 313.8100167 | -0.2889225961 | 0.1193223235 | 0.009582886025 | 0.03403616174 |
| DIDO1 | 1750.776988 | -0.28905034 | 0.0901971222 | 0.000868637034 | 0.004453559507 |
| RAD17 | 806.6504497 | -0.2899975083 | 0.08142960641 | 0.000233836556 | 0.001412410748 |
| SBF2 | 2980.876678 | -0.290170876 | 0.112538598 | 0.006217217757 | 0.02374760266 |
| TMEM135 | 1022.470402 | -0.2904967581 | 0.09876852943 | 0.002074455967 | 0.009462798681 |
| GTF2H4 | 469.790796 | -0.290617644 | 0.1277798171 | 0.01407957443 | 0.04686581347 |
| SLC30A6 | 891.4022021 | -0.2911102619 | 0.1148538455 | 0.006979467091 | 0.02611702826 |
| EXOC3 | 1592.243361 | -0.2914180694 | 0.08794082253 | 0.000587265327 | 0.00316345639 |
| KIAA0319L | 1718.252062 | -0.2918439392 | 0.1211510775 | 0.009795927181 | 0.03466878055 |

| | baseMean | log2FoldChange | lfcSE | pvalue | padj |
|---|---|---|---|---|---|
| TAPT1 | 237.2845894 | -0.2919801391 | 0.1297855799 | 0.01495659354 | 0.04923473827 |
| SOS1 | 597.3838902 | -0.2920539088 | 0.1297662155 | 0.01500807431 | 0.04936215126 |
| PEX6 | 1162.343849 | -0.2923811747 | 0.1262790208 | 0.01277054245 | 0.04318433034 |
| IVD | 1563.40285 | -0.2925340958 | 0.1011861182 | 0.002388814172 | 0.01065200785 |
| FBXW8 | 673.4142079 | -0.2927038204 | 0.09291350211 | 0.00102613728 | 0.005163284206 |
| LPIN3 | 363.4289952 | -0.2931823153 | 0.1217949766 | 0.009866879058 | 0.03487843359 |
| TBC1D31 | 530.4717863 | -0.2933648058 | 0.1286073893 | 0.01369682662 | 0.0457349297 |
| GYS1 | 1751.71434 | -0.2941112598 | 0.1156159453 | 0.006747990024 | 0.02542838967 |
| MYCBP2 | 3935.877118 | -0.2947143628 | 0.1147001203 | 0.00619040659 | 0.02366340532 |
| LPAR1 | 9110.871916 | -0.2951554319 | 0.08898049237 | 0.000577504057 | 0.003117642463 |
| CRKL | 2452.407886 | -0.2953097362 | 0.06518152667 | 3.81E-06 | 3.47E-05 |
| ATG4C | 400.2987768 | -0.2954171237 | 0.1251757949 | 0.01109160887 | 0.03851281819 |
| RWDD2B | 860.5366786 | -0.2962321001 | 0.1223157512 | 0.009346612587 | 0.03336411287 |
| HPS4 | 842.3048648 | -0.296570227 | 0.09422742987 | 0.001030254597 | 0.005180500064 |
| MAP1A | 2911.611891 | -0.2966271477 | 0.0992738753 | 0.001741247831 | 0.008148841058 |
| ZBTB14 | 420.8967938 | -0.2966318397 | 0.1179177176 | 0.007253120224 | 0.02695076292 |
| SFXN5 | 474.3905043 | -0.2968630949 | 0.1103111427 | 0.004370552685 | 0.01776989056 |
| PCTP | 758.1460733 | -0.2971440846 | 0.1126040799 | 0.005098797374 | 0.02021730868 |
| DPH5 | 1437.42907 | -0.2973477194 | 0.08597761038 | 0.000339251828 | 0.001951746708 |
| POT1 | 794.4989382 | -0.2975861213 | 0.1255648653 | 0.01072363945 | 0.03745349936 |
| SOGA3 | 276.1672859 | -0.2982564553 | 0.1125762758 | 0.004938965743 | 0.01969369602 |
| ZFYVE21 | 1374.431236 | -0.2984253166 | 0.08865012569 | 0.000478441331 | 0.002644214172 |
| GDAP2 | 907.3331089 | -0.2985153847 | 0.1270331152 | 0.01112635865 | 0.03862169331 |
| ASH1L | 2116.565323 | -0.2985179528 | 0.1028073669 | 0.002276926056 | 0.01022659187 |
| PRMT2 | 3543.474411 | -0.2986462007 | 0.08369476899 | 0.000225252235 | 0.001364990042 |
| COQ5 | 1494.543719 | -0.2986858174 | 0.09672593157 | 0.001246509115 | 0.006127904088 |
| SLC2A1 | 3410.634603 | -0.2987159984 | 0.08630526805 | 0.000336721896 | 0.001940190525 |
| SNX24 | 391.4568171 | -0.2988191964 | 0.1031049106 | 0.00230861305 | 0.01033772623 |
| CEPT1 | 832.8776917 | -0.2991340427 | 0.1001294013 | 0.00173798958 | 0.008139958119 |
| LSM14B | 932.3389346 | -0.2992910828 | 0.09898216022 | 0.001530093911 | 0.007311157971 |
| DCAF5 | 1265.22644 | -0.2999481741 | 0.09142846651 | 0.000642977608 | 0.003421713868 |
| TBC1D9 | 850.2930029 | -0.300069362 | 0.09737719721 | 0.001274354439 | 0.00624345615 |
| TIAL1 | 1207.368739 | -0.3006488525 | 0.09367013876 | 0.000822200742 | 0.004249131155 |
| TUBGCP6 | 799.6663039 | -0.3013328562 | 0.1104709711 | 0.003870171019 | 0.01608404815 |
| FIG4 | 516.9436913 | -0.3013368928 | 0.1206473433 | 0.007524871904 | 0.02777338071 |
| ARAP3 | 527.4243556 | -0.301725216 | 0.1103237041 | 0.003790411214 | 0.01580381758 |
| MED23 | 1569.612292 | -0.3017295184 | 0.0894832526 | 0.000462578486 | 0.002564159002 |
| C1orf216 | 869.0019542 | -0.301918803 | 0.1035118916 | 0.002163261352 | 0.009793239871 |
| LPIN2 | 2665.29819 | -0.3027943405 | 0.08910349363 | 0.000419361400 | 0.002358848467 |
| WASH1 | 439.1106375 | -0.3033032837 | 0.1129500387 | 0.004322264651 | 0.01760716782 |

| | baseMean | log2FoldChange | lfcSE | pvalue | padj |
|---|---|---|---|---|---|
| VPRBP | 1162.54838 | -0.3034213834 | 0.1264072518 | 0.01002474095 | 0.03537766485 |
| OARD1 | 429.7526257 | -0.3038797245 | 0.1158457279 | 0.005202342779 | 0.02055125541 |
| FAM208A | 1898.976199 | -0.3045119436 | 0.1089687561 | 0.003132474305 | 0.01344075214 |
| ZBTB5 | 410.8722226 | -0.304519491 | 0.1239849467 | 0.00829938539 | 0.03014626716 |
| NUDCD3 | 3266.688285 | -0.304546616 | 0.1126028075 | 0.00413175059 | 0.01697506472 |
| PIGV | 481.4293385 | -0.3047940626 | 0.09389856252 | 0.000716544926 | 0.003764515669 |
| HINFP | 461.7873786 | -0.3056167858 | 0.1340412835 | 0.01313936882 | 0.04421063557 |
| CMTR2 | 705.9853282 | -0.3065243088 | 0.1158727425 | 0.004844728791 | 0.01937501127 |
| RPE | 2197.939365 | -0.3066264222 | 0.1157481933 | 0.004776372246 | 0.01915824525 |
| SUPT3H | 317.6334998 | -0.3068348393 | 0.1130822727 | 0.003964717524 | 0.01639741089 |
| STBD1 | 687.0992843 | -0.3069687423 | 0.1203494778 | 0.006325337021 | 0.02408643041 |
| GTF2I | 5118.453012 | -0.3070734625 | 0.1008051111 | 0.001397596621 | 0.006762696161 |
| PPM1K | 305.4398279 | -0.3072777881 | 0.1339347237 | 0.01266030766 | 0.04290901905 |
| THAP5 | 561.9025379 | -0.3074172104 | 0.106443061 | 0.002307257069 | 0.01033476248 |
| ARHGEF10L | 316.7613444 | -0.3075399471 | 0.1279599814 | 0.009478026109 | 0.03372809052 |
| TRAPPC2 | 438.7895263 | -0.3075807477 | 0.09808888895 | 0.001033973091 | 0.005195688611 |
| APC | 1990.175214 | -0.3077032669 | 0.1067709903 | 0.002398727238 | 0.01069300894 |
| DDX58 | 602.2286599 | -0.3078591825 | 0.1276186312 | 0.00915413657 | 0.03277132469 |
| ZNF41 | 387.9894274 | -0.3079322183 | 0.1145314136 | 0.004266268422 | 0.01740855007 |
| TTLL4 | 888.5245548 | -0.3080275794 | 0.116707115 | 0.004910517389 | 0.01960651472 |
| ANAPC2 | 626.6009272 | -0.3083623751 | 0.1023820953 | 0.001525881098 | 0.007293368756 |
| DENND2A | 1756.748904 | -0.3084297198 | 0.1190041702 | 0.005573837744 | 0.02171914948 |
| PRIM2 | 1596.434382 | -0.3085692653 | 0.1295960288 | 0.01003351293 | 0.03539184412 |
| GOPC | 1825.725346 | -0.3087615904 | 0.09440403373 | 0.000646037349 | 0.003432851571 |
| TTLL5 | 1009.26665 | -0.3097361127 | 0.07680748504 | 3.38E-05 | 0.000252783543 |
| DPY19L4 | 1112.833524 | -0.3102944141 | 0.1171608181 | 0.004751703393 | 0.01908500454 |
| SOGA2 | 1197.625735 | -0.3103450059 | 0.0853409914 | 0.000167299990 | 0.001056822047 |
| FAM120B | 1157.153647 | -0.3104339653 | 0.1043302341 | 0.001743743393 | 0.008156643224 |
| ZNF768 | 859.0972992 | -0.3104647058 | 0.105029378 | 0.001852983173 | 0.008608133059 |
| ZNF343 | 634.2386261 | -0.3111886328 | 0.117866613 | 0.004865362606 | 0.01944251871 |
| VPS36 | 1408.787457 | -0.311667522 | 0.08914752803 | 0.000283235888 | 0.001670138274 |
| SP4 | 262.1381815 | -0.3117033706 | 0.1391545094 | 0.01427457555 | 0.0473666493 |
| PREPL | 2410.043445 | -0.3117153102 | 0.09939352931 | 0.001015095637 | 0.005118103267 |
| FBXW2 | 2121.662079 | -0.3122857578 | 0.06605405362 | 1.39E-06 | 1.38E-05 |
| SKP2 | 2271.533339 | -0.3122883324 | 0.1075307521 | 0.002168178915 | 0.009810742861 |
| GLE1 | 1579.957926 | -0.3122955888 | 0.1101662712 | 0.002702079631 | 0.01184666185 |
| RGP1 | 1348.514092 | -0.3124766521 | 0.09851311246 | 0.000901904846 | 0.004601940117 |
| KIAA0408 | 5.98757897 | -0.3130151018 | 0.6135257511 | 0.000127833948 | 0.000836624026 |
| C6orf1 | 651.9388424 | -0.3130731166 | 0.1139978619 | 0.003535113791 | 0.01493171893 |
| HFE | 355.5175319 | -0.3132336823 | 0.1095412987 | 0.002491527945 | 0.0110603338 |

| | baseMean | log2FoldChange | lfcSE | pvalue | padj |
|---|---|---|---|---|---|
| RAB3GAP2 | 2448.924787 | -0.3132820803 | 0.0915836597 | 0.000369300833 | 0.002096271484 |
| ALKBH2 | 317.1675782 | -0.3133987415 | 0.1304531581 | 0.009307905386 | 0.03324188134 |
| INPP5F | 1194.129591 | -0.3134196424 | 0.08155507931 | 7.34E-05 | 0.000510731888 |
| DCAF10 | 773.2300324 | -0.3134340093 | 0.1043920324 | 0.001584091894 | 0.007530505814 |
| HIF1AN | 2276.895128 | -0.3135856103 | 0.06338214828 | 4.61E-07 | 5.08E-06 |
| NEK1 | 579.4306809 | -0.3137507302 | 0.1012566676 | 0.001138444856 | 0.005663316225 |
| TDRD3 | 522.2572477 | -0.3138125215 | 0.1128867165 | 0.003172064801 | 0.01359495476 |
| WDR81 | 640.7413892 | -0.3141147512 | 0.1312902525 | 0.00928316016 | 0.03316941964 |
| SCAPER | 502.5504106 | -0.3143639357 | 0.1020409518 | 0.001220362535 | 0.006014557359 |
| ARNT | 1518.383555 | -0.3143938077 | 0.08832494274 | 0.000223068209 | 0.001355064287 |
| MAU2 | 1075.613493 | -0.314866789 | 0.1018776197 | 0.001172705223 | 0.005818196621 |
| MTIF3 | 1087.088574 | -0.315509356 | 0.09388369337 | 0.000460853431 | 0.002555548136 |
| BTBD10 | 1580.652301 | -0.3155403893 | 0.1317073748 | 0.009379156246 | 0.03345621882 |
| USP24 | 4089.731646 | -0.3157831332 | 0.1053962209 | 0.001599724099 | 0.007592697515 |
| CCDC132 | 874.2482724 | -0.3159179368 | 0.115425623 | 0.003590871871 | 0.01511150121 |
| ZFYVE20 | 1288.823264 | -0.316042848 | 0.0834807657 | 9.16E-05 | 0.000622306846 |
| C2CD3 | 854.7578156 | -0.3164112387 | 0.1399267375 | 0.01329582357 | 0.04460602008 |
| IFT80 | 1212.600509 | -0.316948161 | 0.1392856871 | 0.01278324445 | 0.04320765643 |
| SNAPC3 | 1360.499751 | -0.3177853715 | 0.07596911216 | 1.72E-05 | 0.000138153482 |
| FBXL18 | 379.6162997 | -0.3178999759 | 0.1185163378 | 0.004198800748 | 0.01717613706 |
| MUT | 1339.142374 | -0.3187942082 | 0.08144372693 | 5.41E-05 | 0.000386867438 |
| SLC25A30 | 553.6526019 | -0.3191786342 | 0.1275031987 | 0.006944614018 | 0.0260073797 |
| EML2 | 702.9362405 | -0.3193740028 | 0.08987184129 | 0.000223780672 | 0.001357177362 |
| SAYSD1 | 306.6825762 | -0.319606563 | 0.1061562233 | 0.001507415587 | 0.007221335479 |
| OBFC1 | 865.8636217 | -0.3197773456 | 0.1076513477 | 0.001721410398 | 0.008069924249 |
| TUBG2 | 208.4251206 | -0.3201572932 | 0.1387441137 | 0.01166058201 | 0.04014205908 |
| YTHDC2 | 942.2131123 | -0.3206953941 | 0.1130303903 | 0.002599440175 | 0.0114573904 |
| PAN2 | 645.9315235 | -0.3209730012 | 0.09804774511 | 0.000618593391 | 0.003305899859 |
| HDAC6 | 1337.680282 | -0.321845022 | 0.06642546524 | 7.53E-07 | 7.92E-06 |
| MUS81 | 1176.333528 | -0.3222324642 | 0.1061745222 | 0.001378744753 | 0.006683136126 |
| ITPR1 | 325.7567887 | -0.3229664623 | 0.146417395 | 0.0149372873 | 0.04919293754 |
| CBX5 | 6876.014107 | -0.3231231411 | 0.1238391257 | 0.005450659278 | 0.02128931427 |
| LRSAM1 | 335.8607067 | -0.3231806615 | 0.1173025577 | 0.003329866885 | 0.01419364101 |
| PTPN18 | 655.7493075 | -0.3235133383 | 0.09096506838 | 0.000217541294 | 0.001327447676 |
| CDK19 | 531.8871312 | -0.3235388955 | 0.1415916853 | 0.01221833388 | 0.04174363772 |
| LCMT2 | 441.7171773 | -0.3236284455 | 0.1125132958 | 0.002260662993 | 0.01016581435 |
| INIP | 456.0511071 | -0.3244007745 | 0.1295464738 | 0.006791582447 | 0.02554177091 |
| PPCDC | 266.9863252 | -0.3245460469 | 0.1462414644 | 0.0142998689 | 0.04740831622 |
| NIT1 | 856.9028271 | -0.3246357859 | 0.08530221077 | 8.21E-05 | 0.000563948925 |
| CBR4 | 415.1024769 | -0.3246367966 | 0.1118262376 | 0.002074330842 | 0.009462798681 |

| | baseMean | log2FoldChange | lfcSE | pvalue | padj |
|---|---|---|---|---|---|
| UBXN2B | 318.651907 | -0.324767447 | 0.1462779014 | 0.01424749491 | 0.04731897204 |
| CTR9 | 2429.718814 | -0.3251992847 | 0.08423082498 | 6.57E-05 | 0.000459940919 |
| USP19 | 1890.234783 | -0.3253522095 | 0.08208837106 | 4.30E-05 | 0.000313994648 |
| ZNF160 | 927.6335667 | -0.325625553 | 0.1325648939 | 0.007722797526 | 0.02835628418 |
| TDRD7 | 308.2164039 | -0.3257895228 | 0.1271353865 | 0.005762288881 | 0.02233074418 |
| USP28 | 1350.470051 | -0.3258456863 | 0.08053344212 | 3.03E-05 | 0.000229667566 |
| NGLY1 | 1020.764438 | -0.3262236012 | 0.1029932279 | 0.000872872097 | 0.004472192932 |
| ZNF624 | 170.9298429 | -0.3264809253 | 0.1361857209 | 0.009019394381 | 0.03238238798 |
| CYP20A1 | 1124.865595 | -0.3265727464 | 0.08670062602 | 9.52E-05 | 0.000643958866 |
| PHC3 | 1338.999873 | -0.3268993986 | 0.1252789523 | 0.005005969502 | 0.019912872 |
| PEX1 | 786.1355335 | -0.3269008309 | 0.08202512887 | 3.82E-05 | 0.000281631737 |
| APPBP2 | 1811.894155 | -0.3273712913 | 0.09419852738 | 0.000290238491 | 0.001708047786 |
| TIGD6 | 251.2728633 | -0.3276407892 | 0.1245012813 | 0.004675308124 | 0.01881877877 |
| IMPACT | 1146.340092 | -0.3276602285 | 0.08919755824 | 0.000137144547 | 0.000886260925 |
| AP3M2 | 933.3280419 | -0.3281281622 | 0.10015951 | 0.000592100686 | 0.003186045216 |
| EFHC1 | 639.2924534 | -0.3284708687 | 0.09884241652 | 0.000505986363 | 0.002776863608 |
| PPP1R21 | 551.5972585 | -0.3285484322 | 0.09249052676 | 0.000219006652 | 0.001334748282 |
| GZF1 | 835.6422309 | -0.3287157346 | 0.09477734769 | 0.000298282061 | 0.001749161724 |
| NIPAL3 | 1740.560648 | -0.3289513626 | 0.08718639464 | 9.20E-05 | 0.000624345676 |
| LACE1 | 230.186073 | -0.3291376144 | 0.1494857121 | 0.01471344886 | 0.04859201997 |
| ZFP64 | 655.7958328 | -0.3292331069 | 0.1059552668 | 0.001055457079 | 0.005291144931 |
| UBE3B | 1424.520652 | -0.3292988475 | 0.08382630283 | 4.93E-05 | 0.000354634026 |
| TMEM234 | 453.0959209 | -0.3293815496 | 0.08947866959 | 0.000132858306 | 0.000863811060 |
| LOC100996497 | 128.2690317 | -0.3295177343 | 0.1442917114 | 0.0120298238 | 0.04122256538 |
| CHTF8 | 3909.433525 | -0.3297985308 | 0.09019734875 | 0.000145695851 | 0.000934222880 |
| ZSCAN12 | 550.6862382 | -0.3300354151 | 0.09044030201 | 0.000150285593 | 0.000960344289 |
| DHTKD1 | 1130.989617 | -0.3301008656 | 0.07256264137 | 3.12E-06 | 2.89E-05 |
| TYW5 | 224.2617216 | -0.3307389329 | 0.1292976715 | 0.005746398723 | 0.02229237379 |
| VPS13D | 2055.097767 | -0.3310153801 | 0.1303941465 | 0.006040545173 | 0.02321571427 |
| TCAIM | 1185.478519 | -0.331072627 | 0.09536918119 | 0.000292206661 | 0.001718950999 |
| PCNXL2 | 600.8053351 | -0.331282549 | 0.0861375992 | 6.86E-05 | 0.000479124900 |
| HMBOX1 | 582.2482201 | -0.3313989377 | 0.1062725884 | 0.001018692058 | 0.005132760087 |
| TMEM42 | 452.4456062 | -0.3318098971 | 0.1043848408 | 0.000829604479 | 0.004284419385 |
| JADE1 | 1977.066429 | -0.3318779094 | 0.13032739 | 0.005855833814 | 0.0226461064 |
| FAM179B | 806.5969667 | -0.3320085951 | 0.1139469804 | 0.001974049407 | 0.009071488156 |
| CDK10 | 841.5322613 | -0.3325362535 | 0.09539084937 | 0.000275470576 | 0.001630821018 |
| ZSCAN29 | 649.7717354 | -0.3325506579 | 0.09834229436 | 0.000403108958 | 0.00227344291 |
| BLOC1S5 | 529.6688821 | -0.33278392 | 0.1048704742 | 0.000841635487 | 0.004332401722 |
| TAOK2 | 1990.162829 | -0.3328742378 | 0.08084962791 | 2.19E-05 | 0.000171446829 |
| LOC100506548 | 1992.991776 | -0.3330488455 | 0.09391116212 | 0.000218111702 | 0.001330383094 |

| | baseMean | log2FoldChange | lfcSE | pvalue | padj |
|---|---|---|---|---|---|
| DLEU1 | 319.2875476 | -0.3333417541 | 0.1389148695 | 0.008647271307 | 0.03122707313 |
| CHD6 | 1735.109172 | -0.3339932891 | 0.1252574972 | 0.004137870574 | 0.01698614695 |
| RNF41 | 2320.277152 | -0.3342346415 | 0.0901410549 | 0.000117583558 | 0.000778089602 |
| DPH7 | 705.4988891 | -0.3342817831 | 0.1156901034 | 0.002113024524 | 0.009609292041 |
| TAF6L | 385.3125623 | -0.3346199084 | 0.1456036537 | 0.01129195334 | 0.03911723902 |
| RAB33B | 279.7378171 | -0.3347293336 | 0.1271866555 | 0.004593347563 | 0.01855898163 |
| ZC3H8 | 152.5923447 | -0.3350939683 | 0.1479352915 | 0.01228786371 | 0.04190426083 |
| NARG2 | 1533.266094 | -0.3352228666 | 0.08930664983 | 9.83E-05 | 0.000663532212 |
| MTA3 | 1089.846299 | -0.3353472438 | 0.1475185975 | 0.01197900627 | 0.0410595767 |
| GTF2H2C | 716.1849408 | -0.335526324 | 0.1327097713 | 0.006201768112 | 0.02370075088 |
| TSPYL5 | 1572.009972 | -0.3357110211 | 0.06521341152 | 1.50E-07 | 1.82E-06 |
| ARMCX1 | 1041.408295 | -0.3357476244 | 0.1147643232 | 0.001869332355 | 0.008673259404 |
| MTOR | 3626.208136 | -0.3365083668 | 0.06201309707 | 3.29E-08 | 4.50E-07 |
| HELZ | 1138.916315 | -0.3365581696 | 0.1148608194 | 0.00188902485 | 0.008756441776 |
| FBXL2 | 390.0069517 | -0.3367120143 | 0.1382293672 | 0.0078526028 | 0.028783211 |
| ENTPD5 | 255.3409907 | -0.3373492723 | 0.1136300904 | 0.001626910409 | 0.007697193862 |
| NOMO3 | 632.1745478 | -0.3378731559 | 0.1195669421 | 0.002560336018 | 0.01131182284 |
| TPCN2 | 267.903028 | -0.3379295058 | 0.1550159982 | 0.0150085606 | 0.04936215126 |
| RBM26 | 1220.220816 | -0.3384486915 | 0.1111363405 | 0.001263059387 | 0.006196273875 |
| SEMA5A | 1402.055596 | -0.3388907495 | 0.1146375172 | 0.001684646291 | 0.007930034341 |
| ARRDC1 | 241.1676258 | -0.3390838247 | 0.1471939383 | 0.01099686911 | 0.03820167619 |
| TNPO2 | 2718.958942 | -0.3393299084 | 0.1024614045 | 5.05E-04 | 0.00277523735 |
| DCP2 | 1024.947282 | -0.3396931796 | 0.1156074308 | 0.00177876769 | 0.008299521995 |
| DUSP18 | 190.0246837 | -0.3397189846 | 0.1385779909 | 0.007438927416 | 0.02751756281 |
| ACAD10 | 542.5764784 | -0.3397488633 | 0.1443890278 | 0.009652464516 | 0.03425877096 |
| EFS | 491.4380263 | -0.3400078811 | 0.1458466377 | 0.0102173377 | 0.03593809142 |
| FBXO4 | 241.1054153 | -0.3401456422 | 0.1422820821 | 0.008734575081 | 0.03148888339 |
| NPIPA1 | 445.4624011 | -0.3401732742 | 0.1415497327 | 0.008445344963 | 0.0306017866 |
| RUFY1 | 1129.222368 | -0.3408220227 | 0.09584133854 | 0.000205889080 | 0.001260996509 |
| NBPF9 | 1145.463199 | -0.3408243045 | 0.1194693646 | 0.002312937846 | 0.01035086612 |
| AASS | 1228.696669 | -0.3408367408 | 0.1101750819 | 0.001070451105 | 0.005360524713 |
| TNFAIP8L1 | 373.2313243 | -0.3412090087 | 0.1206016851 | 0.002454992978 | 0.01092420516 |
| TBCK | 870.0746709 | -0.3412574824 | 0.1134951516 | 0.001417764604 | 0.006835847537 |
| MPI | 1592.300529 | -0.3412659766 | 0.0691845687 | 4.56E-07 | 5.03E-06 |
| CDIP1 | 462.2662368 | -0.3413261789 | 0.1162565414 | 0.00178071032 | 0.008303475087 |
| PNMAL1 | 650.5462354 | -0.3414293269 | 0.1096646015 | 0.000990893206 | 0.00500794601 |
| AARS2 | 779.553698 | -0.3418299311 | 0.1268548782 | 0.003708756725 | 0.01553296171 |
| SNPH | 628.117728 | -0.3419037962 | 0.1158124604 | 0.001680761424 | 0.007919258492 |
| PTGR2 | 117.4781106 | -0.3423554374 | 0.1518030948 | 0.01231074362 | 0.04195345885 |
| TRMT2B | 392.458264 | -0.3423710211 | 0.1560981887 | 0.01425518184 | 0.04733394346 |

| | baseMean | log2FoldChange | lfcSE | pvalue | padj |
|---|---|---|---|---|---|
| GOLGA2P7 | 594.0583181 | -0.3423976866 | 0.1521835403 | 0.01237823092 | 0.04213522637 |
| TNFRSF19 | 295.9295671 | -0.3423979166 | 0.124851268 | 0.003209646664 | 0.0137402039 |
| AP4M1 | 755.0478159 | -0.3424253511 | 0.107775308 | 0.000816105750 | 0.004222028674 |
| ATF7 | 337.9593114 | -0.3425339211 | 0.1092980127 | 0.000934912469 | 0.004750823123 |
| CTCF | 1488.654767 | -0.3426839498 | 0.08692956942 | 4.43E-05 | 0.000322392047 |
| LARP6 | 1123.412631 | -0.3431079693 | 0.0812905043 | 1.34E-05 | 1.10E-04 |
| RGL1 | 754.9173177 | -0.3432214699 | 0.1233627162 | 0.002841401944 | 0.0123918083 |
| ZNF22 | 1453.528574 | -0.3433393951 | 0.08746540582 | 4.72E-05 | 0.000341267707 |
| ZMIZ2 | 254.545718 | -0.3438799314 | 0.1515930036 | 0.01183395401 | 0.04063554917 |
| HDHD2 | 878.2588885 | -0.3440156926 | 0.1139098118 | 0.001335409317 | 0.006501932412 |
| ZSCAN18 | 625.8166675 | -0.344570448 | 0.1576305228 | 0.01439796619 | 0.04771228991 |
| TUBE1 | 482.9605862 | -0.3445814566 | 0.1031508664 | 0.000447152639 | 0.002499119986 |
| PGPEP1 | 662.7026205 | -0.3450092506 | 0.1378410655 | 0.00636177287 | 0.02419423659 |
| RALGAPB | 1883.947002 | -0.3458097387 | 0.1061689089 | 0.000605080791 | 0.003245334258 |
| POLR1A | 1860.34933 | -0.3464799218 | 0.1588778758 | 0.01448993271 | 0.04798501069 |
| DDX60L | 404.9770964 | -0.3472280442 | 0.1240284267 | 0.002670804741 | 0.01173373024 |
| TRMT2A | 733.9261921 | -0.3476638558 | 0.1017388841 | 0.000337539565 | 0.001943397754 |
| TMEM260 | 524.6496995 | -0.3478709365 | 0.1431023287 | 0.007618018943 | 0.02804071299 |
| NME6 | 579.5570769 | -0.3487204317 | 0.1410882229 | 0.00679140818 | 0.02554177091 |
| FAM105A | 2063.116524 | -0.3491032371 | 0.109964698 | 0.000787785190 | 0.004097048039 |
| UBE2D4 | 312.5864465 | -0.349264448 | 0.136523076 | 0.005345239614 | 0.02098214411 |
| NEK9 | 2374.137633 | -0.3493496346 | 0.08546401571 | 2.35E-05 | 0.000183248303 |
| HEATR5A | 1841.146062 | -0.3494825973 | 0.09753788407 | 0.000180691440 | 0.001128961334 |
| TTBK2 | 364.250163 | -0.3495488105 | 0.1311369872 | 0.003952295169 | 0.01636421657 |
| ANKRD46 | 224.5216918 | -0.3495933943 | 0.1294749334 | 0.003562692617 | 0.01502263675 |
| C5orf51 | 754.6858904 | -0.3496988852 | 0.1097734215 | 0.000769817106 | 0.004010429287 |
| HERC2 | 1844.7978 | -0.3506612345 | 0.08556771777 | 2.23E-05 | 0.000174546926 |
| UNC119B | 1554.074393 | -0.350834543 | 0.07386912362 | 1.10E-06 | 1.11E-05 |
| WBP1L | 1147.992461 | -0.351104781 | 0.07996586899 | 6.12E-06 | 5.35E-05 |
| DZIP3 | 862.2300135 | -0.3514572405 | 0.146747613 | 0.008283866263 | 0.03009723884 |
| ZNF2 | 214.2919967 | -0.3516218821 | 0.1359949434 | 0.00489243258 | 0.01953954632 |
| ZNF654 | 488.0825825 | -0.3517774992 | 0.1237365115 | 0.002305726341 | 0.01033412387 |
| NFS1 | 760.3510688 | -0.3529232977 | 0.09673424711 | 0.000138982730 | 0.000896583134 |
| ARMCX6 | 1240.150248 | -0.3529964875 | 0.1034765445 | 0.000336125183 | 0.001937502073 |
| SETD4 | 443.8407613 | -0.353306735 | 0.1001089896 | 0.000218619257 | 0.001332932894 |
| PRSS12 | 1497.078698 | -0.3535472158 | 0.113501853 | 0.000947505679 | 0.004809891598 |
| C2orf43 | 1103.456629 | -0.3535903487 | 0.1031601282 | 0.000317102301 | 0.001842113216 |
| RBMS2 | 1848.192824 | -0.3536029871 | 0.06984241435 | 2.24E-07 | 2.61E-06 |
| FBXL7 | 232.8756801 | -0.3537263032 | 0.1602147535 | 0.01323821014 | 0.04445279901 |
| CC2D1B | 1457.113254 | -0.3544390628 | 0.1055020152 | 0.000429472131 | 0.002410256526 |

| | baseMean | log2FoldChange | lfcSE | pvalue | padj |
|---|---|---|---|---|---|
| NSL1 | 970.3114806 | -0.3544836887 | 0.08603075654 | 1.99E-05 | 0.000157671383 |
| PIP4K2B | 2748.757306 | -0.3547202248 | 0.06256173353 | 7.98E-09 | 1.22E-07 |
| PRKD1 | 706.2171243 | -0.3549252457 | 0.09223654773 | 6.30E-05 | 0.000443689268 |
| HEXIM1 | 436.901798 | -0.3562505283 | 0.1424038665 | 0.006081605085 | 0.02334941158 |
| MFAP1 | 1917.213579 | -0.3566473732 | 0.07715471152 | 2.02E-06 | 1.94E-05 |
| CTDSPL | 1528.945945 | -0.3567172633 | 0.1224812371 | 0.001756559746 | 0.00821143424 |
| ECHDC3 | 360.6467956 | -0.3568172171 | 0.1364445852 | 0.004402419844 | 0.01787521197 |
| SLC2A11 | 127.5954892 | -0.3570946023 | 0.164294518 | 0.01423434187 | 0.04729638833 |
| SMG9 | 675.6339839 | -0.3575829333 | 0.1101258058 | 0.000596641102 | 0.003203530248 |
| VAMP1 | 202.8665905 | -0.3576116264 | 0.148825798 | 0.007938756182 | 0.02901782863 |
| HMGCS1 | 9492.390624 | -0.358030333 | 0.1427395889 | 0.00589188857 | 0.02275359421 |
| C1orf63 | 787.1307106 | -0.3581068652 | 0.1019276511 | 0.000227989242 | 0.001378210245 |
| TTI2 | 599.0597406 | -0.3582967488 | 0.1519527221 | 0.008868412353 | 0.03193271863 |
| WDFY3 | 2338.935253 | -0.358390326 | 0.1095923698 | 0.000557670136 | 0.00302042585 |
| ATP2B4 | 4862.300028 | -0.359219997 | 0.1242952875 | 0.001923088551 | 0.00888667456 |
| ZNF197 | 803.9625363 | -0.3592344458 | 0.0910631989 | 4.16E-05 | 0.000304927696 |
| PLXNB1 | 816.9759908 | -0.3593017399 | 0.1120306676 | 6.83E-04 | 0.003601812302 |
| CLK1 | 1523.420968 | -0.3594005787 | 0.1034175908 | 0.000258586374 | 0.001538191185 |
| AR | 1080.290984 | -0.3596539968 | 0.1179100732 | 0.001156538728 | 0.005739901706 |
| ZFYVE1 | 822.2140349 | -0.3600177129 | 0.1318434463 | 0.003125698055 | 0.01341554291 |
| SESN1 | 555.7742306 | -0.3605571157 | 0.108421055 | 0.000447334878 | 0.002499200375 |
| NRBP2 | 483.5376997 | -0.3605657563 | 0.1407691298 | 0.005088639088 | 0.02018240473 |
| USP48 | 1529.475508 | -0.360592507 | 0.09781804473 | 0.000116793424 | 0.000774237445 |
| ZNF331 | 369.2673504 | -0.3606088223 | 0.1516081213 | 0.008353267691 | 0.0303124062 |
| AUH | 207.4980212 | -0.3612006896 | 0.1461446905 | 0.006535675923 | 0.02478595996 |
| PPP1R37 | 685.8545377 | -0.3614373327 | 0.1252588972 | 0.00194522555 | 0.008966706877 |
| ANKRD50 | 1411.574285 | -0.3624629166 | 0.09102118418 | 3.53E-05 | 0.000263461062 |
| INTS9 | 600.5332657 | -0.3626206173 | 0.1305577614 | 0.002695965597 | 0.01182769176 |
| GAS8 | 350.9163998 | -0.3626281514 | 0.1233691311 | 0.001623398695 | 0.00768449703 |
| C22orf39 | 474.7054229 | -0.3629439314 | 0.1092599583 | 0.000454061703 | 0.002526354523 |
| EFCAB7 | 191.7954286 | -0.3630569791 | 0.1637039421 | 0.01235086974 | 0.04207094475 |
| ZBTB47 | 794.0612107 | -0.3633370459 | 0.1412367164 | 0.004792490538 | 0.01920220442 |
| CYP1B1 | 167.1770082 | -0.3634460586 | 0.1554384772 | 0.009117709534 | 0.03266447961 |
| PAX8-AS1 | 977.4237176 | -0.3635171847 | 0.09290087662 | 4.69E-05 | 3.40E-04 |
| ORC5 | 714.5578085 | -0.3635710477 | 0.1640192412 | 0.01241192548 | 0.04221131988 |
| KIAA0907 | 974.0238355 | -0.3637832802 | 0.09384415387 | 5.43E-05 | 0.000387809846 |
| GIT2 | 1431.807316 | -0.363816028 | 0.08765674805 | 1.70E-05 | 0.000136715976 |
| ITFG2 | 762.088863 | -0.3642375032 | 0.1200922013 | 0.001196299364 | 0.005915543418 |
| F8A1 | 184.6796677 | -0.3642479372 | 0.1516491538 | 0.007704462676 | 0.02830292247 |
| RSRC2 | 2171.874153 | -0.3646360536 | 0.1004075971 | 0.000142379678 | 0.000915719667 |

| | baseMean | log2FoldChange | lfcSE | pvalue | padj |
|---|---|---|---|---|---|
| KRIT1 | 908.0772375 | -0.3648610529 | 0.1017994316 | 0.000170913417 | 0.001077362354 |
| TECPR2 | 1050.10157 | -0.3654918422 | 0.09444599081 | 5.55E-05 | 0.000396061849 |
| APOL6 | 1618.327155 | -0.3659273479 | 0.1682960633 | 0.0136777493 | 0.04568147361 |
| INTS2 | 890.2123087 | -0.3662477574 | 0.1025859659 | 0.000179470069 | 0.001122272096 |
| VPS13A | 1214.429759 | -0.3669238868 | 0.1674095113 | 0.01304121376 | 0.04393994833 |
| TBC1D2 | 1595.856459 | -0.3669464558 | 0.1197027914 | 0.001069139072 | 0.005357632418 |
| FAM109B | 800.4160227 | -0.3669986187 | 0.1107237326 | 0.000456993280 | 0.002537923521 |
| HK2 | 2439.726782 | -0.3670573946 | 0.1553126661 | 0.008461789987 | 0.03064645855 |
| ABCB6 | 857.7451273 | -0.367197212 | 0.1037454993 | 0.000200454088 | 0.001235331508 |
| C16orf70 | 526.8073079 | -0.3672222539 | 0.1046251361 | 0.000223910883 | 0.001357414144 |
| BIVM | 606.4789884 | -0.367612792 | 0.1178549387 | 0.000895864457 | 0.004575823641 |
| SPECC1L | 3354.696252 | -0.3676520194 | 0.07129315945 | 1.29E-07 | 1.59E-06 |
| AP3M1 | 1757.124692 | -0.3683594579 | 0.1158156493 | 0.000723567619 | 0.00379873 |
| MTERFD1 | 675.0274345 | -0.3685447083 | 0.1392372145 | 0.003877097529 | 0.01610661415 |
| ANKFY1 | 2092.884816 | -0.3686776342 | 0.0806329605 | 2.47E-06 | 2.33E-05 |
| C8orf48 | 122.8206608 | -0.3689018168 | 0.1669300317 | 0.01240391866 | 0.04219372743 |
| C15orf38 | 1013.383556 | -0.3689168274 | 0.08705261549 | 1.15E-05 | 9.57E-05 |
| APH1B | 618.8532572 | -0.3691211654 | 0.1158531256 | 0.000706598826 | 0.003717508807 |
| RPAP2 | 592.619187 | -0.3697274439 | 0.1037058076 | 0.000178606650 | 0.001119223239 |
| LOC220729 | 203.0962393 | -0.3699386319 | 0.1296239678 | 0.002080743949 | 0.009485669523 |
| CEP57 | 1568.654336 | -0.3704181509 | 0.06719397288 | 1.82E-08 | 2.62E-07 |
| MIA3 | 1990.219912 | -0.3711227461 | 0.1107680621 | 0.000406023498 | 0.002288146809 |
| LYST | 482.5935726 | -0.3715187389 | 0.1497503369 | 0.006105765887 | 0.02342405264 |
| CDK2AP2 | 637.0920769 | -0.3715398079 | 0.1429014893 | 0.004374898499 | 0.01778265459 |
| RAB4B | 402.8817301 | -0.3716235957 | 0.1743555105 | 0.0147702545 | 0.04872907582 |
| EXOC1 | 1766.211319 | -0.3717645563 | 0.08870539915 | 1.41E-05 | 0.000115266439 |
| FLRT2 | 599.5083248 | -0.3720172727 | 0.1189229239 | 0.000853271693 | 0.004385351136 |
| ABLIM3 | 3254.853773 | -0.3724592028 | 0.1238182363 | 0.001134437641 | 0.00564715548 |
| USP54 | 286.7067405 | -0.3734871614 | 0.138550745 | 0.003291492121 | 0.01405420883 |
| WIPF2 | 662.7805489 | -0.3745956138 | 0.08483799702 | 5.05E-06 | 4.49E-05 |
| ZNF555 | 345.3199981 | -0.3746328173 | 0.1329398572 | 0.002289206675 | 0.01027245274 |
| ZNF354C | 512.1828472 | -0.3747315535 | 0.1147926015 | 0.000530945963 | 0.002897820544 |
| GPT2 | 1215.685772 | -0.375058295 | 0.1040345019 | 0.000151459864 | 0.000966862136 |
| PDK3 | 276.8527644 | -0.375301478 | 0.1437832986 | 0.004198486537 | 0.01717613706 |
| NFIA | 355.6159147 | -0.3755595665 | 0.1195459719 | 0.000804060851 | 0.004169109929 |
| PRDM8 | 187.3803934 | -0.3757449992 | 0.1295618117 | 0.001771126525 | 0.00826655261 |
| ORC2 | 796.3793835 | -0.3767711238 | 0.1492363711 | 0.005297714145 | 0.02082853602 |
| DPF2 | 1387.660477 | -0.3770729316 | 0.0930431562 | 2.48E-05 | 0.000191793880 |
| CTBP1-AS2 | 159.3017263 | -0.377185722 | 0.1468337534 | 0.004678928818 | 0.01882317513 |
| OSGEPL1 | 339.0379238 | -0.3779298374 | 0.1679355764 | 0.01080283092 | 0.03767705541 |

| | baseMean | log2FoldChange | lfcSE | pvalue | padj |
|---|---|---|---|---|---|
| GBA2 | 1113.684234 | -0.3779742971 | 0.07008742659 | 3.49E-08 | 4.76E-07 |
| SYTL4 | 217.8886752 | -0.3783937337 | 0.1440389247 | 0.003960212195 | 0.01638788198 |
| PROSC | 1103.529077 | -0.3792065873 | 0.128934515 | 0.001522929775 | 0.007281599687 |
| SNX19 | 3249.521893 | -0.3793237775 | 0.07782553382 | 5.41E-07 | 5.91E-06 |
| MSMO1 | 5349.16336 | -0.3800953553 | 0.1570123489 | 0.006894725271 | 0.02585186318 |
| ZNF532 | 2312.751422 | -0.3801846808 | 0.1271285165 | 0.001304396181 | 0.006365504665 |
| CCDC127 | 553.1163167 | -0.3810560002 | 0.1126952405 | 0.000342629718 | 0.00196965787 |
| CCDC77 | 434.2183301 | -0.3811471 | 0.140192105 | 0.003000773815 | 0.01295780781 |
| ZNF347 | 333.6917556 | -0.3820138898 | 0.1343290999 | 0.002055196873 | 0.009392211861 |
| MTRF1 | 218.688252 | -0.3822615224 | 0.1332475317 | 0.001901866317 | 0.008807741086 |
| INPP5B | 1108.899158 | -0.3825274737 | 0.09395314679 | 2.25E-05 | 0.000175923601 |
| LYSMD1 | 292.9957766 | -0.3825451102 | 0.1288798461 | 0.001378910214 | 0.006683136126 |
| TRAFD1 | 2587.512026 | -0.3825670992 | 0.1123825939 | 0.000313081140 | 0.001821445884 |
| UTP14C | 1030.776733 | -0.3826036756 | 0.07938192043 | 7.09E-07 | 7.52E-06 |
| ZNF841 | 514.0442972 | -0.3832360945 | 0.1351296383 | 0.002095975914 | 0.009549261133 |
| TNRC6C | 180.7921905 | -0.3836924296 | 0.1281285587 | 0.001275883777 | 0.006246837735 |
| RNF20 | 2272.689554 | -0.3841886167 | 0.08091079537 | 9.98E-07 | 1.02E-05 |
| ULK3 | 562.1203755 | -0.3843450559 | 0.1017508482 | 7.45E-05 | 0.000516344553 |
| SMUG1 | 754.0990708 | -0.384443131 | 0.09310962042 | 1.75E-05 | 1.40E-04 |
| FAM149B1 | 767.3267055 | -0.3847526993 | 0.1079411642 | 0.000174722293 | 0.001099045304 |
| ZNF10 | 115.8962554 | -0.384953088 | 0.1705868443 | 0.01039986582 | 0.03648529739 |
| TOR2A | 175.5737719 | -0.3853862489 | 0.170940189 | 0.01041632468 | 0.03652582622 |
| ZNF260 | 679.248812 | -0.3856352728 | 0.1067799441 | 0.000143411959 | 0.000921165080 |
| R3HDM2 | 440.5014617 | -0.3859493663 | 0.1028350651 | 8.33E-05 | 0.000571276602 |
| C1orf35 | 297.3459435 | -0.3869229485 | 0.1428931718 | 0.003025909658 | 0.01304365052 |
| LOC150776 | 229.9071132 | -0.3873638106 | 0.1282254118 | 0.0011543411 | 0.005730905179 |
| A1BG-AS1 | 178.2875895 | -0.3881549324 | 0.1753000019 | 0.01140699249 | 0.03941951442 |
| RNF170 | 599.5835733 | -0.3881957018 | 0.1041471342 | 9.10E-05 | 0.000619178945 |
| ASTE1 | 326.3465623 | -0.3884007192 | 0.1071995856 | 0.000136128955 | 0.000881994786 |
| LYPLAL1 | 377.2999697 | -0.3888115979 | 0.1124785131 | 0.000253487864 | 0.001512697721 |
| TMTC4 | 627.3085322 | -0.3888326557 | 0.08593908294 | 2.93E-06 | 2.74E-05 |
| EDA2R | 327.1554666 | -0.3889821922 | 0.1278570829 | 0.001069439421 | 0.005357632418 |
| NMRK1 | 278.758322 | -0.389136498 | 0.1323573919 | 0.001482155999 | 0.007118651828 |
| ZNF700 | 429.2940827 | -0.3896408169 | 0.1160959551 | 0.000363603916 | 0.00207183265 |
| IKBKB | 1629.326039 | -0.3900747993 | 0.08064199643 | 6.30E-07 | 6.74E-06 |
| ANKS3 | 263.5005897 | -0.3903957894 | 0.1227861173 | 0.000669944776 | 0.003544707811 |
| RNASE4 | 182.7702355 | -0.3909904049 | 0.1749151109 | 0.01068896493 | 0.03735868518 |
| ZKSCAN4 | 241.8278673 | -0.3910768557 | 0.121479753 | 0.000584487036 | 0.003149629927 |
| NSUN4 | 609.0303893 | -0.3912128012 | 0.1090032814 | 0.000153411238 | 0.000976963185 |
| CYTH3 | 1370.550656 | -0.3913901759 | 0.1028043954 | 6.49E-05 | 0.000455290067 |

| | baseMean | log2FoldChange | lfcSE | pvalue | padj |
|---|---|---|---|---|---|
| IFIT3 | 350.1792804 | -0.3916291878 | 0.1783063986 | 0.01167488638 | 0.04018201185 |
| HEMK1 | 1094.908557 | -0.3916661408 | 0.09547895097 | 1.91E-05 | 0.000152523963 |
| LOC728407 | 131.6526597 | -0.3916668081 | 0.1803990565 | 0.01250965774 | 0.04247577971 |
| RALGAPA2 | 354.6969842 | -0.3917801937 | 0.1357584681 | 0.001741528124 | 0.008148841058 |
| IRAK1BP1 | 211.130906 | -0.3920969861 | 0.1340408709 | 0.001540744839 | 0.007359688774 |
| ZCCHC9 | 912.2960931 | -0.3925145545 | 0.1009804598 | 4.71E-05 | 0.000340715382 |
| VLDLR | 506.2422869 | -0.3926102786 | 0.1822695076 | 0.01256567582 | 0.04263681571 |
| ZFP30 | 397.4838 | -0.3926121188 | 0.1110588173 | 1.87E-04 | 0.001163985649 |
| ZNF250 | 378.3713527 | -0.3930204593 | 0.1159484958 | 0.000321106799 | 0.001862040321 |
| UTRN | 2526.785465 | -0.3930644665 | 0.1305880388 | 0.001153697213 | 0.005729619014 |
| POLDIP3 | 3137.455483 | -0.393078704 | 0.08991607295 | 5.78E-06 | 5.08E-05 |
| FN1 | 119745.7514 | -0.3936522567 | 0.1383115981 | 0.001665276822 | 0.007861225935 |
| DRP2 | 274.0830129 | -0.3938708982 | 0.1462199235 | 0.003078085757 | 0.01324172749 |
| FUT11 | 922.715123 | -0.3939008211 | 0.1104056774 | 1.64E-04 | 1.04E-03 |
| SH3D19 | 1869.290361 | -0.3942568568 | 0.1771678836 | 0.01077172672 | 0.03759499277 |
| HDDC3 | 416.9191727 | -0.3944418118 | 0.1586202513 | 0.005401216617 | 0.02114612522 |
| OIP5-AS1 | 903.2089781 | -0.3946933975 | 0.1024457958 | 5.35E-05 | 0.000382423539 |
| NBPF15 | 1391.825892 | -0.394810416 | 0.1296879753 | 0.001040165136 | 0.00522151676 |
| KMT2E-AS1 | 186.3565564 | -0.3953845764 | 0.1546384671 | 0.004480140633 | 0.01816085322 |
| ZNF25 | 373.4222594 | -0.3961995165 | 0.125008077 | 0.000684379505 | 0.003608260075 |
| MAGEF1 | 542.5313866 | -0.3963566838 | 0.1689128139 | 0.007793362003 | 0.02859422545 |
| EED | 672.4990345 | -0.3964347187 | 0.1458002431 | 0.002805764865 | 0.01223997453 |
| RAD51D | 375.8867754 | -0.3964421927 | 0.108704693 | 0.000119769843 | 0.000791501644 |
| CROT | 488.0630419 | -0.3965020103 | 0.1017393133 | 4.44E-05 | 0.000322926747 |
| MGARP | 165.4615632 | -0.3968763379 | 0.1875655399 | 0.01380726647 | 0.04604174478 |
| DTX3L | 1508.29468 | -0.3968862488 | 0.09633227941 | 1.74E-05 | 0.000139591512 |
| BHLHE40 | 1221.538067 | -0.3982441433 | 0.1288076921 | 0.000876037543 | 0.004486867212 |
| PMS1 | 698.302636 | -0.3983468278 | 0.1577419423 | 0.004609628906 | 0.01859462606 |
| ARMCX3 | 1793.453609 | -0.3988055011 | 0.08548492231 | 1.43E-06 | 1.41E-05 |
| UGGT1 | 4928.539448 | -0.3991914185 | 0.1287204976 | 0.000859631169 | 0.004413464991 |
| ZNF527 | 137.5958056 | -0.3995911908 | 0.1612842548 | 0.005565296063 | 0.02169154269 |
| C1orf50 | 295.9546701 | -0.3996611518 | 0.1092864335 | 0.000114594973 | 0.000761698463 |
| DNAJA3 | 2459.600334 | -0.4001339822 | 0.110666819 | 0.000134268750 | 0.000871839261 |
| FAM73B | 581.7911166 | -0.4006667296 | 0.1069789434 | 8.10E-05 | 0.000557111349 |
| ASB16-AS1 | 81.47577675 | -0.4008455065 | 0.1852146748 | 0.01220083392 | 0.04169341663 |
| ADCK1 | 102.5715387 | -0.4012750811 | 0.1628035601 | 0.005733772172 | 0.02225498797 |
| KLHDC10 | 1484.647597 | -0.4018872236 | 0.0725834219 | 1.44E-08 | 2.10E-07 |
| UBR4 | 6321.784452 | -0.4022970667 | 0.1106884304 | 0.000122678780 | 0.000806786379 |
| ACAD11 | 894.9995052 | -0.4024333615 | 0.07892778612 | 1.58E-07 | 1.91E-06 |
| HTR7P1 | 125.5924773 | -0.4025280554 | 0.1683709523 | 0.00688741959 | 0.02583693381 |

| | baseMean | log2FoldChange | lfcSE | pvalue | padj |
|---|---|---|---|---|---|
| MICU3 | 265.3959965 | -0.4030140944 | 0.1581690251 | 0.004529223164 | 0.01833984327 |
| NISCH | 2227.210752 | -0.4032275769 | 0.1317551689 | 0.000954747337 | 0.00484005213 |
| MUTYH | 291.9508406 | -0.4034389064 | 0.1230262981 | 0.000456629915 | 0.002536851794 |
| ASTN2 | 159.1919698 | -0.404119655 | 0.1442075437 | 0.002163339777 | 0.009793239871 |
| COA5 | 448.4551498 | -0.404282365 | 0.1410055887 | 0.001760620915 | 0.008225254094 |
| FAM219B | 1437.082511 | -0.4044113302 | 0.08537979465 | 9.89E-07 | 1.01E-05 |
| TNKS | 1646.658625 | -0.4047017969 | 0.1083148912 | 8.36E-05 | 0.000573140865 |
| ZNF205 | 253.0627998 | -0.4049225496 | 0.1849878837 | 0.01116489813 | 0.03874019304 |
| ANAPC4 | 772.602635 | -0.4049664775 | 0.1399666855 | 0.001607061673 | 0.007622663666 |
| ZNF767 | 298.9026971 | -0.4051238026 | 0.1655515353 | 0.005881772436 | 0.02272922283 |
| MAPK14 | 2330.932996 | -0.4051386056 | 0.06965228396 | 2.76E-09 | 4.50E-08 |
| LRCH4 | 437.8263275 | -0.4051436196 | 0.1172535931 | 0.000240463813 | 0.00144598777 |
| UBN2 | 357.4554938 | -0.4053279054 | 0.1476422523 | 0.002543309488 | 0.01124327048 |
| CCDC125 | 166.4963184 | -0.4058665552 | 0.1605792562 | 0.004727212163 | 0.0189968858 |
| PKNOX2 | 179.5846946 | -0.4062523174 | 0.173593191 | 0.007687082887 | 0.02825156225 |
| BTN3A3 | 475.7633163 | -0.4066206976 | 0.1506146822 | 0.002890892393 | 0.01256715236 |
| SUGP2 | 2617.662218 | -0.4071704722 | 0.09302943278 | 5.28E-06 | 4.67E-05 |
| NCOA2 | 1159.425883 | -0.4074076592 | 0.1200385112 | 0.000304886181 | 0.001782273401 |
| C4orf21 | 513.1949375 | -0.4075915917 | 0.1897313091 | 0.0122877198 | 0.04190426083 |
| TRIM24 | 521.9390474 | -0.4076474142 | 0.103529976 | 3.65E-05 | 0.000270453763 |
| ARSD | 522.9591789 | -0.4076559854 | 0.1442550183 | 0.001976819321 | 0.009081414031 |
| CLUAP1 | 1136.196474 | -0.4078996466 | 0.08283822398 | 3.83E-07 | 4.30E-06 |
| ACP6 | 213.263295 | -0.407989603 | 0.136682061 | 0.001198079717 | 0.005922380115 |
| TOM1L2 | 806.5887504 | -0.4082033171 | 0.1101249979 | 9.22E-05 | 0.000625915479 |
| SOX12 | 551.3166429 | -0.408992714 | 0.1453275779 | 0.001879445448 | 0.008717465194 |
| ZFP36L2 | 305.2173369 | -0.409439741 | 0.139513823 | 0.001401140999 | 0.006767891031 |
| CCDC113 | 198.0376468 | -0.4095627521 | 0.1572932811 | 0.003720857057 | 0.01557050048 |
| NUP43 | 1842.087375 | -0.4095832768 | 0.1593360357 | 0.00411518764 | 0.01693037545 |
| ZCCHC8 | 756.0065202 | -0.4096932948 | 0.09746920313 | 1.16E-05 | 9.64E-05 |
| FANCB | 286.5965813 | -0.4097908566 | 0.193136483 | 0.01298676185 | 0.04377629548 |
| TTC9C | 829.9652342 | -0.4098160206 | 0.1207975669 | 0.000294233446 | 0.001729507217 |
| CCDC97 | 291.5564075 | -0.4098669656 | 0.1189977873 | 0.000245882845 | 0.001475594393 |
| LINC00674 | 913.3103219 | -0.4100018075 | 0.1341023439 | 0.000942611027 | 0.004786676532 |
| C7orf25 | 478.3076373 | -0.4105415365 | 0.1008291503 | 2.05E-05 | 1.62E-04 |
| ITPR2 | 846.6297045 | -0.4105498332 | 0.1580116495 | 0.003787402296 | 0.01579569546 |
| BCL9 | 127.2983631 | -0.4110397998 | 0.1585385206 | 0.003870585299 | 0.01608404815 |
| EHMT2 | 1508.325861 | -0.4121729662 | 0.1170780582 | 0.000183290961 | 0.001144242818 |
| TMEM63A | 358.703347 | -0.4122486721 | 0.1336341196 | 0.000850636051 | 0.004374825618 |
| STX7 | 1352.42142 | -0.4126465082 | 0.08100665173 | 1.57E-07 | 1.90E-06 |
| DHRS13 | 155.8018865 | -0.4127977693 | 0.191420219 | 0.01175643615 | 0.04041597271 |

| | baseMean | log2FoldChange | lfcSE | pvalue | padj |
|---|---|---|---|---|---|
| ARL10 | 138.1900471 | -0.413410883 | 0.147608418 | 0.002110224704 | 0.00960242531 |
| ERV3-1 | 217.5014222 | -0.4137699342 | 0.1981979249 | 0.01378470306 | 0.04598370627 |
| CXXC1 | 957.0477454 | -0.413889563 | 0.1242053439 | 3.62E-04 | 0.002066639542 |
| ZMYM2 | 2274.940385 | -0.4146204414 | 0.1607319619 | 0.00395415789 | 0.01636737749 |
| LOXL1-AS1 | 233.4902729 | -0.4152111736 | 0.1427639244 | 0.001482724912 | 0.007119087784 |
| BTN3A1 | 638.1990948 | -0.4153953954 | 0.09480147117 | 5.08E-06 | 4.51E-05 |
| CLSTN3 | 254.6578593 | -0.4155669426 | 0.1834885634 | 0.009010682278 | 0.03236671856 |
| DPF3 | 301.0730703 | -0.4155769641 | 0.2004366327 | 0.01408977376 | 0.04688928062 |
| DCLRE1C | 445.0856384 | -0.4161526172 | 0.1260888075 | 0.000401303964 | 0.002264120773 |
| CYB5RL | 122.865516 | -0.4163311704 | 0.185158564 | 0.009359732838 | 0.03340293917 |
| VPS13C | 2519.174092 | -0.4163618152 | 0.1528390598 | 0.00258812979 | 0.01142106237 |
| IPP | 418.0245995 | -0.4163760774 | 0.1021967513 | 1.99E-05 | 0.000157671383 |
| AARS | 9094.882733 | -0.4170822098 | 0.06401989564 | 3.17E-11 | 6.77E-10 |
| CLEC2D | 80.86768465 | -0.4175324879 | 0.1903273555 | 0.01061035991 | 0.0371362597 |
| ZNF75D | 309.7701653 | -0.4176968764 | 0.1379709949 | 0.001012626941 | 0.005107385678 |
| USP20 | 417.8545235 | -0.4184257349 | 0.1460496752 | 0.001683025244 | 0.007924909191 |
| BTG2 | 485.7038399 | -0.4185504556 | 0.1027375691 | 1.98E-05 | 0.000156919014 |
| ADPRHL2 | 1234.775188 | -0.4191634961 | 0.1166015235 | 1.38E-04 | 8.92E-04 |
| ZNF558 | 435.7394762 | -0.4192628354 | 0.1512099902 | 0.002215470847 | 0.009998831597 |
| SEC22A | 625.436572 | -0.4197814367 | 0.1151640729 | 0.000119912290 | 0.000791826441 |
| MTUS1 | 215.1724265 | -0.4198898024 | 0.1710814442 | 0.005427530537 | 0.02122681959 |
| ZNF462 | 909.9182819 | -0.4204954818 | 0.1288185341 | 0.000451733347 | 0.00251810476 |
| ZNF585A | 288.0462959 | -0.420914431 | 0.1305808734 | 0.000521311389 | 0.002849414566 |
| LOC202181 | 218.697724 | -0.4209979416 | 0.1683503957 | 0.004770535109 | 0.01913998848 |
| BPTF | 1615.074266 | -0.4213435552 | 0.08082536223 | 8.11E-08 | 1.03E-06 |
| NDST2 | 434.8081 | -0.4223670495 | 0.1318169098 | 0.000553623710 | 0.003002879205 |
| IFT172 | 908.5000644 | -0.4228298584 | 0.09530057414 | 3.87E-06 | 3.52E-05 |
| DYNC2H1 | 2765.377189 | -0.4229787246 | 0.1355069711 | 0.000727070546 | 0.003814430361 |
| PLEKHO2 | 1574.110399 | -0.4232809391 | 0.1255682742 | 0.000304685217 | 0.001781798195 |
| TENM3 | 703.2407042 | -0.4240437483 | 0.1283833879 | 0.000389326057 | 0.002202384372 |
| KIAA1467 | 213.1398874 | -0.4243597785 | 0.1427627056 | 0.001176088366 | 0.005833037868 |
| NFX1 | 1742.081625 | -0.4250231435 | 0.08859361529 | 6.74E-07 | 7.18E-06 |
| ZFYVE26 | 1212.134156 | -0.4250478602 | 0.09382736129 | 2.49E-06 | 2.35E-05 |
| ZNF844 | 343.6862258 | -0.4253152431 | 0.1670035256 | 0.004141671562 | 0.01699238024 |
| KIAA0586 | 1037.112763 | -0.4254325095 | 0.1699457722 | 0.004662228622 | 0.01878136416 |
| GPR155 | 163.9321293 | -0.4264802799 | 0.1998919013 | 0.01181449072 | 0.0405874371 |
| MEN1 | 586.6830887 | -0.4265278018 | 0.1211816854 | 0.000175397307 | 0.001102124659 |
| ZFP37 | 87.08505954 | -0.4265585778 | 0.1951957207 | 0.01056656532 | 0.03699167435 |
| CPS1 | 305.5007621 | -0.4267788777 | 0.13555478 | 0.000657380038 | 0.003486901103 |
| SPG11 | 2778.323488 | -0.4267926171 | 0.08982842516 | 8.54E-07 | 8.88E-06 |

| | baseMean | log2FoldChange | lfcSE | pvalue | padj |
|---|---|---|---|---|---|
| G2E3 | 1007.584511 | -0.4268569814 | 0.1676136854 | 0.004130469492 | 0.01697506472 |
| ELMO2 | 1453.045824 | -0.427185835 | 0.07022243473 | 4.88E-10 | 8.70E-09 |
| FIGNL1 | 805.921141 | -0.4272563393 | 0.1882030227 | 0.008403424992 | 0.03047956022 |
| DIP2A | 869.5356713 | -0.4274399511 | 0.08954284733 | 7.67E-07 | 8.06E-06 |
| MORC2 | 1098.088531 | -0.4274940324 | 0.104673469 | 1.82E-05 | 0.000145755074 |
| NREP | 1199.940827 | -0.4279347894 | 0.1645373382 | 0.003557543915 | 0.01500517602 |
| SIRT5 | 399.8947043 | -0.4279932616 | 0.1278553451 | 0.000326208689 | 0.001888383041 |
| MDM4 | 861.6152265 | -0.4280766702 | 0.1218201484 | 1.79E-04 | 1.12E-03 |
| PRDM5 | 406.7714595 | -0.4289363148 | 0.1185966202 | 1.22E-04 | 0.000800771960 |
| NR2C2 | 851.2705395 | -0.4289377693 | 0.08807403364 | 4.69E-07 | 5.16E-06 |
| CBFA2T2 | 311.0929951 | -0.4296533195 | 0.1245593917 | 0.000225788466 | 0.001367242321 |
| ERCC4 | 678.5999082 | -0.4296694873 | 0.09114944102 | 1.02E-06 | 1.04E-05 |
| COL12A1 | 22673.69721 | -0.4297895436 | 0.1618426082 | 0.002586446934 | 0.01141701998 |
| HOOK2 | 112.2647441 | -0.4302286378 | 0.205172003 | 0.0126786858 | 0.04295175266 |
| TIAM2 | 225.7681043 | -0.4307058625 | 0.1629079681 | 0.003098943844 | 0.01331222588 |
| SLC46A1 | 100.8614705 | -0.4307083852 | 0.2109983228 | 0.01427806921 | 0.04736768548 |
| USF1 | 1402.735204 | -0.4316763265 | 0.09197819334 | 1.11E-06 | 1.12E-05 |
| RAB40C | 462.899256 | -0.4318157811 | 0.1465764562 | 0.001246653333 | 0.006127904088 |
| LOC730102 | 133.2686343 | -0.4319919174 | 0.1855702684 | 0.007243971856 | 0.02693021148 |
| HEIH | 456.7786015 | -0.4325171708 | 0.1736028821 | 0.00483417633 | 0.01933800413 |
| SREBF1 | 814.4819461 | -0.4326439227 | 0.1291840066 | 0.000320470732 | 0.001860229528 |
| ZNF891 | 73.43366758 | -0.4338670381 | 0.1955247926 | 0.009393270056 | 0.03349853841 |
| PDK1 | 1173.002535 | -0.4340859137 | 0.1329040293 | 0.000447595384 | 0.002499717808 |
| MPHOSPH9 | 1344.604097 | -0.4342218672 | 0.1022865367 | 8.89E-06 | 7.58E-05 |
| FIBIN | 3936.829739 | -0.434332252 | 0.1107190477 | 3.52E-05 | 0.000262365882 |
| RASA1 | 1760.409843 | -0.4348472642 | 0.1382016082 | 0.000640072490 | 0.003408454689 |
| TBC1D24 | 409.9597016 | -0.4348757213 | 0.1210642899 | 0.000130534958 | 0.000850934763 |
| ZNF678 | 279.1400201 | -0.435783715 | 0.1433969121 | 0.000909274043 | 0.004634776184 |
| TMEM216 | 279.4385735 | -0.4366864886 | 0.1647542417 | 0.00298711927 | 0.01291007803 |
| BNIP3 | 6687.110135 | -0.4369589893 | 0.1416884618 | 0.000780376006 | 0.004062593831 |
| MPP5 | 1456.571509 | -0.4373396754 | 0.1755777296 | 0.004610987079 | 0.0185950668 |
| ZNF709 | 113.6846136 | -0.4378369288 | 0.1788739975 | 0.005210251151 | 0.02056067569 |
| PPIL2 | 1131.919361 | -0.4381340606 | 0.07809957418 | 8.28E-09 | 1.26E-07 |
| NPIPA8 | 179.2882437 | -0.4382258351 | 0.1572225045 | 0.001972860449 | 0.009068823473 |
| COG8 | 1038.167062 | -0.4383658096 | 0.09652357184 | 2.25E-06 | 2.14E-05 |
| AKAP10 | 1008.176765 | -0.4386341175 | 0.101645314 | 6.38E-06 | 5.57E-05 |
| ZNF169 | 63.33879038 | -0.4389118816 | 0.2194379268 | 0.01522602374 | 0.04996699744 |
| WDR92 | 448.6411672 | -0.4389353561 | 0.1414782655 | 0.000731233632 | 0.003829524286 |
| LETMD1 | 1049.187026 | -0.4390022806 | 0.132847135 | 0.000366189223 | 0.002083374606 |
| BET1 | 754.2601712 | -0.4392325216 | 0.1132810879 | 4.17E-05 | 0.000305510602 |

| | baseMean | log2FoldChange | lfcSE | pvalue | padj |
|---|---|---|---|---|---|
| SREK1 | 1744.967873 | -0.439409203 | 0.1123911898 | 3.66E-05 | 0.000271254669 |
| FAM200A | 223.3212541 | -0.4396752954 | 0.1275808533 | 0.000220666509 | 0.001342321094 |
| TTI1 | 1464.190741 | -0.4400633219 | 0.09110622315 | 5.49E-07 | 5.97E-06 |
| RP4-639F20.1 | 107.0053775 | -0.4404353273 | 0.1922191983 | 0.007661499428 | 0.02817289824 |
| NR2F2 | 532.9359945 | -0.4405774061 | 0.09597566986 | 1.78E-06 | 1.73E-05 |
| VRK3 | 784.6706101 | -0.4406254241 | 0.07938858776 | 1.16E-08 | 1.72E-07 |
| ZNF226 | 494.9174675 | -0.441368708 | 0.1070734621 | 1.49E-05 | 0.000121792033 |
| N6AMT1 | 243.4722982 | -0.4413707101 | 0.1410337948 | 0.000662627716 | 0.003508486512 |
| LOC650368 | 159.3854417 | -0.4413754842 | 0.1395840886 | 0.000594137108 | 0.003195848049 |
| L2HGDH | 145.4411388 | -0.4421067772 | 0.1789398563 | 0.004797110102 | 0.01921482036 |
| AGAP4 | 112.0892086 | -0.4421861838 | 0.1653536304 | 0.002728572897 | 0.01194171718 |
| PPP1R3D | 139.2521731 | -0.4425841569 | 0.160254841 | 0.002112718916 | 0.009609292041 |
| OPA3 | 866.9880729 | -0.4427028166 | 0.1139856147 | 3.60E-05 | 0.000267733674 |
| ZNF287 | 289.6850984 | -0.4431255246 | 0.1527139112 | 0.00137488889 | 0.006672333989 |
| PRICKLE3 | 96.50455458 | -0.4434568124 | 0.1685103037 | 0.003091636293 | 0.01329617931 |
| HSPA8 | 94746.76123 | -0.4436613047 | 0.1333166208 | 0.000331083059 | 0.001913624099 |
| RAB3D | 591.4771251 | -0.4437295304 | 0.1832269336 | 0.005379689594 | 0.02107847325 |
| RBMS3 | 1416.646735 | -0.4437970214 | 0.1461956602 | 0.000877250756 | 0.004489991924 |
| SAMD9L | 176.7135086 | -0.4443437395 | 0.1909927718 | 0.006894871966 | 0.02585186318 |
| RMND1 | 740.9162391 | -0.4465534264 | 0.1192002293 | 6.89E-05 | 0.000480824619 |
| DST | 30180.21247 | -0.4472875487 | 0.1642625108 | 0.002301028005 | 0.01031617162 |
| GAL3ST4 | 136.8450537 | -0.4476494198 | 0.192082822 | 0.00675289494 | 0.02542838967 |
| GPATCH1 | 381.7755785 | -0.4478553032 | 0.1101189606 | 1.83E-05 | 0.000146513359 |
| DIS3L | 990.1825597 | -0.4481568266 | 0.08846454033 | 1.60E-07 | 1.94E-06 |
| ALG10B | 158.8413045 | -0.4482466548 | 0.1594620344 | 0.001793995947 | 0.008360189562 |
| LOC730101 | 307.6756002 | -0.448521743 | 0.1705202493 | 0.003023308096 | 0.01303621032 |
| LOC283788 | 208.1683041 | -0.448812601 | 0.1588097106 | 0.001704315217 | 0.008009958733 |
| ZFAT | 433.6296732 | -0.4490140144 | 0.114066236 | 3.16E-05 | 0.000237366563 |
| TMEM186 | 272.6599842 | -0.4490230911 | 0.13340838 | 0.000284495454 | 0.00167623776 |
| FAM200B | 437.9454973 | -0.4494500011 | 0.126245416 | 0.000140426076 | 0.000904326925 |
| C2orf68 | 605.9690544 | -0.4498365293 | 0.09430133216 | 7.17E-07 | 7.58E-06 |
| LZTR1 | 1330.976734 | -0.4499203369 | 0.09544062581 | 9.43E-07 | 9.69E-06 |
| ZNF788 | 391.8029093 | -0.4499733232 | 0.1249888653 | 0.000119449827 | 0.000789737339 |
| LOC339803 | 304.247787 | -0.4499758912 | 0.113338374 | 2.75E-05 | 2.11E-04 |
| POC5 | 307.2899317 | -0.4506733784 | 0.1721866114 | 0.003094994677 | 0.01330677902 |
| GOLGA1 | 572.9885458 | -0.4517777614 | 0.09648143161 | 1.10E-06 | 1.12E-05 |
| PPP1R10 | 1761.602264 | -0.4522342571 | 0.1146512246 | 2.99E-05 | 0.000227174119 |
| FLJ20021 | 116.365257 | -0.4525541998 | 0.2113440934 | 0.01040338291 | 0.03648903843 |
| ZNF417 | 407.8214128 | -0.4527945483 | 0.1302720779 | 0.000189199122 | 0.001173255198 |
| C20orf194 | 929.0464879 | -0.4528579268 | 0.1236057869 | 9.31E-05 | 0.000631008748 |

| | baseMean | log2FoldChange | lfcSE | pvalue | padj |
|---|---|---|---|---|---|
| STK38 | 1451.432151 | -0.4528608234 | 0.08059395107 | 7.56E-09 | 1.16E-07 |
| MGME1 | 1083.246509 | -0.4531319625 | 0.1593430174 | 0.001579231129 | 0.007512195618 |
| IVNS1ABP | 2002.76275 | -0.4534689374 | 0.151589307 | 0.000998448938 | 0.005039290252 |
| SPICE1 | 552.5564681 | -0.4547645216 | 0.1039071283 | 4.51E-06 | 4.05E-05 |
| EYA3 | 1200.091613 | -0.4563209308 | 0.08806427143 | 8.50E-08 | 1.08E-06 |
| GGA3 | 523.6803059 | -0.4568707228 | 0.1086359869 | 9.64E-06 | 8.12E-05 |
| ANKRD54 | 573.176972 | -0.4572387087 | 0.113111426 | 1.97E-05 | 0.000156329129 |
| CYB561D1 | 306.3707515 | -0.4573330821 | 0.1242221707 | 8.55E-05 | 0.000584465218 |
| MKS1 | 418.1535084 | -0.4573894726 | 0.1093273667 | 1.07E-05 | 8.98E-05 |
| RABGAP1 | 2472.510934 | -0.4578745577 | 0.1594310624 | 0.001414552841 | 0.006822571187 |
| ALDOC | 2686.952971 | -0.4582670455 | 0.2200150147 | 0.01165465077 | 0.04013091936 |
| KIF15 | 1314.573947 | -0.4589156438 | 0.1750723244 | 0.002917620765 | 0.0126574754 |
| TRNT1 | 674.2057123 | -0.459243306 | 0.123746769 | 7.58E-05 | 0.000524343783 |
| ANKHD1-EIF4EBF | 746.1591356 | -0.4593690449 | 0.1623938074 | 0.00161525011 | 0.007653968362 |
| ZNF133 | 654.9424617 | -0.4593737072 | 0.1073536883 | 6.99E-06 | 6.06E-05 |
| TNK2 | 260.688555 | -0.4594742171 | 0.1417145982 | 0.000425557443 | 0.002390088563 |
| MOCS1 | 656.5345011 | -0.4604415555 | 0.09185662338 | 2.04E-07 | 2.41E-06 |
| E2F8 | 111.8054492 | -0.4608410315 | 0.2078522749 | 0.008444468945 | 0.0306017866 |
| CHRNB1 | 124.3700626 | -0.4615130291 | 0.194563809 | 0.005752434394 | 0.02230416555 |
| LOC101928123 | 167.2568731 | -0.4618068234 | 0.1817210446 | 0.003699440768 | 0.01550266637 |
| LOC145783 | 92.36183029 | -0.4621906989 | 0.2307247498 | 0.01364709766 | 0.04558932848 |
| ZNF234 | 296.0215088 | -0.4635212929 | 0.1345271968 | 0.000204494131 | 0.001253484202 |
| RAD50 | 1911.357374 | -0.4637195423 | 0.08808179266 | 5.30E-08 | 7.03E-07 |
| DEPDC5 | 466.1103192 | -0.4637646189 | 0.126465422 | 8.81E-05 | 0.000600420763 |
| FAM193B | 534.689585 | -0.4639092706 | 0.1296714425 | 0.000123931819 | 0.000813948328 |
| TMCO6 | 202.5118918 | -0.4643047221 | 0.1582443911 | 0.001150566059 | 0.005715975324 |
| TRMT13 | 460.1356561 | -0.46432859 | 0.1574840401 | 0.001095402581 | 0.005463802019 |
| LOC100506691 | 94.1878677 | -0.4646619689 | 0.2259020649 | 0.01204783324 | 0.04127477891 |
| SECISBP2 | 1009.608298 | -0.4649908902 | 0.08856543175 | 5.66E-08 | 7.47E-07 |
| INVS | 819.0211466 | -0.4655767966 | 0.07823741366 | 1.01E-09 | 1.73E-08 |
| TNRC6A | 2234.346361 | -0.4660237321 | 0.1184427648 | 2.99E-05 | 0.000226871311 |
| TK2 | 922.8430132 | -0.4667731171 | 0.1070948166 | 4.79E-06 | 4.29E-05 |
| HMGCR | 6812.643486 | -0.4671811857 | 0.1219406028 | 4.78E-05 | 0.000344654187 |
| STX17 | 726.8115375 | -0.4673570529 | 0.1306335171 | 0.000121356251 | 0.000799501427 |
| TRAF3IP2-AS1 | 135.1712053 | -0.4675797358 | 0.1952259201 | 0.005316934937 | 0.02088755786 |
| PMS2 | 590.3733378 | -0.4676009517 | 0.09346636396 | 2.08E-07 | 2.44E-06 |
| CDK15 | 828.7849722 | -0.4677044532 | 0.2245960845 | 0.01096975559 | 0.03814308522 |
| ZDHHC21 | 232.3875279 | -0.4678078763 | 0.1737429023 | 0.002359001199 | 0.01054116712 |
| PIGM | 241.5369757 | -0.4680997905 | 0.1137283264 | 1.40E-05 | 0.000114749106 |
| CLDN15 | 44.17816946 | -0.4682838317 | 0.2311224752 | 0.01282937893 | 0.04334373134 |

| | baseMean | log2FoldChange | lfcSE | pvalue | padj |
|---|---|---|---|---|---|
| EPOR | 200.6071852 | -0.4685717287 | 0.1700713283 | 0.001940204204 | 0.008951874929 |
| B3GNTL1 | 245.7543295 | -0.4686508479 | 0.1706688847 | 0.001996281571 | 0.00915387629 |
| CCDC117 | 651.0233161 | -0.46877518 | 0.08680535519 | 2.49E-08 | 3.47E-07 |
| RBM12B | 974.456964 | -0.4691184932 | 0.1085907393 | 5.66E-06 | 4.99E-05 |
| ZNF718 | 148.6658494 | -0.4713198659 | 0.1513271632 | 0.000631693477 | 0.003368654792 |
| SLC25A36 | 1029.731209 | -0.4715080495 | 0.1123503702 | 9.73E-06 | 8.20E-05 |
| CENPBD1 | 224.3547842 | -0.4718043586 | 0.1230831152 | 4.43E-05 | 3.22E-04 |
| FRYL | 2655.151526 | -0.4729928679 | 0.1496422147 | 0.000531369045 | 0.002898711115 |
| ACSS1 | 125.3912585 | -0.4732722168 | 0.1740626836 | 0.002130163297 | 0.009670223894 |
| DSTNP2 | 78.24027651 | -0.4735035822 | 0.2035707621 | 0.006210540705 | 0.02372818593 |
| ZNF569 | 242.4710358 | -0.4742926475 | 0.1658885997 | 0.001393924155 | 0.00674711858 |
| SUPT20H | 1833.443222 | -0.474782203 | 0.09133789107 | 7.09E-08 | 9.12E-07 |
| RNF122 | 104.5738602 | -0.4749658927 | 0.2360882141 | 0.01266617066 | 0.04291912037 |
| ZNF81 | 227.6948584 | -0.4750822412 | 0.1700101025 | 0.001694252642 | 0.007970214088 |
| ISL2 | 118.4138809 | -0.4753625726 | 0.2282604516 | 0.01083707934 | 0.03776814703 |
| TAF1 | 1457.315597 | -0.4758270797 | 0.07463284727 | 6.70E-11 | 1.37E-09 |
| NAA40 | 656.2587965 | -0.4760087885 | 0.1057390768 | 2.39E-06 | 2.26E-05 |
| CEP57L1 | 550.855979 | -0.4760216342 | 0.1523859018 | 0.000596577815 | 0.003203530248 |
| CNTRL | 917.2566461 | -0.4760612816 | 0.144126016 | 0.000323438481 | 0.001873803715 |
| RP5-1103G7.4 | 115.6072122 | -0.4764632985 | 0.2371556864 | 0.01274557515 | 0.04311948838 |
| DYNC1H1 | 14559.77087 | -0.4765783127 | 0.119866675 | 2.41E-05 | 0.000187296602 |
| CLDN7 | 107.5938651 | -0.4772298866 | 0.2082184763 | 0.006618106375 | 0.02503480331 |
| HECTD4 | 665.6320775 | -0.4772357188 | 0.1100028869 | 5.09E-06 | 4.51E-05 |
| CCNG2 | 509.9391835 | -0.4774677405 | 0.2080674223 | 0.006543629321 | 0.02479717408 |
| APOBEC3F | 150.8441811 | -0.4775270106 | 0.2298642298 | 0.01098419696 | 0.03818438211 |
| THAP3 | 622.8047487 | -0.4775752149 | 0.1071694672 | 2.95E-06 | 2.76E-05 |
| ZNF613 | 168.3179017 | -0.4779734634 | 0.1418307504 | 0.000253671628 | 0.001513187852 |
| ZNF480 | 647.4611866 | -0.4780333829 | 0.1398381398 | 0.000212917885 | 0.001300301229 |
| OXLD1 | 254.1456132 | -0.4785229423 | 0.1906320472 | 0.003783799074 | 0.01578656072 |
| LRRC20 | 294.3858307 | -0.478981387 | 0.1715579387 | 0.001676644782 | 0.007902662493 |
| GPC2 | 110.0232248 | -0.4790000446 | 0.2065487958 | 0.006158597644 | 0.02359633565 |
| GANC | 296.1674936 | -0.4791857374 | 0.1140908085 | 9.41E-06 | 7.96E-05 |
| APBB3 | 187.7692651 | -0.4797327288 | 0.1844033439 | 0.002910901044 | 0.01263568677 |
| BCL2 | 58.10957389 | -0.4797906989 | 0.233681503 | 0.01150494686 | 0.03969806578 |
| PLEKHA8 | 567.1675014 | -0.479920526 | 0.1045238062 | 1.52E-06 | 1.50E-05 |
| ZCCHC7 | 711.2215883 | -0.4799239468 | 0.08702605394 | 1.25E-08 | 1.85E-07 |
| SLC12A6 | 308.3323858 | -0.4802760597 | 0.1214584966 | 2.67E-05 | 0.000204673064 |
| SLITRK5 | 108.297457 | -0.4803806126 | 0.2298245018 | 0.01050666888 | 0.03680795129 |
| LOC101928409 | 114.8348128 | -0.4806302581 | 0.1764374483 | 0.002038400142 | 0.009333661839 |
| RHOT1 | 1057.926088 | -0.4806313299 | 0.09329549672 | 9.07E-08 | 1.15E-06 |

| | baseMean | log2FoldChange | lfcSE | pvalue | padj |
|---|---|---|---|---|---|
| MEX3B | 204.1795849 | -0.4812683875 | 0.1329322079 | 0.000100163433 | 0.000675116961 |
| PCNT | 1193.028206 | -0.4813042145 | 0.1376908052 | 0.000159003189 | 0.001009121262 |
| UBR2 | 1951.740406 | -0.4823346087 | 0.08271483179 | 1.97E-09 | 3.28E-08 |
| SALL2 | 202.9307136 | -0.4833304506 | 0.1878975043 | 0.003098196966 | 0.01331222588 |
| ZNF582 | 64.70716228 | -0.4836158072 | 0.2331444741 | 0.01076265328 | 0.0375721324 |
| EPM2AIP1 | 1044.992382 | -0.485016426 | 0.1360641532 | 0.000121736890 | 0.000800945893 |
| ATP8B3 | 69.62937712 | -0.4852796165 | 0.2088133572 | 0.005987685904 | 0.02306017988 |
| HILPDA | 922.0598315 | -0.48572784 | 0.1991577606 | 0.004370564794 | 0.01776989056 |
| NUDT16 | 1401.953493 | -0.4859212702 | 0.07502601532 | 3.32E-11 | 7.09E-10 |
| NFKBIL1 | 320.2467512 | -0.486783298 | 0.219508507 | 0.00760416456 | 0.02800356323 |
| SPATA7 | 343.4462239 | -0.4868183245 | 0.1469313777 | 0.000299879924 | 0.001757839449 |
| FBF1 | 191.3822993 | -0.4868472218 | 0.1323367425 | 7.80E-05 | 0.000538342998 |
| MAP3K14 | 283.1216403 | -0.4868596595 | 0.1364723004 | 0.000120203777 | 0.000792798577 |
| SLC22A5 | 92.80614397 | -0.4869389546 | 0.1988312085 | 0.004292879383 | 0.01750182945 |
| ANKRD36B | 598.5581541 | -0.4870443914 | 0.1017276973 | 5.85E-07 | 6.31E-06 |
| CDAN1 | 646.0096922 | -0.4874337627 | 0.1017965874 | 5.77E-07 | 6.26E-06 |
| CARD8 | 456.5908703 | -0.4876060289 | 0.1078178899 | 2.11E-06 | 2.02E-05 |
| GEMIN8 | 548.9059515 | -0.4879645876 | 0.08978003629 | 1.93E-08 | 2.76E-07 |
| NCK1 | 1275.920813 | -0.4881085772 | 0.1062253 | 1.46E-06 | 1.44E-05 |
| ABCC5 | 537.3155191 | -0.4881591479 | 0.1490127968 | 0.000331729158 | 0.001916614451 |
| LNX2 | 448.337757 | -0.4884792555 | 0.1222877833 | 2.17E-05 | 0.000170411003 |
| ZNF83 | 572.4419511 | -0.4890153201 | 0.1201598534 | 1.58E-05 | 0.000128591383 |
| BCL9L | 350.5496823 | -0.4891575747 | 0.1535093365 | 0.000463720551 | 0.002569533045 |
| LOC102723773 | 437.7332546 | -0.4894929218 | 0.1068954072 | 1.59E-06 | 1.56E-05 |
| INSC | 158.0722055 | -0.4902733532 | 0.2459361311 | 0.01241801556 | 0.04222238724 |
| NEU3 | 225.874221 | -0.490467742 | 0.1455110882 | 0.000241966092 | 0.001454434053 |
| EXD3 | 118.7749057 | -0.4905384901 | 0.2411229285 | 0.01139582665 | 0.03940178623 |
| WDSUB1 | 191.6436652 | -0.4910200364 | 0.1460153719 | 0.000248877679 | 0.001491763191 |
| MED22 | 1040.797334 | -0.4913169538 | 0.102386399 | 5.44E-07 | 5.94E-06 |
| ADCK5 | 175.8682312 | -0.4916018373 | 0.1627015246 | 0.000787819768 | 0.004097048039 |
| ENTHD2 | 288.2599717 | -0.4916888144 | 0.1164245288 | 8.07E-06 | 6.93E-05 |
| LUC7L | 999.6645115 | -0.4918100426 | 0.0893419454 | 1.26E-08 | 1.86E-07 |
| C18orf54 | 289.344696 | -0.4918386869 | 0.2424562388 | 0.01144157166 | 0.03951601957 |
| MID1IP1 | 1410.282042 | -0.4924994913 | 0.1200753883 | 1.36E-05 | 0.000111467915 |
| ZNF43 | 303.7082866 | -0.4925588971 | 0.110298897 | 2.70E-06 | 2.54E-05 |
| TFB1M | 1067.066498 | -0.4927745916 | 0.1705402849 | 0.00118884381 | 0.005884539725 |
| BOLA1 | 281.844305 | -0.4931920061 | 0.1775946714 | 0.001665991288 | 0.007862105953 |
| ANKRD36 | 882.9650495 | -0.4934518737 | 0.1541601399 | 0.000432514921 | 0.002426418487 |
| NUDT6 | 170.5946497 | -0.4939413781 | 0.1871765571 | 0.002481391802 | 0.01102519921 |
| CDC42EP2 | 771.3911839 | -0.4945286819 | 0.157221308 | 0.000519483835 | 0.002841511695 |

| | baseMean | log2FoldChange | lfcSE | pvalue | padj |
|---|---|---|---|---|---|
| ZNF737 | 227.4551583 | -0.4945949534 | 0.1451443106 | 0.000209737332 | 0.001282982391 |
| LOC80154 | 213.4089186 | -0.4946643848 | 0.1716215091 | 0.00120802949 | 0.005963644258 |
| FAT4 | 2802.132863 | -0.4953664163 | 0.1521310311 | 0.000354218761 | 0.002029227834 |
| TMEM106A | 138.8185388 | -0.4961764778 | 0.1852612621 | 0.002200655086 | 0.009941005332 |
| MINA | 2263.016896 | -0.4965152868 | 0.0779823475 | 6.62E-11 | 1.35E-09 |
| NOL3 | 349.7165144 | -0.4965919878 | 0.1436953573 | 0.000173903971 | 0.001094360202 |
| NVL | 971.3307969 | -0.4974790207 | 0.1023132197 | 3.87E-07 | 4.33E-06 |
| METTL15 | 815.0296878 | -0.4975239655 | 0.09057663287 | 1.34E-08 | 1.97E-07 |
| GTSE1 | 1041.353963 | -0.4981944328 | 0.185379031 | 0.002130319993 | 0.009670223894 |
| TAB1 | 724.71827 | -0.4985258549 | 0.1290645905 | 3.62E-05 | 0.000269015221 |
| ZNF37A | 699.1345616 | -0.4987800997 | 0.1275657985 | 3.03E-05 | 0.000229383998 |
| SLFN12 | 539.6815956 | -0.4990065705 | 0.1157633237 | 5.34E-06 | 4.71E-05 |
| CABYR | 213.1072604 | -0.4992067122 | 0.1267250793 | 2.64E-05 | 2.03E-04 |
| GJC2 | 39.06071828 | -0.499602426 | 0.2467633489 | 0.01139791076 | 0.03940178623 |
| MKL2 | 352.5115019 | -0.5000463539 | 0.1595712687 | 0.000531869208 | 0.002899670684 |
| ANKRD26 | 712.4805979 | -0.5003319529 | 0.1125187503 | 2.87E-06 | 2.69E-05 |
| TRIM65 | 1084.521134 | -0.5005172225 | 0.09643392315 | 6.97E-08 | 9.00E-07 |
| TSTD2 | 767.8311365 | -0.5006324646 | 0.09463788557 | 4.12E-08 | 5.56E-07 |
| LOC729603 | 42.12999985 | -0.5011643253 | 0.2543218234 | 0.01265550584 | 0.04290251057 |
| PIDD | 286.4043683 | -0.5016212925 | 0.1741836198 | 0.001178890282 | 0.005844987482 |
| LOC100996442 | 6.373613107 | -0.5022991736 | 1.094643706 | 0.01081186888 | 0.03769974609 |
| ZNF318 | 615.4970074 | -0.5023428984 | 0.1073544173 | 9.51E-07 | 9.76E-06 |
| LYRM7 | 497.871215 | -0.5030028459 | 0.1171898875 | 5.74E-06 | 5.05E-05 |
| FPGT-TNNI3K | 52.75552426 | -0.5031245753 | 0.2681607043 | 0.01515855826 | 0.04980047969 |
| ABCC10 | 501.4609361 | -0.503138178 | 0.107500557 | 9.35E-07 | 9.63E-06 |
| ARHGEF9 | 537.5535124 | -0.5034098478 | 0.1203642005 | 9.34E-06 | 7.92E-05 |
| KIAA0753 | 900.0541057 | -0.5037144892 | 0.1207658195 | 9.70E-06 | 8.18E-05 |
| NR2C1 | 575.3013135 | -0.5037505105 | 0.1483645903 | 0.000212420032 | 0.001297793135 |
| SETDB1 | 1167.839097 | -0.5040775004 | 0.08074937023 | 1.44E-10 | 2.78E-09 |
| FOXN3-AS1 | 65.7663752 | -0.5040827319 | 0.2288165187 | 0.007433360794 | 0.02750380439 |
| ACBD4 | 94.19868848 | -0.5043168005 | 0.2454190318 | 0.01031383827 | 0.03623471873 |
| TXNIP | 15306.49245 | -0.5047847875 | 0.2265122003 | 0.007351292883 | 0.02727470714 |
| ANGEL2 | 889.0584297 | -0.5051418278 | 0.08774988115 | 2.73E-09 | 4.45E-08 |
| ZNF600 | 179.3549343 | -0.5052354749 | 0.1395936842 | 9.28E-05 | 0.000629488469 |
| ARHGEF28 | 3255.227135 | -0.5052815179 | 0.1666656665 | 0.000729870828 | 0.003826509809 |
| KLHL12 | 1719.703324 | -0.5065451452 | 0.06048209358 | 1.86E-17 | 7.59E-16 |
| RHEBP2 | 83.72475905 | -0.507216883 | 0.2111344348 | 0.004521049085 | 0.01831172465 |
| SCD | 25082.58125 | -0.5079705152 | 0.1158229473 | 3.61E-06 | 3.30E-05 |
| C8orf58 | 529.7600296 | -0.508124378 | 0.1096929636 | 1.15E-06 | 1.16E-05 |
| MYO9A | 3014.898736 | -0.5086631651 | 0.09396338436 | 1.96E-08 | 2.80E-07 |

| | baseMean | log2FoldChange | lfcSE | pvalue | padj |
|---|---|---|---|---|---|
| FANCL | 359.9258593 | -0.5091461631 | 0.1372361628 | 6.40E-05 | 0.000449415216 |
| MBD6 | 81.64290433 | -0.5097589163 | 0.2188474918 | 0.00540926116 | 0.02116092733 |
| SRGAP2 | 1109.596922 | -0.5103295045 | 0.06621296932 | 4.26E-15 | 1.40E-13 |
| MBOAT1 | 246.5656911 | -0.5109145767 | 0.1889450109 | 0.001938931692 | 0.008951551616 |
| ATHL1 | 401.7910803 | -0.5114622353 | 0.2299545606 | 0.006824110626 | 0.02563819912 |
| ZNF747 | 204.1304014 | -0.511661493 | 0.1962226454 | 0.002537705044 | 0.01122516055 |
| NFYC-AS1 | 90.11462725 | -0.5119123817 | 0.2383283255 | 0.008153595048 | 0.0296963984 |
| AKAP9 | 2130.257555 | -0.5121119238 | 0.1280878741 | 1.96E-05 | 0.000156282609 |
| BMS1P6 | 298.9516788 | -0.5129078187 | 0.1387138162 | 6.68E-05 | 0.000467683787 |
| UAP1L1 | 293.4231037 | -0.5129246969 | 0.1565433917 | 0.000312845675 | 0.001820937942 |
| HEXDC | 215.5693409 | -0.5129880047 | 0.2130936044 | 0.004323513554 | 0.01760730616 |
| TMOD2 | 456.0610039 | -0.5130076414 | 0.1922795841 | 0.002135438369 | 0.009684600022 |
| HERC2P3 | 114.677143 | -0.5131986377 | 0.2181903072 | 0.00497197632 | 0.0197935175 |
| MORN1 | 48.5874846 | -0.5132511582 | 0.276376617 | 0.01496085939 | 0.04923789466 |
| CDPF1 | 240.6113723 | -0.5134562426 | 0.1623111041 | 0.000458497280 | 0.00254437794 |
| PDE5A | 12443.58058 | -0.5136962308 | 0.236297642 | 0.008089847388 | 0.02949307976 |
| ANKRD1 | 577.8046344 | -0.5141761262 | 0.2455188876 | 0.008990777738 | 0.03231081094 |
| LUC7L3 | 1949.182405 | -0.5144276768 | 0.1350198851 | 4.23E-05 | 0.000309668665 |
| RMI2 | 135.5836298 | -0.5145622452 | 0.1746020047 | 0.000919518055 | 0.004682183422 |
| TEP1 | 809.899716 | -0.5148958338 | 0.1052248743 | 3.12E-07 | 3.56E-06 |
| SMARCAL1 | 1082.297032 | -0.5170575413 | 0.1086003307 | 6.01E-07 | 6.47E-06 |
| LDB1 | 2516.669702 | -0.5174632928 | 0.1553544208 | 0.000225914042 | 0.001367330969 |
| TMEM99 | 370.899815 | -0.5175226301 | 0.11430803 | 1.86E-06 | 1.80E-05 |
| CEP350 | 2472.438541 | -0.5175746783 | 0.08801119993 | 1.31E-09 | 2.21E-08 |
| NAT9 | 1111.394407 | -0.5186637654 | 0.07804859599 | 9.37E-12 | 2.15E-10 |
| TCF12 | 4413.938889 | -0.5189444275 | 0.1318386364 | 2.49E-05 | 0.000192414904 |
| FAM156B | 421.865403 | -0.5190307912 | 0.1735123054 | 0.000789770594 | 0.004104326135 |
| TRIM34 | 179.6004054 | -0.5190672162 | 0.176064011 | 0.000905985937 | 0.004619597476 |
| OPHN1 | 274.8893067 | -0.5201697638 | 0.1176707529 | 3.03E-06 | 2.82E-05 |
| THTPA | 373.0702135 | -0.520451252 | 0.1298785311 | 1.84E-05 | 0.000147139385 |
| NOC3L | 1661.060968 | -0.5208036897 | 0.1694121426 | 0.000595989412 | 0.003202340802 |
| BBS2 | 1044.789274 | -0.5208670203 | 0.1573605001 | 0.000269614972 | 0.001599959075 |
| CHD9 | 2869.302085 | -0.5209126766 | 0.120786521 | 4.99E-06 | 4.45E-05 |
| AMZ2P1 | 162.0329129 | -0.5210149047 | 0.1827880007 | 0.001215266605 | 0.005995395784 |
| ZNF641 | 218.4123588 | -0.5211511398 | 0.255556145 | 0.009993180956 | 0.03528301429 |
| DHFRL1 | 218.9927872 | -0.5212725298 | 0.1288740923 | 1.60E-05 | 0.000129143991 |
| NAPB | 173.497502 | -0.5214429455 | 0.1339861871 | 3.03E-05 | 0.000229546782 |
| ZNF783 | 292.6395152 | -0.5218855423 | 0.1268396743 | 1.18E-05 | 9.78E-05 |
| PARP3 | 235.1205029 | -0.5223975161 | 0.1594831595 | 0.000302548270 | 0.001771388594 |
| CXorf23 | 171.7402665 | -0.5238415647 | 0.1710204392 | 0.000618576675 | 0.003305899859 |

| | baseMean | log2FoldChange | lfcSE | pvalue | padj |
|---|---|---|---|---|---|
| USP40 | 1322.805441 | -0.5239638928 | 0.07887969856 | 1.01E-11 | 2.30E-10 |
| LOC100506639 | 88.05151693 | -0.5240666801 | 0.1892710225 | 0.001549639808 | 0.007390322582 |
| ST3GAL4-AS1 | 106.7469857 | -0.5241343721 | 0.2141799264 | 0.003753672488 | 0.01568137757 |
| KIAA1468 | 1085.108473 | -0.524976553 | 0.08045797925 | 2.16E-11 | 4.71E-10 |
| TP73-AS1 | 607.4529565 | -0.5261219441 | 0.1080749187 | 3.45E-07 | 3.92E-06 |
| ZC3H4 | 577.5616458 | -0.5263275543 | 0.1189459393 | 2.91E-06 | 2.72E-05 |
| MSANTD4 | 1072.823042 | -0.5267489562 | 0.09959864302 | 3.80E-08 | 5.15E-07 |
| DISC1 | 378.3387475 | -0.5269755837 | 0.1962187915 | 0.001928663717 | 0.008906908832 |
| PSMD6-AS2 | 38.83620241 | -0.5269934805 | 0.2786611786 | 0.01332225979 | 0.04467457795 |
| LRRC37A4P | 301.2928077 | -0.5274808351 | 0.1354484931 | 2.91E-05 | 0.000221620356 |
| RP11-540A21.2 | 112.3484449 | -0.527658037 | 0.1929641849 | 0.001674978687 | 0.007899511457 |
| MED20 | 1285.847279 | -0.5284789316 | 0.07630911119 | 1.35E-12 | 3.46E-11 |
| ZNF420 | 216.5233224 | -0.5286193487 | 0.1647700993 | 0.000374884740 | 0.002125548584 |
| IRAK4 | 806.5006188 | -0.5287884268 | 0.1113246456 | 6.14E-07 | 6.60E-06 |
| KRT19 | 53.83476394 | -0.5288560733 | 0.2266840829 | 0.004960964008 | 0.01976024428 |
| C2orf42 | 346.4892726 | -0.5289924636 | 0.117515485 | 2.00E-06 | 1.93E-05 |
| RWDD3 | 268.2658636 | -0.5300942245 | 0.1122746086 | 7.08E-07 | 7.52E-06 |
| GAS6-AS2 | 259.593108 | -0.5303319676 | 0.1974387503 | 0.001914498548 | 0.00885247481 |
| PUS7L | 850.4577611 | -0.5304719608 | 0.1917512596 | 0.001489041576 | 0.00714543196 |
| CARKD | 1190.878851 | -0.5307973114 | 0.1022469879 | 6.30E-08 | 8.22E-07 |
| ZC3H10 | 320.6286309 | -0.531013254 | 0.1253996754 | 6.79E-06 | 5.90E-05 |
| TRIM4 | 815.6859254 | -0.5316319423 | 0.1124182426 | 7.18E-07 | 7.59E-06 |
| AFG3L1P | 247.8757318 | -0.531829253 | 0.212862416 | 0.003176475972 | 0.01360602726 |
| KDM4A | 2280.556349 | -0.5323059114 | 0.09884670175 | 2.16E-08 | 3.07E-07 |
| FAM63B | 565.0967275 | -0.5323236044 | 0.1469422551 | 8.31E-05 | 0.000570634997 |
| DISP1 | 136.612133 | -0.5323545816 | 0.1752363474 | 0.000655770734 | 0.003479604585 |
| FBXO48 | 57.73585495 | -0.5330228677 | 0.2213531835 | 0.004075257537 | 0.01678000815 |
| SCAI | 226.6324437 | -0.5335719305 | 0.1471587708 | 8.20E-05 | 0.000563114463 |
| GLIDR | 127.6056074 | -0.533627223 | 0.1963673452 | 0.001734039434 | 0.008124012945 |
| CPT2 | 579.9582733 | -0.5338816849 | 0.1168754127 | 1.45E-06 | 1.43E-05 |
| LOC100129034 | 488.7890924 | -0.5341481223 | 0.1578266539 | 0.000200033250 | 0.001233759348 |
| CCDC142 | 187.9748837 | -0.5344713077 | 0.1760950226 | 0.000648141506 | 0.00344280374 |
| USP30 | 271.098384 | -0.5347642652 | 0.1510720677 | 1.13E-04 | 0.000753647212 |
| CCDC14 | 2070.255214 | -0.5350607718 | 0.1348761022 | 2.08E-05 | 0.000164567307 |
| MUM1 | 752.176838 | -0.5356083933 | 0.1238653154 | 4.46E-06 | 4.01E-05 |
| NEIL3 | 385.5446931 | -0.5359708596 | 0.1629475777 | 0.000276034741 | 0.00163284913 |
| TVP23C-CDRT4 | 165.5763955 | -0.536302011 | 0.2101331019 | 0.002721946066 | 0.01191972205 |
| PSIMCT-1 | 141.1517652 | -0.537172104 | 0.1586972482 | 0.000197747764 | 0.001220674325 |
| CSTF3 | 1430.963836 | -0.5376839439 | 0.1220658692 | 3.11E-06 | 2.88E-05 |
| RMND5B | 457.8007968 | -0.5381438346 | 0.1159095068 | 1.00E-06 | 1.02E-05 |

| | baseMean | log2FoldChange | lfcSE | pvalue | padj |
|---|---|---|---|---|---|
| CRYBG3 | 3129.364111 | -0.5382213946 | 0.1587878152 | 0.000192208569 | 0.001190429864 |
| SYNRG | 1215.97701 | -0.5383885502 | 0.09286210626 | 2.01E-09 | 3.34E-08 |
| COX15 | 1669.963947 | -0.5391357761 | 0.07687768429 | 7.04E-13 | 1.87E-11 |
| GTF2IRD2B | 547.7790587 | -0.5393034948 | 0.2497260131 | 0.007086443834 | 0.02644362462 |
| RAB30 | 213.5286693 | -0.539736975 | 0.1958641109 | 0.001508267734 | 0.007223093691 |
| C14orf132 | 942.3227135 | -0.5402294272 | 0.2528692623 | 0.007480472169 | 0.0276437702 |
| SLC5A3 | 1440.877018 | -0.540494972 | 0.1783748864 | 0.000643408537 | 0.003422547235 |
| STPG1 | 281.6839974 | -0.5405696224 | 0.1832461057 | 0.000832392852 | 0.004292863586 |
| ZMYM3 | 957.2896588 | -0.5407004444 | 0.1008424757 | 2.42E-08 | 3.39E-07 |
| ZNF84 | 821.924327 | -0.5407614898 | 0.1063944697 | 1.09E-07 | 1.36E-06 |
| TNRC6B | 1295.012158 | -0.5421252262 | 0.0823295356 | 1.37E-11 | 3.07E-10 |
| FICD | 116.9270876 | -0.5422116955 | 0.2509821932 | 0.007041821499 | 0.02632329407 |
| IRF2 | 390.9748383 | -0.542638192 | 0.1019541608 | 3.02E-08 | 4.15E-07 |
| ELMOD3 | 396.3932128 | -0.5428199002 | 0.141252648 | 3.38E-05 | 0.000252706976 |
| PDDC1 | 995.4287388 | -0.542839671 | 0.1066750915 | 1.04E-07 | 1.31E-06 |
| ESPL1 | 506.2367455 | -0.5429156191 | 0.2096654377 | 0.002380773149 | 0.01062569886 |
| CEP44 | 513.6586982 | -0.5429231512 | 0.1320104519 | 1.11E-05 | 9.26E-05 |
| TOX | 1201.064919 | -0.544893142 | 0.1387632973 | 2.39E-05 | 0.000185678772 |
| WDR25 | 495.5907955 | -0.5460307381 | 0.1383769667 | 2.20E-05 | 0.000172215668 |
| FBN1 | 21547.13456 | -0.5465317572 | 0.1933231953 | 0.001302924369 | 0.006360406864 |
| PYROXD2 | 203.2995488 | -0.5466639025 | 0.2422037101 | 0.005546900254 | 0.02163683465 |
| BEND4 | 65.54867017 | -0.547049487 | 0.291099821 | 0.01261138308 | 0.04278215601 |
| CNTNAP3B | 606.8392755 | -0.5479160306 | 0.1189917453 | 1.17E-06 | 1.17E-05 |
| NCKIPSD | 641.9457059 | -0.5482166718 | 0.08513564965 | 3.55E-11 | 7.50E-10 |
| NSD1 | 3649.804566 | -0.5486239905 | 0.07254389521 | 1.16E-14 | 3.73E-13 |
| TMEM45A | 628.1383218 | -0.5486639617 | 0.1957866627 | 0.001273506076 | 0.00624135351 |
| UBE2Q2P1 | 71.75992615 | -0.5488824963 | 0.2529284688 | 0.006829851165 | 0.0256468232 |
| ZNF37BP | 287.174916 | -0.5492707961 | 0.195602309 | 0.001254195591 | 0.00615887802 |
| RBM43 | 159.1101471 | -0.5493098312 | 0.1711798395 | 0.000351804121 | 0.002016170735 |
| MTMR4 | 2308.688361 | -0.5494032903 | 0.1127829111 | 3.14E-07 | 3.59E-06 |
| ZNF44 | 188.7176156 | -0.5505766495 | 0.1380344008 | 1.84E-05 | 0.000146800652 |
| RP11-85F14.5 | 54.33866733 | -0.5507721775 | 0.2424019024 | 0.005314199726 | 0.02088232244 |
| MAP10 | 114.6831501 | -0.5520828084 | 0.1732193051 | 0.000376283276 | 0.002132653863 |
| CUL9 | 761.9067881 | -0.5535078266 | 0.1016463749 | 1.47E-08 | 2.15E-07 |
| ZNF300 | 531.0667117 | -0.5535230818 | 0.1599032027 | 0.000136779197 | 0.000884667882 |
| SBF2-AS1 | 503.6897619 | -0.553984391 | 0.2565000727 | 0.006750854422 | 0.02542838967 |
| MSH5-SAPCD1 | 130.0427214 | -0.5540425707 | 0.2443297955 | 0.005241435431 | 0.02065089498 |
| HOXB5 | 38.64763985 | -0.554188807 | 0.304818235 | 0.01392933249 | 0.0463968303 |
| GUSBP11 | 231.2148549 | -0.5543751218 | 0.1944418843 | 0.00108631421 | 0.00542392095 |
| ZNF786 | 142.3462491 | -0.5545106671 | 0.162948084 | 0.000175931898 | 0.001103855895 |

| | baseMean | log2FoldChange | lfcSE | pvalue | padj |
|---|---|---|---|---|---|
| LOC101929479 | 307.1376897 | -0.5546204172 | 0.2081402318 | 0.001852847953 | 0.008608133059 |
| LINC00857 | 72.81436565 | -0.5547211258 | 0.2104858081 | 0.002031091014 | 0.009307760574 |
| ZNF235 | 89.43848257 | -0.5549747087 | 0.2141209531 | 0.002283821199 | 0.01025137589 |
| LMLN | 141.1989833 | -0.5554161598 | 0.1955967258 | 0.001120038768 | 0.005581076713 |
| WEE1 | 955.0808601 | -0.5556117057 | 0.1487092646 | 4.75E-05 | 0.000343196409 |
| LPP-AS2 | 70.05317084 | -0.5557449259 | 0.2256320662 | 0.003209595944 | 0.0137402039 |
| ZNF790 | 122.3917 | -0.5558651779 | 0.2262658406 | 0.003259123449 | 0.01393998536 |
| HOXB4 | 183.234775 | -0.5562239581 | 0.159763924 | 1.30E-04 | 0.000846533131 |
| FRA10AC1 | 296.542774 | -0.5569751459 | 0.1284364577 | 3.97E-06 | 3.61E-05 |
| ING4 | 561.9731455 | -0.5575136194 | 0.1859061699 | 0.000680844907 | 0.003592168612 |
| LIPE | 96.46265163 | -0.5577880767 | 0.2418307597 | 0.004711096319 | 0.01893723356 |
| PAN3-AS1 | 36.04206918 | -0.5578227266 | 0.2821511226 | 0.01009224966 | 0.03555691083 |
| KIAA1328 | 203.3471221 | -0.5579043875 | 0.1692232111 | 0.000250949569 | 0.001500557483 |
| ZNF823 | 210.1262077 | -0.5579119874 | 0.2154891097 | 0.00227085874 | 0.01020857964 |
| IFFO1 | 489.6494409 | -0.5583693955 | 0.11958278 | 8.26E-07 | 8.62E-06 |
| SPOP | 1508.71375 | -0.5584806812 | 0.07101912311 | 1.07E-15 | 3.73E-14 |
| APOL2 | 704.9801165 | -0.5585218311 | 0.1200295198 | 8.93E-07 | 9.22E-06 |
| ZBTB26 | 109.4707995 | -0.5592027187 | 0.1699288333 | 0.000257242549 | 0.001530809078 |
| SPEF2 | 71.32577192 | -0.5594753584 | 0.2357689182 | 0.004031641453 | 0.01663005711 |
| GGACT | 86.40560633 | -0.5597140134 | 0.2240240096 | 0.002881873893 | 0.01253160642 |
| SRBD1 | 1045.558277 | -0.5598285678 | 0.1128524907 | 1.91E-07 | 2.27E-06 |
| WDR27 | 195.3451321 | -0.5601110969 | 0.1618590779 | 0.000137496220 | 0.000887762893 |
| TSTD3 | 97.20778855 | -0.5603987952 | 0.2057297513 | 0.001545840614 | 0.007374566135 |
| CNKSR3 | 227.8917197 | -0.5607199832 | 0.1474807905 | 3.78E-05 | 0.000279628456 |
| FAM198A | 62.3762057 | -0.5615641604 | 0.2590095035 | 0.00654201931 | 0.02479717408 |
| ZNF585B | 326.6826109 | -0.5618795305 | 0.1195200832 | 7.13E-07 | 7.56E-06 |
| TMCC1 | 162.4526742 | -0.5623332362 | 0.1752552775 | 0.000331990993 | 0.001917383205 |
| EHHADH | 293.951332 | -0.5625819776 | 0.1316619545 | 5.17E-06 | 4.58E-05 |
| RPP25L | 530.5467145 | -0.5630269551 | 0.1119056496 | 1.33E-07 | 1.63E-06 |
| AP001258.4 | 48.32272587 | -0.5633024461 | 0.2874238833 | 0.01011050348 | 0.03560437236 |
| STK36 | 542.6924549 | -0.5638617539 | 0.1259008955 | 1.94E-06 | 1.87E-05 |
| ZNF354A | 220.5060055 | -0.565917975 | 0.1306942216 | 3.97E-06 | 3.60E-05 |
| DNHD1 | 148.4715269 | -0.5662337252 | 0.1544847171 | 6.38E-05 | 0.000448709301 |
| ATXN7L1 | 140.7683186 | -0.5664418977 | 0.1608772062 | 0.000108261390 | 0.000724451164 |
| VCAN | 5831.259662 | -0.5675081966 | 0.1960783477 | 0.000901817474 | 0.004601940117 |
| ZNF888 | 190.9864145 | -0.5676466733 | 0.1682427081 | 0.000185908798 | 0.001157672984 |
| NFYB | 772.7313579 | -0.5686873145 | 0.121258521 | 7.29E-07 | 7.69E-06 |
| ZNF418 | 130.5807056 | -0.5686923008 | 0.1686457678 | 0.000188545360 | 0.001170175854 |
| PXMP4 | 435.3483743 | -0.5687253038 | 0.1128764904 | 1.26E-07 | 1.56E-06 |
| PLEKHA2 | 1308.645974 | -0.5692695541 | 0.1004963162 | 3.98E-09 | 6.34E-08 |

| | baseMean | log2FoldChange | lfcSE | pvalue | padj |
|---|---|---|---|---|---|
| LRRC37A3 | 89.44257164 | -0.5696431979 | 0.2065774983 | 0.001378183756 | 0.006683136126 |
| ZNF518A | 644.8645419 | -0.5703875087 | 0.1246185864 | 1.25E-06 | 1.25E-05 |
| RNF213 | 2838.446251 | -0.5706869324 | 0.1280859038 | 2.18E-06 | 2.07E-05 |
| NARFL | 581.0075289 | -0.5712548293 | 0.1414437408 | 1.37E-05 | 0.000112359484 |
| PLA2G6 | 97.19648489 | -0.5713476825 | 0.3044184792 | 0.01148281618 | 0.03963088783 |
| MVK | 1358.4832 | -0.5719052157 | 0.1493589199 | 2.92E-05 | 0.000222007808 |
| DZANK1 | 48.96808501 | -0.5719449982 | 0.3106905869 | 0.01237226888 | 0.04212456238 |
| GPAM | 819.7416146 | -0.5721911011 | 0.1539178588 | 5.05E-05 | 0.000363457588 |
| LOC100132815 | 174.8368125 | -0.5732804692 | 0.1336363609 | 4.67E-06 | 4.19E-05 |
| LOC642423 | 56.451613 | -0.5737846641 | 0.2771438677 | 0.007817001066 | 0.02867389231 |
| RXRB | 1180.422447 | -0.5740269783 | 0.1120612926 | 7.92E-08 | 1.01E-06 |
| LINC00565 | 89.12960054 | -0.5744117171 | 0.1979289123 | 0.000871276092 | 0.004465552405 |
| ZMYM4 | 2804.977302 | -0.5745227136 | 0.07847633877 | 6.80E-14 | 2.02E-12 |
| ZNF346 | 435.7518691 | -0.5747037596 | 0.1028523557 | 6.11E-09 | 9.42E-08 |
| C6orf203 | 317.9203623 | -0.5747381701 | 0.1133661496 | 1.05E-07 | 1.32E-06 |
| CDO1 | 35.12596908 | -0.5754281464 | 0.3085800438 | 0.01169434874 | 0.04022110381 |
| KCTD18 | 424.7019008 | -0.5758973526 | 0.134639073 | 4.85E-06 | 4.33E-05 |
| FAM185A | 138.4623642 | -0.5764747805 | 0.1501574965 | 3.10E-05 | 0.000234380497 |
| MCM3AP-AS1 | 33.91902612 | -0.5770274622 | 0.3207992282 | 0.01315300035 | 0.04423650828 |
| ACACA | 4927.084016 | -0.5772398007 | 0.09377596388 | 1.96E-10 | 3.70E-09 |
| LOC100506714 | 114.335287 | -0.5773795869 | 0.1569642747 | 5.86E-05 | 0.000416179192 |
| TDRKH | 146.4956607 | -0.5774129065 | 0.1630726951 | 9.76E-05 | 0.000659151129 |
| RNF146 | 754.7587621 | -0.5782185595 | 0.08285373964 | 7.98E-13 | 2.10E-11 |
| TIGD1 | 160.9789589 | -0.5788805744 | 0.1482780894 | 2.37E-05 | 0.000184528081 |
| SLC24A1 | 146.8534285 | -0.579945122 | 0.2120325903 | 0.001401341258 | 0.006767891031 |
| XPC | 2069.779885 | -0.5801624666 | 0.1058098505 | 1.06E-08 | 1.59E-07 |
| PAQR7 | 684.9227262 | -0.5802701959 | 0.1576842628 | 5.00E-05 | 0.000359848866 |
| NFYA | 1069.047006 | -0.580902103 | 0.1135523797 | 8.12E-08 | 1.03E-06 |
| TRIM68 | 378.3068624 | -0.5809437784 | 0.1184342846 | 2.41E-07 | 2.80E-06 |
| GPR126 | 94.75493729 | -0.581214875 | 0.3053277548 | 0.01048152491 | 0.03672850658 |
| TBC1D32 | 161.8757259 | -0.5812625411 | 0.1578720082 | 5.65E-05 | 0.000402614960 |
| MXI1 | 325.5718371 | -0.5830472805 | 0.2870683809 | 0.00800122074 | 0.0292128925 |
| POLI | 509.1701173 | -0.5850804853 | 0.13164915 | 2.21E-06 | 2.10E-05 |
| HOXA10 | 562.696372 | -0.5852734461 | 0.1598061318 | 6.02E-05 | 0.000426270248 |
| CREB1 | 1106.531762 | -0.5854744631 | 0.08787590291 | 7.10E-12 | 1.66E-10 |
| VPS37D | 109.1975252 | -0.5856635604 | 0.3011233455 | 0.009500775113 | 0.03378481984 |
| CYP2R1 | 179.2881038 | -0.5875117659 | 0.1474389973 | 1.66E-05 | 0.000133677476 |
| C5orf42 | 1126.492206 | -0.5881208412 | 0.1427022672 | 9.06E-06 | 7.69E-05 |
| INMT | 47.72825448 | -0.5881414766 | 0.3461105269 | 0.0149935582 | 0.04933460508 |
| LOC100129637 | 32.41913056 | -0.5885370335 | 0.3015037533 | 0.009461818372 | 0.03368651692 |

| | baseMean | log2FoldChange | lfcSE | pvalue | padj |
|---|---|---|---|---|---|
| UHRF2 | 1950.172968 | -0.5903475254 | 0.1289614536 | 1.17E-06 | 1.17E-05 |
| ZNF112 | 356.09739 | -0.5919361663 | 0.1271265351 | 7.99E-07 | 8.38E-06 |
| MLLT10 | 908.7665403 | -0.5923975959 | 0.09201001199 | 3.10E-11 | 6.64E-10 |
| POLL | 643.7028278 | -0.5929561599 | 0.1337290823 | 2.27E-06 | 2.15E-05 |
| WRAP73 | 465.5838676 | -0.5937167768 | 0.126721116 | 6.90E-07 | 7.34E-06 |
| AC074286.1 | 44.29863491 | -0.5942425022 | 0.2862579509 | 0.00720611692 | 0.02682967612 |
| LOC389831 | 281.7246734 | -0.595431485 | 0.1311115375 | 1.37E-06 | 1.36E-05 |
| AHNAK | 65342.15547 | -0.5963113247 | 0.1211428881 | 1.99E-07 | 2.37E-06 |
| DNMT3B | 145.528178 | -0.596529268 | 0.1545642043 | 2.69E-05 | 0.000206547995 |
| SYNGAP1 | 93.08939449 | -0.5967643351 | 0.2011876112 | 0.000666016693 | 0.003525176877 |
| TRIM69 | 159.8540443 | -0.5969273198 | 0.1597667188 | 4.36E-05 | 0.000318533353 |
| FBXL4 | 635.7155079 | -0.5971093751 | 0.1036922645 | 2.15E-09 | 3.56E-08 |
| PAPPA2 | 85.58931888 | -0.5973704241 | 0.2341099324 | 0.002212215209 | 0.00998716563 |
| XRRA1 | 368.4908251 | -0.5975976868 | 0.1209418677 | 1.93E-07 | 2.30E-06 |
| ADAL | 162.467284 | -0.599027695 | 0.1666781173 | 7.50E-05 | 0.000519694035 |
| EIF4A2 | 6987.030115 | -0.5996789526 | 0.06620726845 | 3.44E-20 | 1.73E-18 |
| RP11-696N14.1 | 116.5808849 | -0.6002444563 | 0.2593885635 | 0.004019062327 | 0.01658991378 |
| USP44 | 58.25952783 | -0.6004875369 | 0.3034587866 | 0.008415237053 | 0.03051496943 |
| IFIH1 | 57.04355962 | -0.6005077065 | 0.2887688039 | 0.006878339868 | 0.0258158816 |
| CENPI | 839.2660285 | -0.6009842532 | 0.2349353411 | 0.002113903469 | 0.009610353815 |
| HDAC11 | 362.2243448 | -0.6015193042 | 0.207685074 | 0.000804196636 | 0.004169109929 |
| NSUN5P2 | 135.0106432 | -0.6016745613 | 0.1677844602 | 7.73E-05 | 0.000534012745 |
| DOLPP1 | 631.1739392 | -0.6016973447 | 0.1551188156 | 2.43E-05 | 1.88E-04 |
| KIAA1919 | 233.8431211 | -0.6030937792 | 0.1559313372 | 2.55E-05 | 0.000196449146 |
| PGM5P2 | 61.51316191 | -0.60374292 | 0.3532888401 | 0.01378083792 | 0.04598370627 |
| NHLRC3 | 240.3421817 | -0.6038885639 | 0.1651049765 | 5.88E-05 | 0.000417351805 |
| JAKMIP2 | 531.8363346 | -0.6048550146 | 0.2215027291 | 0.001300575849 | 0.006353108206 |
| PAFAH2 | 570.4376887 | -0.605366574 | 0.1026364336 | 9.00E-10 | 1.55E-08 |
| POLG2 | 284.2141562 | -0.6054511793 | 0.1378386611 | 2.65E-06 | 2.49E-05 |
| LOC101060212 | 51.01295408 | -0.605594294 | 0.2824491207 | 0.005924253672 | 0.02285727207 |
| FAM86C2P | 60.38931763 | -0.6055943389 | 0.2237089008 | 0.001409560198 | 0.006805104341 |
| BBC3 | 178.2889694 | -0.6056158227 | 0.2216003116 | 0.001289304466 | 0.006304142851 |
| PRICKLE1 | 85.04613435 | -0.6059153441 | 0.2589470068 | 0.003692113718 | 0.01547631789 |
| D2HGDH | 238.4877995 | -0.6063137005 | 0.1711240379 | 9.72E-05 | 0.000657042346 |
| PAIP2B | 73.28389548 | -0.6076578875 | 0.2461870871 | 0.002697757302 | 0.01183119543 |
| SCYL3 | 334.4694173 | -0.6081449362 | 0.1191706098 | 7.98E-08 | 1.02E-06 |
| ARMCX2 | 1837.143003 | -0.6083554996 | 0.09565052684 | 4.91E-11 | 1.01E-09 |
| NINL | 376.2664721 | -0.6086128425 | 0.183867248 | 0.000204316571 | 0.001252911629 |
| IRF1 | 936.2920292 | -0.6090544947 | 0.1285643787 | 5.11E-07 | 5.59E-06 |
| ZNF552 | 171.9454849 | -0.6091556951 | 0.1747319399 | 0.000108691324 | 0.000726639891 |

| | baseMean | log2FoldChange | lfcSE | pvalue | padj |
|---|---|---|---|---|---|
| RCOR3 | 441.5251182 | -0.6095130551 | 0.1314278052 | 8.29E-07 | 8.64E-06 |
| HEATR5B | 1063.61236 | -0.6103950179 | 0.09897932941 | 1.70E-10 | 3.25E-09 |
| NAIP | 153.7622186 | -0.6121011214 | 0.1487289246 | 9.01E-06 | 7.66E-05 |
| GPR173 | 84.73094661 | -0.6124636829 | 0.1898313692 | 0.000275472822 | 0.001630821018 |
| TRIP11 | 2072.095232 | -0.6136589531 | 0.09678460034 | 5.56E-11 | 1.15E-09 |
| TAF1A | 189.3804271 | -0.6136954366 | 0.1682483204 | 5.90E-05 | 0.000418813140 |
| LOC644656 | 131.95864 | -0.6146130794 | 0.1961756168 | 0.000361924351 | 0.002064632826 |
| ZBTB37 | 199.0974365 | -0.6147045671 | 0.1797181414 | 0.000136890623 | 0.000885004125 |
| METAP1D | 125.2839967 | -0.6150615099 | 0.1541388945 | 1.51E-05 | 0.000123180801 |
| HCG11 | 258.2771363 | -0.6152894812 | 0.1250222046 | 2.02E-07 | 2.40E-06 |
| CTC1 | 171.3128729 | -0.6153110402 | 0.1998946207 | 0.000436413647 | 0.002445526082 |
| SUOX | 474.5280929 | -0.6155031993 | 0.1101848308 | 5.51E-09 | 8.56E-08 |
| CEP250 | 1749.452145 | -0.6159630546 | 0.1451631344 | 4.99E-06 | 4.45E-05 |
| LOC101929112 | 49.69002892 | -0.6164744717 | 0.2604670657 | 0.003371126498 | 0.01434077212 |
| HSDL1 | 736.1059792 | -0.6172553875 | 0.0927983762 | 6.92E-12 | 1.63E-10 |
| CARF | 146.0372697 | -0.6175706302 | 0.2650660548 | 0.003640093187 | 0.01528407994 |
| POLR3B | 866.6235664 | -0.6183068266 | 0.1310549068 | 5.48E-07 | 5.97E-06 |
| SCN9A | 1230.459968 | -0.6183851206 | 0.2557209256 | 0.002925798239 | 0.01268505314 |
| LOC146880 | 59.25071892 | -0.6183863841 | 0.2641029088 | 0.003586786625 | 0.01509857678 |
| IFRD1 | 1490.916799 | -0.6186937216 | 0.1105742811 | 4.96E-09 | 7.77E-08 |
| HDHD3 | 161.4837558 | -0.6188532709 | 0.1528497256 | 1.15E-05 | 9.54E-05 |
| MOSPD2 | 621.7877735 | -0.6191376451 | 0.1065227221 | 1.46E-09 | 2.46E-08 |
| CCDC51 | 548.4626162 | -0.6204857403 | 0.1332087383 | 6.71E-07 | 7.16E-06 |
| LOC101060179 | 88.30153404 | -0.6206029642 | 0.3601267375 | 0.01263111945 | 0.04283539122 |
| MAFG-AS1 | 86.88054854 | -0.6209898115 | 0.2159795152 | 0.000817679389 | 0.004227232092 |
| ZNF860 | 34.00686518 | -0.6210366667 | 0.3076681843 | 0.007390862816 | 0.02738739584 |
| SSH2 | 536.1477264 | -0.6211454929 | 0.1186701089 | 3.84E-08 | 5.21E-07 |
| AC159540.1-2 | 98.56995637 | -0.6233397837 | 0.3562083619 | 0.01195930228 | 0.04101867118 |
| GOLGA6L5P | 68.01900226 | -0.623870487 | 0.2662354809 | 0.003469916861 | 0.01468967647 |
| DCLRE1A | 485.1590204 | -0.6240321649 | 0.2033881115 | 0.000437264908 | 0.00244845326 |
| FAM117A | 90.95919482 | -0.6245743944 | 0.313748039 | 0.007586159099 | 0.02795108212 |
| EPG5 | 2434.087247 | -0.6248892295 | 0.1635117929 | 2.87E-05 | 0.000218866455 |
| NBPF12 | 428.9801522 | -0.624976729 | 0.1612520014 | 2.31E-05 | 0.000180069343 |
| CD27-AS1 | 108.463097 | -0.6255797903 | 0.1929599909 | 0.000248425570 | 0.001489652966 |
| RP5-886K2.3 | 72.74098745 | -0.625661379 | 0.1997487837 | 0.000355944641 | 0.002037547005 |
| HLA-DMB | 22.472773 | -0.6270609639 | 0.3529712391 | 0.01168238908 | 0.04018925394 |
| APOL1 | 68.83966128 | -0.6270926831 | 0.3024664744 | 0.006375471676 | 0.02423395399 |
| EIF3J-AS1 | 149.9360081 | -0.6271319 | 0.1881567198 | 0.000179142089 | 0.001121162914 |
| C11orf45 | 28.54304397 | -0.6281385659 | 0.3221086073 | 0.008302656992 | 0.03015079511 |
| DNAJC27 | 125.9912365 | -0.6319146718 | 0.1710440249 | 4.63E-05 | 0.000335672762 |

| | baseMean | log2FoldChange | lfcSE | pvalue | padj |
|---|---|---|---|---|---|
| KLHL22 | 696.1158226 | -0.6324316267 | 0.1271697774 | 1.46E-07 | 1.79E-06 |
| THAP10 | 64.64697909 | -0.6324408665 | 0.291492325 | 0.005087994379 | 0.02018240473 |
| C5orf45 | 224.1960496 | -0.6329875473 | 0.13283014 | 4.21E-07 | 4.65E-06 |
| FAM86C1 | 258.5207681 | -0.6331256532 | 0.1748892236 | 6.13E-05 | 0.000433314571 |
| C12orf60 | 41.50923912 | -0.6358944966 | 0.326767271 | 0.008071168001 | 0.02943218721 |
| HOXC6 | 75.49723263 | -0.6371912697 | 0.2027700554 | 0.000332577340 | 0.001920024822 |
| LINC00909 | 291.4271678 | -0.6378860231 | 0.117881021 | 1.40E-08 | 2.06E-07 |
| PSRC1 | 1235.397311 | -0.6383500766 | 0.1926584367 | 0.000183878812 | 0.001147431534 |
| CCDC101 | 308.2124528 | -0.6393751218 | 0.1418031193 | 1.39E-06 | 1.38E-05 |
| SLC25A21-AS1 | 64.66696136 | -0.6401137854 | 0.2904204561 | 0.004576724145 | 0.01851204722 |
| ZNF766 | 498.2232703 | -0.6402013563 | 0.1132294719 | 3.48E-09 | 5.57E-08 |
| EDRF1 | 872.1175166 | -0.6402021123 | 0.1306101957 | 2.07E-07 | 2.44E-06 |
| NUDT12 | 426.3734853 | -0.6402374821 | 0.1193603507 | 1.78E-08 | 2.57E-07 |
| TRMT44 | 280.5632747 | -0.6403787317 | 0.1125939927 | 2.88E-09 | 4.68E-08 |
| LOC102606465 | 149.2024423 | -0.640743558 | 0.160530909 | 1.38E-05 | 0.000113458514 |
| AMACR | 301.7679094 | -0.6420609515 | 0.1519671726 | 4.96E-06 | 4.43E-05 |
| SOX13 | 134.4941692 | -0.6425259279 | 0.1920949674 | 0.000164604765 | 0.001041121643 |
| C2CD5 | 825.3279353 | -0.6425709648 | 0.08691255462 | 3.27E-14 | 1.01E-12 |
| RANBP10 | 329.381153 | -0.6426012644 | 0.1127226544 | 2.65E-09 | 4.34E-08 |
| USP21 | 265.481833 | -0.642649678 | 0.1445734331 | 1.85E-06 | 1.79E-05 |
| GBAP1 | 71.54438119 | -0.6435559486 | 0.2005210461 | 2.64E-04 | 1.57E-03 |
| XIST | 3249.380973 | -0.643931158 | 0.196384276 | 0.000203827001 | 0.001251520278 |
| MGC57346 | 187.0901711 | -0.64465814 | 0.1560034129 | 7.72E-06 | 6.65E-05 |
| LOC101929761 | 40.70725215 | -0.6452642022 | 0.2781703658 | 0.003499632916 | 0.01479444477 |
| ARFRP1 | 877.5418113 | -0.6459344839 | 0.1178592039 | 9.30E-09 | 1.40E-07 |
| NBPF11 | 271.2962111 | -0.6460068235 | 0.1401822897 | 8.58E-07 | 8.92E-06 |
| ZNF137P | 26.3983732 | -0.6463426779 | 0.3925492302 | 0.01351320802 | 0.0452232309 |
| NPIPA5 | 201.1996433 | -0.6465460761 | 0.2339850216 | 0.0010478733 | 0.005254895779 |
| LOC102723467 | 93.24128027 | -0.6466155402 | 0.1869623821 | 0.000108617676 | 0.000726508799 |
| MEIS2 | 1356.282962 | -0.6477095336 | 0.1504565031 | 3.18E-06 | 2.94E-05 |
| CLK4 | 345.588741 | -0.648197117 | 0.138605726 | 6.16E-07 | 6.62E-06 |
| NEURL4 | 470.3648702 | -0.6486126207 | 0.1307020753 | 1.48E-07 | 1.80E-06 |
| TMEM136 | 397.0534425 | -0.6504019986 | 0.1016527196 | 3.46E-11 | 7.33E-10 |
| LOC100506621 | 101.7116393 | -0.6508520859 | 0.2407100476 | 0.001236772212 | 0.006085360696 |
| KANSL1L | 222.8659601 | -0.6520713394 | 0.1572964568 | 7.03E-06 | 6.09E-05 |
| DNAH5 | 1210.901755 | -0.6524022891 | 0.1279574321 | 6.95E-08 | 8.97E-07 |
| POLH | 1834.486771 | -0.6529874141 | 0.144910406 | 1.36E-06 | 1.35E-05 |
| HMGCL | 1455.468706 | -0.6529949757 | 0.1053108819 | 1.21E-10 | 2.36E-09 |
| DYRK1B | 155.2360689 | -0.6531583349 | 0.2646838494 | 0.002307045247 | 0.01033476248 |
| PHF21A | 346.3486629 | -0.653229959 | 0.2152802406 | 0.000443658253 | 0.002482385467 |

| | baseMean | log2FoldChange | lfcSE | pvalue | padj |
|---|---|---|---|---|---|
| NTF3 | 77.05641463 | -0.6534545322 | 0.2090455051 | 0.000332973066 | 0.001921564338 |
| HOXA11-AS | 108.1926947 | -0.6540538126 | 0.2522704257 | 0.0016587269 | 0.007835274369 |
| LOC100288842 | 31.08928073 | -0.654711041 | 0.3995137144 | 0.01307767418 | 0.04404286153 |
| SZT2 | 603.4163896 | -0.6548187344 | 0.1017235858 | 2.63E-11 | 5.68E-10 |
| BRCA2 | 631.2194351 | -0.6549955845 | 0.2301636975 | 0.000795800901 | 0.004131338779 |
| CCNL2 | 2374.722519 | -0.655007343 | 0.1376108447 | 4.00E-07 | 4.46E-06 |
| FAM217B | 268.1626564 | -0.6552392667 | 0.1406327088 | 6.50E-07 | 6.95E-06 |
| SOCS1 | 41.80947778 | -0.6562617202 | 0.3361842588 | 0.007498549113 | 0.0276899548 |
| ADAM1A | 56.84509275 | -0.6563394073 | 0.2826698488 | 0.003343949978 | 0.01423736666 |
| NSUN6 | 294.7332402 | -0.6564273595 | 0.1307318522 | 1.08E-07 | 1.36E-06 |
| LRRCC1 | 208.439512 | -0.6589012098 | 0.1433247849 | 8.83E-07 | 9.14E-06 |
| STEAP2 | 84.09349643 | -0.6592286894 | 0.2432294395 | 0.001193165669 | 0.005902007853 |
| RFESD | 41.77800508 | -0.6598509083 | 0.3111351198 | 0.005240651439 | 0.02065089498 |
| MXRA5 | 213.3660374 | -0.6603742405 | 0.3324536775 | 0.006811468965 | 0.02560362571 |
| LOC389765 | 24.7989401 | -0.6605307927 | 0.3537376447 | 0.00879884291 | 0.0316975495 |
| FANCF | 323.3239415 | -0.6610957706 | 0.1037490531 | 3.99E-11 | 8.37E-10 |
| CCDC93 | 1102.78834 | -0.6611660773 | 0.1092897643 | 3.09E-10 | 5.62E-09 |
| MLH3 | 1011.187152 | -0.6631482275 | 0.09414498232 | 4.00E-13 | 1.10E-11 |
| WDR31 | 59.60538207 | -0.663467694 | 0.2308449089 | 0.000730335381 | 0.003827512669 |
| LOC101929500 | 148.0261338 | -0.6641913854 | 0.3126992324 | 0.005063914582 | 0.02010040635 |
| FAN1 | 977.4012255 | -0.6642466055 | 0.09979822744 | 6.00E-12 | 1.42E-10 |
| GCNT1 | 974.3259741 | -0.6646728588 | 0.1924718521 | 1.04E-04 | 0.000697309662 |
| DHRS4-AS1 | 313.512065 | -0.6649018736 | 0.1633299118 | 9.20E-06 | 7.81E-05 |
| FRMD6-AS1 | 59.98885802 | -0.6651964009 | 0.2461584546 | 0.001198553496 | 0.005922755726 |
| EFNA4 | 190.3184084 | -0.665316454 | 0.1669878804 | 1.31E-05 | 0.000107960619 |
| PRPF40B | 317.4482711 | -0.6664398341 | 0.1078584388 | 1.37E-10 | 2.65E-09 |
| LINC00476 | 127.2489131 | -0.6670138423 | 0.1946435073 | 0.000115013971 | 0.000763801524 |
| TIGD2 | 271.1011542 | -0.6670673951 | 0.1477958116 | 1.28E-06 | 1.28E-05 |
| C10orf25 | 63.0617422 | -0.6678480776 | 0.2474596572 | 0.001206618532 | 0.005958654503 |
| ZFP28 | 358.088256 | -0.6682523003 | 0.1259004146 | 2.29E-08 | 3.23E-07 |
| ZBTB25 | 156.9201448 | -0.6690347124 | 0.1926814251 | 9.79E-05 | 0.000661367662 |
| CDKN2C | 452.7177405 | -0.6691714692 | 0.1647050171 | 9.36E-06 | 7.93E-05 |
| TRERF1 | 499.3309675 | -0.6699266167 | 0.110080689 | 2.43E-10 | 4.52E-09 |
| MYOZ3 | 16.98998295 | -0.6700535796 | 0.4320464808 | 0.01456987614 | 0.04820686353 |
| TRIM59 | 859.9908793 | -0.6723707389 | 0.2067866346 | 2.01E-04 | 0.001240096145 |
| SENP7 | 523.4222165 | -0.6724386069 | 0.1413589684 | 3.96E-07 | 4.41E-06 |
| CLHC1 | 63.25031463 | -0.6732244565 | 0.2137352853 | 2.93E-04 | 1.72E-03 |
| MRVI1 | 824.6780333 | -0.6745019885 | 0.3380726113 | 0.006394508267 | 0.02430011067 |
| ENO2 | 1466.55212 | -0.6766880183 | 0.1913664265 | 7.43E-05 | 0.000515703349 |
| LOC101927045 | 39.131709 | -0.6775139945 | 0.2812094817 | 0.002537526804 | 0.01122516055 |

| | baseMean | log2FoldChange | lfcSE | pvalue | padj |
|---|---|---|---|---|---|
| HOXB3 | 311.8013405 | -0.677928303 | 0.1217467212 | 5.22E-09 | 8.17E-08 |
| ZFP82 | 178.1829483 | -0.6779790566 | 0.1838989293 | 4.23E-05 | 0.000309240008 |
| ZNF184 | 427.2579066 | -0.6800241921 | 0.1209915802 | 3.82E-09 | 6.08E-08 |
| MORN4 | 299.4465225 | -0.6800739145 | 0.1897449816 | 6.19E-05 | 0.000437050393 |
| CREBZF | 807.893319 | -0.6824788904 | 0.1549694971 | 2.02E-06 | 1.94E-05 |
| LINC00346 | 61.29130598 | -0.6838576192 | 0.2255440808 | 0.000418272181 | 0.002353610926 |
| CLCN6 | 377.2749153 | -0.6848504213 | 0.1273523494 | 1.49E-08 | 2.17E-07 |
| LYSMD4 | 202.018751 | -0.6854855796 | 0.1392599695 | 1.66E-07 | 2.01E-06 |
| KMT2A | 1184.44561 | -0.6860924475 | 0.1381391048 | 1.32E-07 | 1.63E-06 |
| MAVS | 2595.33228 | -0.6868460092 | 0.0660517686 | 5.20E-26 | 4.21E-24 |
| TRIM6 | 310.5218279 | -0.6872445816 | 0.2235221852 | 0.000356168668 | 0.002038045852 |
| DNAJB2 | 896.4093895 | -0.6874821725 | 0.1515153907 | 1.07E-06 | 1.09E-05 |
| PITX2 | 244.9596424 | -0.6876876124 | 0.2295049657 | 0.000455504206 | 0.002532487725 |
| KGFLP1 | 45.51934194 | -0.6879910912 | 0.3122096124 | 0.003977793815 | 0.01644235761 |
| LRIF1 | 657.3959059 | -0.6888777701 | 0.1030019785 | 4.46E-12 | 1.07E-10 |
| DCP1B | 746.2979437 | -0.6901892843 | 0.1147023299 | 3.32E-10 | 6.02E-09 |
| ENGASE | 326.4345525 | -0.6903417712 | 0.1416272587 | 2.08E-07 | 2.44E-06 |
| KCNQ3 | 57.49765259 | -0.6913845434 | 0.2987168036 | 0.003057648869 | 0.01316141486 |
| CLSTN2 | 31.58573858 | -0.6920085615 | 0.331031972 | 0.005165324639 | 0.02043755476 |
| RNF214 | 543.6519802 | -0.6939237322 | 0.0963184274 | 1.16E-13 | 3.36E-12 |
| CNTNAP1 | 608.4556822 | -0.6944796945 | 0.2124901722 | 0.000184323104 | 0.001149240662 |
| LOC101927451 | 25.30273813 | -0.694647587 | 0.3673388131 | 0.007682266213 | 0.02824228683 |
| ZNF280D | 1112.548364 | -0.6958890585 | 0.07678588472 | 2.55E-20 | 1.30E-18 |
| MAP3K12 | 823.8977987 | -0.6960845793 | 0.155505386 | 1.40E-06 | 1.39E-05 |
| ZMYM1 | 463.2781345 | -0.6961289823 | 0.1567850095 | 1.66E-06 | 1.63E-05 |
| AKNA | 422.3939761 | -0.696516881 | 0.1926121185 | 5.23E-05 | 0.000374905324 |
| ZNF491 | 63.35473859 | -0.6965277664 | 0.2333371853 | 0.000472645235 | 0.002616065024 |
| BORA | 482.1938929 | -0.6969098271 | 0.2038203951 | 0.000107022320 | 0.000716804014 |
| FAM120C | 264.2100256 | -0.6975042143 | 0.1119831082 | 9.33E-11 | 1.86E-09 |
| ZNF436 | 723.1188384 | -0.6986905468 | 0.1382873131 | 8.22E-08 | 1.05E-06 |
| CCDC66 | 338.6017956 | -0.6988608009 | 0.1254033666 | 4.83E-09 | 7.61E-08 |
| GORAB | 300.3966644 | -0.7030764819 | 0.141170757 | 1.17E-07 | 1.46E-06 |
| MSL3P1 | 253.5436753 | -0.7033185166 | 0.1608264783 | 2.21E-06 | 2.10E-05 |
| LOC101930357 | 37.86762175 | -0.7037969584 | 0.3967121137 | 0.009019309435 | 0.03238238798 |
| CELSR2 | 29.17870306 | -0.7067960501 | 0.3843016391 | 0.008063200787 | 0.02941686447 |
| ARHGEF2 | 3896.321635 | -0.7072328611 | 0.07200920427 | 1.79E-23 | 1.17E-21 |
| MRM1 | 213.0845136 | -0.707437508 | 0.1461302326 | 2.36E-07 | 2.74E-06 |
| TRANK1 | 609.4733854 | -0.7075259702 | 0.2973717719 | 0.002446497443 | 0.01088965633 |
| ESPNL | 86.54237523 | -0.7076741767 | 0.3007210508 | 0.002624892722 | 0.01155589229 |
| C2orf44 | 858.3802738 | -0.707719155 | 0.1170965139 | 2.85E-10 | 5.22E-09 |

| | baseMean | log2FoldChange | lfcSE | pvalue | padj |
|---|---|---|---|---|---|
| ANKS1A | 525.8707503 | -0.7089099506 | 0.1221475377 | 1.22E-09 | 2.07E-08 |
| AGAP7 | 41.68695745 | -0.7091678324 | 0.3588317412 | 0.006155735646 | 0.02359144094 |
| NBPF25P | 24.48544418 | -0.7100597172 | 0.3522379005 | 0.005801747292 | 0.02246027442 |
| ZFP41 | 194.2240634 | -0.7109049699 | 0.1478626584 | 2.78E-07 | 3.19E-06 |
| DKFZP434I0714 | 126.3680021 | -0.7129119866 | 0.1847676288 | 1.98E-05 | 0.000157586047 |
| FRS3 | 92.36631889 | -0.7134563651 | 0.2270308574 | 0.000268884572 | 0.001596260923 |
| FBN2 | 5620.407739 | -0.7136747452 | 0.182105113 | 1.52E-05 | 0.000123654652 |
| ZNF30 | 124.9869564 | -0.7146385225 | 0.1554917674 | 7.89E-07 | 8.28E-06 |
| ANKRD23 | 30.6552443 | -0.7156400331 | 0.3176940908 | 0.003336558233 | 0.0142180926 |
| RAB33A | 78.66797052 | -0.7170199731 | 0.2483161352 | 0.000597175154 | 0.003205241843 |
| ZNF442 | 46.6394077 | -0.7183542354 | 0.2825999403 | 0.001615717816 | 0.007653968362 |
| ANO8 | 66.12321131 | -0.7189805496 | 0.2165048969 | 0.000146976156 | 0.000941621341 |
| KAT6B | 999.6224671 | -0.7194670092 | 0.1188228682 | 2.54E-10 | 4.72E-09 |
| TRIM56 | 347.2145952 | -0.7195485854 | 0.1437571872 | 1.01E-07 | 1.28E-06 |
| LOC101928466 | 14.60340193 | -0.7196405974 | 0.4599579381 | 0.0123147218 | 0.04195741257 |
| ZBTB3 | 254.8945413 | -0.7196925663 | 0.1366825823 | 2.54E-08 | 3.55E-07 |
| LINC00342 | 144.9554625 | -0.7201708467 | 0.2076434042 | 8.58E-05 | 0.000586174555 |
| GOLGA8M | 31.55839945 | -0.7204397142 | 0.3517102821 | 0.005209792066 | 0.02056067569 |
| RP5-1050D4.5 | 20.54483751 | -0.7210310526 | 0.450262263 | 0.01138989567 | 0.03940152338 |
| CCDC130 | 491.8666396 | -0.721246222 | 0.1449843956 | 1.15E-07 | 1.44E-06 |
| KBTBD6 | 711.6436772 | -0.7214511855 | 0.1282349095 | 3.32E-09 | 5.33E-08 |
| ZFHX4 | 279.3163346 | -0.7221418718 | 0.1405939689 | 5.04E-08 | 6.72E-07 |
| LOC102724872 | 25.76345241 | -0.7226165098 | 0.4679090331 | 0.01227690715 | 0.04190407043 |
| RP11-890B15.2 | 88.84644125 | -0.7236687253 | 0.2625576266 | 0.000864791928 | 0.004435372728 |
| LOC646903 | 48.74280485 | -0.7277367138 | 0.2679248941 | 0.000978482804 | 0.004950265195 |
| ARVCF | 119.7350037 | -0.7284514761 | 0.2101329903 | 8.44E-05 | 0.00057755926 |
| KIAA1109 | 2909.263467 | -0.728473054 | 0.1505728068 | 2.28E-07 | 2.66E-06 |
| ZNF292 | 1442.259971 | -0.728522826 | 0.1073094574 | 2.07E-12 | 5.19E-11 |
| C5orf34 | 246.0230138 | -0.7287607923 | 0.2623922396 | 0.000792805649 | 0.004117233966 |
| C4orf29 | 219.3876036 | -0.7288619075 | 0.1325351823 | 6.74E-09 | 1.04E-07 |
| ZCCHC4 | 346.4765755 | -0.729579875 | 0.1605905681 | 9.40E-07 | 9.67E-06 |
| ZNF182 | 204.7603981 | -0.7300535532 | 0.1373724341 | 1.91E-08 | 2.74E-07 |
| ZNF623 | 800.3602228 | -0.7304430007 | 0.08552221869 | 2.47E-18 | 1.10E-16 |
| CIDEB | 34.04611107 | -0.731152959 | 0.299865114 | 0.002035993943 | 0.009327358097 |
| CCDC163P | 36.31589097 | -0.7318983819 | 0.2937217046 | 0.001802645717 | 0.008392617909 |
| CCNF | 1185.345799 | -0.7319225472 | 0.2036002379 | 5.17E-05 | 0.000371093978 |
| FAM63A | 126.4125681 | -0.7320407488 | 0.2170718018 | 0.000117299718 | 0.000776901916 |
| MXD3 | 298.91787 | -0.7321397824 | 0.201378982 | 4.39E-05 | 0.000320431004 |
| RMI1 | 299.1163691 | -0.7321685244 | 0.1702458997 | 2.86E-06 | 2.68E-05 |
| RBM4B | 486.9314538 | -0.7335374684 | 0.1005075374 | 5.23E-14 | 1.58E-12 |

| | baseMean | log2FoldChange | lfcSE | pvalue | padj |
|---|---|---|---|---|---|
| MIEF2 | 242.3070333 | -0.7338360602 | 0.1720894726 | 3.37E-06 | 3.09E-05 |
| ZNF441 | 279.2945653 | -0.7343303513 | 0.1152682612 | 3.34E-11 | 7.11E-10 |
| SYNJ1 | 168.7381059 | -0.7352154202 | 0.1374500232 | 1.60E-08 | 2.32E-07 |
| L3MBTL1 | 52.3181947 | -0.735851405 | 0.2532375517 | 0.000549922216 | 0.002986065604 |
| ZNF18 | 343.0867786 | -0.7362606476 | 0.1434161114 | 4.89E-08 | 6.53E-07 |
| ZNF252P | 725.1935291 | -0.7366610938 | 0.1068577214 | 9.69E-13 | 2.52E-11 |
| ZNF512 | 1135.271215 | -0.7384550943 | 0.06865160716 | 1.02E-27 | 9.55E-26 |
| PGBD4 | 81.91549396 | -0.740323646 | 0.2494291271 | 0.000435503797 | 0.002442265926 |
| C3orf18 | 299.3416315 | -0.7414729217 | 0.1888229989 | 1.39E-05 | 0.000113618935 |
| WNK3 | 78.06531446 | -0.7422897338 | 0.2089091504 | 6.05E-05 | 0.000428235248 |
| LOC102724814 | 65.78576039 | -0.7427195529 | 0.2891528481 | 0.001376284036 | 0.00667692832 |
| ZNF280C | 277.017573 | -0.7432222426 | 0.1447441519 | 4.80E-08 | 6.43E-07 |
| MAPK7 | 300.3446031 | -0.7433242816 | 0.1453109772 | 5.31E-08 | 7.03E-07 |
| AGAP9 | 107.7598698 | -0.7438084496 | 0.1632095693 | 8.82E-07 | 9.14E-06 |
| ZNF14 | 178.6111717 | -0.7444516867 | 0.1458569818 | 5.77E-08 | 7.60E-07 |
| ZNF616 | 484.2520313 | -0.7446351606 | 0.134675785 | 5.47E-09 | 8.51E-08 |
| LOC100506730 | 49.46625614 | -0.7461115341 | 0.2992014006 | 0.001695302674 | 0.007972634718 |
| C11orf68 | 1432.838631 | -0.7464093827 | 0.1312267017 | 2.19E-09 | 3.63E-08 |
| KIAA1731 | 1189.253668 | -0.7467398478 | 0.1390931921 | 1.34E-08 | 1.97E-07 |
| PLD1 | 772.0480809 | -0.7467981855 | 0.1779260849 | 4.37E-06 | 3.94E-05 |
| THAP2 | 119.799658 | -0.7482969268 | 0.1661419203 | 1.12E-06 | 1.13E-05 |
| DPY19L2 | 82.80990786 | -0.7489206754 | 0.2498539734 | 0.000389737688 | 0.002203875596 |
| BRWD3 | 632.7882204 | -0.7498053713 | 0.1449757268 | 3.87E-08 | 5.24E-07 |
| LOC101927903 | 89.33937574 | -0.7502819102 | 0.2445370681 | 0.000312522172 | 0.001819766379 |
| DUSP19 | 78.80150798 | -0.7504297409 | 0.2078572458 | 4.79E-05 | 0.000345221753 |
| PABPC1L | 95.92192294 | -0.7530107027 | 0.1912091319 | 1.32E-05 | 0.000108540279 |
| ZNF239 | 282.8681815 | -0.7531890652 | 0.1912861 | 1.29E-05 | 1.07E-04 |
| NLRX1 | 339.7693821 | -0.753726149 | 0.146827466 | 4.70E-08 | 6.30E-07 |
| LYG1 | 11.39018552 | -0.7537583563 | 0.5511139978 | 0.01421840753 | 0.04725398879 |
| SWT1 | 130.6740062 | -0.754640462 | 0.23223387 | 0.000169362425 | 0.00106803776 |
| MSANTD2 | 412.8012151 | -0.7550405891 | 0.1498193824 | 7.48E-08 | 9.62E-07 |
| KCNE1 | 31.26333021 | -0.7556381483 | 0.3921806179 | 0.005917904648 | 0.02283869422 |
| TENC1 | 997.7748869 | -0.7568604658 | 0.2204093869 | 8.79E-05 | 0.000599558505 |
| ZNF785 | 153.8969257 | -0.7586292866 | 0.2291394691 | 0.000137253591 | 0.000886580791 |
| CNTNAP3 | 283.161805 | -0.7590628196 | 0.2288804969 | 0.000132818352 | 0.000863811060 |
| ZNF577 | 131.0351783 | -0.7605794455 | 0.1716266188 | 1.49E-06 | 1.47E-05 |
| ZNF510 | 488.4709392 | -0.7612990004 | 0.1112317956 | 1.29E-12 | 3.31E-11 |
| LBX2-AS1 | 26.47312984 | -0.7614130933 | 0.4149549685 | 0.00686589874 | 0.02577568491 |
| ZNF446 | 192.8443321 | -0.7617591852 | 0.2367584351 | 0.000185088210 | 0.001153527989 |
| C1RL-AS1 | 111.5204633 | -0.7635689374 | 0.2196992055 | 7.53E-05 | 0.000521675385 |

| | baseMean | log2FoldChange | lfcSE | pvalue | padj |
|---|---|---|---|---|---|
| HERC2P9 | 162.7221337 | -0.7655347575 | 0.1556994827 | 1.44E-07 | 1.76E-06 |
| SCAND3 | 35.23402361 | -0.7668405451 | 0.3380018361 | 0.002783856329 | 0.01215864972 |
| POLM | 171.9280035 | -0.7672002814 | 0.1608197926 | 2.97E-07 | 3.40E-06 |
| MAP3K1 | 1316.588592 | -0.7680430439 | 0.1142837036 | 2.85E-12 | 6.96E-11 |
| TSPY26P | 63.43952862 | -0.7698031706 | 0.2954139573 | 0.001181375918 | 0.0058495198 |
| LRRC27 | 217.2412903 | -0.769938442 | 0.1801975355 | 2.99E-06 | 2.78E-05 |
| METTL20 | 67.9019665 | -0.770456903 | 0.3008914503 | 0.001308484149 | 0.006383361894 |
| SLC25A27 | 92.5823858 | -0.7713018918 | 0.4208450793 | 0.006613791772 | 0.02502484008 |
| LOC101060195 | 42.50186466 | -0.7716330517 | 0.3030784182 | 0.001400169933 | 0.006767891031 |
| AHNAK2 | 2663.974057 | -0.7721479948 | 0.07909250769 | 2.77E-23 | 1.79E-21 |
| LOC101928192 | 41.88982109 | -0.7723702093 | 0.2859024864 | 0.000916645080 | 0.004669151079 |
| RNASEL | 325.5661288 | -0.772884131 | 0.1772868664 | 2.00E-06 | 1.92E-05 |
| RUSC1-AS1 | 20.63391073 | -0.773762168 | 0.5475149485 | 0.01226189499 | 0.0418636447 |
| HARBI1 | 121.1720243 | -0.7741633211 | 0.2130884699 | 4.14E-05 | 0.000304162803 |
| INTU | 310.2396755 | -0.7745471861 | 0.115825947 | 3.80E-12 | 9.20E-11 |
| HTATSF1P2 | 169.1094196 | -0.7748708395 | 0.1355556119 | 1.81E-09 | 3.02E-08 |
| ZNF70 | 226.3542592 | -0.7762493565 | 0.135414151 | 1.60E-09 | 2.68E-08 |
| HOXA9 | 296.25923 | -0.7766616406 | 0.1384570758 | 3.24E-09 | 5.21E-08 |
| FZD2 | 837.2241144 | -0.7767896912 | 0.117308709 | 6.38E-12 | 1.50E-10 |
| LOC100507642 | 25.13100634 | -0.7779423938 | 0.4698171641 | 0.008923848986 | 0.03210903517 |
| TTC30A | 344.9467601 | -0.7784018241 | 0.1433079632 | 8.90E-09 | 1.35E-07 |
| CARD9 | 25.07763721 | -0.7789179922 | 0.4339499999 | 0.007067568362 | 0.02639965513 |
| SDPR | 2226.966313 | -0.7795436535 | 0.1065469691 | 4.03E-14 | 1.23E-12 |
| SETDB2 | 427.5939422 | -0.7800685834 | 0.1382671629 | 2.70E-09 | 4.40E-08 |
| EIF1B-AS1 | 14.41042314 | -0.7805847163 | 0.671881666 | 0.01521263233 | 0.04994506785 |
| ZNF337 | 703.2220625 | -0.782121817 | 0.1115557197 | 3.85E-13 | 1.06E-11 |
| BBS10 | 421.634987 | -0.7822130539 | 0.1347860637 | 1.03E-09 | 1.78E-08 |
| STX18-AS1 | 19.48321982 | -0.7822745882 | 0.4370500591 | 0.007269560524 | 0.02699837532 |
| LOC100288069 | 70.99475941 | -0.7830724566 | 0.2424393164 | 0.000168853847 | 0.001065733333 |
| LOC101929264 | 28.51243198 | -0.7833344548 | 0.3382401544 | 0.002474029185 | 0.01099905062 |
| ADM2 | 151.8454603 | -0.783357357 | 0.236612266 | 0.000128545102 | 0.000840539318 |
| CEL | 12.29202567 | -0.7833611844 | 0.5944858636 | 0.01390225185 | 0.04631698987 |
| ZNF485 | 121.8810384 | -0.7847183672 | 0.2347559208 | 0.000115239627 | 0.000764958901 |
| TOB2P1 | 28.38647952 | -0.7847771399 | 0.4919863998 | 0.009374972542 | 0.03344930893 |
| ZNF470 | 239.3793121 | -0.7855620956 | 0.2051337337 | 1.87E-05 | 0.000148951527 |
| RNF34 | 1134.091942 | -0.7863467099 | 0.1045648107 | 8.81E-15 | 2.85E-13 |
| PPARA | 477.1330096 | -0.786764612 | 0.1771414285 | 1.34E-06 | 1.34E-05 |
| BRD8 | 1676.77843 | -0.7880022662 | 0.08167599541 | 8.12E-23 | 5.01E-21 |
| KNOP1 | 847.8357796 | -0.7880799622 | 0.1404148416 | 3.07E-09 | 4.96E-08 |
| HNRNPU-AS1 | 231.3903609 | -0.7888625088 | 0.2759223615 | 0.000539571004 | 0.002936283878 |

| | baseMean | log2FoldChange | lfcSE | pvalue | padj |
|---|---|---|---|---|---|
| AJUBA | 1763.510048 | -0.7890188128 | 0.07425619719 | 3.72E-27 | 3.32E-25 |
| GTSE1-AS1 | 12.32930093 | -0.7891977796 | 0.6180589817 | 0.01399041542 | 0.04658986696 |
| RP11-449D8.1 | 28.96777032 | -0.7901506706 | 0.6976961514 | 0.01462696072 | 0.04834202401 |
| AGFG2 | 247.6388902 | -0.7919809063 | 0.1390224651 | 1.88E-09 | 3.13E-08 |
| ZNF621 | 691.1888538 | -0.7920286285 | 0.09787613747 | 9.56E-17 | 3.60E-15 |
| RP11-307E17.8 | 35.4432767 | -0.7958873895 | 0.3207225302 | 0.001566646304 | 0.007466644308 |
| KIAA1656 | 50.94779518 | -0.7962141359 | 0.2439975886 | 0.000151846109 | 0.000968653264 |
| HSPA1A | 2389.165506 | -0.7974368257 | 0.1109017676 | 1.03E-13 | 3.00E-12 |
| SLC35E2B | 1211.650034 | -0.7980350756 | 0.1291469823 | 8.66E-11 | 1.74E-09 |
| FAM161A | 150.3714572 | -0.7980612941 | 0.2669211117 | 0.000354857396 | 0.002032104532 |
| BRICD5 | 59.94694135 | -0.7987136922 | 0.3131159776 | 0.001267352922 | 0.006215289081 |
| KCTD21 | 505.1620817 | -0.798883636 | 0.09830732858 | 6.94E-17 | 2.68E-15 |
| PRUNE | 232.7387262 | -0.7998897169 | 0.1657541876 | 2.04E-07 | 2.41E-06 |
| LOC101928560 | 27.62762318 | -0.8007398252 | 0.3601928431 | 0.002867895024 | 0.01248175651 |
| C12orf65 | 441.2015144 | -0.8007824476 | 0.1165859248 | 9.94E-13 | 2.57E-11 |
| AASDH | 521.6770867 | -0.8026561891 | 0.1029785758 | 1.01E-15 | 3.55E-14 |
| ZNF563 | 75.18928729 | -0.8028622864 | 0.2400556502 | 0.000109743797 | 0.000732582196 |
| MIAT | 61.86225321 | -0.8032494356 | 0.2372916676 | 9.70E-05 | 0.000655875548 |
| OXSM | 290.9806758 | -0.8050576061 | 0.1518325719 | 1.71E-08 | 2.47E-07 |
| RALGAPA1P | 128.3435935 | -0.8054645525 | 0.1809168611 | 1.26E-06 | 1.26E-05 |
| SPEG | 232.5420996 | -0.8057924708 | 0.2549107242 | 0.000202926257 | 0.001247983911 |
| TUT1 | 436.6356976 | -0.8059930104 | 0.1195296663 | 2.37E-12 | 5.88E-11 |
| KIAA1377 | 70.58654693 | -0.806782805 | 0.2546271118 | 0.000200768401 | 0.001236756609 |
| PRDM15 | 324.0923495 | -0.8071207638 | 0.1190924322 | 1.94E-12 | 4.88E-11 |
| NEAT1 | 1955.266726 | -0.8079024863 | 0.2218699499 | 3.63E-05 | 2.70E-04 |
| ASB8 | 626.5636668 | -0.8079180081 | 0.1006793588 | 1.58E-16 | 5.88E-15 |
| PYGO2 | 658.3402692 | -0.8100827378 | 0.09263646196 | 3.46E-19 | 1.63E-17 |
| TMEM92 | 9.612075622 | -0.8105206939 | 0.6141539578 | 0.01321567293 | 0.04439716281 |
| CCDC144B | 203.4024358 | -0.8110021873 | 0.1727179555 | 3.86E-07 | 4.32E-06 |
| ZNF839 | 310.7522949 | -0.8114573332 | 0.1203102461 | 2.34E-12 | 5.80E-11 |
| ZNF382 | 127.3967076 | -0.8122794647 | 0.1650825547 | 1.27E-07 | 1.58E-06 |
| INS-IGF2 | 53.79425048 | -0.8142606514 | 0.3293001914 | 0.001490941301 | 0.00715161889 |
| MGA | 2007.443086 | -0.8143779204 | 0.1342957236 | 1.99E-10 | 3.74E-09 |
| NPHP3-ACAD11 | 202.8340498 | -0.8146917466 | 0.2969557526 | 0.000708437431 | 0.00372586539 |
| PPIEL | 39.76375339 | -0.8148021898 | 0.369655578 | 0.002902164564 | 0.01260879142 |
| PRORSD1P | 10.40154693 | -0.8166478902 | 0.6651958069 | 0.01382353481 | 0.04607535478 |
| NMNAT1 | 431.2662718 | -0.817354037 | 0.1053991495 | 1.34E-15 | 4.63E-14 |
| LINC00883 | 501.0434344 | -0.817418901 | 0.1723566627 | 3.00E-07 | 3.43E-06 |
| LOC101927382 | 51.2416232 | -0.8183553603 | 0.4708134349 | 0.006944782205 | 0.0260073797 |
| DNAJB1 | 3240.647291 | -0.8185085477 | 0.08188996489 | 2.50E-24 | 1.84E-22 |

| | baseMean | log2FoldChange | lfcSE | pvalue | padj |
|---|---|---|---|---|---|
| MMAA | 181.5082378 | -0.8188913836 | 0.1449290783 | 2.38E-09 | 3.92E-08 |
| HYMAI | 21.94837625 | -0.8204122939 | 0.4322429349 | 0.005434399049 | 0.02124810069 |
| ZNF192P1 | 17.15105377 | -0.8204938189 | 0.4704652475 | 0.007227370937 | 0.02688863715 |
| ZNF525 | 175.6060361 | -0.8224244512 | 0.1325085315 | 8.11E-11 | 1.63E-09 |
| MTL5 | 13.30238247 | -0.8225763296 | 0.5872487671 | 0.01135590764 | 0.03930221035 |
| RIC8B | 551.6339003 | -0.8227836221 | 0.09577516955 | 1.37E-18 | 6.23E-17 |
| ZBTB12 | 111.1311533 | -0.8248149857 | 0.2053963264 | 8.08E-06 | 6.93E-05 |
| TTLL11 | 209.9373991 | -0.824944668 | 0.1958214238 | 3.43E-06 | 3.14E-05 |
| LOC102724187 | 47.34735857 | -0.8257366998 | 0.2847644077 | 0.000453146390 | 0.002524156947 |
| KIF27 | 117.6265911 | -0.826506431 | 0.2229795989 | 2.72E-05 | 0.000208589223 |
| REV3L | 1662.356879 | -0.8268225207 | 0.1722682623 | 2.22E-07 | 2.60E-06 |
| GOLGA8B | 469.4688219 | -0.8269648903 | 0.2279749056 | 3.64E-05 | 0.000270316875 |
| LOC101928489 | 15.14415235 | -0.8283418703 | 0.5242599811 | 0.009056506592 | 0.03249995822 |
| FAM27E3 | 49.62257503 | -0.8298076703 | 0.3831775362 | 0.002960514822 | 0.01282114752 |
| MEG9 | 53.29420126 | -0.8300272167 | 0.3553086269 | 0.002039864933 | 0.009336473713 |
| PARS2 | 197.9888723 | -0.8300456962 | 0.1596483105 | 2.88E-08 | 3.97E-07 |
| FAM220A | 434.5994374 | -0.8301373089 | 0.09771753412 | 2.97E-18 | 1.30E-16 |
| ZNF596 | 125.5238917 | -0.8306336557 | 0.1979884213 | 3.72E-06 | 3.40E-05 |
| C17orf100 | 32.19782873 | -0.8325300664 | 0.3444296421 | 0.001717756866 | 0.008057870816 |
| MAGI3 | 418.4978244 | -0.8331482318 | 0.1247634349 | 3.61E-12 | 8.74E-11 |
| ZNF691 | 208.6560761 | -0.8332224191 | 0.1594186738 | 2.42E-08 | 3.39E-07 |
| ZNF852 | 49.06049748 | -0.8336113382 | 0.2926794218 | 0.000508786615 | 0.002789147244 |
| KLF15 | 28.08246727 | -0.8340727307 | 0.5683211265 | 0.009782185754 | 0.03463661443 |
| SEPSECS | 224.7891826 | -0.8364039723 | 0.1548448101 | 9.58E-09 | 1.44E-07 |
| PLA2G4B | 50.60860298 | -0.8389202213 | 0.3733701011 | 0.002511994425 | 0.01112887122 |
| USP37 | 702.958041 | -0.8394758145 | 0.09971967555 | 5.63E-18 | 2.41E-16 |
| CISH | 55.93703081 | -0.840005595 | 0.4115780703 | 0.003822613374 | 0.0159202491 |
| LOC153684 | 50.51972896 | -0.8404011067 | 0.2588547345 | 0.000144846950 | 0.000929179770 |
| SLC35E2 | 733.0108039 | -0.8408696863 | 0.1400549401 | 2.75E-10 | 5.08E-09 |
| ZNF514 | 343.0412138 | -0.8409627632 | 0.1462485024 | 1.27E-09 | 2.16E-08 |
| AC012309.5 | 20.17106368 | -0.8424044542 | 0.4118321905 | 0.003940953699 | 0.01633088216 |
| USP49 | 136.393435 | -0.8430364507 | 0.1637475846 | 3.67E-08 | 4.99E-07 |
| RASSF7 | 117.9650566 | -0.8433815218 | 0.3164627863 | 0.000834043635 | 0.004298248082 |
| GEN1 | 526.9448794 | -0.8437809452 | 0.2253438307 | 2.25E-05 | 0.000175873374 |
| RNF207 | 29.0092766 | -0.8449708365 | 0.3496302661 | 0.001653367337 | 0.007812436141 |
| PRCD | 24.60674738 | -0.8477859322 | 0.4014851443 | 0.003342944485 | 0.01423715687 |
| IL21R | 123.0621331 | -0.8478216712 | 0.2148857629 | 1.02E-05 | 8.59E-05 |
| LOC101060351 | 37.03347363 | -0.848017262 | 0.4525650956 | 0.005207340787 | 0.02056008936 |
| NHLRC1 | 39.4765687 | -0.8481901662 | 0.3375952193 | 0.001288133271 | 0.006302667192 |
| STC2 | 3274.160063 | -0.8483746458 | 0.08849930557 | 1.32E-22 | 8.08E-21 |

| | baseMean | log2FoldChange | lfcSE | pvalue | padj |
|---|---|---|---|---|---|
| RP11-480I12.7 | 8.479755817 | -0.8492919997 | 0.750503549 | 0.01400849169 | 0.04663963164 |
| HOXA11 | 648.8957669 | -0.8500383834 | 0.1885771773 | 8.60E-07 | 8.94E-06 |
| STARD9 | 721.8789302 | -0.8501236145 | 0.1396534486 | 1.59E-10 | 3.06E-09 |
| LPIN1 | 2717.22331 | -0.8511535039 | 0.0604359767 | 6.91E-46 | 1.49E-43 |
| COLQ | 11.99855772 | -0.852054508 | 0.742776226 | 0.01304459755 | 0.04394140563 |
| MTERFD3 | 187.9850202 | -0.8523348799 | 0.1384765884 | 1.05E-10 | 2.07E-09 |
| ZNF490 | 197.5770783 | -0.8532225465 | 0.1580902099 | 9.27E-09 | 1.40E-07 |
| LOC100270804 | 50.02844545 | -0.8547231946 | 0.2783085753 | 0.000249386295 | 0.001493766691 |
| C17orf97 | 98.26747712 | -0.8548508946 | 0.1773542558 | 1.95E-07 | 2.32E-06 |
| LINC00338 | 105.4054917 | -0.8552182083 | 0.2334358378 | 3.15E-05 | 0.000237043074 |
| RIBC2 | 24.11567847 | -0.8557880392 | 0.4887561184 | 0.006259453632 | 0.02386607046 |
| SGK494 | 29.98642992 | -0.8566630835 | 0.372398446 | 0.002178012634 | 0.009850677431 |
| ZNF221 | 117.9484016 | -0.8581584557 | 0.2085845006 | 4.98E-06 | 4.44E-05 |
| C22orf46 | 217.7884 | -0.8588532176 | 0.1641435158 | 2.24E-08 | 3.17E-07 |
| LOC101929385 | 18.44642322 | -0.8592663739 | 0.4219197298 | 0.003963460044 | 0.01639676482 |
| ARL17A | 287.0633446 | -0.8597299134 | 0.1646803531 | 2.39E-08 | 3.36E-07 |
| AXIN2 | 35.55757384 | -0.8597863134 | 0.3508695576 | 0.00144863073 | 0.006968873325 |
| LIFR-AS1 | 41.03647396 | -0.8598687336 | 0.3040463103 | 0.000523710674 | 0.002861478248 |
| FAM50B | 115.3676098 | -0.8601065064 | 0.1966361158 | 1.58E-06 | 1.55E-05 |
| LOC642852 | 879.153527 | -0.8603013952 | 0.1224550379 | 3.00E-13 | 8.37E-12 |
| NKTR | 1966.371188 | -0.8604360762 | 0.1603587261 | 1.08E-08 | 1.61E-07 |
| NAIF1 | 250.9599976 | -0.8608598801 | 0.1699603331 | 5.37E-08 | 7.11E-07 |
| NIFK-AS1 | 87.55912332 | -0.8621069271 | 0.2097212147 | 5.06E-06 | 4.50E-05 |
| PEX11B | 721.6668772 | -0.863374087 | 0.09988600158 | 8.40E-19 | 3.87E-17 |
| GOLPH3L | 542.774779 | -0.8634986365 | 0.1428786003 | 1.77E-10 | 3.38E-09 |
| TRIM45 | 159.2291316 | -0.864234352 | 0.1619583011 | 1.30E-08 | 1.91E-07 |
| DYNLL1-AS1 | 323.7567796 | -0.8657517972 | 0.1643557785 | 1.85E-08 | 2.66E-07 |
| ERMAP | 481.4931076 | -0.8662361273 | 0.268741912 | 1.45E-04 | 9.27E-04 |
| TRPV1 | 43.65980686 | -0.8684134822 | 0.261964828 | 0.000108734883 | 0.000726639891 |
| STAG3L5P-PVRIG | 251.4395898 | -0.8694779007 | 0.1489771515 | 7.26E-10 | 1.27E-08 |
| KIAA0430 | 900.8169144 | -0.8695770368 | 0.08882403652 | 1.76E-23 | 1.17E-21 |
| ATP6AP1L | 17.30717287 | -0.8696572038 | 0.4672951829 | 0.00536673111 | 0.0210443138 |
| TMEM132B | 20.30548452 | -0.8705500793 | 0.6625106403 | 0.01083909546 | 0.03776814703 |
| KIT | 10.7674583 | -0.87166024 | 0.5506282283 | 0.008551265308 | 0.03093459394 |
| GADD45B | 1190.374023 | -0.8717773221 | 0.09721944808 | 4.23E-20 | 2.12E-18 |
| RAD51-AS1 | 37.5551282 | -0.8721623892 | 0.3301446826 | 0.000861673140 | 0.00442242378 |
| ZBTB48 | 301.6178514 | -0.8736909308 | 0.1524981979 | 1.34E-09 | 2.27E-08 |
| MAMDC4 | 58.03952138 | -0.8738311577 | 0.2416619233 | 3.63E-05 | 0.000269684665 |
| BMS1P4 | 96.78278316 | -0.8749948625 | 0.2236616711 | 1.12E-05 | 9.30E-05 |
| RSBN1L-AS1 | 167.9387435 | -0.8753565075 | 0.2048141339 | 2.37E-06 | 2.24E-05 |

| | baseMean | log2FoldChange | lfcSE | pvalue | padj |
|---|---|---|---|---|---|
| ZNF671 | 209.5060329 | -0.8762535603 | 0.191305745 | 5.78E-07 | 6.27E-06 |
| GOLGA8N | 47.66413283 | -0.876851998 | 0.278416179 | 0.000185371810 | 0.001154812089 |
| RP11-129J12.2 | 13.5146249 | -0.8779936895 | 0.6278168015 | 0.00990162853 | 0.03498465761 |
| AC091729.9 | 75.7084464 | -0.8785349095 | 0.2827173965 | 0.000210161039 | 0.001285046289 |
| ZNF846 | 72.41880712 | -0.8790542849 | 0.259789645 | 8.37E-05 | 0.000573675768 |
| LOC155060 | 28.95095541 | -0.8791509863 | 0.3642475357 | 0.001545686047 | 0.007374566135 |
| LOC100996873 | 10.89780558 | -0.8791600827 | 0.9625775871 | 0.01322315239 | 0.04441225263 |
| APOL3 | 127.2145 | -0.8810771579 | 0.2458006264 | 3.93E-05 | 0.000289331565 |
| E2F2 | 76.35067384 | -0.8811508227 | 0.2610110693 | 8.49E-05 | 0.000580649559 |
| LOC100506124 | 14.6397506 | -0.8817186626 | 0.5409407938 | 0.007590307725 | 0.02795944872 |
| TSC1 | 1396.791016 | -0.881736018 | 0.08099062257 | 1.83E-28 | 1.82E-26 |
| MTHFSD | 304.7221919 | -0.8819254171 | 0.1431307675 | 9.40E-11 | 1.87E-09 |
| MTMR9LP | 35.50676657 | -0.8821996725 | 0.3118726196 | 0.000507483487 | 0.002783028227 |
| LGALS8-AS1 | 9.895967696 | -0.8823252403 | 0.7159131111 | 0.0119471097 | 0.04098629407 |
| NBEAL1 | 986.6040212 | -0.8835870198 | 0.1050804718 | 5.76E-18 | 2.46E-16 |
| ZNF554 | 99.46249816 | -0.8841702346 | 0.2416093118 | 2.97E-05 | 0.00022604457 |
| ANKRD33B | 180.4485746 | -0.8842228867 | 0.1803092794 | 1.20E-07 | 1.50E-06 |
| KCTD1 | 155.8571426 | -0.8853342604 | 0.1464187289 | 2.00E-10 | 3.75E-09 |
| MST1 | 73.71194427 | -0.8856738636 | 0.4042673059 | 0.002524670324 | 0.01117746549 |
| LINC00663 | 20.10466198 | -0.8862525861 | 0.449494706 | 0.004200303246 | 0.01717613706 |
| MAMSTR | 24.06996269 | -0.8868150721 | 0.3619819514 | 0.001439335208 | 0.00692648479 |
| LOC101928762 | 7.500096985 | -0.8879759575 | 0.8315062191 | 0.01339403197 | 0.04487482943 |
| DET1 | 271.0876768 | -0.8883046372 | 0.1287870527 | 7.14E-13 | 1.89E-11 |
| LOC148413 | 362.9817285 | -0.88927399 | 0.1323364248 | 2.38E-12 | 5.88E-11 |
| NNT-AS1 | 128.0346283 | -0.8900070075 | 0.167558406 | 1.42E-08 | 2.08E-07 |
| LOC100190986 | 211.6971925 | -0.8917793554 | 0.2216899491 | 6.83E-06 | 5.93E-05 |
| KRTAP1-5 | 25.52413484 | -0.891869926 | 0.413481889 | 0.002744402083 | 0.01200746477 |
| USP27X-AS1 | 19.99746517 | -0.8923212903 | 0.5942947791 | 0.008655480191 | 0.03124913787 |
| CCDC15 | 224.8943536 | -0.8929739737 | 0.2516386861 | 4.37E-05 | 0.000318533353 |
| ZNF253 | 136.0462824 | -0.8931840288 | 0.2316336695 | 1.34E-05 | 0.000109998339 |
| INSIG2 | 277.7777837 | -0.8932754041 | 0.1660669965 | 9.53E-09 | 1.44E-07 |
| ATF7IP | 1400.50297 | -0.8934707611 | 0.09201325786 | 3.68E-23 | 2.36E-21 |
| LBX2 | 37.21535606 | -0.8935686372 | 0.3000403729 | 3.21E-04 | 0.001862040321 |
| LOC101928548 | 54.20521959 | -0.8937333962 | 0.289876854 | 0.000223531562 | 0.001357177362 |
| FAM86EP | 82.25983183 | -0.8939478667 | 0.1971380984 | 7.16E-07 | 7.58E-06 |
| JRKL | 480.6950533 | -0.8939646805 | 0.1113030828 | 1.30E-16 | 4.84E-15 |
| ZNF117 | 157.7040694 | -0.8958105423 | 0.2372813451 | 1.87E-05 | 1.49E-04 |
| NPIPB4 | 274.4585119 | -0.8962976972 | 0.1633981904 | 5.29E-09 | 8.26E-08 |
| C19orf73 | 12.58246283 | -0.8964155047 | 0.5256844417 | 0.006660692484 | 0.02517670739 |
| FAM13A | 1814.81008 | -0.8967559307 | 0.1923316735 | 3.80E-07 | 4.27E-06 |

| | baseMean | log2FoldChange | lfcSE | pvalue | padj |
|---|---|---|---|---|---|
| LOC100506060 | 45.11193182 | -0.8969223618 | 0.2770285252 | 0.000133521799 | 0.000867367393 |
| NS3BP | 74.02716968 | -0.8970028048 | 0.2296968913 | 1.10E-05 | 9.19E-05 |
| LOC102724455 | 60.30151042 | -0.8975501537 | 0.322263125 | 0.000538312726 | 0.002930507561 |
| LENG9 | 87.59745258 | -0.897784523 | 0.3044213508 | 0.000329949787 | 0.001907814516 |
| TAS2R31 | 14.81373914 | -0.8982843994 | 0.5703391288 | 0.007932064891 | 0.02900338607 |
| CSRP2BP | 586.9418259 | -0.9003025629 | 0.1132465587 | 2.43E-16 | 8.93E-15 |
| GLI4 | 57.51646266 | -0.900389407 | 0.3079936472 | 0.000362780787 | 0.002067933824 |
| SNAI3-AS1 | 156.7354371 | -0.9006274292 | 0.2132513481 | 2.87E-06 | 2.69E-05 |
| ZNF232 | 174.2505247 | -0.9009930301 | 0.1751519403 | 3.34E-08 | 4.57E-07 |
| VILL | 173.0509221 | -0.9011064229 | 0.1955470398 | 5.01E-07 | 5.49E-06 |
| HIF3A | 160.2839936 | -0.9014127747 | 0.2017061302 | 9.53E-07 | 9.76E-06 |
| EME1 | 307.5123571 | -0.9018233608 | 0.3052936531 | 0.000320167402 | 0.001859193625 |
| DOPEY1 | 614.4034317 | -0.9018401191 | 0.1489844657 | 1.80E-10 | 3.42E-09 |
| XAF1 | 177.405013 | -0.9021929447 | 0.3017934292 | 2.88E-04 | 1.70E-03 |
| SUSD2 | 50.5385658 | -0.9036964201 | 0.339010604 | 0.000767616183 | 0.004001763778 |
| FRG1B | 44.90457667 | -0.9038536541 | 0.284176538 | 0.000161635256 | 0.001024515682 |
| LOC100506634 | 87.99971867 | -0.9043321669 | 0.2014470758 | 8.74E-07 | 9.07E-06 |
| LOC101928946 | 11.97505837 | -0.9075130248 | 0.655385905 | 0.009829826418 | 0.03477222275 |
| ZC3H6 | 224.7993729 | -0.9078104602 | 0.1349217872 | 2.24E-12 | 5.57E-11 |
| FBXL8 | 23.96470867 | -0.9092220089 | 0.581261553 | 0.007530193794 | 0.02778225935 |
| FLJ38668 | 26.75973511 | -0.9098239396 | 0.3366612662 | 0.000694926710 | 0.003662571256 |
| ZSCAN26 | 417.2353367 | -0.910459425 | 0.1085983256 | 6.71E-18 | 2.84E-16 |
| ALPK1 | 478.0326711 | -0.9118601523 | 0.1893921276 | 1.76E-07 | 2.11E-06 |
| ZNF607 | 165.8693681 | -0.9119532653 | 0.1349279856 | 1.84E-12 | 4.64E-11 |
| COCH | 8.699925351 | -0.9122686997 | 0.6563484039 | 0.01016749499 | 0.03578814018 |
| ERCC6 | 196.1867731 | -0.9132915106 | 0.1618458389 | 2.08E-09 | 3.46E-08 |
| PTCD2 | 342.5903171 | -0.9140392873 | 0.1351525735 | 1.71E-12 | 4.33E-11 |
| INHBE | 150.2110698 | -0.9145211354 | 0.2837755892 | 0.000132294654 | 0.000861274646 |
| RUNX1-IT1 | 6.962563259 | -0.9149805413 | 0.8737366451 | 0.01333325342 | 0.04470137586 |
| GOLGA6L10 | 17.35817144 | -0.9154181059 | 0.4681148748 | 0.004013867036 | 0.01658226035 |
| CCDC8 | 1216.513341 | -0.9165128732 | 0.1423795586 | 1.50E-11 | 3.34E-10 |
| FAM35DP | 32.15717888 | -0.9172686501 | 0.5329563958 | 0.005761941856 | 0.02233074418 |
| MEG3 | 1607.416633 | -0.9173827197 | 0.1975111712 | 3.90E-07 | 4.35E-06 |
| ZNF517 | 55.55200342 | -0.9174678204 | 0.311544447 | 0.000326702688 | 0.001890507707 |
| NYAP1 | 32.41115541 | -0.9193826408 | 0.4874661277 | 0.004402471777 | 0.01787521197 |
| PPP1R3E | 53.4678988 | -0.9195161554 | 0.2701082722 | 7.18E-05 | 0.000499733661 |
| S100PBP | 938.950777 | -0.9195203847 | 0.0881837159 | 2.44E-26 | 2.04E-24 |
| LOC100272217 | 53.006486 | -0.9199377617 | 0.2972476698 | 0.000203124122 | 0.001248684997 |
| RP11-384K6.6 | 69.0843788 | -0.9204846751 | 0.2819681026 | 0.000116564089 | 0.000773405853 |
| TTC23 | 683.3021898 | -0.9208444845 | 0.1188998796 | 1.23E-15 | 4.28E-14 |

| | baseMean | log2FoldChange | lfcSE | pvalue | padj |
|---|---|---|---|---|---|
| WASH5P | 190.7515422 | -0.9214750985 | 0.2123208064 | 1.62E-06 | 1.59E-05 |
| RAB3A | 15.70669321 | -0.9230031726 | 0.487313582 | 0.004595552054 | 0.01855898163 |
| LOC101928403 | 23.49680768 | -0.9239995229 | 0.3932852928 | 0.001710542761 | 0.008031621309 |
| KBTBD3 | 76.46901028 | -0.9266219001 | 0.1988367591 | 3.87E-07 | 4.34E-06 |
| ZKSCAN1 | 1893.793042 | -0.9278261715 | 0.1030857521 | 2.84E-20 | 1.45E-18 |
| ZNF488 | 22.73530108 | -0.9294249963 | 0.381935092 | 0.001386001464 | 0.006713134611 |
| CRYGS | 12.20627205 | -0.9307888947 | 0.6988357654 | 0.009669515191 | 0.0343111086 |
| ZNF19 | 51.8786589 | -0.9311695761 | 0.25803818 | 3.34E-05 | 0.000249921961 |
| PEG3 | 76.54455824 | -0.9324224445 | 0.3686908492 | 0.001024347538 | 0.005159509637 |
| ZNF445 | 948.328672 | -0.933715502 | 0.1019326575 | 6.69E-21 | 3.58E-19 |
| ARMC7 | 103.3161999 | -0.9349399247 | 0.210193945 | 9.83E-07 | 1.00E-05 |
| LOC100288152 | 69.17122103 | -0.9355073811 | 0.3084131031 | 0.000238773951 | 0.00143814942 |
| PHOSPHO2 | 17.46175982 | -0.9368374732 | 0.4465014996 | 0.002974361075 | 0.01286240547 |
| CCDC87 | 10.02936982 | -0.9377752618 | 0.611323749 | 0.008026315715 | 0.02929733138 |
| ZNF501 | 173.5961162 | -0.9384505607 | 0.1473027141 | 2.29E-11 | 4.99E-10 |
| CAPN3 | 16.20374937 | -0.9390324235 | 0.5151427297 | 0.004952572302 | 0.01973941596 |
| FER1L4 | 37.93505217 | -0.9404825263 | 0.8181093784 | 0.01022019004 | 0.03593963381 |
| SLC16A12 | 69.90731756 | -0.9406656484 | 0.2620926006 | 3.57E-05 | 0.000265953086 |
| ACACB | 85.10493025 | -0.9427431733 | 0.215276022 | 1.33E-06 | 1.33E-05 |
| THNSL1 | 215.2857101 | -0.9434652557 | 0.1964110982 | 1.76E-07 | 2.11E-06 |
| ZNF818P | 133.4948032 | -0.9440923051 | 0.2001224655 | 2.70E-07 | 3.11E-06 |
| ZNF605 | 417.755586 | -0.9443561341 | 0.1501857955 | 3.85E-11 | 8.10E-10 |
| RCOR2 | 259.5964618 | -0.9447276912 | 0.2088366468 | 6.80E-07 | 7.24E-06 |
| LINC00265 | 35.30886986 | -0.9457810931 | 0.2918791095 | 0.000123732881 | 0.000813359323 |
| MAB21L2 | 5818.963662 | -0.945887811 | 0.1241537483 | 3.71E-15 | 1.22E-13 |
| ADM | 3615.308992 | -0.9481275345 | 0.1515158473 | 3.72E-11 | 7.85E-10 |
| MORF4L2-AS1 | 6.836396625 | -0.9487018995 | 0.9364493837 | 0.01256522023 | 0.04263681571 |
| PTGER4 | 487.454165 | -0.9503662498 | 0.1018642675 | 1.32E-21 | 7.37E-20 |
| HSPA1B | 2669.387365 | -0.9506337437 | 0.08783637318 | 3.25E-28 | 3.14E-26 |
| ZNF454 | 45.05856439 | -0.950957202 | 0.3010370135 | 1.59E-04 | 0.0010087357 |
| LOC100499484-C | 48.02607588 | -0.9512401219 | 0.4529058072 | 0.002695848089 | 0.01182769176 |
| ZNF79 | 378.8494432 | -0.9514134926 | 0.1170140535 | 5.14E-17 | 2.02E-15 |
| PUS10 | 139.3139878 | -0.9514399529 | 0.1551682304 | 1.05E-10 | 2.06E-09 |
| ZNF674 | 118.0622171 | -0.9515870131 | 0.1663329068 | 1.27E-09 | 2.16E-08 |
| LOC101060376 | 118.5388548 | -0.9524633415 | 0.2370683081 | 6.31E-06 | 5.52E-05 |
| PRPH2 | 209.4406586 | -0.954154092 | 0.2212718033 | 1.74E-06 | 1.70E-05 |
| ZNF780A | 485.7512849 | -0.9554440349 | 0.1071396861 | 5.89E-20 | 2.87E-18 |
| MMACHC | 233.1287973 | -0.9574432128 | 0.1740705821 | 4.39E-09 | 6.94E-08 |
| TSPAN10 | 10.84274922 | -0.9579352862 | 0.7315013412 | 0.009515367246 | 0.03382863011 |
| ZNF774 | 12.66862688 | -0.9581078608 | 0.6678300547 | 0.008209079951 | 0.02986925498 |

| | baseMean | log2FoldChange | lfcSE | pvalue | padj |
|---|---|---|---|---|---|
| RTKN | 271.1535137 | -0.958496927 | 0.1460994768 | 6.38E-12 | 1.50E-10 |
| AGAP5 | 84.48253641 | -0.9592692933 | 0.2355711399 | 5.00E-06 | 4.45E-05 |
| TCEANC2 | 266.7806455 | -0.9607059352 | 0.1205978197 | 1.95E-16 | 7.22E-15 |
| RBM5 | 3142.62488 | -0.9611795937 | 0.09085894163 | 4.37E-27 | 3.85E-25 |
| TRIM46 | 83.7688665 | -0.9612131387 | 0.2177001759 | 1.11E-06 | 1.12E-05 |
| LINC00886 | 18.46930429 | -0.9615632931 | 0.425595863 | 0.001986202288 | 0.009119262242 |
| TSC22D3 | 3477.315378 | -0.9616564809 | 0.2671845939 | 3.15E-05 | 0.000237252027 |
| FAM46C | 263.4023386 | -0.9618200019 | 0.2939061407 | 0.000103520026 | 0.000695536855 |
| FAM178A | 1652.249163 | -0.9630471677 | 0.07967238792 | 1.52E-34 | 2.15E-32 |
| LOC100130093 | 40.89801812 | -0.964256879 | 0.3508143706 | 0.000541342812 | 0.002944849521 |
| PTGFR | 138.9227592 | -0.966634532 | 0.191835411 | 5.27E-08 | 7.00E-07 |
| THAP6 | 289.6184064 | -0.9700868359 | 0.160105846 | 1.57E-10 | 3.02E-09 |
| HEATR6 | 1126.911646 | -0.9751871901 | 0.08768618431 | 1.19E-29 | 1.28E-27 |
| SH3TC2 | 103.2871507 | -0.9758820405 | 0.1861622273 | 1.85E-08 | 2.66E-07 |
| PGBD2 | 134.3221814 | -0.9783650348 | 0.1938071438 | 4.86E-08 | 6.50E-07 |
| KLHDC1 | 28.48656052 | -0.9795839435 | 0.3388086483 | 0.000364567417 | 0.002074940473 |
| TSC22D1-AS1 | 8.799859881 | -0.9799318055 | 1.114096188 | 0.01140833418 | 0.03941951442 |
| DBP | 19.2128156 | -0.9824509322 | 0.756034076 | 0.008733282368 | 0.03148888339 |
| CCDC41-AS1 | 11.57795046 | -0.9833621101 | 0.5787112667 | 0.005744862504 | 0.02229222252 |
| SEC31B | 66.86370356 | -0.9841358561 | 0.3066604676 | 0.000126356422 | 0.000828046112 |
| ZNF285 | 123.8844128 | -0.9843761609 | 0.167202143 | 4.69E-10 | 8.38E-09 |
| LOC101927249 | 19.17859222 | -0.9861220136 | 0.8390662993 | 0.009437301505 | 0.0336126401 |
| TICRR | 388.5694031 | -0.9866961484 | 0.190647717 | 2.46E-08 | 3.44E-07 |
| DNM3OS | 514.9051259 | -0.9876883129 | 0.1368222689 | 5.74E-14 | 1.72E-12 |
| TYW1B | 58.66389917 | -0.9903117583 | 0.2438258269 | 5.03E-06 | 4.48E-05 |
| CDKN1B | 700.1192871 | -0.9921595582 | 0.2033934433 | 1.15E-07 | 1.43E-06 |
| C21orf58 | 216.2252228 | -0.9926136295 | 0.1535227637 | 1.11E-11 | 2.52E-10 |
| ANKZF1 | 698.5158738 | -0.9969335404 | 0.1446212406 | 6.08E-13 | 1.63E-11 |
| ELAC1 | 169.4708699 | -0.9977079002 | 0.1541042414 | 1.05E-11 | 2.40E-10 |
| PEX11A | 239.4835699 | -0.9981908628 | 0.1620616726 | 8.06E-11 | 1.63E-09 |
| C21orf67 | 29.13110913 | -1.000168936 | 0.4320739766 | 0.001590577923 | 0.007556513943 |
| ZNF573 | 85.46907486 | -1.000174308 | 0.2051622826 | 1.18E-07 | 1.46E-06 |
| LOC257396 | 104.4334991 | -1.002372973 | 0.1751713577 | 1.22E-09 | 2.07E-08 |
| C15orf52 | 267.7425994 | -1.003081777 | 0.1332300168 | 5.69E-15 | 1.86E-13 |
| VIPAS39 | 939.062663 | -1.003463541 | 0.09595897841 | 1.55E-26 | 1.34E-24 |
| LOC102724873 | 93.77615775 | -1.004874362 | 0.1886795245 | 1.12E-08 | 1.67E-07 |
| MIR503HG | 74.71470363 | -1.005211337 | 0.2293312086 | 1.25E-06 | 1.25E-05 |
| ZNF717 | 189.1413814 | -1.006222837 | 0.1397218752 | 6.57E-14 | 1.95E-12 |
| MITF | 886.2692677 | -1.006478493 | 0.1100810161 | 6.90E-21 | 3.67E-19 |
| ZNF302 | 440.5567643 | -1.00660804 | 0.1461522451 | 6.31E-13 | 1.68E-11 |

| | baseMean | log2FoldChange | lfcSE | pvalue | padj |
|---|---|---|---|---|---|
| PRKAR2A-AS1 | 11.19008151 | -1.007545064 | 0.6396813289 | 0.006728182037 | 0.02537383545 |
| ZNF594 | 313.0578415 | -1.009535315 | 0.1059280909 | 1.84E-22 | 1.10E-20 |
| PIK3C2B | 42.54340781 | -1.01103536 | 0.3227638127 | 0.000159298918 | 0.001010567364 |
| C17orf80 | 823.7554629 | -1.011848443 | 0.08208908828 | 7.39E-36 | 1.08E-33 |
| PIF1 | 305.3561732 | -1.014504382 | 0.1634402208 | 5.80E-11 | 1.19E-09 |
| ZNF658 | 252.1910537 | -1.017004255 | 0.1454971147 | 3.06E-13 | 8.52E-12 |
| KLHL3 | 36.60672078 | -1.018379104 | 0.3160662277 | 0.000119925355 | 0.000791826441 |
| IGIP | 32.31890059 | -1.018444085 | 0.3725967721 | 0.000533447643 | 0.002907211555 |
| C7orf13 | 101.1793864 | -1.021636391 | 0.2044796401 | 6.07E-08 | 7.96E-07 |
| OGT | 3383.094508 | -1.022230855 | 0.1470100274 | 3.85E-13 | 1.06E-11 |
| SLC25A42 | 171.324077 | -1.023040808 | 0.2470188009 | 3.36E-06 | 3.09E-05 |
| RCSD1 | 21.48635741 | -1.024271004 | 0.9398913127 | 0.00929969345 | 0.03322052202 |
| BCDIN3D-AS1 | 12.87014093 | -1.02498255 | 0.8458345611 | 0.008938846913 | 0.03215522873 |
| FSTL4 | 47.93643182 | -1.025882629 | 0.4254107497 | 0.001179745589 | 0.005845694816 |
| DPY19L2P2 | 42.14031681 | -1.026492475 | 0.2708203687 | 1.51E-05 | 0.000123344392 |
| HSP90AB4P | 15.20838838 | -1.027154907 | 0.4729840354 | 0.002222922374 | 0.01002334683 |
| MGC16142 | 13.61974963 | -1.029420098 | 0.5093787146 | 0.00303899577 | 0.01308490689 |
| ZEB1-AS1 | 79.18842297 | -1.029717427 | 0.2211288536 | 3.36E-07 | 3.83E-06 |
| DDIT3 | 1193.009243 | -1.031489185 | 0.1537699038 | 2.10E-12 | 5.25E-11 |
| ZNF660 | 122.0108306 | -1.032323754 | 0.2376513189 | 1.36E-06 | 1.35E-05 |
| DBT | 627.5695513 | -1.032471534 | 0.0996189209 | 3.97E-26 | 3.23E-24 |
| DNAJC3-AS1 | 25.05195127 | -1.032857878 | 0.3821647019 | 0.000563718095 | 0.003047639336 |
| FLJ38717 | 65.18877565 | -1.032996831 | 0.2749835652 | 1.60E-05 | 0.000129143991 |
| ZFP3 | 97.20397855 | -1.03651716 | 0.2951885083 | 4.09E-05 | 0.000300001539 |
| LINC01011 | 26.3880149 | -1.040891454 | 0.3576241565 | 0.000309645817 | 0.001805135701 |
| VMAC | 44.71509147 | -1.042853311 | 0.3546479539 | 0.000276590543 | 0.001634189127 |
| ZGLP1 | 19.01513505 | -1.044509091 | 0.4298449986 | 0.001179818183 | 0.005845694816 |
| TADA2A | 583.7025269 | -1.045892998 | 0.1056466593 | 4.50E-24 | 3.24E-22 |
| ANP32A-IT1 | 12.90717327 | -1.046572948 | 0.6371641211 | 0.005484127391 | 0.02140319075 |
| HOMEZ | 357.2232145 | -1.047650347 | 0.116230409 | 2.21E-20 | 1.13E-18 |
| EFNA3 | 34.70287703 | -1.048289328 | 0.5053770668 | 0.00243432629 | 0.01083872133 |
| ZFP90 | 843.3339679 | -1.048820066 | 0.1073431326 | 1.63E-23 | 1.09E-21 |
| LINC00260 | 8.563619538 | -1.052805348 | 0.7149646883 | 0.007252790175 | 0.02695076292 |
| PILRB | 98.84174144 | -1.054183219 | 0.2553709306 | 3.43E-06 | 3.14E-05 |
| PABPC5 | 13.33238283 | -1.056827469 | 0.6896093093 | 0.006338704272 | 0.02411882645 |
| ZNF69 | 123.3862158 | -1.056996098 | 0.1626371397 | 8.79E-12 | 2.03E-10 |
| LOC399900 | 12.67362417 | -1.057968886 | 0.6710765915 | 0.00588796189 | 0.02274672148 |
| PPAPDC2 | 225.6312698 | -1.0611039 | 0.155246613 | 8.68E-13 | 2.28E-11 |
| LINC01000 | 351.9434728 | -1.063118983 | 0.1370972263 | 9.22E-16 | 3.25E-14 |
| ZNF75A | 159.4575457 | -1.065435916 | 0.1694187847 | 3.35E-11 | 7.11E-10 |

| | baseMean | log2FoldChange | lfcSE | pvalue | padj |
|---|---|---|---|---|---|
| YEATS2 | 2538.136359 | -1.06576589 | 0.0743985785 | 1.66E-47 | 3.92E-45 |
| BANP | 369.8369318 | -1.065865805 | 0.1531107635 | 3.42E-13 | 9.49E-12 |
| LOC441124 | 14.49303921 | -1.066959181 | 0.5320793701 | 0.00292417276 | 0.01268220455 |
| FPGT | 373.1716955 | -1.06730191 | 0.1135337497 | 5.75E-22 | 3.35E-20 |
| SH3BP5-AS1 | 71.75265557 | -1.067595604 | 0.2257468565 | 2.23E-07 | 2.61E-06 |
| ZNF702P | 105.4775683 | -1.068668293 | 0.196652758 | 5.75E-09 | 8.90E-08 |
| RUNX1T1 | 79.39235797 | -1.068675825 | 0.2916104188 | 2.22E-05 | 0.000173620847 |
| MDM1 | 251.6437952 | -1.070464427 | 0.1447194081 | 1.43E-14 | 4.54E-13 |
| ZNF396 | 19.55101363 | -1.070836138 | 0.5577368871 | 0.003282161919 | 0.01402643766 |
| SEC16B | 82.56648201 | -1.071102567 | 0.2329243281 | 4.04E-07 | 4.49E-06 |
| HELB | 83.06992236 | -1.071399485 | 0.2061071806 | 2.03E-08 | 2.89E-07 |
| ZSCAN30 | 452.5414501 | -1.073043434 | 0.1260578323 | 1.82E-18 | 8.18E-17 |
| APOLD1 | 136.5200903 | -1.073952168 | 0.1830522857 | 4.48E-10 | 8.03E-09 |
| ANKRD37 | 115.0252654 | -1.074547484 | 0.292901311 | 2.11E-05 | 0.000166008795 |
| EZH1 | 533.6892997 | -1.081397661 | 0.1162515453 | 1.48E-21 | 8.21E-20 |
| ZBTB40 | 1007.607536 | -1.082170411 | 0.09722658668 | 9.33E-30 | 1.02E-27 |
| C1orf213 | 129.4400466 | -1.082748171 | 0.1879420795 | 8.14E-10 | 1.42E-08 |
| DNM1P41 | 9.376753017 | -1.083814122 | 1.121756768 | 0.009415838537 | 0.03357098179 |
| LINC00894 | 29.40188872 | -1.088508643 | 0.353863604 | 0.000175036011 | 0.001100553705 |
| FAM218A | 27.5137353 | -1.089274559 | 0.3470838121 | 0.000140831805 | 0.000906547669 |
| TOB1-AS1 | 9.738091119 | -1.09115301 | 0.6488824109 | 0.005239457341 | 0.02065089498 |
| GATSL3 | 124.0579171 | -1.091944881 | 0.278138182 | 7.62E-06 | 6.58E-05 |
| CTTNBP2 | 32.00921142 | -1.092727542 | 0.3997506419 | 0.000479186411 | 0.002647349342 |
| LOC100287808 | 13.57151463 | -1.093762391 | 0.6545999394 | 0.004916352791 | 0.01961929153 |
| SMYD4 | 410.6877585 | -1.093829279 | 0.1307333645 | 5.97E-18 | 2.53E-16 |
| NAPEPLD | 126.3813889 | -1.095059394 | 0.184687634 | 3.05E-10 | 5.56E-09 |
| SLX4 | 235.7697645 | -1.096766605 | 0.1581255778 | 4.01E-13 | 1.10E-11 |
| FAM124A | 13.8427588 | -1.097075779 | 0.5315702123 | 0.002594174437 | 0.01144434465 |
| LINC01004 | 141.1700765 | -1.097433983 | 0.1819083018 | 1.62E-10 | 3.11E-09 |
| LOC101930375 | 11.19589659 | -1.097911739 | 0.5817338901 | 0.003576893134 | 0.0150654489 |
| HSP90B2P | 12.28490318 | -1.10492407 | 0.5535283555 | 0.002936363095 | 0.01272395522 |
| CCDC121 | 45.01259663 | -1.105046331 | 0.3144969956 | 3.69E-05 | 0.000273568222 |
| PANK1 | 265.4179601 | -1.106109507 | 0.1196656019 | 2.45E-21 | 1.36E-19 |
| AKAP3 | 12.77824174 | -1.108088849 | 0.586495414 | 0.003443401907 | 0.01460249436 |
| MIRLET7BHG | 147.6925584 | -1.110579102 | 0.1711053396 | 8.26E-12 | 1.91E-10 |
| LINC01106 | 14.62007506 | -1.111035243 | 0.5581558195 | 0.002795443943 | 0.01220210052 |
| SCAND2P | 128.7090201 | -1.111300346 | 0.2314128136 | 1.43E-07 | 1.76E-06 |
| GOLGA8A | 887.2106566 | -1.113293875 | 0.2249248478 | 6.76E-08 | 8.78E-07 |
| LINC00087 | 84.11055129 | -1.114808233 | 0.3418450082 | 8.80E-05 | 0.000600113174 |
| NSUN5P1 | 147.2713829 | -1.117319836 | 0.1596018925 | 2.62E-13 | 7.35E-12 |

| | baseMean | log2FoldChange | lfcSE | pvalue | padj |
|---|---|---|---|---|---|
| HOXA6 | 39.49840973 | -1.117465825 | 0.3949739811 | 0.000343289058 | 0.001970485244 |
| FRY | 1337.437472 | -1.118457649 | 0.2389740784 | 2.44E-07 | 2.83E-06 |
| ZNF606 | 202.4081833 | -1.120418693 | 0.1308896746 | 1.18E-18 | 5.36E-17 |
| STK32A | 10.81404057 | -1.121381553 | 0.74249229 | 0.006161222178 | 0.02360031824 |
| ATP1A1OS | 66.66234158 | -1.123631726 | 0.2872880383 | 7.68E-06 | 6.62E-05 |
| USP32P1 | 80.87018282 | -1.123801987 | 0.3642373764 | 0.000152293999 | 0.000971094370 |
| TRMT1L | 600.0444398 | -1.124516863 | 0.1288460615 | 2.50E-19 | 1.19E-17 |
| GNRH1 | 53.79466456 | -1.124739485 | 0.3847349485 | 0.000253914695 | 0.001513772294 |
| GOLGA6L9 | 141.0172532 | -1.125111522 | 0.183959721 | 9.23E-11 | 1.85E-09 |
| LOC101928588 | 91.25539289 | -1.125113323 | 0.2771439972 | 4.17E-06 | 3.77E-05 |
| BCDIN3D | 151.6430843 | -1.127186668 | 0.154214278 | 2.62E-14 | 8.16E-13 |
| ZNF619 | 206.1278672 | -1.127681609 | 0.1683050587 | 2.02E-12 | 5.07E-11 |
| C19orf44 | 159.4098194 | -1.128385205 | 0.1533409418 | 1.78E-14 | 5.60E-13 |
| TTC30B | 144.9827847 | -1.128649896 | 0.1680014083 | 1.78E-12 | 4.51E-11 |
| ZNF630 | 44.05425267 | -1.12971807 | 0.2899995382 | 8.46E-06 | 7.24E-05 |
| LOC100996741 | 40.2494701 | -1.131977484 | 0.3193292146 | 3.36E-05 | 0.000251638815 |
| MED18 | 493.9033721 | -1.137387858 | 0.1155003136 | 6.75E-24 | 4.70E-22 |
| LOC286437 | 50.67215932 | -1.13869068 | 0.2621275444 | 1.29E-06 | 1.29E-05 |
| LOC102724135 | 274.1106735 | -1.140125872 | 0.1186701025 | 7.42E-23 | 4.60E-21 |
| LOC100131564 | 161.0558222 | -1.141518864 | 0.2232805307 | 2.93E-08 | 4.04E-07 |
| LOC102724757 | 21.67193965 | -1.141892766 | 0.4496207063 | 0.000767496421 | 0.004001763778 |
| FAM161B | 144.1072753 | -1.143798933 | 0.1762691447 | 8.18E-12 | 1.90E-10 |
| GPR63 | 443.562056 | -1.145112628 | 0.2771841924 | 3.02E-06 | 2.81E-05 |
| ZNF33A | 521.546257 | -1.147116029 | 0.1300126121 | 1.09E-19 | 5.26E-18 |
| MSS51 | 45.72969808 | -1.151188834 | 0.3119484814 | 1.84E-05 | 0.000147112147 |
| ZNF713 | 30.56196593 | -1.153522527 | 0.3044263085 | 1.33E-05 | 0.000109595527 |
| KIAA1614 | 239.1605762 | -1.154459108 | 0.222443242 | 1.89E-08 | 2.71E-07 |
| EXPH5 | 30.00362292 | -1.155824148 | 0.3626924614 | 0.000112530055 | 0.000749311266 |
| CDC42EP3 | 4129.975742 | -1.155836989 | 0.1554374579 | 9.04E-15 | 2.91E-13 |
| ZSCAN23 | 95.883962 | -1.15897233 | 0.1924427747 | 1.69E-10 | 3.23E-09 |
| LOC100129518 | 17.76874294 | -1.159355453 | 0.5642882481 | 0.002242748556 | 0.01009745487 |
| MGC16275 | 23.40546389 | -1.161414379 | 0.4074319743 | 0.000316619385 | 0.00184074425 |
| RECQL5 | 454.2908845 | -1.162921205 | 0.1250724829 | 1.35E-21 | 7.50E-20 |
| RASL11A | 200.0160601 | -1.163029328 | 0.4585919547 | 0.000704945137 | 0.003714058087 |
| CCDC180 | 23.07104437 | -1.166008753 | 0.4193784518 | 0.000385822457 | 0.00218422455 |
| CRIPAK | 11.25322948 | -1.166223639 | 0.7221674787 | 0.004962380789 | 0.01976060112 |
| KCNMB3 | 39.89771399 | -1.166385273 | 0.3803934998 | 0.000163016558 | 0.001032391974 |
| CTAGE8 | 7.376235851 | -1.166711576 | 1.689446046 | 0.00977666297 | 0.03462879305 |
| FLJ44342 | 230.9039393 | -1.169125042 | 0.2330821318 | 4.58E-08 | 6.15E-07 |
| OSBPL7 | 129.9175705 | -1.171165235 | 0.2977222523 | 6.79E-06 | 5.90E-05 |

| | baseMean | log2FoldChange | lfcSE | pvalue | padj |
|---|---|---|---|---|---|
| ZNF608 | 986.1044802 | -1.17145241 | 0.1863141306 | 2.89E-11 | 6.23E-10 |
| PNPLA3 | 229.8348326 | -1.173406749 | 0.1624177687 | 4.73E-14 | 1.44E-12 |
| LDLRAD4 | 8.227223888 | -1.173539434 | 1.428841895 | 0.009047363405 | 0.03247497438 |
| TSSK6 | 9.653664768 | -1.173782246 | 0.7647743226 | 0.005642097568 | 0.02195640112 |
| KIAA1407 | 71.94426348 | -1.174103805 | 0.3033805585 | 8.98E-06 | 7.64E-05 |
| PTCH1 | 246.4896117 | -1.175330875 | 0.2289628649 | 2.49E-08 | 3.47E-07 |
| PLGLB2 | 13.32719517 | -1.180882497 | 0.64313368 | 0.003445276537 | 0.01460249436 |
| BTBD8 | 34.8657016 | -1.182087265 | 0.3447248429 | 4.81E-05 | 0.000346367066 |
| ZNF879 | 104.6746506 | -1.183396607 | 0.1821048758 | 7.60E-12 | 1.77E-10 |
| ZNF341 | 81.55203487 | -1.184378445 | 0.1944175702 | 1.02E-10 | 2.02E-09 |
| ATF7IP2 | 32.3659515 | -1.184871343 | 0.3552213727 | 6.83E-05 | 0.000477150406 |
| PAQR8 | 206.727621 | -1.188015126 | 0.18912744 | 3.10E-11 | 6.64E-10 |
| ZNF214 | 106.2867821 | -1.189068578 | 0.1820519703 | 6.16E-12 | 1.46E-10 |
| SPIN3 | 174.0834565 | -1.191683401 | 0.1473042089 | 5.79E-17 | 2.25E-15 |
| LOC102723566 | 10.80013872 | -1.192327858 | 0.6178214801 | 0.002972885539 | 0.01285975967 |
| ZNF780B | 324.8286864 | -1.203369476 | 0.1604464574 | 5.86E-15 | 1.91E-13 |
| ZNF391 | 41.7535187 | -1.207628595 | 0.2979216311 | 4.29E-06 | 3.87E-05 |
| LOC100128398 | 23.45283398 | -1.210459716 | 0.4112635066 | 0.000236491149 | 0.00142613071 |
| CBX2 | 136.0726667 | -1.211579865 | 0.2867878696 | 1.85E-06 | 1.79E-05 |
| KRCC1 | 724.4025607 | -1.213791045 | 0.1179319612 | 6.92E-26 | 5.54E-24 |
| PEX12 | 198.9709746 | -1.21718949 | 0.1465838 | 9.00E-18 | 3.80E-16 |
| LOC102724005 | 15.20544189 | -1.220187293 | 0.6886862514 | 0.003552491675 | 0.01498880964 |
| LINC00847 | 163.8995841 | -1.22307746 | 0.1700499634 | 5.84E-14 | 1.74E-12 |
| SH3D21 | 153.0814216 | -1.223240665 | 0.2379816192 | 2.31E-08 | 3.25E-07 |
| DNAH11 | 66.22054739 | -1.227683143 | 0.2331715408 | 1.24E-08 | 1.83E-07 |
| PPP1R3C | 2234.480412 | -1.229058791 | 0.4654051376 | 0.000491143596 | 0.002706379352 |
| GOLGA8O | 31.39633757 | -1.231145038 | 0.7683733287 | 0.004282238907 | 0.01746323065 |
| DNAJC28 | 15.92290101 | -1.232479907 | 0.5564614412 | 0.001521824655 | 0.007278653159 |
| ZNF692 | 241.3502134 | -1.233086784 | 0.1760318974 | 2.13E-13 | 6.02E-12 |
| LOC101929046 | 6.203874221 | -1.234130458 | 0.8848597815 | 0.00698138038 | 0.02611702826 |
| LOC100506022 | 36.616344 | -1.238207845 | 0.3136196368 | 6.47E-06 | 5.65E-05 |
| ZNF547 | 62.31214977 | -1.250919988 | 0.2551816962 | 7.75E-08 | 9.94E-07 |
| SLC16A13 | 19.63398673 | -1.255407717 | 0.4727150498 | 0.000498945097 | 0.002743276796 |
| FLJ37201 | 17.07481395 | -1.255536022 | 0.4632393042 | 0.000458304159 | 0.002544254521 |
| LOC102724281 | 13.09358531 | -1.256207646 | 0.6466126729 | 0.002696164002 | 0.01182769176 |
| KBTBD7 | 222.830532 | -1.260299129 | 0.1538980937 | 2.28E-17 | 9.20E-16 |
| KCNQ1OT1 | 1718.388906 | -1.260412106 | 0.2302905157 | 3.46E-09 | 5.55E-08 |
| LOC730668 | 6.219372643 | -1.260537235 | 1.157457045 | 0.008585389135 | 0.03103371178 |
| LOC101928524 | 12.7413909 | -1.260698055 | 0.5949682385 | 0.001943869048 | 0.008963228943 |
| RAD52 | 195.6774801 | -1.261344746 | 0.1447838746 | 2.53E-19 | 1.20E-17 |

| | baseMean | log2FoldChange | lfcSE | pvalue | padj |
|---|---|---|---|---|---|
| KLRAP1 | 16.50309929 | -1.262166116 | 0.5187001835 | 0.000925255562 | 0.004704962453 |
| FAM72D | 111.1254414 | -1.265074756 | 0.3068920262 | 2.77E-06 | 2.60E-05 |
| C17orf103 | 105.2659633 | -1.265869963 | 0.4409652471 | 0.000253972945 | 0.001513772294 |
| SNAI2 | 2213.00318 | -1.266407174 | 0.2566515939 | 6.25E-08 | 8.17E-07 |
| PARD6B | 19.6707226 | -1.272987274 | 0.584695365 | 0.0015805069 | 0.007515863056 |
| LOC100133331 | 20.00829475 | -1.275969156 | 0.4185194323 | 0.000162351338 | 0.001028616631 |
| MDFI | 28.97786181 | -1.279420889 | 0.4327641127 | 0.000204849490 | 0.001255145703 |
| ZNF181 | 219.7734237 | -1.279721339 | 0.1694227418 | 3.61E-15 | 1.20E-13 |
| C5orf63 | 86.82914381 | -1.279916296 | 0.2100090934 | 9.56E-11 | 1.90E-09 |
| LOC100134822 | 22.01183755 | -1.281342067 | 0.4286584846 | 0.000186477733 | 0.001160730343 |
| HCFC2 | 718.1718574 | -1.282424915 | 0.08663849343 | 1.27E-50 | 3.65E-48 |
| LOC102723820 | 25.61633772 | -1.282500634 | 0.402580822 | 0.000102652719 | 0.000690792676 |
| KCTD7 | 400.6949716 | -1.286779321 | 0.1326570494 | 2.63E-23 | 1.71E-21 |
| LOC101926980 | 31.62273521 | -1.286782891 | 0.3341720004 | 8.97E-06 | 7.64E-05 |
| SH3GL1P2 | 8.934207253 | -1.289887205 | 0.747639098 | 0.003949649438 | 0.01635781098 |
| RAD51L3-RFFL | 38.83474009 | -1.293888991 | 0.4117306545 | 0.000109634583 | 0.000732323603 |
| STARD5 | 103.6324483 | -1.294145629 | 0.2256766803 | 8.17E-10 | 1.42E-08 |
| LOC101060181 | 19.38654696 | -1.298322809 | 0.4283808835 | 0.000167288599 | 0.001056822047 |
| LOC101928592 | 13.97977879 | -1.299170173 | 0.6621472804 | 0.002376674315 | 0.01061058587 |
| LINC00506 | 7.060343283 | -1.299200841 | 0.9331508062 | 0.006369028919 | 0.02421564647 |
| C10orf10 | 101.2711306 | -1.299862982 | 0.4001082324 | 7.45E-05 | 0.000516344553 |
| ZNF154 | 61.18230658 | -1.300866644 | 0.3526542035 | 1.59E-05 | 0.000129143991 |
| RP11-466F5.8 | 20.84765995 | -1.301541344 | 0.5290294185 | 0.000834302972 | 0.004298248082 |
| LOC100287896 | 28.86262753 | -1.301697391 | 0.4100342604 | 0.000102973404 | 0.000692176530 |
| THAP8 | 136.0612408 | -1.30448719 | 0.2382150087 | 3.43E-09 | 5.50E-08 |
| DNASE1 | 97.68025866 | -1.30633109 | 0.2148235496 | 9.65E-11 | 1.91E-09 |
| FAM13C | 46.19586854 | -1.308972185 | 0.5941513735 | 0.001369006314 | 0.006648119704 |
| ZNF658B | 68.76912623 | -1.312934828 | 0.3199987991 | 2.95E-06 | 2.76E-05 |
| CA9 | 44.46019162 | -1.313174859 | 0.4617616182 | 0.000270578381 | 0.001603120381 |
| IP6K3 | 271.369766 | -1.315942932 | 0.2065554678 | 1.50E-11 | 3.34E-10 |
| ZNF571 | 42.51820112 | -1.317558966 | 0.302982428 | 1.05E-06 | 1.06E-05 |
| TNFRSF25 | 17.15565922 | -1.317998526 | 0.4482381051 | 0.000221338108 | 0.001345103304 |
| C1orf220 | 21.7027715 | -1.321308391 | 0.6654358197 | 0.002168524062 | 0.009810742861 |
| IL16 | 105.1906106 | -1.321491967 | 0.5865706476 | 0.001181293169 | 0.0058495198 |
| ZFP2 | 18.55718308 | -1.322058216 | 0.4711179469 | 3.29E-04 | 1.90E-03 |
| ZNF772 | 293.4060172 | -1.329362091 | 0.1477914026 | 1.94E-20 | 1.00E-18 |
| PPFIBP2 | 58.66677648 | -1.33163481 | 0.284193758 | 2.18E-07 | 2.55E-06 |
| LOC729468 | 9.844630737 | -1.334988866 | 0.8052299572 | 0.004093982774 | 0.01684779147 |
| ZNF429 | 59.730661 | -1.33533184 | 0.2236074933 | 1.99E-10 | 3.74E-09 |
| ZFP62 | 575.560294 | -1.337795934 | 0.1214689768 | 2.71E-29 | 2.86E-27 |

| | baseMean | log2FoldChange | lfcSE | pvalue | padj |
|---|---|---|---|---|---|
| LOC102724624 | 11.83313371 | -1.340274849 | 0.5843403269 | 0.001235630536 | 0.006081753075 |
| STOX2 | 46.49884465 | -1.341224452 | 0.6491633263 | 0.001757683957 | 0.008214110622 |
| FIGN | 214.1854604 | -1.341753282 | 0.138442736 | 2.69E-23 | 1.74E-21 |
| GPR20 | 99.96876928 | -1.343364255 | 0.4706818438 | 0.000252509227 | 0.001508066543 |
| DMGDH | 11.07397587 | -1.347730729 | 0.672664346 | 0.002257283177 | 0.01015368254 |
| ZNF471 | 144.5497073 | -1.349354637 | 0.2308892539 | 3.93E-10 | 7.07E-09 |
| LOC440300 | 101.4759868 | -1.351837689 | 0.2136109021 | 1.91E-11 | 4.19E-10 |
| HEXIM2 | 99.4322987 | -1.35448915 | 0.2854575185 | 1.50E-07 | 1.83E-06 |
| LOC100289230 | 52.20676919 | -1.354928091 | 0.2660914944 | 2.80E-08 | 3.88E-07 |
| LOC100499484 | 39.20688534 | -1.358337172 | 0.3318375964 | 3.08E-06 | 2.86E-05 |
| TRHDE-AS1 | 252.6659141 | -1.361768704 | 0.2120311917 | 1.03E-11 | 2.35E-10 |
| PRKCQ | 20.12136303 | -1.362725724 | 0.4099698582 | 6.30E-05 | 0.000443689268 |
| RP11-611D20.2 | 18.2151091 | -1.364194547 | 0.4847356344 | 0.000303967085 | 0.00177829703 |
| ZNF681 | 64.34327568 | -1.365710006 | 0.2691634876 | 3.01E-08 | 4.14E-07 |
| UBA6-AS1 | 198.2268111 | -1.367636114 | 0.1314280377 | 1.96E-26 | 1.66E-24 |
| LOC102724335 | 38.8177019 | -1.368228276 | 0.2860138584 | 1.34E-07 | 1.65E-06 |
| SENP8 | 125.537 | -1.37212113 | 0.1812522926 | 3.15E-15 | 1.05E-13 |
| PRSS30P | 23.76871028 | -1.372292698 | 0.4788138164 | 0.000255155161 | 0.001520210165 |
| PIK3R3 | 526.1880552 | -1.376524044 | 0.1268257945 | 1.49E-28 | 1.49E-26 |
| ZNF589 | 205.9128935 | -1.37773051 | 0.1993735064 | 3.65E-13 | 1.01E-11 |
| GSDMB | 94.4128178 | -1.381920586 | 0.2091801501 | 3.21E-12 | 7.80E-11 |
| CBX8 | 195.1780502 | -1.389687165 | 0.1944146928 | 6.51E-14 | 1.94E-12 |
| NUDT18 | 173.9152405 | -1.391541109 | 0.1730822532 | 7.45E-17 | 2.85E-15 |
| LOC101929140 | 10.57498837 | -1.392102497 | 0.9316933134 | 0.004751539974 | 0.01908500454 |
| ZNF763 | 21.91008272 | -1.40128682 | 0.5202726139 | 0.000405334540 | 0.002285129105 |
| LOC100996662 | 11.72509858 | -1.41091574 | 0.59673154 | 0.000994787203 | 0.005024215288 |
| LINC00865 | 55.27847144 | -1.41154934 | 0.2902260016 | 8.38E-08 | 1.07E-06 |
| JRK | 739.1668985 | -1.411584345 | 0.1295060518 | 9.29E-29 | 9.47E-27 |
| LOC100506127 | 38.36798628 | -1.415933158 | 0.3261463462 | 9.98E-07 | 1.02E-05 |
| LOC101928188 | 32.10685296 | -1.425273939 | 0.4271510455 | 5.21E-05 | 0.000373892579 |
| NBPF1 | 476.3133064 | -1.425812567 | 0.123386108 | 5.19E-32 | 6.33E-30 |
| CLK2 | 532.2975573 | -1.433799252 | 0.1027873534 | 2.42E-45 | 5.07E-43 |
| PPP1R3G | 30.68062972 | -1.434945309 | 0.5006805327 | 0.000236083771 | 0.001424250923 |
| AHSA2 | 500.2889125 | -1.436300514 | 0.1845959785 | 5.43E-16 | 1.94E-14 |
| CCDC89 | 98.69904649 | -1.443698666 | 0.205507098 | 1.64E-13 | 4.70E-12 |
| LOC101929130 | 23.46267324 | -1.446556646 | 0.4671096905 | 0.000120285579 | 0.000792798577 |
| LOC100379224 | 42.35263101 | -1.451271497 | 0.2769740236 | 1.22E-08 | 1.81E-07 |
| LOC102724250 | 755.4007806 | -1.451947799 | 0.08397780344 | 4.77E-68 | 2.09E-65 |
| RIMBP3B | 14.15661156 | -1.452702099 | 0.5855362681 | 0.000706333444 | 0.003717426177 |
| CCDC183-AS1 | 9.689794276 | -1.455942591 | 0.6471142523 | 0.001331503466 | 0.006485036016 |

| | baseMean | log2FoldChange | lfcSE | pvalue | padj |
|---|---|---|---|---|---|
| KRBA2 | 207.8548543 | -1.457765759 | 0.1348520838 | 2.63E-28 | 2.55E-26 |
| MINOS1P1 | 11.29294475 | -1.461281643 | 0.6592963005 | 0.001317835823 | 0.006426877684 |
| ZSCAN20 | 266.8509841 | -1.467350298 | 0.1296208415 | 7.99E-31 | 9.29E-29 |
| EFNA1 | 14.91422172 | -1.467401115 | 0.9217237997 | 0.003626665457 | 0.0152363497 |
| ARHGEF39 | 123.7918018 | -1.473320729 | 0.1924637924 | 1.39E-15 | 4.78E-14 |
| HSPA1L | 41.16350648 | -1.484175711 | 0.2970396276 | 4.45E-08 | 6.00E-07 |
| RP11-983P16.4 | 18.55694933 | -1.484454957 | 0.4231413792 | 3.18E-05 | 0.000239212655 |
| USP51 | 39.86063689 | -1.485754498 | 0.32908937 | 4.62E-07 | 5.09E-06 |
| KIAA1652 | 16.05293794 | -1.487230397 | 0.5024279757 | 0.000187779187 | 0.00116590672 |
| ZNF862 | 262.426634 | -1.491281508 | 0.194024699 | 1.10E-15 | 3.85E-14 |
| ZNF793 | 78.98814673 | -1.491461957 | 0.257683881 | 4.96E-10 | 8.82E-09 |
| GOLGA8R | 28.02116732 | -1.494670204 | 0.3597185935 | 2.31E-06 | 2.19E-05 |
| CCDC36 | 95.87017438 | -1.49698161 | 0.193965361 | 9.13E-16 | 3.23E-14 |
| ZNF33B | 362.496769 | -1.49954052 | 0.1704983877 | 1.12E-19 | 5.36E-18 |
| SKIDA1 | 22.70644679 | -1.501083943 | 0.4207548662 | 2.38E-05 | 0.000184887794 |
| FLJ30403 | 7.152954712 | -1.501532246 | 0.7202595375 | 0.001990529045 | 0.00913593926 |
| PFKFB4 | 877.1056315 | -1.502236376 | 0.2532585141 | 2.07E-10 | 3.88E-09 |
| NPFF | 8.116208467 | -1.502366576 | 0.6645666554 | 0.001353742725 | 0.006580435987 |
| HERC2P2 | 1120.063699 | -1.503153561 | 0.1602048635 | 4.73E-22 | 2.77E-20 |
| LOC101929824 | 6.170419307 | -1.504942127 | 1.124896759 | 0.006010954022 | 0.02311996768 |
| APLN | 80.0349853 | -1.519540167 | 0.4302396527 | 2.35E-05 | 0.000182832154 |
| ZNF662 | 61.73950283 | -1.521907258 | 0.2315906201 | 3.97E-12 | 9.58E-11 |
| LOC100128288 | 20.93068042 | -1.525467699 | 0.5020055296 | 0.000143643082 | 0.000922251767 |
| ZNF135 | 168.4136787 | -1.528569583 | 0.1673318587 | 4.89E-21 | 2.68E-19 |
| CROCCP3 | 69.64020668 | -1.530765127 | 0.2422005373 | 1.90E-11 | 4.18E-10 |
| N4BP2L2-IT2 | 39.59762904 | -1.533438991 | 0.3868472565 | 4.65E-06 | 4.17E-05 |
| GIN1 | 169.9469229 | -1.535164067 | 0.1409996272 | 1.06E-28 | 1.08E-26 |
| WDR52 | 50.45350411 | -1.536304097 | 0.3714539524 | 2.28E-06 | 2.16E-05 |
| C1orf51 | 82.98969501 | -1.538991566 | 0.2328034363 | 2.77E-12 | 6.81E-11 |
| RP5-1057J7.6 | 83.08080096 | -1.539567401 | 0.2602541944 | 2.39E-10 | 4.45E-09 |
| FAM227A | 53.6785738 | -1.540287602 | 0.2883959518 | 6.42E-09 | 9.89E-08 |
| ZNF311 | 60.44440132 | -1.547020929 | 0.2388726901 | 7.42E-12 | 1.73E-10 |
| GDPGP1 | 14.60239872 | -1.549395541 | 0.5514215592 | 0.000279897693 | 0.001651762489 |
| LOC100129917 | 135.8880611 | -1.551812643 | 0.1914442758 | 4.00E-17 | 1.58E-15 |
| IFIT2 | 62.08247665 | -1.55871378 | 0.2653664064 | 3.06E-10 | 5.58E-09 |
| TRIM66 | 101.6422062 | -1.563708876 | 0.193877817 | 5.81E-17 | 2.25E-15 |
| HCG8 | 7.374067262 | -1.564318542 | 0.7634804599 | 0.002063484337 | 0.009421410088 |
| BATF2 | 13.28968147 | -1.5664299 | 0.7406810585 | 0.001494141288 | 0.007160048163 |
| GDF5 | 390.7192717 | -1.566846119 | 0.3098319636 | 2.69E-08 | 3.74E-07 |
| WDR5B | 257.8562489 | -1.569665712 | 0.1324315128 | 1.55E-33 | 2.06E-31 |

| | baseMean | log2FoldChange | lfcSE | pvalue | padj |
|---|---|---|---|---|---|
| SLX4IP | 74.99088007 | -1.569946771 | 0.2201382327 | 7.57E-14 | 2.24E-12 |
| ZNF248 | 283.1046759 | -1.580528713 | 0.1181682539 | 6.82E-42 | 1.22E-39 |
| LOC100507520 | 125.1282409 | -1.583465161 | 0.1975596906 | 7.41E-17 | 2.84E-15 |
| C6orf223 | 7.371486092 | -1.585554324 | 0.875150902 | 0.002926533518 | 0.01268505314 |
| LOC101928042 | 10.52164894 | -1.587276813 | 0.585816263 | 0.000411738703 | 0.002318599678 |
| ATP6V0E2-AS1 | 40.0854236 | -1.594205826 | 0.3046408461 | 1.13E-08 | 1.69E-07 |
| GRAMD2 | 20.78643008 | -1.598790713 | 0.4241249652 | 1.04E-05 | 8.70E-05 |
| KCNJ2 | 91.51396475 | -1.607978778 | 0.3041099367 | 7.90E-09 | 1.21E-07 |
| PNMAL2 | 43.49703583 | -1.611782256 | 0.4981999337 | 6.55E-05 | 0.000458800877 |
| LOC728743 | 76.95155772 | -1.613967102 | 0.3266567106 | 5.18E-08 | 6.88E-07 |
| MIR210HG | 262.5711142 | -1.615854123 | 0.2137115776 | 2.69E-15 | 9.06E-14 |
| ARHGEF37 | 33.32742418 | -1.625280058 | 0.5519775414 | 0.000164181204 | 0.001038883959 |
| HSD17B7P2 | 47.13761536 | -1.629822184 | 0.2692883185 | 1.05E-10 | 2.07E-09 |
| ZNF397 | 262.538568 | -1.629926704 | 0.1615419096 | 4.32E-25 | 3.30E-23 |
| BAMBI | 438.9973127 | -1.630129394 | 0.3483016244 | 1.72E-07 | 2.07E-06 |
| MTHFR | 540.5936186 | -1.652068585 | 0.2291135481 | 3.73E-14 | 1.15E-12 |
| ZSCAN31 | 42.8154604 | -1.652450553 | 0.335595239 | 5.41E-08 | 7.16E-07 |
| BMS1P2 | 35.76988934 | -1.658516966 | 0.4206444663 | 4.87E-06 | 4.36E-05 |
| RASL11B | 70.20947552 | -1.663127149 | 0.3367535934 | 5.03E-08 | 6.71E-07 |
| RAPGEF4 | 6.660270116 | -1.663754265 | 0.8198761824 | 0.002076771105 | 0.009470457881 |
| NIM1K | 23.13912496 | -1.669088215 | 0.5988851523 | 0.000256161593 | 0.001525595986 |
| ZNF610 | 57.45182003 | -1.671046957 | 0.2466823269 | 9.32E-13 | 2.43E-11 |
| PKDCC | 2685.925745 | -1.671113402 | 0.1960396144 | 1.03E-18 | 4.73E-17 |
| SMAD6 | 73.37284077 | -1.67179049 | 0.3391124267 | 5.17E-08 | 6.88E-07 |
| ZNF425 | 73.95181426 | -1.677769331 | 0.3256148904 | 1.69E-08 | 2.45E-07 |
| TUBA3FP | 17.5070232 | -1.684365424 | 0.4920458577 | 3.79E-05 | 0.000279750642 |
| LOC101928714 | 13.11534458 | -1.702992094 | 0.5942453173 | 0.00023589255 | 0.00142367417 |
| LURAP1L | 351.3996053 | -1.709568947 | 0.2776986836 | 4.77E-11 | 9.87E-10 |
| LOC101927610 | 17.28211031 | -1.711362559 | 0.4480979209 | 9.24E-06 | 7.83E-05 |
| LOC100272216 | 24.58551039 | -1.711983945 | 0.384077511 | 5.47E-07 | 5.96E-06 |
| SYTL5 | 10.61159277 | -1.718995997 | 0.5761384001 | 0.000178757877 | 0.001119228782 |
| GPR146 | 24.22128389 | -1.722969626 | 0.8337506474 | 0.001372367325 | 0.006662268374 |
| RP5-1180C10.2 | 44.48203799 | -1.734465059 | 0.2996322937 | 4.79E-10 | 8.55E-09 |
| NKX2-3 | 870.2279393 | -1.735171268 | 0.2692820056 | 7.29E-12 | 1.71E-10 |
| C17orf59 | 183.5912916 | -1.735328399 | 0.2149113389 | 4.41E-17 | 1.74E-15 |
| ZNF132 | 174.6994958 | -1.74280467 | 0.1635152697 | 1.18E-27 | 1.10E-25 |
| LOC101930129 | 8.074638479 | -1.75404278 | 0.9126304805 | 0.002193514988 | 0.00991175862 |
| LOC100507131 | 23.2607583 | -1.770067072 | 0.3878586351 | 3.46E-07 | 3.93E-06 |
| ZKSCAN7 | 135.5594124 | -1.776867796 | 0.1845751681 | 4.17E-23 | 2.64E-21 |
| IGDCC3 | 6.583300699 | -1.777889883 | 1.145750033 | 0.003917402524 | 0.01624232976 |

| | baseMean | log2FoldChange | lfcSE | pvalue | padj |
|---|---|---|---|---|---|
| ZNF334 | 90.00693255 | -1.779139668 | 0.2864789294 | 3.52E-11 | 7.45E-10 |
| CSMD3 | 61.50630634 | -1.798655206 | 0.3765306182 | 1.06E-07 | 1.33E-06 |
| LOC101929005 | 10.61763505 | -1.802705705 | 0.604958157 | 0.000171076495 | 0.001077933957 |
| TCEANC | 35.57472055 | -1.805447055 | 0.3869971673 | 2.05E-07 | 2.42E-06 |
| HOXD4 | 111.5424597 | -1.809518386 | 0.2046269317 | 6.77E-20 | 3.29E-18 |
| SH3GL1P1 | 13.12150645 | -1.867486232 | 0.5107563771 | 1.76E-05 | 1.41E-04 |
| TMEM79 | 79.38737582 | -1.876296366 | 0.2539882794 | 9.53E-15 | 3.06E-13 |
| ZNF177 | 31.37480053 | -1.887415227 | 0.4679589182 | 2.98E-06 | 2.78E-05 |
| SPACA6P-AS | 9.912154115 | -1.898275031 | 0.8027902275 | 0.000802937850 | 0.004165484895 |
| ZKSCAN3 | 166.6413778 | -1.92667051 | 0.1619943135 | 9.23E-34 | 1.24E-31 |
| ZNF439 | 56.2392475 | -1.940521955 | 0.2717524285 | 6.96E-14 | 2.06E-12 |
| ZNF300P1 | 77.11859557 | -1.945126052 | 0.2188116724 | 4.49E-20 | 2.23E-18 |
| FAM157A | 9.707825314 | -1.946939454 | 0.7632480009 | 0.000538303539 | 0.002930507561 |
| TMEM100 | 152.155126 | -1.963756801 | 0.3497011208 | 1.13E-09 | 1.93E-08 |
| ZNF572 | 54.80246248 | -1.967384962 | 0.358787415 | 2.63E-09 | 4.31E-08 |
| ZSWIM3 | 144.5158247 | -1.967623825 | 0.190604797 | 3.59E-26 | 2.94E-24 |
| LOC100506571 | 21.66026015 | -1.97880721 | 0.5725610923 | 2.72E-05 | 0.000208589223 |
| LOC102723469 | 8.043097873 | -1.985764712 | 0.6950822365 | 0.000270462727 | 0.001603073072 |
| PPFIA4 | 119.4019948 | -1.995551239 | 0.4260213769 | 1.50E-07 | 1.82E-06 |
| C5orf54 | 180.323534 | -2.001772701 | 0.1599735218 | 3.91E-37 | 5.82E-35 |
| OTUB2 | 47.497715 | -2.002494886 | 0.3694570983 | 3.47E-09 | 5.55E-08 |
| KCNIP2 | 10.72425286 | -2.031297952 | 0.6648145986 | 0.000124771951 | 0.000818395613 |
| ZNF815P | 13.34469523 | -2.045279141 | 0.5636797456 | 1.76E-05 | 0.000141389389 |
| NPY4R | 13.22707624 | -2.046612346 | 0.6341015966 | 6.54E-05 | 0.000458534537 |
| MIR143HG | 61.37805173 | -2.056929424 | 0.5619359445 | 1.17E-05 | 9.68E-05 |
| LOC554206 | 9.853587829 | -2.064314033 | 0.7741518092 | 0.000368042147 | 0.002089923544 |
| RP11-82L2.1 | 10.09860634 | -2.138034534 | 0.6745052504 | 9.73E-05 | 0.000657919810 |
| ZNF546 | 102.4956458 | -2.150303128 | 0.1867254904 | 8.40E-32 | 1.02E-29 |
| HCG27 | 14.40275762 | -2.151881314 | 0.5911278726 | 1.56E-05 | 0.000126689796 |
| SLFNL1 | 9.120116774 | -2.15389563 | 0.7380416084 | 0.000196389228 | 0.001213294282 |
| RHOU | 71.22347631 | -2.1602617 | 0.5004386319 | 8.08E-07 | 8.46E-06 |
| SIRT4 | 21.42225544 | -2.163634615 | 0.4238193102 | 2.23E-08 | 3.15E-07 |
| LINC00654 | 59.70941438 | -2.182151363 | 0.281227816 | 5.77E-16 | 2.06E-14 |
| CSRNP3 | 32.67973631 | -2.184234344 | 0.3839422597 | 8.47E-10 | 1.47E-08 |
| RNF138P1 | 36.46177639 | -2.195461387 | 0.451922555 | 6.50E-08 | 8.46E-07 |
| DDIT4 | 6472.195958 | -2.197847562 | 0.2268516041 | 2.17E-23 | 1.42E-21 |
| BMP4 | 580.1599025 | -2.220252412 | 0.5210298471 | 9.40E-07 | 9.67E-06 |
| EPHB3 | 36.60790586 | -2.240375417 | 0.7232726877 | 8.04E-05 | 0.000554040622 |
| LOC102724356 | 11.58519522 | -2.248685268 | 0.5892998218 | 9.20E-06 | 7.81E-05 |
| RIMBP3 | 15.1433969 | -2.298121158 | 0.5527493153 | 2.10E-06 | 2.00E-05 |

| | baseMean | log2FoldChange | lfcSE | pvalue | padj |
|---|---|---|---|---|---|
| C1orf172 | 10.40956082 | -2.368322295 | 0.6101807467 | 8.16E-06 | 7.00E-05 |
| SOWAHD | 8.47188925 | -2.37047982 | 1.017194636 | 0.000809510114 | 0.0041951953 |
| FAXDC2 | 91.58111492 | -2.387901029 | 0.5244863056 | 2.59E-07 | 2.99E-06 |
| PPT2-EGFL8 | 25.01309986 | -2.399451671 | 0.5121679419 | 1.82E-07 | 2.17E-06 |
| PSMD5-AS1 | 101.1104162 | -2.423751391 | 0.2398057492 | 3.45E-25 | 2.65E-23 |
| LOC102723772 | 26.15995985 | -2.550906198 | 0.4057895717 | 2.53E-11 | 5.49E-10 |
| ADAMTS15 | 32.45358743 | -2.631359621 | 0.8907864679 | 0.000120260153 | 0.000792798577 |
| LOC100996273 | 17.69530652 | -2.651259075 | 0.5652065762 | 1.72E-07 | 2.07E-06 |
| FAM46B | 264.3812489 | -2.695833479 | 0.2039290466 | 3.49E-41 | 6.12E-39 |
| ZNF449 | 256.6678129 | -2.748739356 | 0.1443731826 | 4.69E-82 | 2.79E-79 |
| LINC00638 | 14.1682653 | -2.762312822 | 0.6117371432 | 4.84E-07 | 5.32E-06 |
| LOC100506472 | 10.23022894 | -2.834031947 | 0.7104676509 | 5.77E-06 | 5.08E-05 |
| LOC729652 | 6.505753937 | -2.930292063 | 0.9648409071 | 0.000187486210 | 0.001165545799 |
| ZFP14 | 109.0368235 | -2.958549548 | 0.2771089792 | 1.02E-27 | 9.55E-26 |
| ZNF233 | 28.32533197 | -3.124757208 | 0.495417625 | 1.93E-11 | 4.24E-10 |
| LOC101928230 | 7.203354225 | -3.463320487 | 1.139673943 | 0.000201270522 | 0.001239336974 |
| LOC100128882 | 16.37145284 | -5.553312122 | 1.010890185 | 2.32E-08 | 3.27E-07 |
